# Insect REPAT proteins mediate cross-kingdom communication in microbe–insect–plant interactions

**DOI:** 10.64898/2026.08.03.742523

**Authors:** Rosa Esmeralda Becerra, Adrià Mengual-Martí, Ada Frattini, Victor Flors, Chaymaa Riahi, Meritxell Pérez-Hedo, Alberto Urbaneja, Salvador Herrero, Cristina Crava

## Abstract

Plant–herbivore interactions are embedded within complex ecological networks in which microorganisms associated with either partner can reshape interactions across trophic levels. Yet how microbial infections of herbivorous insects alter the molecular signals they deliver to plants remains largely unexplored. Here, we identify REPAT proteins as key molecular mediators linking microbial infection in caterpillars to plant defense responses. Comparative genomic analyses revealed that REPATs comprise an insect protein family that has expanded in Lepidoptera, with a conserved β-REPAT lineage and rapidly diversifying α- and γ-REPAT lineages, the latter encompassing most REPATs previously implicated in plant interactions. Proteomic analyses of caterpillar oral secretions revealed that viruses with contrasting infection strategies exerted opposite effects on REPAT abundance. Infection with the lethal baculovirus Spodoptera exigua multiple nucleopolyhedrovirus (SeMNPV) increased the abundance of multiple REPAT proteins, whereas infection with the persistent, covert iflavirus Spodoptera exigua iflavirus 1 (SeIV1) consistently reduced REPAT abundance across dietary conditions. These changes were associated with reciprocal effects on plant defense: herbivory by SeMNPV-infected larvae attenuated jasmonate-associated responses in tomato, whereas SeIV1-infected larvae enhanced defense activation in pepper. Together, our findings reveal that viral infection of the herbivore can propagate across trophic levels and reshape plant responses to herbivory by altering the repertoire of effectors delivered during feeding, uncovering a mechanism through which insect–microbe interactions can influence plant defense.

## Introduction

In caterpillars, the insect gut serves as the primary interface with the environment, acting as the main route of entry for both nutrients and a diverse array of entomopathogens (Wagner and Hoyt, 2022). This role is particularly critical for polyphagous Lepidoptera, such as Noctuidae spp., which feed on a wide variety of host plants and are consequently exposed to a vast spectrum of nutritional components, plant defensive allelochemicals, and diverse microbial threats (Hilliou et al., 2021; Kergoat et al., 2021; Kshatriya and Gershenzon, 2024). To cope with these changing environments, these insects deploy plastic physiological responses to maintain gut homeostasis by balancing nutrient uptake, epithelial renewal, and immune defenses.

Caterpillar adaptation to different host plants is mediated by the secretion of elicitors and effectors that modulate plant defense responses, as well as by flexible regulation of digestive enzymes and detoxification pathways, enhancing nutrient accessibility (Heckel, 2018; Jones et al., 2022; Kshatriya and Gershenzon, 2024; Wagner and Hoyt, 2022). During herbivory, plants perceive tissue damage and insect-derived cues, triggering the activation of plant defense mechanisms aimed at deterring attackers (Schuman and Baldwin, 2016). These interactions are largely mediated by components of caterpillar oral secretion (OS), a complex mixture of saliva and gut regurgitant deposited onto wounded plant tissues during insect feeding (Peiffer and Felton, 2009; Rivera-Vega et al., 2017). OS components can act either as elicitors that activate defense responses or as effectors that suppress or attenuate defense signaling (Jones et al., 2022).

In recent years, a few members of a group of orally secreted proteins known as REPAT have been identified as key effectors in caterpillar OS (Chen et al., 2019, 2023; García-Marin et al., 2025; Wang et al., 2024; Zhang et al., 2024). For example, *Helicoverpa armigera* (Lepidoptera: Noctuidae) effector HARP1 enter plant cells through feeding wounds and stabilize JAZ repressors to suppress jasmonate (JA) signaling (Chen et al., 2019; Yan et al., 2023), whereas another REPAT protein, HAS1, targets bHLH transcription factors and cooperates with HARP1 to attenuate downstream defense gene activation (Chen et al., 2023). In *Spodoptera exigua* (Lepidoptera: Noctuidae), REPAT38 acts similarly to HARP1 (Chen et al., 2019) whereas VLPR4 in *Spodoptera frugiperda* (Lepidoptera: Noctuidae) has a role similar to that of HAS1 (Zhang et al., 2024). In a distantly related species, the tomato leaf miner, *Tuta absoluta* (Lepidoptera: Gelechiidae), a REPAT protein downregulates plant herbivory response by suppressing multiple defense hormones, including JA, ethylene, and abscisic acid (X. Wang et al., 2024). Collectively, these findings across evolutionarily divergent species point to a conserved effector function for orally secreted REPATs in plant–insect interactions.

Intriguingly, the REPAT function extends beyond plant-insect interactions. These proteins were originally identified as strongly induced genes in *S. exigua* in response to pathogen challenge (REPAT: REsponse to PAThogens) (Hernández-Martínez et al., 2010; Herrero et al., 2007; Navarro-Cerrillo et al., 2013a, 2012). REPATs are small secreted proteins (<20 kDa), predominantly expressed in the larval midgut, and several undergo N-linked glycosylation (Herrero et al., 2007; Navarro-Cerrillo et al., 2013a). Many contain a Multiprotein Bridge Factor 2 (MBF2) domain (Pfam PF15868), associated with transcriptional co-activation via TFIIA interaction (Li et al., 1997; Liu et al., 2000).

At the transcriptional level, *repat* genes respond to diverse midgut challenges, including bacterial pathogens (Hernández-Martínez et al., 2010; Herrero et al., 2007; Navarro-Cerrillo et al., 2013a, 2012; Tanaka et al., 2010; Y. Wang et al., 2024; C.-Y. Zhou et al., 2016) baculoviruses (Chen et al., 2020; Herrero et al., 2007; Hrithik et al., 2021; Jakubowska et al., 2013), entomopathogenic fungi (Hrithik et al., 2021; Y. Wang et al., 2024), and epithelial damage caused by pore-forming toxins such as Vip3A and Cry1C (Ayra-Pardo et al., 2019; Bel et al., 2013; Ren et al., 2020), suggesting that midgut epithelial disruption and cellular stress, rather than pathogen presence alone, are the main triggers of this response, possibly mediated through eicosanoid signaling (Hrithik et al., 2021). Functional studies support a role in immunity, including reduced baculovirus virulence upon *repat1* overexpression and increased susceptibility to bacterial infection following RNAi knockdown in *S. exigua* and *S. frugiperda* (Herrero et al., 2007; Hrithik et al., 2021; Y. Wang et al., 2024), potentially involving Toll pathway regulation (Y. Wang et al., 2024). Altogether, these findings position REPATs as a major component of the insect response to gut-associated biotic stress.

We hypothesize that REPAT proteins integrate immune and ecological functions across different biological scales. Within the larval gut, they contribute to responses to biotic stress, while a subset is secreted in OS where they act as effectors that manipulate host plant signaling. To explore this dual functionality at the interface of insect immunity and plant–insect interactions, we first investigate the evolution of the *repat* gene family across insects, revealing expansion and diversification within Lepidoptera, particularly in generalist species. We next assess their presence in OS across two virus*-S. exigua*-host plant systems and correlate their abundance in OS with plant defense responses. Altogether, our study reveals REPAT proteins as key molecular mediators of microbe-insect-plant interactions, showing that microbial infection can reshape plant defense by altering the repertoire of insect effectors delivered during feeding.

## Results

### Genome-wide discovery of *repat* genes across insects

REPAT proteins identified to date fall into two structural groups that differ in the presence or absence of the MBF2 domain (NCBI database accessed October 2025) (**Figure 1A,1B**). Despite this distinction, AlphaFold predicts that REPAT proteins adopt a conserved fold characterized by a core of seven β-sheets (**Figure 1C**), even though their sequences have diverged substantially. To identify novel repat genes across insects, we constructed a Hidden Markov Model (HMM) based on an alignment of the conserved *S. exigua* MBF2. We combined it with iterative BLAST searches against genomic and protein datasets from 20 invertebrate species. This approach identified 430 bona fide *repat* genes from 16 insect species (**Figure 1D, Supplementary Dataset 1**), including 39 of the 42 previously described *S. exigua repat* transcripts (Navarro-Cerrillo et al., 2013a) and the 10 *Bombyx mori mbf2* genes previously reported (C.-Y. Zhou et al., 2016), extending their annotation to 81 and 30 genes in *S. exigua* and *B. mori*, respectively.

**Figure 1.**
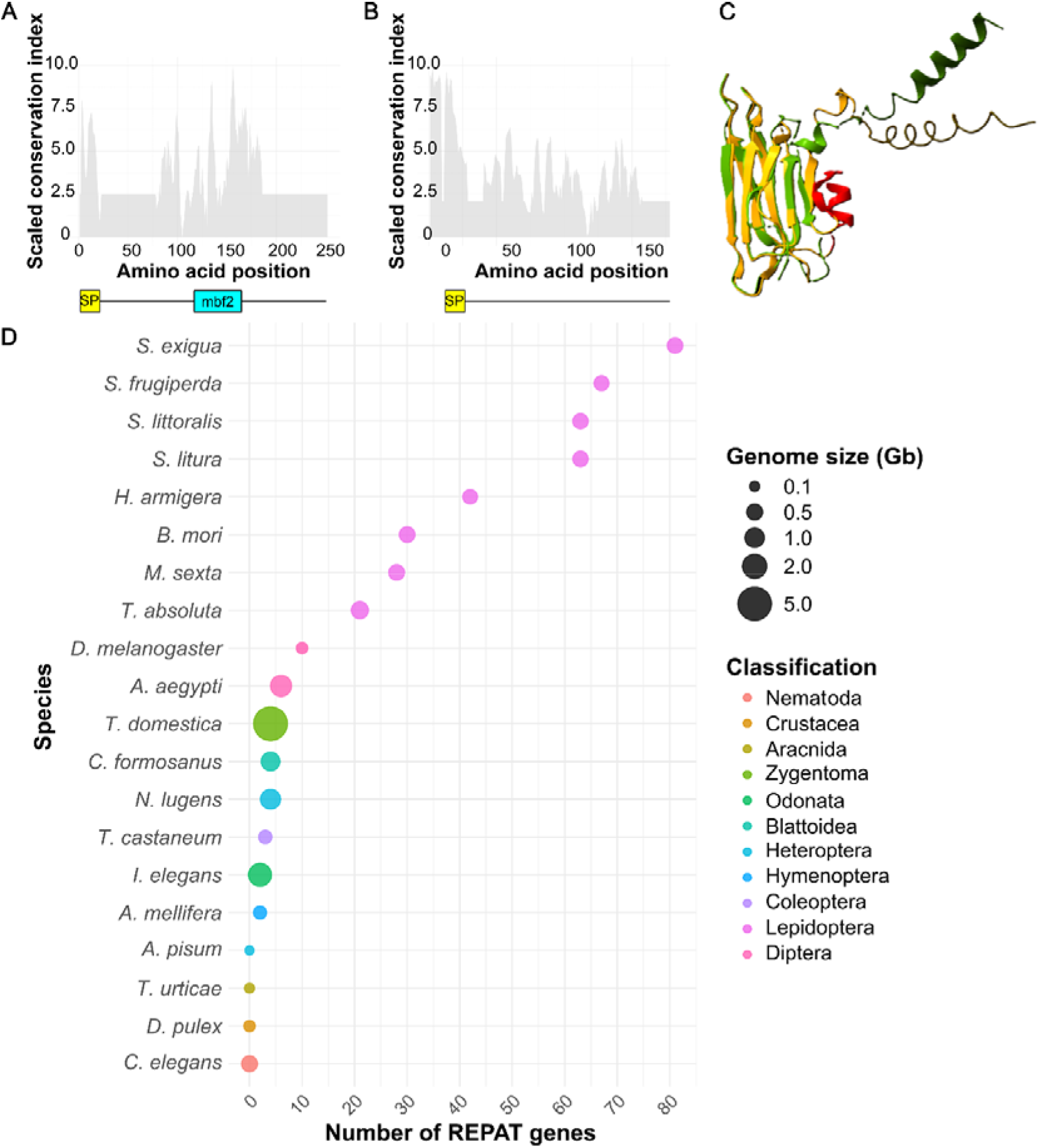
Phylogenomic survey of *repat* genes. (**A**) Scaled conservation index derived from the alignment of *S. exigua* REPAT proteins containing the MBF2 domain available in NCBI (accessed in October 2025). SP, signal peptide. (**B**) Scaled conservation index derived from the alignment of *S. exigua* REPAT proteins lacking the MBF2 domain deposited in NCBI (accessed in October 2025). Abbreviation: SP, signal peptide. (**C**) Structural superposition of REPAT1 (accession no. ACI90726.1; designated Sexi-αREPAT12 in this study) (yellow), which lacks the MBF2 domain, and REPAT33 (accession no. AFH57153.1; designated Sexi-γREPAT1 in this study) (green), which contains the MBF2 domain, highlighting the conserved seven β-sheet core shared by both proteins despite only 24% amino acid sequence identity. The α-helix highlighted in red is a structural feature specific to α-REPAT proteins. (**D**) Bubble plot showing the number of *repat* genes identified across 20 genomes as a function of genome size, illustrating the lack of correlation between these two variables. Bubble color indicates taxonomic classification.

*repat* genes were broadly distributed across insects, including basal lineages such as Zygentoma, but were not detected in the non-insect ecdysozoans included in our survey (e.g., *Daphnia pulex*, *Caenorhabditis elegans*, and *Tetranychus urticae*), suggesting that the gene family likely originated within the insect lineage after its divergence from other ecdysozoan groups. The size of the *repat* gene family varied markedly among species and did not correlate with genome size. Lepidopteran genomes showed a pronounced expansion (21-81 genes), with the highest numbers observed in *Spodoptera* spp. (**Figure 1D**). Notably, MBF2-lacking *repat* sequences were found exclusively within the *Spodoptera* genus, which also exhibited the largest expansion of this gene family (63–81 genes) (**Figure 2A**). These results identify *repat* as an insect-specific gene family that underwent an extensive expansion during lepidopteran evolution.

**Figure 2.**
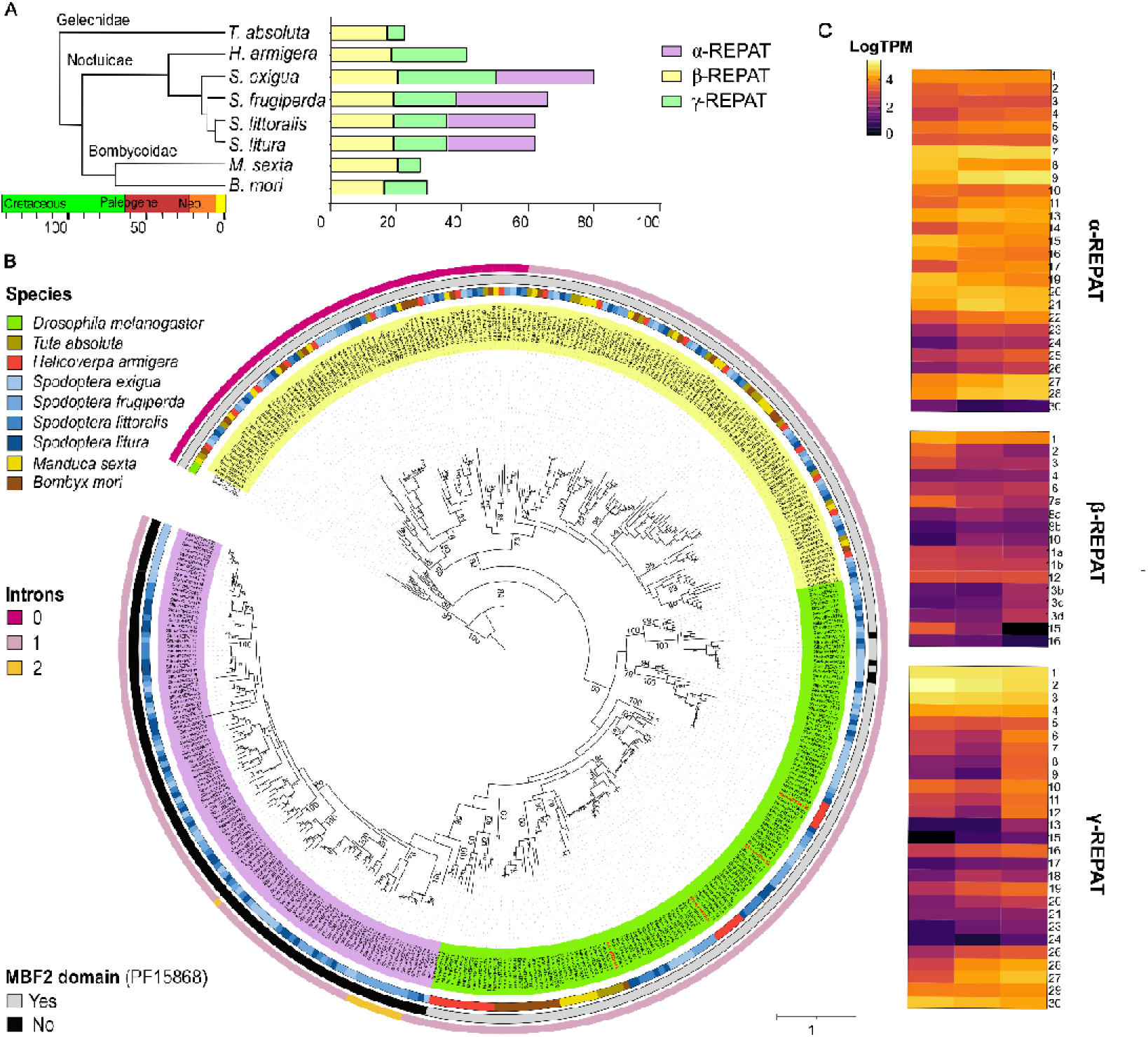
Evolutionary origins and diversification of lepidopteran *repat* genes. (**A**) Stacked-bar plot showing the number of *α-repat*, *β-repat* and *γ-repat* genes identified in the indicated lepidopteran species. A time-calibrated phylogeny adapted from Kergoat et al. (2021), illustrating the relationships among the analyzed species, is shown on the left. (**B**) Phylogenetic relationships of the lepidopteran *repat* coding sequences. The sequences were aligned with PRANK and the tree was built with RAxML under the GTRGAMMA model of substitution with 500 bootstrap replicates, using two *D. melanogaster* sequences as outgroup. The scale bar indicates the expected number of substitutions per site. Background color indicates *α-*, *β-* and *γ-repat* groups, and external circles indicate the species of origin, the number of introns, and the presence or absence of the MBF2 domain. Sequences highlighted in red correspond to REPAT proteins previously characterized as plant defense effectors: Harm-γREPAT9 = HARP1 (Chen et al., 2019), Harm-γREPAT21 = HAS1 (Chen et al., 2023), Sfru-γREPAT12 = VRLP4 (Zhang et al., 2024), and Tuta-γREPAT1 = REPAT38 (X. Wang et al., 2024) (**C**) Heatmap showing log-transformed TPM values for the 70 (out of 81) *S. exigua repat* genes expressed in the gut of fifth-instar larvae (≥1 TPM). Columns represent the mean of three biological replicates (n = 3).

### Evolution and expansion of lepidopteran *repat* genes

To investigate the evolutionary diversification of *repat* genes across Lepidoptera, we reconstructed the phylogenetic relationships among all identified sequences and integrated information on MBF2 domain content, chromosomal localization, and gene structure (**Supplementary dataset 1**). This analysis expanded the classification previously proposed by Navarro-Cerrillo et al. (2013) and identified three major REPAT groups: *α-*, *β-*, and *γ-repat* (**Figure 2**). Whereas *β-* and *γ-repats* are broadly distributed across Lepidoptera and contain the MBF2 domain, α*-repats* are restricted to the genus *Spodoptera* and lack a canonical MBF2 domain. (**Figure 2A**).

*β-repat* genes form a well-supported monophyletic clade displaying clear one-to-one orthology among lepidopteran species (**Figure 2B, Supplementary Figure 1A**). Gene number is remarkably conserved (17–21 copies), and most genes retain conserved chromosomal synteny despite being distributed across multiple chromosomes (**Supplementary Figure 1B**). Many intronless *β-repats* occur in tandem arrays, suggesting that local duplication events contributed to their expansion (**Supplementary Figure 1D**). Reconciliation of gene and species trees revealed moderate gene turnover, with 15 inferred gene gains and 9 gene losses across Lepidoptera (**Supplementary Figure 1C**).

In contrast, *α-* and *γ-repat* genes are predominantly organized as tandem clusters on a single chromosome (**Supplementary Figure 2**). *α-repats* form a well-supported monophyletic group restricted to *Spodoptera*, where they have undergone substantial expansion (27–30 copies). *γ-repats* are present across all analyzed species but exhibit considerable variation in copy number, ranging from five genes in the specialist *P. absoluta* to thirty genes in the generalist *S. exigua*. Phylogenetic relationships within both groups remain poorly resolved, consistent with extensive lineage-specific duplication events (**Figure 2B**). Notably, all REPAT proteins previously characterized as plant defense effectors, including HARP1, HAS1, VRLP4, and REPAT38, belong to the γ-REPAT group (**Figure 2B**).

### Large dynamic range of *repat* expression in the caterpillar gut

To characterize *repat* expression in the caterpillar gut, we analyzed gut transcriptomes from three biological replicates of fifth instar *S. exigua* caterpillars. *repat* expression (≥1 TPM) was detected for 70 of the 81 annotated genes, indicating that most family members are transcriptionally active in the gut.

Expression levels varied markedly among *repat* groups (**Figure 2C**). Twenty-seven of the 30 *α-repat* genes were expressed exhibiting high expression levels, with a median of 8,491 TPM (IQR = 24,009). Similarly, most *β-repats* were detected (17 of 21 genes), but transcript abundances were generally low (median = 462 TPM; IQR = 1,076). Within this group, however, *β-repat1* dominated transcriptional output, reaching 15,031 TPM, 27-fold higher than the average of the remaining members. *γ-repats* showed intermediate but highly variable expression levels, with 16 of 30 genes expressed and a median of 2,129 TPM (IQR = 13,893). The most abundant *γ-repat* transcripts were *γ-repat1-5*, which are orthologs of the *H. armigera* gene encoding the effector HARP1 (*harm-γ-repat9*). Overall, *repat* transcripts were broadly expressed in the *S. exigua* gut, but their abundance differed markedly among evolutionary groups, with *α-repats* showing constitutively high expression, *β-repats* generally expressed at low levels, and *γ-repats* exhibiting substantial heterogeneity in transcript abundance.

### Baculovirus infection increases oral REPAT abundance and decreases tomato anti-herbivory defenses

Previous transcriptomic studies showed that infection by the DNA virus Spodoptera exigua multinucleopolyhedrous virus (SeMNPV) induces the expression of multiple *repat* genes in the larval gut of *S. exigua* (Chen et al., 2020; Jakubowska et al., 2013). We therefore asked whether changes in the REPAT composition of caterpillar OS accompany this transcriptional response.

REPAT proteins detected in the OS predominantly belonged to the γ-REPAT group, followed by α-REPATs, whereas only a single β-REPAT was identified (**Figure 3A**). SeMNPV infection increased the abundance of several REPAT proteins in the OS (**Figure 3B**). Among γ-REPATs, the strongest increases were observed for γ-REPAT1–5, which are orthologous to the *H. armigera* effector HARP1 (Harm-γREPAT9) (**Figure 3A**, **Figure 2B, Supplementary Figure 3**). Likewise, members of the γ-REPAT10–13 clade, orthologous to the effector HAS1 (Harm-γREPAT21), also increased in abundance, although statistical significance was only detected for γ-REPAT11 (**Supplementary figure 4**). In addition, four α-REPAT proteins showed increased abundance following SeMNPV infection (**Figure 3A**).

**Figure 3.**
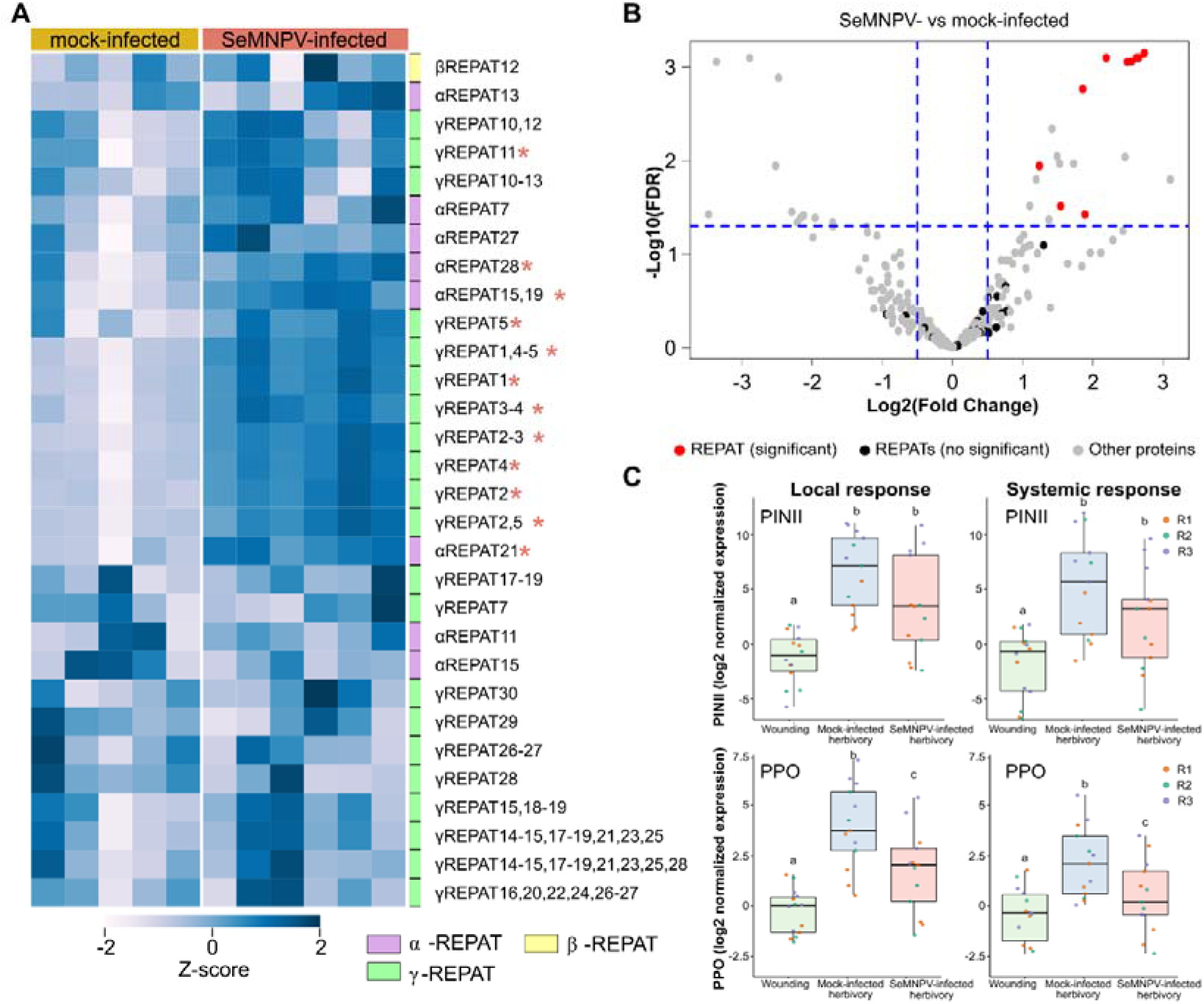
Baculovirus infection increases REPAT abundance in caterpillar oral secretion and decrease tomato defense responses. (**A**) Heatmap showing Z-score-normalized LFQ (label-free quantification) values of REPAT proteins detected in the oral secretions (OS) of mock- and Spodoptera exigua multiple nucleopolyhedrovirus (SeMNPV)-infected fifth-instar larvae. Asterisks indicate proteins showing significant differences in abundance (t test, P < 0.05). (**B**) Volcano plot of differential protein abundance in caterpillar OS. Red dots indicate REPAT proteins significantly enriched in SeMNPV-infected larvae, black dots indicate non-significant REPAT proteins, and gray dots represent all other proteins. (**C**) Relative expression of the jasmonate-responsive genes *PINII* (*proteinase inhibitor II*) and *PPO* (*polyphenol oxidase*) in tomato leaflets subjected to mechanical wounding or 3 h of herbivory by mock- or SeMNPV-infected larvae. Expression was measured 24 h after treatment in locally damaged leaflets (left panels) and adjacent systemic leaflets (right panels). Data represents three independent experiments (R1-R3), with five biological replicate per treatment. Differences were analyzed by ANCOVA followed by Tukey’s HSD test. Different letters indicate significant differences (*P* < 0.05).

Because several infection-induced REPATs belong to clades containing proteins previously implicated in the suppression of plant defenses, we next examined markers of tomato defense responses following herbivory by mock- or SeMNPV-infected larvae, which consumed similar amounts of leaf tissue independently of infection status (**Supplementary Figure 5**). At end of the three-hour feeding period, expression of the JA biosynthetic gene *OPR3* was reduced in leaves exposed to SeMNPV-infected caterpillars (**Supplementary Figure 6**). Similarly, twenty-four hours after feeding, the JA-responsive gene *PPO* showed reduced expression compared with leaves attacked by mock-infected larvae, whereas the other JA-responsive gene, *PINII,* did not show a statistically significant difference, although an overall trend toward reduced expression was observed (**Figure 3C**). These changes in locally damaged leaves were mirrored by similar changes in systemic adjacent leaves. In contrast, markers of salicylic acid signaling remained unaffected (**Supplementary Figure 6**). These findings establish a parallel between larval infection status -associated with an enrichment of orally secreted REPAT proteins - and the modulation of JA-associated responses in tomato.

### SeIV1 infection elicits opposite changes in REPAT abundance and pepper anti-herbivory defenses

To ask whether the association between REPAT abundance and plant responses extends beyond the SeMNPV-tomato system, we examined an independent pathosystem involving pepper and the Spodoptera exigua iflavirus 1 (SeIV1), an RNA virus that establishes persistent, asymptomatic infections with minor effects on host fitness (Millán-Leiva et al., 2012; Virto et al., 2014). Previous findings showed that the feeding substrate influences the REPAT composition of larval OS (García-Marin et al., 2025). We therefore profiled the OS of SeIV1-free and SeIV1-infected *S. exigua* larvae maintained on either artificial diet or pepper leaves to determine whether SeIV1 infection was associated with changes in REPAT composition under both feeding regimes. SeIV1 infection altered REPAT composition under both dietary conditions, although these changes were smaller than those associated with diet (**Figure 4A**, **Figure 4B**). In both dietary regimens, SeIV1 infection was associated with a general reduction in REPAT abundance. **(Figure 4C, D)**. Specifically, several α-REPATs (α-REPAT5, α-REPAT10, α-REPAT29, and α-REPAT30) and γ-REPAT27 decreased in abundance in SeIV1-infected larvae. Among REPATs orthologous to the previously characterized plant effectors HARP1 and HAS1, only HARP1 orthologs showed significant reductions, particularly γ-REPAT2 and γ-REPAT5 in pepper-fed larvae (**Supplementary Figures 3 and 4**).

**Figure 4.**
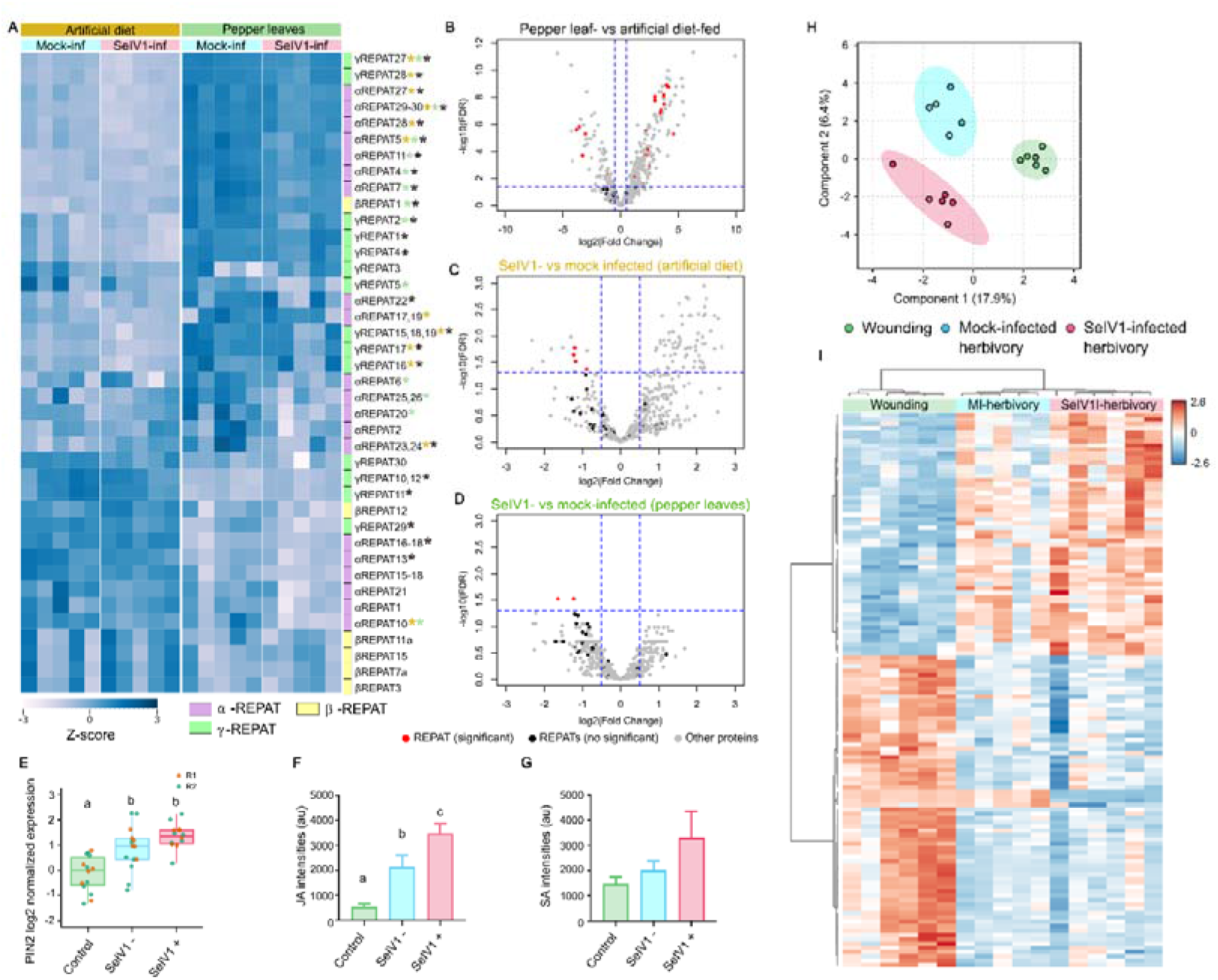
SeIV1 infection decreases REPAT abundance in caterpillar oral secretion and enhances pepper defense responses. (**A**) Heatmap showing Z-score-normalized LFQ (label-free quantification) values of REPAT proteins detected in the oral secretions (OS) of mock- and Spodoptera exigua iflavirus 1 (SeIV1)-infected fifth-instar larvae reared on artificial diet or pepper leaves. Asterisks indicate significant differences in protein abundance (t test, P < 0.05) associated with diet (black), SeIV1 infection in artificial diet (yellow), or SeIV1 infection in pepper-fed larvae (green). (**B–D**) Volcano plots showing differential protein abundance in caterpillar OS according to diet (**B**), SeIV1 infection in larvae reared on artificial diet (**C**), and SeIV1 infection in larvae fed on pepper leaves (**D**). Red dots indicate significantly differentially abundant REPAT proteins, black dots indicate non-significant REPAT proteins, and gray dots represent all other proteins detected in the OS. (**E**) Relative expression of a jasmonate (JA)-responsive defense gene (*PINII*) in pepper leaves subjected to mechanical wounding or herbivory by free- or SeIV1-infected larvae. (**F**) Relative abundance of JA in un-attacked pepper leaves or subjected to herbivory by free- or SeIV1-infected neonate larvae. (**G**) Relative abundance of salicylic acid in un-attacked pepper leaves or subjected to herbivory by free- or SeIV1-infected neonate larvae (**H**) Principal component analysis (PCA) of metabolomic profiles from pepper leaves subjected to mechanical wounding or herbivory by free- or SeIV1-infected larvae. (I) Heatmap showing the relative abundance of metabolites differentially accumulated in wounded leaves and in leaves damaged by free- or SeIV1-infected caterpillars.

We next determined whether the oral-secretion phenotype observed in fifth-instar larvae was accompanied, at the pathosystem level, by altered responses of pepper plants following herbivory by vertically infected neonates. Herbivory increased JA levels approximately 1.5-fold in plants fed upon by SeIV1-infected larvae, whereas salicylic acid levels remained unchanged (**Figure 4F**). In contrast, the JA-responsive PIN2 gene JA-responsive did not reached significantly higher levels in plants attacked by SeIV1-infected larvae (**Figure 4E**). However, metabolomic analyses revealed that herbivory substantially altered the overall metabolic profile of pepper plants, with additional differences observed between plants attacked by SeIV1-free and SeIV1-infected larvae (**Figure 4G, H**). Together, these results reveal that SeMNPV and SeIV1 infections produce opposite effects on REPAT abundance in OS, which parallel to opposite changes in JA-dependent plant defenses. This reciprocal pattern strongly supports REPAT proteins as mediators, linking insect infection status to host plant responses.

## Discussion

Plant–herbivore interactions are embedded within a complex ecological network in which microorganisms associated with either partner can influence the outcome of their interaction (Martínez-Medina et al., 2024; Shao et al., 2024). Here we show that viral infection of the herbivore rewires this molecular dialogue by altering the abundance of REPAT proteins in caterpillar OS, leading to opposite changes in plant defense responses. These findings identify REPAT proteins as candidate mediators connecting insect infection status with host defense signaling and reveal a previously unrecognized mechanism underlying microbe–insect–plant interactions.

Our evolutionary analyses suggest that REPAT proteins most likely originated within insects and subsequently diversified extensively in Lepidoptera, giving rise to two evolutionary and functionally distinct components. *β-repats* display clear one-to-one orthology, indicating strong evolutionary constraint and likely conservation of the ancestral function associated with the MBF2 transcriptional cofactor originally described in *B. mori* (Li et al., 1997; Liu et al., 2000; C. Zhou et al., 2016). The presence of intronless *β-repats* orthologs across both lepidopteran and non-lepidopteran insects further suggests that this subgroup may approximate the ancestral REPAT repertoire. In contrast, the rapidly evolving *α-* and *γ-repat* lineages have undergone lineage-specific expansion, tandem organization, and variation in copy number among species. Strikingly, nearly all REPATs implicated in plant interactions, including previously characterized effectors (Chen et al., 2019, 2023; X. Wang et al., 2024; Zhang et al., 2024) and the virus-responsive oral-secreted REPATs identified here, belong to the α- and γ-REPAT lineages. Although α-REPATs lack a detectable MBF2 domain, AlphaFold predicts an almost identical three-dimensional fold, suggesting that structural conservation has been maintained despite extensive sequence divergence.

Although the selective forces driving *α-* and *γ-repat* diversification remain unclear, their evolutionary dynamics, including tandem expansion, copy-number variation, and plastic regulation by both, host plants (García-Marin et al., 2025; Zhang et al., 2024) and pathogen infection, are hallmarks of gene families operating at the interface between organisms, where rapid adaptation to changing biotic interactions is favored (Hernandez et al., 2025). Consistent with this idea, generalist Lepidoptera harbor substantially larger *α-* and *γ-repat* repertoires than specialist species, suggesting that diversification of these lineages may have contributed to adaptation to diverse ecological niches (Hughes et al., 2018). This architecture resembles that of several gene families involved in biotic interactions, including detoxification CYP450 enzymes, whose repertoires expand with host breadth (Dermauw et al., 2020), and rapidly evolving plant nucleotide-binding leucine-rich repeat receptors (known as NLR) involved in pathogen recognition (Tamborski and Krasileva, 2020). Collectively, these observations suggest that REPAT evolution partitioned the family into a conserved ancestral module and a rapidly evolving interface specialized for ecological interactions. While β-REPATs likely retained ancestral functions, diversification of α- and γ-REPATs appears to have provided the molecular substrate for adaptation to changing interactions with both microbes and host plants.

A striking finding of this study is that two viruses with fundamentally different life histories (Mattia et al., 2025; Rohrmann, 2019) exert opposite effects on REPAT abundance in caterpillar OS. Infection with the lethal baculovirus SeMNPV increased the abundance of multiple REPAT proteins, whereas infection with the covert iflavirus SeIV1 consistently reduced their abundance under two different dietary conditions. Remarkably, these opposite changes were mirrored by reciprocal effects on plant defense activation, with SeMNPV attenuating JA-mediated responses and SeIV1 enhancing them. Notably, previous work has shown that direct application of a baculovirus-killed cadavers to tomato plants does not alter their responses to herbivory, further supporting the possibility that the contrasting effects observed here are mediated indirectly through changes in the herbivore (Jones et al., 2025). The induction of REPATs by SeMNPV is consistent with previous transcriptomic studies reporting strong REPAT activation following baculovirus infection (Chen et al., 2020; Herrero et al., 2007; Jakubowska et al., 2013). Baculoviruses initiate infection in the midgut before disseminating systemically (Rohrmann, 2019), a process expected to strongly activate stress- and damage-associated pathways that induce REPAT expression. By contrast, SeIV1 follows a fundamentally different infection strategy, establishing persistent, largely asymptomatic infections that are efficiently transmitted vertically and appear to coexist with their host over multiple generations (Jakubowska et al., 2014; Virto et al., 2014). Under this lifestyle, repression of REPATs may reflect a stable virus–host equilibrium in which immune activity is dampened rather than activated, potentially contributing to a cellular environment more compatible with long-term persistence. Thus, viruses with contrasting infection strategies appear to shift the REPAT repertoire in opposite directions, with corresponding consequences for host plant defense activation.

Because baculoviruses are well known to manipulate host behaviour and physiology to maximize viral transmission (Gasque et al., 2019; Ikeda et al., 2015), increased REPAT abundance and the associated attenuation of plant defenses could represent an additional component of this extended phenotype. By attenuating JA-mediated defenses, increased REPAT abundance could improve larval feeding efficiency during infection, thereby providing additional resources for viral replication before host death. However, the observation that the evolutionarily unrelated iflavirus SeIV1 also modifies REPAT abundance, albeit in the opposite direction, suggests that REPAT regulation may represent a general consequence of how different viral lifestyles shape host physiology, rather than a genuine manipulation by baculoviruses. An alternative, and not mutually exclusive, interpretation is that plants exploit REPAT proteins as indicators of herbivore physiological state. Inducible defenses are metabolically costly, and plants are expected to adjust their investment according to the threat posed by an attacking herbivore (Cipollini et al., 2014; Züst and Agrawal, 2017). A caterpillar undergoing a lethal baculovirus infection represents a transient threat whose feeding period will soon terminate, whereas a persistently infected SeIV1 carrier remains capable of sustained herbivory. Under this scenario, dampening defenses against terminally infected larvae while mounting stronger responses against persistently infected herbivores could optimize the cost–benefit balance of defense allocation

In conclusion, by combining comparative genomics, proteomics, and plant functional assays, we identify REPAT proteins as a molecular interface linking the infection status of herbivorous insects to plant defense responses. Regardless of the underlying mechanism, our results demonstrate that microbial infection can reshape plant–herbivore interactions by modifying the repertoire of insect effectors delivered during feeding. Rather than acting exclusively within the insect host, the consequences of viral infection extend across trophic levels, altering the molecular information transmitted to the plant and ultimately its defensive phenotype. These findings broaden our understanding of how microorganisms influence ecological interactions and establish REPAT proteins as a mechanistic link connecting microbes, insects, and plants.

## Materials and methods

### Gene identification and manual annotation

Genome assemblies and annotated protein datasets from the species analyzed were retrieved from the sources listed in Table S1. *repat* genes were identified using a custom Hidden Markov Model (HMM) constructed from PF15868 (MBF2 transcriptional cofactor) domain sequences of *Spodoptera exigua* and *Bombyx mori* retrieved from UniProt (The UniProt Consortium, 2025). The HMM was constructed from the alignment of the ten annotated MBF2-containing REPAT proteins from *B. mori* (C.-Y. Zhou et al., 2016) together with the 26 conserved MBF2-containing REPATs from *S. exigua* deposited in NCBI (accessed October 2025) and used to screen all predicted protein datasets.

Significant HMM hits (E-value < 1 × 10⁻⁵) were used as queries in iterative PSI-BLAST searches until convergence. All retrieved sequences, together with previously reported *S. exigua* REPAT proteins (Hernández-Martínez et al., 2010; Herrero et al., 2007; Navarro-Cerrillo et al., 2013b), were subsequently used as queries in TBLASTN searches against genomic assemblies. Genomic regions surrounding each significant hit (E-value < 1 × 10⁻³) were manually inspected to define exon–intron structures by comparison with known *S. exigua repat* genes, following canonical GT–AG splice junctions.

Predicted proteins were validated by assessing the presence of signal peptides using SignalP 6.0 (Teufel et al., 2022) and MBF2 domains using the Pfam database.

### Protein structure comparison

Structural comparisons between REPAT pairs were performed using predicted protein structures retrieved from the AlphaFold Protein Structure Database (Bertoni et al., 2026). The two protein models were superimposed in UCSF ChimeraX (version 1.12) using the MatchMaker tool with default parameters (Meng et al., 2023). The resulting structural alignment was used to compare the overall protein folds and identify conserved and divergent structural features.

### Phylogenetic reconstruction and gene nomenclature

Coding sequences of all lepidopteran *repat* genes were aligned using PRANK (Löytynoja, 2014) implemented in TranslatorX (Abascal et al., 2010). Alignments were manually inspected and curated to remove poorly aligned regions before phylogenetic reconstruction (Dataset S2). Because the C-terminal region of *β-repat13* orthologs was highly divergent, only the conserved region encompassing the MBF2 domain was included in the analysis. Maximum-likelihood phylogenies were reconstructed with RAxML (Stamatakis, 2014) under the GTRGAMMA substitution model using 500 bootstrap replicates. Trees were visualized and edited with iTOL (Letunic and Bork, 2024).

Orthology assignments integrated phylogenetic position, conserved gene structure, and chromosomal synteny. When one-to-one orthology could not be confidently resolved, genes were assigned to broader orthologous groups containing multiple lineage-specific paralogs. *repat* genes were classified into α-, β-, and γ-groups according to their phylogenetic position, gene structure, and chromosomal location. Within each group, genes were numbered according to their genomic position. Gene names were preceded by a four-letter species abbreviation consisting of the first letter of the genus followed by the first three letters of the species name (e.g., *Spodoptera exigua*, Sexi; *Tuta absoluta*, Tabs). Orthologs of *S. exigua* retained the same numerical designation (e.g., *Slitu-β-repat4*, *Bmor-β-repat15*). When multiple lineage-specific paralogs occurred within the same orthologous group, they were distinguished by letter suffixes.

### Gene birth-death analyses

Evolutionary gains and losses of *β-repat* genes were inferred using CAFE (De Bie et al., 2006). Gene copy numbers for each orthologous group (Supplementary Figure 1B) were analyzed under a maximum-likelihood birth–death model to estimate ancestral family sizes, lineage-specific duplication and loss events, and the global turnover rate (λ). Divergence times for the species tree were obtained from published Lepidoptera phylogenies (Kawahara et al., 2019; Kergoat et al., 2021).

### Insect rearing and virus infections

A laboratory colony of *S. exigua* originally obtained from Andermatt Biocontrol AG (Switzerland) has been maintained at the University of Valencia (Spain) for more than 200 generations. Unless otherwise indicated, larvae were reared on artificial diet (Elvira et al., 2010) at 25 ± 2 °C, 70% relative humidity, and a 16:8 h light:dark photoperiod.

Viral infections were established using the droplet-feeding method (Hughes et al., 1986). Briefly, larvae were placed individually in Petri dishes containing 4 μL droplets arranged in a circle. Each droplet consisted of 10% sucrose in phosphate-buffered saline (PBS, pH 7.4), 10% phenol red as a feeding tracer, and the appropriate viral suspension. After visual confirmation of droplet ingestion, larvae were transferred individually to bioassay trays containing artificial diet and maintained under the rearing conditions described above. Mock-treated larvae received the same solution without virus.

For SeMNPV infections, newly molted fourth-instar larvae were infected with the SP2 isolate (Caballero et al., 1992) at a concentration of 5 × 10⁷ occlusion bodies (OBs)/mL. For SeIV1 infections, newly molted second-instar larvae were infected with 1 × 10⁹ viral genome equivalents/mL (Millán-Leiva et al., 2012).

### RNA extraction, sequencing and expression analysis

Midguts from newly molted fifth-instar *S. exigua* larvae were hand-dissected, pooled in groups of 10 larvae, immediately frozen in liquid nitrogen, and stored in 1.5-mL microcentrifuge tubes. Three independent biological replicates were collected. Gut tissues were homogenized using RNase-free disposable pestles, and total RNA was extracted with TriPure™ Isolation Reagent (Roche) according to the manufacturer’s instructions.

RNA quality assessment, library preparation, and sequencing were performed by Macrogen Inc. (Seoul, South Korea). Libraries were prepared using the TruSeq Stranded Total RNA LT Sample Preparation Kit with Ribo-Zero (Illumina) and sequenced on an Illumina NovaSeq 6000 platform to generate 150-bp paired-end reads. Raw sequencing data have been deposited in the NCBI Sequence Read Archive under BioProject accession PRJNA1046378.

Raw reads were quality-filtered with Trimmomatic (Bolger et al., 2014), and residual ribosomal RNA reads were removed using SortMeRNA (Kopylova et al., 2012). Filtered reads were aligned to the *S. exigua* reference genome (GCA_902829305.4) using HISAT2 (Kim et al., 2019). Between 70 and 78% of reads mapped uniquely to the genome or annotated splice junctions, yielding approximately 12.7–14.7 million uniquely mapped reads per sample.

The manually curated repat annotation generated in this study was incorporated into the official S. exigua gene annotation (13,002 annotated genes; October 2022 release). Gene-level read counts were obtained using featureCounts (Liao et al., 2014). TPM values were calculated from raw counts and gene lengths using edgeR (Robinson et al., 2010).

### Oral secretion collection

Oral secretions (OS) were collected from newly molted fifth-instar *S. exigua* larvae for proteomic analyses of SeMNPV- and SeIV1-infected insects. For SeMNPV experiments, larvae were maintained on artificial diet and OS was collected 72 h post-infection. For SeIV1 experiments, larvae were fed either artificial diet or pepper leaves for 48 h before OS collection. OS was collected as previously described (García-Marin et al., 2025). Briefly, larvae were gently held between the thumb and index finger, and the mouthparts were lightly stimulated with a 10-µL pipette tip to induce regurgitation. Secretions were pooled on ice throughout the collection process. Each biological replicate consisted of pooled OS from 10 larvae in the SeMNPV experiments and from 50 larvae in the SeIV1 experiments.

Samples were centrifuged at 1,000 × g for 5 min at 4 °C to remove debris. The supernatants were immediately frozen in liquid nitrogen and stored at −80 °C until proteomic analysis. Five biological replicates were collected for each condition, except for SeMNPV-infected larvae, for which six biological replicates were analyzed.

### Proteomic sample preparation

Oral secretion proteins from SeIV1-infected larvae were precipitated using the 2-D Clean-Up Kit (Cytiva) according to the manufacturer’s instructions. Protein pellets were resuspended in 25 μL of 1× Laemmli buffer and denatured at 95 °C for 5 min. SeMNPV samples were processed directly without precipitation and diluted in Laemmli buffer before denaturation.

Protein extracts (15 μg for SeIV1 samples and 30 μg for SeMNPV samples) were separated by one-dimensional SDS-PAGE. Gel bands were excised and digested overnight at 37 °C with sequencing-grade trypsin (Promega) in 50 mM ammonium bicarbonate (Shevchenko et al., 1996). Digestion was stopped by addition of trifluoroacetic acid (TFA, 1% final concentration), followed by two acetonitrile extraction steps. Peptides were dried in a SpeedVac and resuspended in 2% acetonitrile/0.1% TFA (SeIV1) or 0.1% TFA (SeMNPV).

### Mass spectrometry

Because the SeMNPV and SeIV1 proteomic datasets were generated independently, peptide samples were analyzed using two data-independent acquisition (DIA) mass spectrometry workflows optimized for each experiment. For SeMNPV samples, 2.5 μL of each peptide digest was diluted to 20 μL with 0.1% formic acid (FA) and loaded onto an Evotip Pure tip (Evosep) according to the manufacturer’s instructions. Peptides were separated on an Endurance analytical column (15 cm × 150 μm, 1.5 μm; Evosep) using an Evosep One system with the 30 samples-per-day method.

Eluted peptides were ionized using a CaptiveSpray source (Bruker) operated at 1,600 V and 180 °C and analyzed on a timsTOF fleX mass spectrometer (Bruker) operating in diaPASEF mode. TIMS was configured with a mobility range of 0.6–1.6 Vs cm⁻² (1/K₀), a ramp time of 100 ms, a duty cycle of 100%, and a ramp rate of 9.42 Hz. MS survey scans were acquired over an m/z range of 100–1,700.

For SeIV1 samples, 5 μL of each peptide digest was loaded onto a trap column (3 μm C18-CL, 350 μm × 0.5 mm; Eksigent Technologies) using an Ekspert nanoLC 425 system (Eksigent Technologies) and desalted with 0.1% TFA at 5 μL min⁻¹ for 3 min. Peptides were separated on an analytical column (3 μm C18-CL, 120 Å, 75 μm × 150 mm; Eksigent Technologies) equilibrated in 5% acetonitrile containing 0.1% formic acid.

A linear gradient from 7 to 40% solvent B (acetonitrile, 0.1% FA) in solvent A (0.1% FA) was applied over 60 min at a flow rate of 300 nL min⁻¹. Peptides were analyzed using a TripleTOF 6600+ mass spectrometer (SCIEX) equipped with an Optiflow nanoelectrospray source operated at 3.0 kV and 200 °C. Data were acquired in SWATH mode using a TOF MS survey scan (350–1,250 m/z, 0.05 s), followed by product ion scans collected across 100 variable isolation windows (400–1,250 m/z, 0.08 s each), resulting in a total cycle time of 2.79 s.

### Statistical analysis of proteomic data

Protein abundance data were analyzed in R (version 4.5.2). Label-free quantitates (LFQ) intensities were log2-transformed and proteins with valid measurements in fewer than 70% of samples were excluded. Missing values were imputed using a down-shifted Gaussian distribution. Differential abundance analysis was performed using the limma package (Ritchie et al., 2015), applying an FDR < 0.05 and log2FC > 0.5 as significance thresholds. Heatmaps were generated from Z-score-transformed abundances. Individual REPAT proteins were additionally compared using Welch’s t-test.

### Tomato herbivory assays

Tomato plants (*Solanum lycopersicum* cv. Moneymaker) were germinated in coconut fiber pellets (Jiffy) and transplanted two weeks after germination into 12 × 12 cm pots containing a 2:1 mixture of peat and perlite. Plants were grown in a controlled greenhouse at 24 ± 2 °C (University of Valencia, Spain). Four-week-old plants were transferred to a controlled growth chamber (24 ± 2 °C, 16 h light:8 h dark photoperiod) three days before the experiments.

To induce defense responses, plants were exposed for 3 h to feeding by newly molted fifth-instar *S. exigua* larvae that were either mock-treated or infected with SeMNPV (72 h post-infection). Two larvae were confined using clip cages to the terminal leaflet of the third and fourth true leaves. Mechanically wounded plants (tracing wheel plus clip cage) served as controls. Each plant represented one biological replicate, with five plants analyzed per treatment. Three independent experiments were performed.

After the 3-h feeding period, larvae and clip cages were removed. Terminal leaflets from the third true leaf were photographed for leaf area quantification using LeafByte (Getman-Pickering et al., 2020), immediately frozen in liquid nitrogen, and stored at −80 °C for RNA extraction. Twenty-four hours after the start of the experiment, both the terminal and adjacent leaflets of the fourth true leaf were collected to evaluate systemic defense responses. Larvae removed from plants were maintained individually on artificial diet to confirm SeMNPV infection.

Total RNA was extracted from 50 mg of frozen leaf tissue using TRIdity GTM reagent (AppliChem) following the manufacturer’s instructions. RNA (1 μg) was treated with DNase I (Thermo Fisher Scientific) and reverse-transcribed using PrimeScript RT Reagent (Takara Bio). Quantitative real-time PCR (qRT-PCR) was performed on a StepOnePlus Real-Time PCR System (Applied Biosystems) using HOT FIREpol EvaGreen qPCR Mix Plus (ROX) (Solis Biodyne). Primer sequences are listed in Table S2. Relative gene expression was calculated as the 2^−ΔCt (Livak and Schmittgen, 2001), with SlEF used as the reference gene. Differences in gene expression among treatments were analyzed using analysis of covariance (ANCOVA), with treatment included as a fixed categorical factor and experiment included as a covariate. Post hoc comparisons among treatments were performed using Tukey’s honestly significant difference (HSD) test. Prior to ANCOVA, a two-way ANOVA was conducted to test for treatment × experiment interactions.

### Pepper herbivory assay, gene expression, and metabolomic analyses

Pepper plants (*Capsicum annuum* cv. Lipari) were grown in a controlled growth chamber at the Instituto Valenciano de Investigaciones Agrarias (IVIA, Spain) under 25 ± 2 °C, 65 ± 10% relative humidity, and a 14 h light:10 h dark photoperiod. Seeds were germinated and transplanted individually into plastic pots (8 x 8 x 8 cm) two weeks after germination. Fully developed plants (approximately 20 cm in height) were used for all experiments.

Three experimental treatments were established: (i) uninfested control plants, (ii) plants infested with ten SeIV1-free *S. exigua* neonates, and (iii) plants infested with ten *S. exigua* neonates vertically infected with SeIV1. Each biological replicate consisted of six individually caged plants (BugDorm), and the complete experiment was independently repeated twice.

Vertically infected neonates were obtained from adults that had been experimentally infected with SeIV1 as second-instar larvae using the droplet-feeding procedure described above. Following adult emergence, infected moths were allowed to mate, and vertical transmission of SeIV1 to the offspring was confirmed by RT-qPCR as previously described (Mengual-Martí et al., 2022).

Forty-eight hours after infestation, the apical portion of each plant was harvested, frozen in liquid nitrogen, and ground to a fine powder. The resulting material was divided for gene expression and metabolomic analyses.

For gene expression analyses, total RNA was extracted using NZYol reagent (NZYTech, Lisbon, Portugal). One microgram of RNA was treated with TURBO DNA-free™ Kit (Thermo Fisher Scientific) and reverse transcribed using PrimeScript™ RT Reagent Kit (Takara Bio). Quantitative PCR was performed on a LightCycler® 480 System (Roche) using NZYSpeedy qPCR Green Master Mix (NZYTech). Expression of the jasmonate-responsive marker gene *PIN2* (Proteinase Inhibitor II) was quantified. Primer sequences are listed in Table S2. Gene expression was normalized against *CaEF1* (Bouagga et al., 2018), and transcript abundance was calculated as the 2^−ΔCt method (Livak and Schmittgen, 2001). Differences in gene expression among treatments were analyzed using analysis of covariance (ANCOVA), with treatment included as a fixed categorical factor and experiment included as a covariate. Post hoc comparisons among treatments were performed using Tukey’s honestly significant difference (HSD) test. Prior to ANCOVA, a two-way ANOVA was conducted to test for treatment × experiment interactions.

For untargeted metabolomic analysis, metabolites were extracted from 30 mg of freeze-dried tissue with 1 mL of 10% methanol, incubated on ice for 30 min, centrifuged at 15,000 × *g* for 15 min at 4 °C, and filtered through 0.2 μm regenerated cellulose filters. Aliquots (20 μL) were analyzed on an Acquity UPLC system (Waters, Milford, MA, USA) coupled to a QTOF Premier mass spectrometer operating in both positive and negative electrospray ionization modes. Data were acquired using MassLynx v4.2 (Waters). Chromatographic features were detected, aligned, and corrected using XCMS in R. Peak intensities were normalized to sample dry weight, and positive- and negative-ion datasets were combined, median-normalized, cube-root transformed, and Pareto-scaled using MetaboAnalyst before multivariate analyses.

## Acknowledgements

This study was supported by projects (PID2024-162058O-BC32, PID2024-162058OB-C33, PID2024-158774OB-I00, and CNS2025-165452), funded by the Spanish Research Agency and Ministry of Science, Innovation and Universities (MICIU/AEI/10.13039/501100011033), by the Human Frontier Scientific Program (project No. RGY0052/2022) and by the IVIA-52202B project from the Valencian Institute of Agricultural Research (IVIA) of the Valencian Government (GVA) (this project is eligible for co-financing by the European Union through the ERDF Operational Program). C. Crava and M. Pérez-Hedo were recipients of a Ramón y Cajal grant from the Spanish Research Agency and the Ministry of Science, Innovation and Universities (No. MRR/RYC2021-033098-I and RYC2022-036061-I, respectively). R. Becerra was supported by a Santiago Grisolía fellowship from the Generalitat Valenciana (No. CIGRIS/2021/077). We thank Rosa María González-Martínez, for their excellent help with insect rearing and laboratory management.

