## Supplementary Figures for "Insect REPAT proteins mediate cross-kingdom communication in microbe–insect–plant interactions"

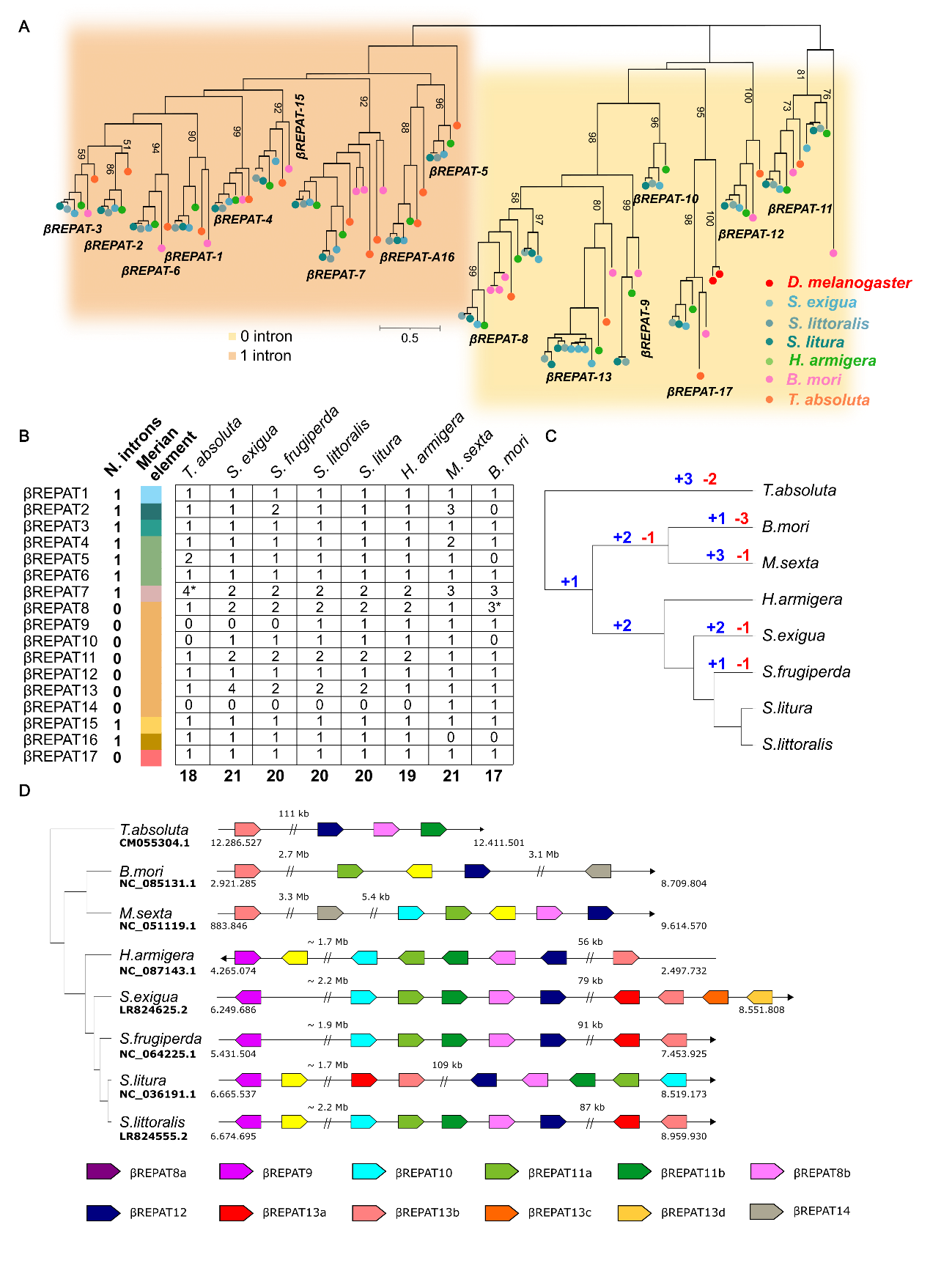
**Supplementary Figure 1**. **Evolutionary origins and diversification of lepidopteran *β-repat* genes**

(**A**) Phylogenetic relationships of the lepidopteran *β-repat* coding sequences. The sequences were aligned with PRANK and the tree was built with RAxML under the GTRGAMMA model of substitution with 500 bootstrap replicates. The scale bar indicates the expected number of substitutions per site. (**B**) Presence–absence matrix of *β-repat* genes across seven lepidopteran species. Merian elements indicate the chromosomal element in which each gene is located. Asterisks denote orthologous groups containing one or more copies located on different Merian elements relative to the remaining members of the group (**C**) Gene gain and loss events inferred for *β-repat* genes and mapped onto the lepidopteran species tree adapted from (Kawahara et al., 2019; Kergoat et al., 2021). CAFE inferred a global gene family turnover rate of λ = 0.241569 across the phylogeny, with a total of 15 inferred gene gain events (gene birth rate, B = 0.0072 gene⁻¹ Myr⁻¹) and 9 gene loss events (gene death rate, D = 0.0043 gene⁻¹ Myr⁻¹). (**D**) Tandem organization of the intronless *β-repat8–14* genes, which are located on the same chromosome in all analyzed species, suggesting that tandem duplication through non-allelic homologous recombination has contributed to the expansion of this gene cluster during lepidopteran evolution.

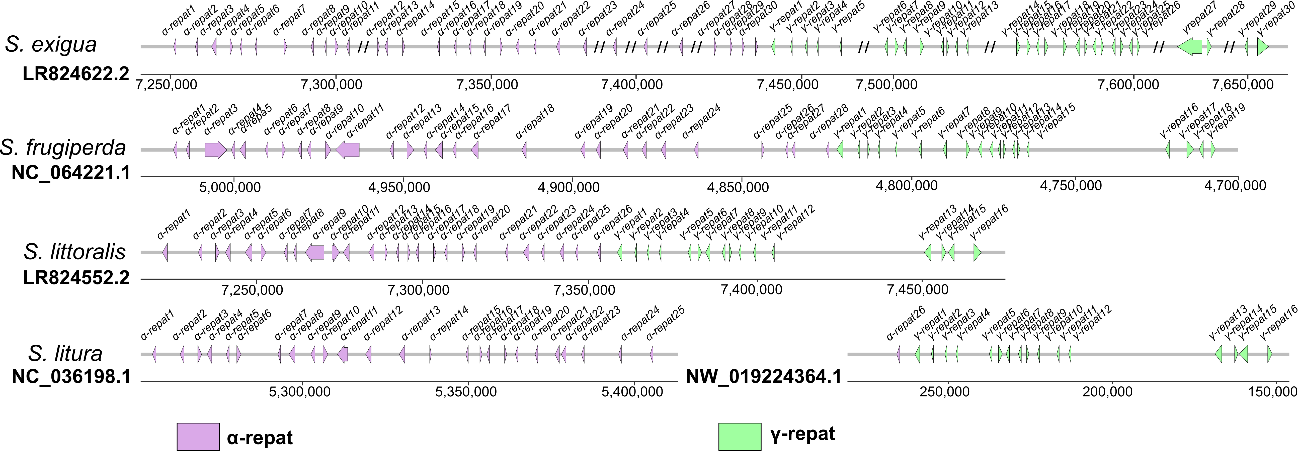

**Supplementary Figure 2. Tandem organization of *α-* and *γ-repat* genes across *Spodoptera* genomes**

*α-repat* and *γ- repat* genes are predominantly organized in tandem arrays located on a single chromosome in most analyzed species where a subset of *γ- repat* genes is found on additional chromosomes, and *S. litura* and *S. littoralis* which possess a single *α-repat* gene located in a different chromosome (Supplementary dataset 1).

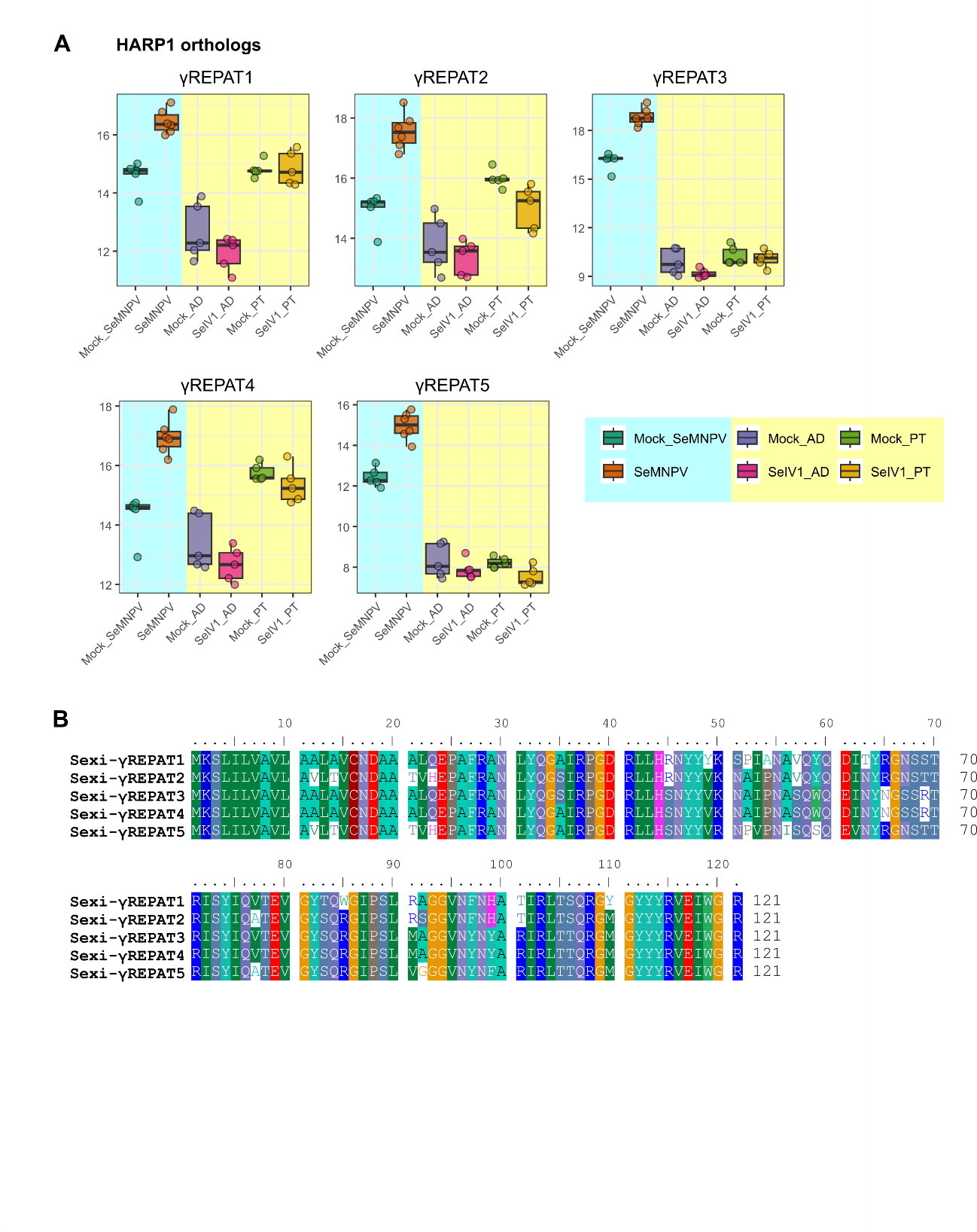
Supplementary Figure 3. Differential regulation and sequence conservation of HARP1 orthologs in *S. exigua*.

**(A)** Boxplots showing the LFQ (label-free quantity) abundances of the five S. exigua orthologs of the plant defense effector HARP1 following infection with SeMNPV (blue shading) or SeIV1 (yellow shading). Data for the two viruses were obtained from independent proteomic experiments. Protein abundances were quantified by quantitative proteomics of larval oral secretions. AD, artificial diet; PT, pepper-fed larvae. **(B)** Amino acid sequence alignment of the five S. exigua HARP1 orthologs. Shading indicates residues conserved in at least 60% of the sequences.

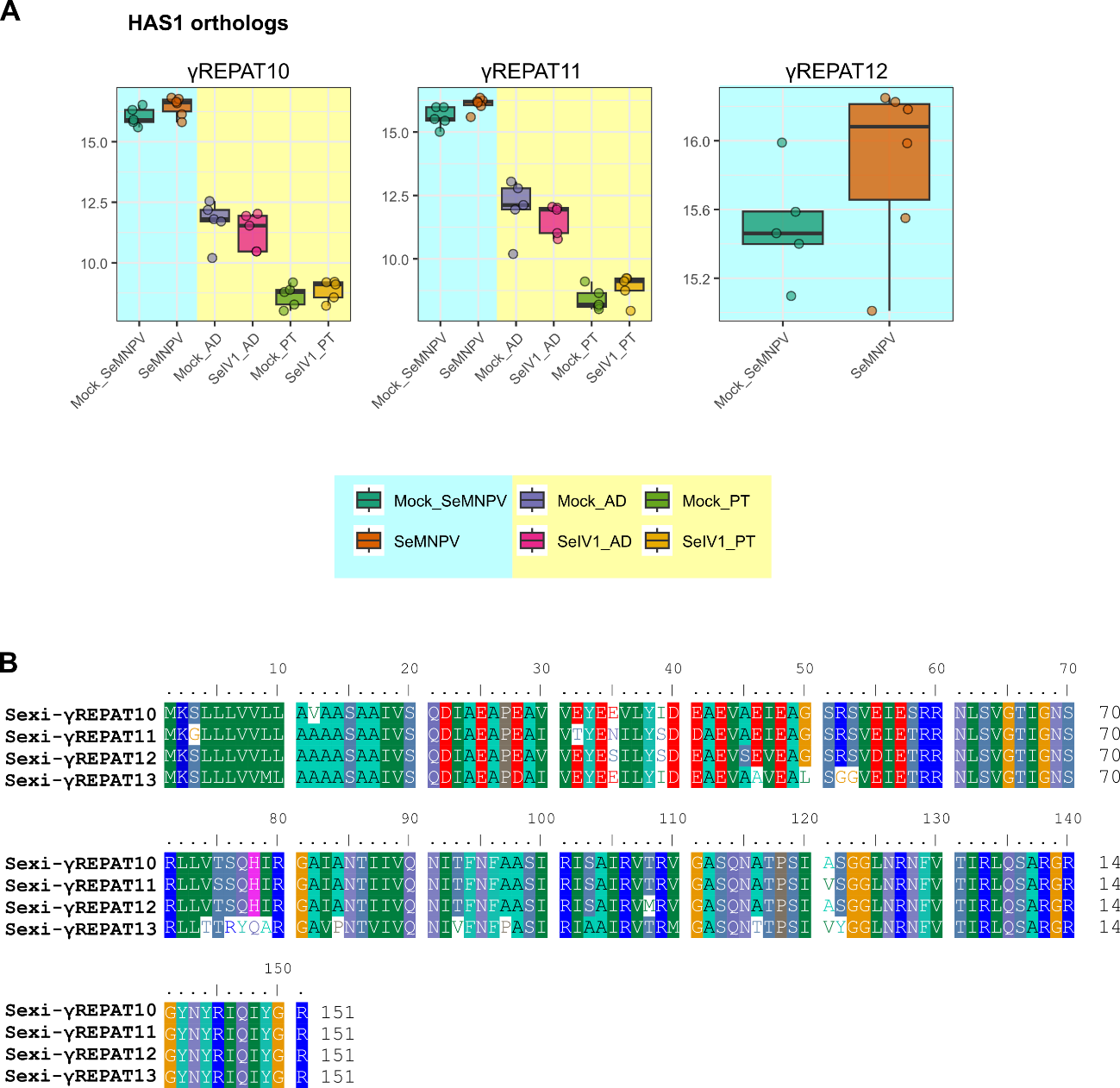

Supplementary Figure 4. Differential regulation and sequence conservation of HAS1 orthologs in *S. exigua*.

**(A)** Boxplots showing the LFQ (label-free quantity) abundances of the four S. exigua orthologs of the plant defense effector HAS1 following infection with SeMNPV (blue shading) or SeIV1 (yellow shading). Data for the two viruses were obtained from independent proteomic experiments. Protein abundances were quantified by quantitative proteomics of larval oral secretions. AD, artificial diet; PT, pepper-fed larvae. **(B)** Amino acid sequence alignment of the four S. exigua HAS1 orthologs. Shading indicates residues conserved in at least 60% of the sequences.

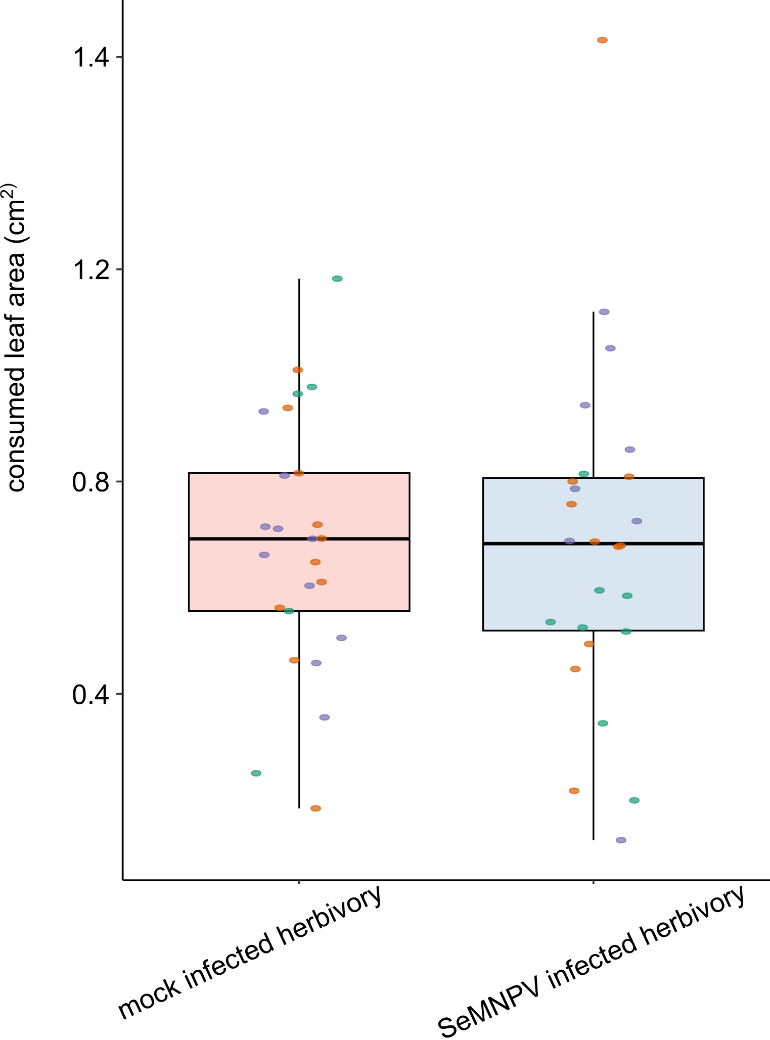

Supplementary Figure 5 SeMNPV infection does not alter leaf consumption by *S. exigua* larvae.

Boxplot showing the leaf area consumed by individual clip-caged *S. exigua* larvae during a 3-h feeding period. Leaf area consumed did not differ significantly between mock- and SeMNPV-infected larvae.

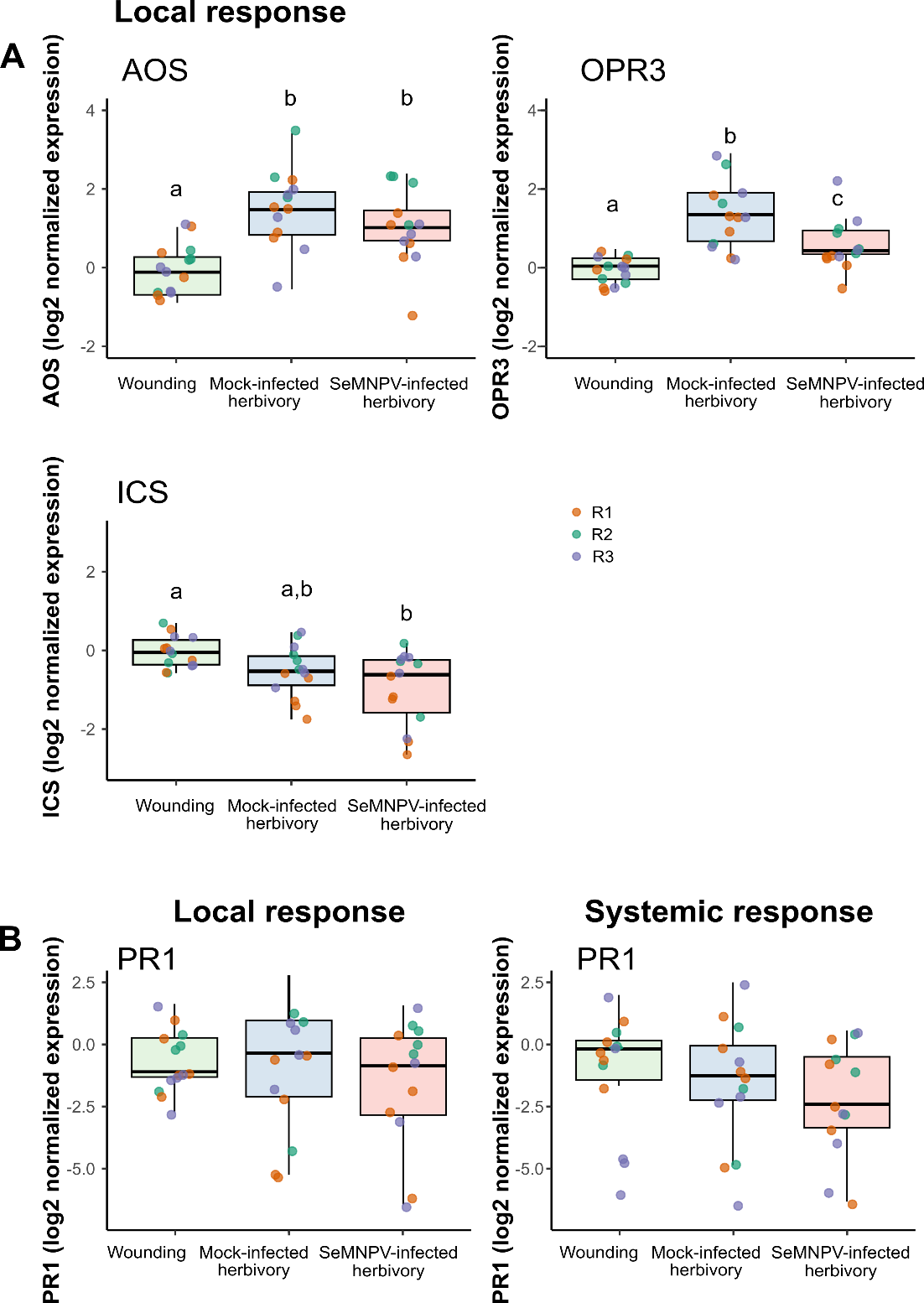

Supplementary Figure 6 Herbivory by SeMNPV-infected larvae reduce jasmonate biosynthetic responses without affecting salicylic acid signaling in tomato.

(**A**) Relative expression of the jasmonate (JA) biosynthetic genes AOS (allene oxide synthase) and OPR3 (12-oxophytodienoate reductase 3), and the salicylic acid (SA)-related gene ICS (isochorismate synthase) in tomato leaflets subjected to mechanical wounding or 3 h of herbivory by mock- or SeMNPV-infected larvae. Gene expression was measured in locally damaged leaflets at the end of the treatment. (**B**) Relative expression of the SA-responsive marker PR1 (pathogenesis-related protein 1) in tomato leaflets subjected to mechanical wounding or 3 h of herbivory by mock- or SeMNPV-infected larvae. Expression was measured 24 h after treatment in locally damaged leaflets (left panels) and adjacent systemic leaflets (right panels).

### Supplementary tables

**Table S1. Sources of genomic and protein sequence data.**

| Species | Genome version | Proteome version | Repository |
| --- | --- | --- | --- |
| *S. exigua* | GCA_902829305.4 | Yes | NCBI |
| *S. frugiperda* | GCF_023101765.2 | Yes | NCBI |
| *S. litura* | GCF_002706865.1 | Yes | NCBI |
| *S. littoralis* | GCA_902850265.1 | Yes | NCBI |
| *H. armigera* | GCF_030705265.1 | Yes | NCBI |
| *B. mori* | GCF_030269925.1 | Yes | NCBI |
| *M. sexta* | GCF_014839805.1 | Yes | NCBI |
| *P. absoluta* | GCA_027580185.1 | Yes | NCBI |
| *D. melanogaster* | dmel_r6.66_FB2025_05 | Yes | Flybase |
| *A. aegypti* | GCF_002204515.2 | Yes | NCBI |
| *T. castaneum* | GCF_031307605.1 | Yes | NCBI |
| *A. mellifera* | GCF_003254395.2 | Yes | NCBI |
| *A. pisum* | GCF_005508785.2 | Yes | NCBI |
| *N. lugens* | GCF_014356525.2 | Yes | NCBI |
| *C. formosanus* | GCA_013340265.1 | Yes | NCBI |
| *I. elegans* | GCF_921293095.1 | Yes | NCBI |
| *T. domestica* | GCF_964235325.1 | Yes | NCBI |
| *T. urticae* | GCF_000239435.1 | Yes | NCBI |
| *D. pulex* | GCF_021134715.1 | Yes | NCBI |
| *C. elegans* | GCF_000002985.6 | Yes | NCBI |

**Supplementary Table S2.** List of the primers used for RT-qPCR.

| **Primer name** | **Target gene (abbreviation)** | **Sequence (5’ – 3’)** | **Reference** |
| --- | --- | --- | --- |
| **Tomato (*Solanum lycopersicum*)** | | | |
| SlEF_F | Elongation factor 1 alpha (*SlEF*) | GATTGGTGGTATTGGAACTGTC | (Martinez-Medina et al., 2013) |
| SlEF_R |  | AGCTTCGTGGTGCATCTC | (Martinez-Medina et al., 2013) |
| AOS_F | Allene oxide synthase (*AOS*) | AACAGTGTGCCGGAAAAGAC | This study |
| AOS_R |  | AATGGAGATGCACCGACTTC | This study |
| OPR3_F | 12-oxophytodienoate reductase 3  (*OPR3*) | GCATATGGGCAAACTGAAGC | This study |
| OPR3_R |  | ATGAATGTCCCCTGATACGC | This study |
| PIN2_F | Wound-induced proteinase inhibitor 2  (*PIN2*) | GAAAATCGTTAATTTATCCCAC | (Martinez-Medina et al., 2013) |
| PIN2_R |  | ACATACAAACTTTCCATCTTTA | (Martinez-Medina et al., 2013) |
| PPO_F | Polyphenol oxidase  (PPO) | CACCACCAACTTTTGGAACC | This study |
| PPO_R |  | TACGCTGGCGATAACTGAAC | This study |
| ICS_F | Isochorismate synthase  (*ICS*) | CGCCTCAGCATTTGTAGTGA | This study |
| ICS_R |  | CTTGAACACACAGCCTCCAA | This study |
| PR1_R | Pathogenesis related protein 1  (*PR1*) | GTGGGATCGGATTGATATCCT | (Martinez-Medina et al., 2013) |
| PR1_F |  | CCTAAGCCACGATACCATGAA | (Martinez-Medina et al., 2013) |
| **Pepper (*Capsicum annum*)** | | | |
| CaEF1a_F | Elongation factor 1 alpha  (CaEF1) | TGAAGAATGGTGATGCTGGC | (Bouagga et al., 2018) |
| CaEF1a_R |  | GACAACACCAACAGCAACAG | (Bouagga et al., 2018) |
| CaPIN2_F | Proteinase inhibitor II  (PIN2) | CTTGCCCCAAGAATTGTGAT | (Bouagga et al., 2018) |
| CaPIN2_R |  | GCCCTAGCGTATTACGGAGA | (Bouagga et al., 2018) |
