## Supplementary Dataset S2 for "Insect REPAT proteins mediate cross-kingdom communication in microbe–insect–plant interactions"

>Bmor-betaREPAT1

---------MT------S-F----------K---L-I-SCA--V--L-------L-----L------V-----------L---T---------------I---------------------------------------------------------------------------I-------------SATD-------------------------------------------------------------------------------------------------------------------------------------------------------------------------------------------------------L---------QVG-V----------SVN--------N----------------------------------------E--LAY--V-----------KNV------------------N-RN-AVPL-----K-V-RTKNVFY----SR-----N-----D--------T--------------RV-IRG------------I---------S---A-----I-D-----------------L--D-A--G-N--AK-P--TITSGGVGFSFTNIKIKS-ERG--EGLNY-Q-------VQI-F---------------I--------------------------------------------------------------------------------------------------------------------------------------------------------------------------------------------------------------------------------------------------------------------------------------------------------------------------------

>Bmor-betaREPAT11a

----MSKY--------------------------L-F----ICL--I-------C-----A---Y--I-----------S---T---------------G---------------------------------------------------------------------------H------G---K-----T-------------------------------------------------------------------------------------------------------------------------------------------------------------------------------------------------------V---------IVG-Q----------TYDYNNV----V----------------------------------------R--KVH--E-----------QSV------------------E-AN-AIPL-----F-K-RVQEVFF----EF-----P-----Q--------P-E------------QK-IKG------------I---------A---I-----K-D-----------------L--E-N--G-L--AE-P--SITRGGLGFNFVNIKLKS-DRG--SGFKF-V-------IEI-Y---------------A--------------------------------------------------------------------------------------------------------------------------------------------------------------------------------------------------------------------------------------------------------------------------------------------------------------------------------

>Bmor-betaREPAT12

---------MR------A-F-------TV-L---L-I------A--C-------L-----L---S--A-----------C---S---------------A---------------------------------------------------------------------------Q------S---H-----D-------------------------------------------------------------------------------------------------------------------------------------------------------------------------------------------------------L---------SLG-Q----------LTY--------R----------------------------------------D--IVL--Y-----------KIN------------------EYKY-GFPF-----I-V-RTSEIEY-----P-EPGQQ-----N--------F--------------AY-ISA------------I---------Y---V-----K-DHY---------------T--D-G--N-G--GY-P--IVKSGGVGQKFVKLKLKS-QRG--HGFNF-T-------VTI-Y---------------G--------------------------------------------------------------------------------------------------------------------------------------------------------------K-----------------------------------------------------------------------------------------------Y-----------------------------------------------------------------

>Bmor-betaREPAT13

---------MY------R-L----------V---C-F------L--V-------------L------V-----------G---A---------------V---------------------------------------------------------------------------A------A---N-----D-------------------------------------------------------------------------------------------------------------------------------------------------------------------------------------------------------I---------VIG-H---------E--N--------G----------------------------------------K--KIY--D-----------SNR------------------T----ASPA-----IWR-RADKITV----NA-----T-----D--------N--------------EV-ISG------------V---------I---I-----T-D-----------------LRAE-K----D--GE-V--QIVEGGQGQKNVTFELKS-PAV-FRGYEF-H-------VEV-Y---------------S--------------------------------------------------------------------------------------------------------------------------------------------------------------V----P-ED--------E--------------------IQ----------------AE-VNTS-----------------------TGLQDAKL--------------------------------------------------------------------

>Bmor-betaREPAT14

--MNIYIK--------------------------I-I------I--L-------T-----S---S--C-----------A---S---------------I---------------------------------------------------------------------------L------C---Y-----D-------------------------------------------------------------------------------------------------------------------------------------------------------------------------------------------------------L---------YLG-S----------DFG--------A----------------------------------------R--MSY--------------REV------------------K-KQ-GLLF-----F-K-RSESLTF----EY-----P-----E--------G--------------TN-ITG------------I---------A---C-----V-D-----------------L--SPN--K-M--GS-V--TVEEGGIGFNSVSLYLES-ERS--ASLRY-T-------VEI-Y---------------I----------------------------------------------------------------------------PSFIKHTSDQNRIGVVFR-YKI------------------------------------------------------------------------------------------------------------------------------------------------------------------------------------------------------------------------------

>Bmor-betaREPAT15

---------MK-S----L-V----------L---L-----------C-------L-----S---L--C-----------V---C---------------V---------------------------------------------------------------------------H------S---Y-----H-------------------------------------------------------------------------------------------------------------------------------------------------------------------------------------------------------F---------LVGDI----------SSS----------------------------------------------------LAH--H-----------KLV------------------Q-YN-AIPF-----M-K-RVKNYFY----SS-----A-----D--------N--------------KI-ITG------------I---------Q---A-----L-D-----------------S--L-N--S-K--AT-V--NITAGGVGYPYVNMRMKS-ERG--SGLSY-D-------IGI-Y-VNQNY-LRI-----N--------------------------------------------------------------------------------------------------------------------------------------------------------------------------------------------------------------------------------------------------------------------------------------------------------------------------------

>Bmor-betaREPAT17

---------MK------I-L----------V---K-L------I--F-------I-----I---SN-V-----------D---A------RIF------C---------------------------------------------------------------------------E------K---N-----K-------------------------------------------------------------------------------------------------------------------------------------------------------------------------------------------------------M---------PL-------------------------------------------------------------------------------------EVG------------------EWHK-DYKQ-----M-D--SAIVEN-----S-----P-----V--------T--------------IN-EGQ------------L---------Y---V-----Y-ENYFPGHTIKYIHVDNLAI--R-S--C-G--AS-A--SIKNGGVGTSSVLIVLHA-DSN--QEIRS-V-------IDI-W---------------G-------V-K--------NY--E---KTT--RRAL--NP----ESLKNMKSLYLFKELRAVNHNG----GF----W----------------------------------------------------------------------------------R-----------------------------------------------------------------------------------------------------------------------------------------------------------------

>Bmor-betaREPAT3

---------MK-F-A--P-V----------L---I-F------I--F-------L-----S---L--V-----------C---V---------------I---------------------------------------------------------------------------Q------C---K-----D-------------------------------------------------------------------------------------------------------------------------------------------------------------------------------------------------------L---------IVG-T----------SFN--------K----------------------------------------R--LLW--Q-----------EKA------------------E-YN-AIPL-----K-K-RVKEVFF----SD-----P-----G--------Q--------------QL-IMG------------I---------I---A-----R-D-----------------L--D-H--S-D--AE-A--SITAGGIGFSYANIKLKS-PRG--SGLNY-Q-------LEI-Y---------------T--------------------------------------------------------------------------------------------------------------------------------------------------------------------------------------------------------------------------------------------------------------------------------------------------------------------------------

>Bmor-betaREPAT4

---------MR------L-V----------L---L-L------L--L-------S-----V---S--V-----------V---C---------------F---------------------------------------------------------------------------C---G--A---Y-----D-------------------------------------------------------------------------------------------------------------------------------------------------------------------------------------------------------M---------IVG-D----------TVH--------R----------------------------------------K--MVF--H-----------QRV------------------K-DF-AIPF-----K-K-RIKTLSY----TD-----P-----E--------K--------------RI-IKG------------V---------A---A-----I-D-----------------N--D-F--S-H--AS-A--NITEGGVGYSFVTVRMKS-QRH--HPLNF-E-------VEI-Y---------------L--------------------------------------------------------------------------------------------------------------------------------------------------------------------------------------------------------------------------------------------------------------------------------------------------------------------------------

>Bmor-betaREPAT6

---------MN------Y-K----------I-----------------------------V---F--V-----------L-LSV---------------VF--------------------------------------------------------------------------V------L---N-IEAKD-------------------------------------------------------------------------------------------------------------------------------------------------------------------------------------------------------F---------VDG-T----------KVN--------N----------------------------------------L--LIS--T-----------GKV------------------V-VK-GYPL-----I-K-RDKDYVY----VD-----P-----K--------L--------------RV-IKG------------I---------I---A-----R-D-----------------L--S-R--T-K--AE-V--TVTSGGVGATNVTLHLKS-ERG--EGLNY-L-------ILI-F---------------G---N-S--------------------------------------------------------------------------------------------------------------------------------------------------------------------------------------------------------------------------------------------------------------------------------------------------------------------------

>Bmor-betaREPAT7a

---------MR------A-I----------I---L-L------F--L-------I-----F---F--F-----------D---F---------------R---------------------------------------------------------------------------E------C---S-----H-------------------------------------------------------------------------------------------------------------------------------------------------------------------------------------------------------T---------FVG-T----------SVM--------R----------------------------------------P--LVY--K-----------TNA------------------T-YS-ALTF-----R-K-RIEFMNY----TI-----P-----K-----SG-N--------------SV-IQG------------I---------L---A-----Y-D-----------------R--T-H--S-G--AT-A--NITEGGIGYNKVKIRMKS-DRG--NEINY-E-------IYI-Y---------------A--------------------------------------------------------------------------------------------------------------------------------------------------------------------------------------------------------------------------------------------------------------------------------------------------------------------------------

>Bmor-betaREPAT7b

---------MK------L-Q----------I---V-V------L--L-------L-----A---A--A-----------V---I---------------V---------------------------------------------------------------------------E------C---G-----H-------------------------------------------------------------------------------------------------------------------------------------------------------------------------------------------------------T---------FVG-T----------SVN--------R----------------------------------------P--LVY--H-----------HDV------------------Q-YS-SKMF-----R-K-RVENLHF----SL-----P-----H---VPSIFG--------------RS-IQG------------I---------L---A-----F-D-----------------K--T-Y--S-T--AS-A--NITQGGIGYNFVNLRMKS-ERG--SKIHY-D-------VYI-F---------------A--------------------------------------------------------------------------------------------------------------------------------------------------------------------------------------------------------------------------------------------------------------------------------------------------------------------------------

>Bmor-betaREPAT7c

---------MK------L-Q----------I---I-I------L--V-------L-----A---A--V-----------V---I---------------V---------------------------------------------------------------------------E------C---G-----H-------------------------------------------------------------------------------------------------------------------------------------------------------------------------------------------------------L---------FVG-T----------NIN--------R----------------------------------------P--MVY--H-----------HNA------------------K-YD-AKLF-----R-K-RVENLHY----VL-----P-----Q---VPSTIG--------------KS-IQG------------I---------L---A-----Y-D-----------------K--T-H--S-T--AS-A--NITQGGIWFTFVNLRMKS-ERV--NKLNY-D-------VYI-Y---------------V--------------------------------------------------------------------------------------------------------------------------------------------------------------------------------------------------------------------------------------------------------------------------------------------------------------------------------

>Bmor-betaREPAT8a1

---------MY----A-K-I-VV-------I-----S------V--A-------L-----L---Q--L----A------S---I---------------A---------------------------------------------------------------------------E------S---N-----D-------------------------------------------------------------------------------------------------------------------------------------------------------------------------------------------------------I---------RLG-I---------DTPR--------S----------------------------------------R--KIY--S-----------EIK------------------E----ADPA-----LWK-RTDDIII----NA-----P-----N--------N--------------EL-ITA------------V---------Y---I-----T-D-----------------LREY-K----D--GE-A--FIESGGVGQKSVTIGLKS-PTI-LRGYKF-E-------IEV-Y---------------A--------------------------------------------------------------------------------------------------------------------------------------------------------------S-------D--------P-----NAGYFSKG--YAPYL------------------QE-TQYA-----------------------R-------KF------------------------------------------------------------------

>Bmor-betaREPAT8a2

---------MY----A-K-I-VV-------I-----S------V--A-------L-----L---Q--L----A------S---I---------------V---------------------------------------------------------------------------E------S---N-----D-------------------------------------------------------------------------------------------------------------------------------------------------------------------------------------------------------I---------RLG-I--------IDTPR--------S----------------------------------------R--KIY--S-----------EIV------------------E----AVPG-----LSK-RRPNIKI----NA-----P-----H--------S--------------GL-ITA------------V---------Y---I-----T-D-----------------LREY-K----D--GE-A--FITSGGVGQKSVTIGLKT-PTI-LRGYKF-Q-------IEV-Y---------------A--------------------------------------------------------------------------------------------------------------------------------------------------------------T-------D--------P-----NAG----------------------------------------------------------------------------------------------------------------------------------------

>Bmor-betaREPAT8a3

---------MY----A-K-I-VV-------I-----S------V--A-------L-----L---Q--L----A------S---I---------------V---------------------------------------------------------------------------E------S---N-----D-------------------------------------------------------------------------------------------------------------------------------------------------------------------------------------------------------I---------RLG-I--------IDTPR--------S----------------------------------------R--KIY--S-----------EIV------------------E----AVPG-----LSK-RRPNIKI----NA-----P-----H--------S--------------GL-ITA------------V---------Y---I-----T-D-----------------LREY-K----D--GE-A--FITSGGVGQKSVTIGLKT-PTI-LRGYKF-Q-------IEV-Y---------------A--------------------------------------------------------------------------------------------------------------------------------------------------------------T-------D--------P-----NAG----------------------------------------------------------------------------------------------------------------------------------------

>Bmor-betaREPAT9

---------MG------L-P----------L---I-F------A--L-------F-----L---I--I-----------G---T---------------T---------------------------------------------------------------------------R------S---N-----D-------------------------------------------------------------------------------------------------------------------------------------------------------------------------------------------------------L---------MLG-Y---------PDVS--------S----------------------------------------K--LIY--T-----------KVH------------------Q----ENPA-----IWI-RSDVFTV----NC-----S-----K--------N--------------EV-INA------------I---------S---I-----L-D-----------------LRPD-K----W--GE-A--YIKAGGIGEKFVTIELDS-PSV-FRGFNF-W-------VDV-Y---------------A--------------------------------------------------------------------------------------------------------------------------------------------------------------I----Q-TN--------S--FLFNHG--------------------------------------------------------------------KK------------------------------------------------------------------

>Bmor-gammaREPAT1

---------MN-----TI-L----------V---F-A---------L-------V---VLA---A--------------T---N---------A---------------------------------------------------------------------------------------------------A-----I--L--------P---------------------------------------------------------------------------------------------S---------------------------------------------ST------------RA---------------------N---L---------IVG-Y----------VST--------G----------------------------------------D--RLL--Y-----------RSY------------------VYQP-ARPN-----T-I-QYQDIVY----RG-----N-----A--------T--------------TR-ITA------------I---------Q---A-----T-E-----------------V--G-A--TQY--AS-A--WILSGGLSQNNVTVRFQS-ARG--YGYYY-L-------FDV-W---------------G--------------------------------------------------------------------------------------------------------------------------------------------------------------R-----------------------------------------------------------------------------------------------------------------------------------------------------------------

>Bmor-gammaREPAT10

---------MK-----LC-V----------V---F-A------F--A-------A-----V---A--V-----------V---A---------A---------------------------------------------------------------------------------------------------S-----G-----------A----------------------------------------------------------------------D--------------------VKS---------------------------------------------GS------------RS---------------------N---L---------SVG-F----------IGP--------Q----------------------------------------D--RLL--R-CNVRTVF---SAR-------DSLVWQPTRRGFTFM-V------------------------S------------------------LH------PP------A------------P---------V---L-----T-Q-----------------S--G-L--RKC--AT-V--QISRGGVGFEFVSFEFRN-APG--RGFTY-D-------LEI-W---------------G--------------------------------------------------------------------------------------------------------------------------------------------------------------R-----------------------------------------------------------------------------------------------------------------------------------------------------------------

>Bmor-gammaREPAT11

---------MK-----LC-V----------V---F-A------F--A-------A-----V---A--V-----------V---A---------A---------------------------------------------------------------------------------------------------S-----G-----------A----------------------------------------------------------------------D--------------------VKS---------------------------------------------GS------------RS---------------------N---L---------SVG-F----------IGP--------Q----------------------------------------D--RLL--R----RLVS---YSR------------------FSRM-A------------------------T-----D-----P--------S--WLHFYGLFAPPGTR-INA------------V---------R---A-----T-Q-----------------V--G----SQS--PT-V--QISRGGVGFEFVSFEFRN-APG--RGFTY-D-------LEI-W---------------G--------------------------------------------------------------------------------------------------------------------------------------------------------------R-----------------------------------------------------------------------------------------------------------------------------------------------------------------

>Bmor-gammaREPAT12

---------MK-----ST-L----------L---L-L------L--V-------V-----V---A--A-----------V---S---------A---------------------------------------------------------------------------------------------------V-----E--K--------E----------------------------------------------------------------------D--------------------FDS---------------------------------------------FR------------RS---------------------T---L---------NLG-V----------IGN--------Q----------------------------------------D--RLL--R-----------SEN------------------FVRA-S------------------------M-----V-----S--------S--WAHPLTLFAPTGTR-INA------------I---------R---A-----R-E-----------------V--G-A--SQS--GS-I--RLTHGGLGSTFAAFDITN-AAG--RGLNY-R-------VEI-W---------------G--------------------------------------------------------------------------------------------------------------------------------------------------------------R-----------------------------------------------------------------------------------------------------------------------------------------------------------------

>Bmor-gammaREPAT13

---------MK-----FT-L----------L---F-V------L--V-------I-----V---T--A-----------V---A---------A---------------------------------------------------------------------------------------------------D-----E--E--------A----------------------------------------------------------------------G--------------------VRS---------------------------------------------NG------------RP---------------------D---L---------VIG-T----------ITS--------Q----------------------------------------D--SLI--R-----------SFR------------------FNRP-A------------------------S-----P-----S--------S--QSHFHRIAMAAGVR-ISA------------I---------T---A-----R-E-----------------V--G-Q--TQN--PT-I--RVAGGGIGFNNVNLEARN-ARG--RGFSY-N-------VQI-W---------------G--------------------------------------------------------------------------------------------------------------------------------------------------------------R-----------------------------------------------------------------------------------------------------------------------------------------------------------------

>Bmor-gammaREPAT2

---------MK-----FA-L----------L---F-V------L--T-------A-----I---V--A-----------V---S---------T---------------------------------------------------------------------------------------------------N-----E--V--------A----------------------------------------------------------------------D--------------------PRS---------------------------------------------TD------------RS---------------------N--AL---------SIG-S----------ITS--------S----------------------------------------D--RLL--R-----------SFV------------------VSRA-A------------------------T-----T-----N--------S--RAVNVRYTAPAGRR-VRA------------V---------R---V-----R----------------------G-S--TQF--AS-V--RRTGGGVGFTFVNLQFVN-APR--RGFSF-T-------VQV-W---------------G--------------------------------------------------------------------------------------------------------------------------------------------------------------R-----------------------------------------------------------------------------------------------------------------------------------------------------------------

>Bmor-gammaREPAT3

---------MK-----FA-L----------L---F-V------L--T-------A-----I---V--A-----------V---S---------T---------------------------------------------------------------------------------------------------N-----E--V--------A----------------------------------------------------------------------D--------------------PRS---------------------------------------------TD------------RA---------------------N--AL---------SIG-S----------ITS--------S----------------------------------------D--RLL--R-----------SFV------------------VSRA-A------------------------T-----I-----N--------S--RVVNVRFTAPAGRR-VRA------------V---------R---V-----R----------------------G-S--TQF--AR-V--RRTGGGVGFTFVNLQFVN-APR--RGFRF-T-------VQI-W---------------G--------------------------------------------------------------------------------------------------------------------------------------------------------------R-----------------------------------------------------------------------------------------------------------------------------------------------------------------

>Bmor-gammaREPAT4

---------MK-----FA-L----------L---F-V------L--T-------A-----I---V--A-----------V---S---------T---------------------------------------------------------------------------------------------------N-----E--V--------A----------------------------------------------------------------------D--------------------PRS---------------------------------------------TD------------RA---------------------N--AL---------SIG-S----------ITS--------S----------------------------------------D--RLL--R-----------SFV------------------VSRA-A------------------------T-----I-----N--------W--RVLNVRFTAPAGRR-VRA------------V---------R---V-----R----------------------G-S--TQF--TR-V--RRTGGGVGFTFVNLQFVN-APR--RAFFF-T-------VQI-W---------------G--------------------------------------------------------------------------------------------------------------------------------------------------------------R-----------------------------------------------------------------------------------------------------------------------------------------------------------------

>Bmor-gammaREPAT5

---------MK-----FA-L----------L---F-V------L--T-------A-----V---V--A-----------V---S---------T---------------------------------------------------------------------------------------------------D-----E--V--------A----------------------------------------------------------------------D--------------------PSS---------------------------------------------TS------------RA---------------------N--GL---------AIG-S----------IAP--------N----------------------------------------D--RLL--R-----------SFV------------------VTRA-A------------------------T-----A-----S--------S--QAVNVRYTAPAGRR-VRA------------V---------R---A-----R-E-----------------V--G-V--SQF--AT-V--RLVGGGVGSTFVTLQFRN-SAR--RGYHF-N-------VQI-W---------------G--------------------------------------------------------------------------------------------------------------------------------------------------------------R-----------------------------------------------------------------------------------------------------------------------------------------------------------------

>Bmor-gammaREPAT6

---------MK-----FA-L----------L---F-V------L--T-------A-----V---V--A-----------V---S---------T---------------------------------------------------------------------------------------------------D-----E--V--------A----------------------------------------------------------------------D--------------------PSS---------------------------------------------TN------------RA---------------------N--GL---------SIG-S----------IAP--------N----------------------------------------D--RLL--R-----------SFV------------------VTRA-A------------------------T-----A-----S--------S--QAVNVRYTAPAGRR-VRA------------V---------R---A-----R-E-----------------V--G-V--SQF--AT-V--RLVGGGVGSTFVTLQFRN-SAR--RGYHF-N-------VQI-W---------------G--------------------------------------------------------------------------------------------------------------------------------------------------------------R-----------------------------------------------------------------------------------------------------------------------------------------------------------------

>Bmor-gammaREPAT7

---------MK-----FA-L----------L---F-V------L--T-------A-----V---V--A-----------V---S---------T---------------------------------------------------------------------------------------------------D-----E--V--------A----------------------------------------------------------------------D--------------------PSS---------------------------------------------TS------------RA---------------------N--GL---------AIG-S----------IAP--------N----------------------------------------D--RLL--R-----------SFV------------------VTRA-A------------------------T-----A-----S--------S--QAVNVRYTAPAGRR-VRA------------V---------R---A-----R-E-----------------V--G-V--SQF--AT-V--RLVGGGVGSTFVTLQFRN-SAR--RGYHF-N-------VQI-W---------------G--------------------------------------------------------------------------------------------------------------------------------------------------------------R-----------------------------------------------------------------------------------------------------------------------------------------------------------------

>Bmor-gammaREPAT8

---------MK-----FA-L----------L---F-V------L--T-------A-----V---V--A-----------V---S---------T---------------------------------------------------------------------------------------------------N-----E--V--------A----------------------------------------------------------------------D--------------------PSS---------------------------------------------TN------------RA---------------------L--VL---------SIG-T----------IAS--------N----------------------------------------D--RLL--R-----------SFS------------------VTRG-V------------------------T-----A-----T--------S--QIVDLRYTAPNSMR-IRA------------I---------R---V-----R-Q-----------------V--G-A--SQF--AT-V--RVVGGGVGWNTVTLRFIN-SPR--RGFNF-N-------VQI-W---------------G--------------------------------------------------------------------------------------------------------------------------------------------------------------C-------------------------------------------------------------------------------------------------FG--------------------------------------------------------------

>Bmor-gammaREPAT9

---------MK-----L--I----------L---F-V------L--A-------V-----T---A--V-----------V---S---------A---------------------------------------------------------------------------------------------------N-----E--E--------A----------------------------------------------------------------------N--------------------VKS---------------------------------------------SR------------RS---------------------E---L---------NLG-F----------ISF--------Q----------------------------------------D--RLL-LR----RYV--------------------------TRT-S------------------------S-----S-----T--------S--WSYFFWHQEPFGTR-ITA------------I---------R---A-----R-E-----------------S--G-I--SQS--PT-V--RVFSGGVGFNTVNIEFRN-APG--RGYRY-Q-------LEI-W---------------G--------------------------------------------------------------------------------------------------------------------------------------------------------------R-----------------------------------------------------------------------------------------------------------------------------------------------------------------

>Dmel_CG13323

---------MR------L-L----------L---I-L------S--V-------V-----L---A--V-----I-----L---G---------------C---------------------------------------------------------------------------H------A---Y-----G-------------------------------------------------------------------------------------------------------------------------------------------------------------------------------------------------------A---------TWG-R----------RNN--------N----------------------------------------D--YLL--S-----------RTT------------------EVRN-PIKN-----N-Y-WNVNVNY-----P-----A-----G--------F--------------YN-ISA------------V---------I---V-----Y-DNF---------------K--N-N--S-G--AS-P--SLYSGGPGYRFATVNLRG-QVN--RGINS-T-------VEI-W---------------G--------------------------------------------------------------------------------------------------------------------------------------------------------------R-----------------------------------------------------------------------------------------------------------------------------------------------------------------

>Dmel_CG13324

---------MR------L-L----------L---I-L------S--V-------V-----L---A--V-----I-----L---G---------------C---------------------------------------------------------------------------H------A---Y-----S-------------------------------------------------------------------------------------------------------------------------------------------------------------------------------------------------------A---------TWG-R----------RVN--------N----------------------------------------D--FLL--S-----------RTR------------------EVRN-PIKN-----N-Y-WNVNVNY-----P-----N-----G--------F--------------YN-ISA------------V---------I---V-----Y-DNF---------------K--N-N--S-G--AS-P--SLYSGGPGYRFATVNLRG-QVN--RGIDS-T-------VEI-W---------------G--------------------------------------------------------------------------------------------------------------------------------------------------------------R-----------------------------------------------------------------------------------------------------------------------------------------------------------------

>Harm-gammaREPAT1

---------MK-----SF-I----------L---I-A------V--L------ST-----L---S--V------CL-G-A---A---------------V---------------------------------------------------------------------------QR-----S---D---------------------------------------------------------------------------------------------------------------------------------------------------------------------------------------------------------------------------I----------PVD--------IGV--------A-KP-----------------------G--D--WVL--V-----------QAE------------------TRAN-PLEN-----Y-I-TSITVRT----RL-----D-----P--------R--------------YQ-VNM------------L---------R---L-----I-D-----------------R--N-P--AQG--AV-I--SNI-GGVGTNEVSFIVTS-QPH--DGFHI-T-------SQL-W---------------G--------------------------------------------------------------------------------------------------EDLTAG--TGAVR---SLEPT-ECIMTVDNGKTIQTYYCNEMVHWS--------------Q-----------------------------------------------------------------------------------------------------------------------------------------------------------------

>Harm-gammaREPAT22

---------MK-----SC-I----------L---I-A------L--L------AA-----L---A--S------CVHA-A---A---------------V---------------------------------------------------------------------------P------T---D---------------------------------------------------------------------------------------------------------------------------------------------------------------------------------------------------------------------------V----------PTN----PPTVAGE--------V-QP-----------------------G--D--SRL--A-----------MGE------------------HMRS-GEPD-----E-V-VIMHVVY----RL-----S-----A--------S--------------HK-VNA------------V---------I---M-----T-E-----------------L--G-A--AQG--GA-P--AFV-GGLGTNEISVILVS-ARG--RGLHY-R-------VEL-W---------------G--------------------------------------------------------------------------------------------------DDAS-G--A--GS---N--PP-TVTVL----------EL---------------------------------------------------------------------------------------------------------------------------------------------------------------------------------------

>Harm-gammaREPAT2

---------MK-----NF-I----------V---I-A------L--L------ST-----L---S--V------CL-G-A---V---------------V---------------------------------------------------------------------------QN-----A---Q---------------------------------------------------------------------------------------------------------------------------------------------------------------------------------------------------------------------------F----------PIN--------VGV--------L-EP-----------------------G--D--YVM--V-----------RGG------------------NRGN-PQPN-----S-V-SQVTVRH----QL-----H-----P--------K--------------YK-VNM------------V---------K---F-----L-D-----------------Q--A-P--ETG--GF-I--SFV-GGIGTNFVTFVFTS-QRG--GGYNF-T-------YQL-W---------------G--------------------------------------------------------------------------------------------------HDIT-G--ADGLR---SLEPS-ECTMTVDDGKTVQTYYCNEMIH----------------------------------------------------------------------------------------------------------------------------------------------------------------------------------

>Harm-gammaREPAT3

---------MK-----NF-I----------V---I-A------L--L------ST-----L---S--V------CL-G-A---V---------------V---------------------------------------------------------------------------QN-----A---Q---------------------------------------------------------------------------------------------------------------------------------------------------------------------------------------------------------------------------F----------PIN--------VGV--------L-EP-----------------------G--D--WVM--I-----------SGE------------------SRAI-QHPN-----L-VINHITVRT----RL-----H-----P--------D--------------YK-VNM------------A---------K---F-----I-D-----------------R--A-P--ETG--GF-I--SFV-GGIGTNYVTFIVSA-QRG--GGFHF-S-------YQL-W---------------G--------------------------------------------------------------------------------------------------HDTT-G--AEIRH---SLESI-ECTMTVDDGKTVKTYNCNEFVH----------------------------------------------------------------------------------------------------------------------------------------------------------------------------------

>Harm-gammaREPAT4

---------MK-----SF-I----------L---I-T---------L------AA-----L---S--V------CL-S-V---A---------------D---------------------------------------------------------------------------QE-----S---H---------------------------------------------------------------------------------------------------------------------------------------------------------------------------------------------------------------------------T----------S-S--------VGK--------Q-QP-----------------------G--D--WLM--F-----------RFE------------------NTAY-PIPD-----S-R-RRHTVQV----QL-----P-----S--------S--------------AQ-VNY------------V---------T---F-----E-D-----------------K--D-P--SSG--GF-V--SYI-GGFANNHITVMIQS-QWG--GGFDF-V-------TDI-W---------------G--------------------------------------------------------------------------------------------------QDPE-S--SQPVKLMSSAVQT-NCTMTVDDGKTIQTFDCNMNLH----------------------------------------------------------------------------------------------------------------------------------------------------------------------------------

>Harm-gammaREPAT5

---------MK-----SC-I----------L---V-A------L--L------AA-----L---A--S------CVHA-A---A---------------V---------------------------------------------------------------------------P--------------------------------------------------------------------------------------------------------------------------------------------------------------------------------------------------------------------------------------------------TN----APTVVGA--------V-QP-----------------------G--D--WLM--T-----------LGT------------------TTSV-AQPD-----Y-V-ITKRLRY----TL-----G-----S--------M--------------YQ-VNA------------V---------I---I-----T-E-----------------Q--G-V--PQG--GA-P--AFV-GGLGTNDITVVLVS-ARG--RGLHY-R-------IEL-W---------------G--------------------------------------------------------------------------------------------------YDTT-G--ADLGP---D--RP-QITIA----------EF---------------------------------------------------------------------------------------------------------------------------------------------------------------------------------------

>Harm-gammaREPAT6

---------MK-----TL-V----------L---F-A------L--L------AV-----V---P--V------CL-G-G---A---------------L---------------------------------------------------------------------------P------S---N---------------------------------------------------------------------------------------------------------------------------------------------------------------------------------------------------------------------------I----------PTT--------VGV--------A-EP-----------------------G--D--REF--F-----------RTV------------------ARAG-ARPG-----H-V-NTVTITP----VI-----S-----W--------N--------------YI-ANA------------V---------S---F-----K-E-----------------Q--G-T--PQG--AY-V--SSV-TGLGTNFVRYSVTS-QSG--KGLHV-I-------ADL-W---------------G--------------------------------------------------------------------------------------------------RDVT-ADRSSASE---MLEQHFKCTMTVSDGVTEETREC---------------------------------------------------------------------------------------------------------------------------------------------------------------------------------------

>Harm-gammaREPAT7

---------MR-----SY-I----------L---V-A------L--L------AA-----L---A--S------CLDA-A---V---------------L---------------------------------------------------------------------------P------T------------------------------------------------------------------------------------------------------------------------------------------------------------------------------------------------------------------------------------------PIN----APTVAGE--------V-QL-----------------------G--D--WVV--L-----------RAM------------------FRTS-GLPN-----E-S-TTRNLRY----SM-----S-----E--------K--------------YQ-VNA------------V---------S---I-----V-E-----------------V--G-P--PQG--GA-P--AFV-GGLGTNVIDVVLVS-ARG--RGFEY-N-------IEL-W---------------G--------------------------------------------------------------------------------------------------HDTT-G--ADPGP---N--PP-DVLIT----------EL---------------------------------------------------------------------------------------------------------------------------------------------------------------------------------------

>Harm-gammaREPAT8

---------MK-----SY-I----------L---V-A------L--L------AA-----L---A--S------CLDA-A---V---------------L---------------------------------------------------------------------------P------T------------------------------------------------------------------------------------------------------------------------------------------------------------------------------------------------------------------------------------------PIN----APTVAGE--------V-QP-----------------------G--D--WVL--L-----------RGW------------------IRTAPSTSN-----Q-A-LDRRLRY----TL-----S-----E--------R--------------YQ-VNA------------V---------S---I-----V-E-----------------V--G-P--PQG--GA-P--AFV-GGLGTNVIDVVLVA-ARG--RGFEY-N-------IEL-W---------------G--------------------------------------------------------------------------------------------------HDTT-G--AYPGP---N--PP-DVLIT----------EL---------------------------------------------------------------------------------------------------------------------------------------------------------------------------------------

>Harm-gammaREPAT23

---------MK-----SC-V----------L---I-A------L--L------AA-----L---A--S------CVHA-A---A---------------V---------------------------------------------------------------------------P--------------------------------------------------------------------------------------------------------------------------------------------------------------------------------------------------------------------------------------------------TN----APTVVGA--------V-QP-----------------------G--D--WLM--G-----------IGT------------------TNSA-AQPD-----Y-V-ITKRLRY----TL-----G-----S--------M--------------YR-VNA------------V---------I---I-----R-E-----------------L--G-A--AQG--GA-P--AFV-GGLGSNDVTVVLVS-ARG--RGLHY-R-------VEL-W---------------G--------------------------------------------------------------------------------------------------HDTS-G--ADLGP---D--RP-LITIA----------EL---------------------------------------------------------------------------------------------------------------------------------------------------------------------------------------

>Harm-betaREPAT1

---------MS------V-I----------S---H-V------L--L-------L-----L------V-----------L---C---------------V---------------------------------------------------------------------------A------L---Y--KAED-------------------------------------------------------------------------------------------------------------------------------------------------------------------------------------------------------L---------LLG-S----------SAN--------A----------------------------------------Q--LGY--Q-----------ETV------------------K-LG-SVPL-----K-T-RTKNVFY----NS-----N-----N--------T--------------RI-IKA------------I---------S---A-----I-D-----------------I--D-K--S-K--TK-P--AVTAGGVGSSFVNVRFKS-EKG--VGLNY-L-------VQI-F---------------V--------------------------------------------------------------------------------------------------------------------------------------------------------------------------------------------------------------------------------------------------------------------------------------------------------------------------------

>Harm-betaREPAT10

MA-------SK------L-L----------I---T-A------L--L-------I-----F---A--A-----------A---L---------------V-SAQ-----------------------------------------------------------------------A------R---G-----N-------------------------------------------------------------------------------------------------------------------------------------------------------------------------------------------------------F---------FVG-N----------TFG--------A----------------------------------------H--LIY--H-----------ETH------------------S-KY-AIPF-----I-V-RDEDFTV----TG-----V-----D--------N--------------EI-IEA------------I---------S---V-----S-D-----------------L--E-G----K--GL-S--YVKSGGIGQHNVTIHLES-QRS--HGYKF-L-------VEV-Y---------------A--------------------------------------------------------------------------------------------------------------------------------------------------------------I-----------------------------------------------------------------------------------------------------------------------------------------------------------------

>Harm-betaREPAT11a

----MFKI--------------------------L-L---------L-------C-----A---V--I-----------S---T---------------V---------------------------------------------------------------------------C------A---K-----S-------------------------------------------------------------------------------------------------------------------------------------------------------------------------------------------------------V---------IVG-Q----------TWEQNGT---------------------------------------------R--KVY--E-----------TVV------------------S-AN-SIPF-----L-K-REKFVNF----EL-----S-----N--------G-D-------------K-IKG------------I---------A---I-----K-D-----------------M--D-N--A-L--AE-P--SITQGGIGFGFANIKLKS-ERG--SGFNF-Q-------IEI-Y---------------G--------------------------------------------------------------------------------------------------------------------------------------------------------------------------------------------------------------------------------------------------------------------------------------------------------------------------------

>Harm-betaREPAT11b

----MLKE--------------------------F-I------L--V-------V-----T---L--L-----------A---L---------------V---------------------------------------------------------------------------Q------S---S-----S-------------------------------------------------------------------------------------------------------------------------------------------------------------------------------------------------------V---------YVG-H----------VYDQTGL----V----------------------------------------R--KVY--D-----------KMI------------------E-ST-AIPL-----I-K-REEEVYF----EY-----P-----A--------G-D------------QK-IKG------------I---------A---I-----K-D-----------------L--H-D--G-K--AE-A--SITRGGLGFGFVNIKLKS-ERG--SGLKY-L-------IEI-F---------------A--------------------------------------------------------------------------------------------------------------------------------------------------------------------------------------------------------------------------------------------------------------------------------------------------------------------------------

>Harm-betaREPAT12

---------MR------T-S-------SI-A---I-L------L--C-------L-----I---G--A-----------A---C---------------S---------------------------------------------------------------------------Q------S---H-----D-------------------------------------------------------------------------------------------------------------------------------------------------------------------------------------------------------L---------VIG-Q----------AQY--------G----------------------------------------D--VIV--Y-----------KAD------------------EVRY-GFPF-----I-V-RQSILEY-----P-EQGQK-----N--------Y--------------AY-IKA------------I---------Y---I-----K-DND---------------R--D-G--T-G--GY-P--SISAGGVGQRFVKIKLKS-QRG--SGLNF-T-------VTI-Y---------------G--------------------------------------------------------------------------------------------------------------------------------------------------------------R-----------------------------------------------------------------------------------------------YV----------------------------------------------------------------

>Harm-betaREPAT13

----------M------K-I----------I---L-A------L--F-------G-----F------A-----------A---V---------------A---------------------------------------------------------------------------S------A---A-----D-------------------------------------------------------------------------------------------------------------------------------------------------------------------------------------------------------L---------TVG-N---------E--G--------G----------------------------------------S--LIF--E-----------QNV------------------T----ASPT-----IWK-QTRNLTV----NA-----T-----D--------D--------------AV-ISR------------V---------V---V-----I-D-----------------NRPE-K----D--GE-A--KVVEGGEGEKNVTIELKG-PAV-FRGFDF-T-------IQV-F---------------A--------------------------------------------------------------------------------------------------------------------------------------------------------------A----P-EN--------E-----NTQH--------PKATG----------------HD-TQQT-----------------------Q---------------------------------------------------------------------------

>Harm-betaREPAT15

---------MRYL----I-I----------L-----T------V--C-------L-----F---V-A--------------------------------V---------------------------------------------------------------------------Y------A---D-----H-------------------------------------------------------------------------------------------------------------------------------------------------------------------------------------------------------L---------IVGNV----------SNR--------V----------------------------------------V--LAN--Q-----------TKV------------------E-YN-AIPF-----M-K-RVKYFFY----SQ-----T-----D--------N--------------RL-IQG------------I---------Q---A-----L-D-----------------T--L-H--S-K--AS-I--NITAGGVGYSYVNLRLKS-ERG--RGLSY-D-------IGI-Y-VQDYFYH----------------------------------------------------------------------------------------------------------------------------------------------------------------------------------------------------------------------------------------------------------------------------------------------------------------------------------------

>Harm-betaREPAT16

---------MK------S-T---LILC---V---L-L------T--T-------L-----A---S--V---------------L---------------S---------------------------------------------------------------------------S------TYFFG-----K---------------------------------------------------------------------------------------------------------------------------------------------------------------------------------------------------------------------E----------TKD--------T----------------------------------------K--IIH--R-----------DKL------------------R-YE-AIPL-----K-K-RIKTYTY----NG-----T-----A------------------------P-IKA------------I---------S---C-----Y-D-----------------F--Q-N--S-E--AS-V--NITSGGIGYNYVELRLKS-QRS--YRLDF-A-------IEI-Y---------------A--------------------------------------------------------------------------------------------------------------------------------------------------------------------------------------------------------------------------------------------------------------------------------------------------------------------------------

>Harm-betaREPAT17

---------MD------L-L----------L---I-I------L--F-------V-----I---T--A-----------D---A------RMF------C---------------------------------------------------------------------------E------K---T-----K-------------------------------------------------------------------------------------------------------------------------------------------------------------------------------------------------------A---------PL-------------------------------------------------------------------------------------ELG------------------QWQY-SLKH-----M-D--SAVVEN-----P-----A-----V--------S--------------IN-EGQ------------V---------Y---V-----Y-ENYFPGYTIRYIHVDNLAI--R-T--C-G--AS-A--SIKAGGVGASSVLIVLRA-YIN--QEIRS-V-------IDI-W---------------G-------EKK-----VSNNT--N---ALN--NTQLSLNP----EEVKNINYNYLFKDTRALNHNR------QNLH-----------------------------------------------------------------------------------------------------------------------------------------------------------------------------------------------------------------------------------------------------

>Harm-betaREPAT2

---------MK-C-V--I-V----------L---S-V------F--V-------L-----I---A--I-----------S---S---------------C---------------------------------------------------------------------------L------A---N-----D-------------------------------------------------------------------------------------------------------------------------------------------------------------------------------------------------------L---------IIG-T----------SYN--------T----------------------------------------R--LVW--Q-----------QKA------------------D-YM-GFPF-----K-K-RVKEVFY----SD-----P-----G--------Q--------------QI-IKG------------V---------I---A-----R-D-----------------I--D-H--S-A--GS-A--TVTAGGVGFSFVNIRLKS-ERG--SALNY-Q-------IEI-Y---------------V--------------------------------------------------------------------------------------------------------------------------------------------------------------------------------------------------------------------------------------------------------------------------------------------------------------------------------

>Harm-betaREPAT3

---------MK-T-A--P-V----------L---L-V------V--L-------L-----S---I--I-----------C---V---------------V---------------------------------------------------------------------------Q------C---K-----D-------------------------------------------------------------------------------------------------------------------------------------------------------------------------------------------------------L---------IVG-S----------SFN--------N----------------------------------------R--LIH--Q-----------QKA------------------E-YN-AIPL-----K-K-RVKEVFF----SD-----P-----G--------Q--------------QI-IKG------------I---------I---L-----R-D-----------------L--D-H--T-E--AI-P--SVTAGGVGFSYANIRLKS-ERG--SGLNY-Q-------VEI-Y---------------V--------------------------------------------------------------------------------------------------------------------------------------------------------------------------------------------------------------------------------------------------------------------------------------------------------------------------------

>Harm-betaREPAT4

---------MR------F-V----------L---S-L------L--I-------L-----S---C--L-----------L---C---------------V---------------------------------------------------------------------------F----VLC---S-----D-------------------------------------------------------------------------------------------------------------------------------------------------------------------------------------------------------M---------IIG-D----------TVH--------R----------------------------------------K--MVF--H-----------QRV------------------K-DF-AIPF-----K-K-RIQTLTY----TD-----P-----E--------K--------------RT-IKG------------V---------A---A-----I-D-----------------N--D-F--S-H--AS-A--NITDGGVGYSYVTVRMKS-QRH--HPLNF-E-------VEI-Y---------------V--------------------------------------------------------------------------------------------------------------------------------------------------------------------------------------------------------------------------------------------------------------------------------------------------------------------------------

>Harm-betaREPAT5

---------MK-IQ---V-L----------V---F-V------V--C-------L-----F---V--A-----------A---T---------------C--------------------------------------------------------------------------------------D-----H-------------------------------------------------------------------------------------------------------------------------------------------------------------------------------------------------------L---------ILGNT----------NNN--------Q----------------------------------------N--MIH--H-----------TTA------------------H-YT-AIPF-----I-K-RVKNVFY----SG--------------------H--------------SV-INS------------I---------M---A-----Y-D-----------------N--K-H--T-N--AT-A--AVTAGGIGYTYVNLRLKS-ERG--KELDY-D-------IGI-Y---------------A--------------------------------------------------------------------------------------------------------------------------------------------------------------------------------------------------------------------------------------------------------------------------------------------------------------------------------

>Harm-betaREPAT6

---------MK------C-V----------I-----------------------------I---F--VCVFL-------L---C---------------V---------------------------------------------------------------------------C------L---N--EGKD-------------------------------------------------------------------------------------------------------------------------------------------------------------------------------------------------------F---------TVG-T----------RAN--------N----------------------------------------L--LIS--T-----------EKV------------------K-YR-SLPL-----I-R-RDKDYTY----ID-----P-----K--------E--------------RI-IKG------------I---------I---A-----R-D-----------------L--S-R--T-D--TE-V--TITSGGIGATNVTLHLQS-GRG--EELNY-L-------ILI-F---------------S---NIK--------------------------------------------------------------------------------------------------------------------------------------------------------------------------------------------------------------------------------------------------------------------------------------------------------------------------

>Harm-betaREPAT7a

---------MK------V-L----------S---I-V------F--L-------L-----A---A--V-----------V---LA--------------V---------------------------------------------------------------------------E------C---S-----H-------------------------------------------------------------------------------------------------------------------------------------------------------------------------------------------------------T---------FLG-T----------NVL--------R----------------------------------------P--LVY--H-----------HSV------------------Q-YS-SKVF-----R-K-RVENMYF----AL-----P-----A---VPSNAG--------------RA-IQG------------I---------L---A-----F-D-----------------M--T-H--T-G--AS-A--NVTQGGLGFNFVNLRLKS-DRG--EKINF-D-------VYI-Y---------------A--------------------------------------------------------------------------------------------------------------------------------------------------------------------------------------------------------------------------------------------------------------------------------------------------------------------------------

>Harm-betaREPAT7b

---------MK------I-Y----------T---L-V------L--L-------L-----A---A--V-----------V---F---------------V---------------------------------------------------------------------------E------C---G-----H-------------------------------------------------------------------------------------------------------------------------------------------------------------------------------------------------------T---------FMG-T----------SFL--------R----------------------------------------P--QVY--H-----------KEV------------------R-FP-SRFF-----Q-K-RVEYLNY----TL-----P-----AYRGYPGAPG--------------GS-IQG------------I---------L---A-----F-D-----------------M--T-H--S-G--AS-A--NVTRGGIGHNFVDIRMKS-DRN--KELHY-D-------VYI-Y---------------A--------------------------------------------------------------------------------------------------------------------------------------------------------------------------------------------------------------------------------------------------------------------------------------------------------------------------------

>Harm-betaREPAT8a

---------MD----F-KFL----------I---L-L------L--S-------L-----L---F--L-----------T---S---------------V---------------------------------------------------------------------------K------S---N-----D-------------------------------------------------------------------------------------------------------------------------------------------------------------------------------------------------------I---------TLG-F---------ASPY--------A----------------------------------------K--KIY--S-----------AVK------------------I----ANPA-----LWR-REDDVMV----SS-----P-----N--------H--------------EL-ISA------------V---------Y---I-----T-D-----------------LRPD-K----E--GE-A--YIEAGGLGSKTVTIALKS-PSI-LRGYKF-A-------IEV-Y---------------A--------------------------------------------------------------------------------------------------------------------------------------------------------------Q----S-VQPGFGQSYPS-----NPGYVRGG--LSGYA---------------------GTYASTGYPTG----SIYP--HVH---Y-------KK------------------------------------------------------------------

>Harm-betaREPAT8b

---------MY----F-K-L----------L-----T------V--L-------V-----L---Y--L-----------S---L---------------A---------------------------------------------------------------------------E------S---N-----D-------------------------------------------------------------------------------------------------------------------------------------------------------------------------------------------------------I---------RIG-H---------TTYR--------S----------------------------------------K--KIY--S-----------EIK------------------E----ANPA-----LWT-RTDDIII----NA-----P-----H--------N--------------EV-IDA------------I---------Y---I-----T-D-----------------LREN-K----D--GK-A--SIVSGGVGMKSVTITLKS-PAI-LRGYKF-E-------IEV-F---------------S--------------------------------------------------------------------------------------------------------------------------------------------------------------D-------N--------P-----NQGYY-KD--SHYFG------------------AD-TQYA-----------------------R-------KH------------------------------------------------------------------

>Harm-betaREPAT9

---------MI------Q-I----------I---F-F------L--N-------V-----I---T--L-----------I---T---------------V---------------------------------------------------------------------------S------C---H-----D-------------------------------------------------------------------------------------------------------------------------------------------------------------------------------------------------------V---------FLG-Y---------ADLK--------S----------------------------------------R--LIY--S-----------KGH------------------E----QSPA-----VWM-RSDVFTV----NC-----S-----R--------N--------------EV-ISA------------I---------Q---I-----I-D-----------------LRID-K----L--GE-V--NIRDGGIGAKFVTIELYS-PGT-YLGLNF-W-------VEV-F---------------A--------------------------------------------------------------------------------------------------------------------------------------------------------------L----K-TS--------Y--FLYNHG--------------------------------------------------------------------K-------------------------------------------------------------------

>Harm-gammaREPAT9

---------MK-----SL-I----------L---V-A------V--L-------A---ALA---V--C-----------S---D---------A---------------------------------------------------------------------------------------------------A-----A-LQ-------QN---------------------------------------------------------------------------------------------P---------------------------------------------AF------------RA---------------------N---M---------YQG-A----------IRP--------G----------------------------------------D--RLL--Y-----------RNY------------------YYKA-PIAN-----A-V-QYQDITY----RG-----S-----S--------S--------------TR-ISF------------I---------Q---A-----V-E-----------------V--G-Q--TQW--GQ-P--SLRSGGVNFSNATIRLTS-ARG--WGYYY-M-------IEI-W---------------G--------------------------------------------------------------------------------------------------------------------------------------------------------------R-----------------------------------------------------------------------------------------------------------------------------------------------------------------

>Harm-gammaREPAT10

---------MK-----SA-V----------V---L-A------V--L-------A---ALA---V--S-----------S---A---------A---------------------------------------------------------------------------------------------------A-----A--A--------E---------------------------------------------------------------------------------------------R---------------------------------------------AA------------RV---------------------D---L---------NHG-Y----------ARP--------D----------------------------------------D--ILL--Y-----------SNT------------------VLRA-PVAN-----E-V-QSEDIAY----SG--------------------S--------------AR-ITA------------I---------R---A-----T-E-----------------V--G-Q--TQW--AI-P--SVRSGGVGRNFATVRLTS-ARG--FGYHY-N-------IEI-W---------------G--------------------------------------------------------------------------------------------------------------------------------------------------------------R-----------------------------------------------------------------------------------------------------------------------------------------------------------------

>Harm-gammaREPAT11

---------MK-----SA-V----------V---F-A------V--L-------A---ALA---V--S-----------S---A---------A---------------------------------------------------------------------------------------------------A-----A--A--------E---------------------------------------------------------------------------------------------R---------------------------------------------AA------------RV---------------------D---L---------NHG-Y----------ARP--------D----------------------------------------D--ILL--Y-----------SNT------------------VLRA-PVAN-----E-V-QSEDIAY----SG--------------------S--------------TR-ITA------------I---------R---A-----T-E-----------------V--G-Q--TQW--AI-P--SVRSGGVGRNFATVRLTS-ARG--FGYHY-S-------IEI-W---------------G--------------------------------------------------------------------------------------------------------------------------------------------------------------R-----------------------------------------------------------------------------------------------------------------------------------------------------------------

>Harm-gammaREPAT12

---------MK-----TL-S----------L---I-V------V--L-------A---ALA---V--C-----------T---Q---------S--------------------------------------------------------------------------------------------------------------------------------------------------------------------------------------------------------------------------------------------------------------------------------A---------------------N---L---------SQG-H----------PSA--------F----------------------------------------C--RLL--Y-----------RNY------------------YRRN-AIPN-----A-V-QYQDVTY----RG-----N-----S--------T--------------VR-ISS------------I---------R---A-----T-E-----------------V--G-G--TQF--GV-P--SIRAGGVNRNNVTIRLTS-ARG--RG-----------------------------------------------------------------------------------------------------------------------------------------------------------------------------------------------K---------------------------------------------------------------------------------------------------KRQQ----------------------------------------------------------

>Harm-gammaREPAT13

---------MK-----TL-S----------L---I-V------V--L-------A---ALA---V--C-----------T---Q---------S--------------------------------------------------------------------------------------------------------------------------------------------------------------------------------------------------------------------------------------------------------------------------------A---------------------N---L---------SQG-H----------PSA--------F----------------------------------------C--RLL--Y-----------RNY------------------YRRN-AIPN-----A-V-QYQDVTY----RST----N-----S--------T--------------VR-ISS------------I---------R---A-----T-E-----------------V--G-G--TQF--AS-P--SIRAGGVNRNNVTIRLTS-ARG--RGYYY-M-------VEI-W---------------G--------------------------------------------------------------------------------------------------------------------------------------------------------------R-----------------------------------------------------------------------------------------------------------------------------------------------------------------

>Harm-gammaREPAT14

---------MK-----TL-S----------L---I-V------V--L-------A---ALA---V--C-----------T---Q---------S--------------------------------------------------------------------------------------------------------------------------------------------------------------------------------------------------------------------------------------------------------------------------------A---------------------N---L---------SQG-H----------PSA--------F----------------------------------------C--RLL--Y-----------RNY------------------YRRN-AIPN-----A-V-QYQDVTY----RST----N-----S--------T--------------VR-ISS------------I---------R---A-----T-E-----------------V--G-G--TQF--GV-P--SIRAGGVNRNNVTIRLTS-ARG--RGYYY-M-------VEI-W---------------G--------------------------------------------------------------------------------------------------------------------------------------------------------------R-----------------------------------------------------------------------------------------------------------------------------------------------------------------

>Harm-gammaREPAT17

---------MK-----SL-L----------V---V-M------L--L-------A-----A---A--A-----------T---A---------A---------------------------------------------------------------------------------------------------SIDSEI----------------------------------------------------------------------------------------------------------PEVVVTYEDI---ND---------------TDVADV-TT----TAVAS------------RN---------------------N---I---------NVG-V----------IGN--------S-------------------------------------------RLL--S-----------RTH------------------HVRS-GIPG-----H-R-HIQDITF----RA-----N-----N--------G--------------LR-FTA------------I---------R---V-----T-H-----------------Y--G-N--YQG--AS-A--GIRSGGVGSTVVTIRLQS-RVH--FGYQY-R-------IEL-Y---------------G--------------------------------------------------------------------------------------------------------------------------------------------------------------R-----------------------------------------------------------------------------------------------------------------------------------------------------------------

>Harm-gammaREPAT18

---------MK-----KL-L----------V---L-L------L--L-------A-----G---A--A-----------T---A---------A---------------------------------------------------------------------------------------------------CIHTDLV--Q--------A---------------------------------------------------------------------------------------------PEMIVTYEEV---DG---------------DTVTDV-PV----TAVAT------------RT---------------------N---I---------SVG-T----------IGN--------S-------------------------------------------RLI--A-----------RSV------------------HVQN-AVAK-----A-R-HVRDISF----RS-----D-----D--------P--------------IK-ISA------------I---------R---V-----T-H-----------------V--G-S--VDG--AT-S--SIKSGGVGNSFVVIRLQT-AVG--RGYHY-T-------IEI-Y---------------G--------------------------------------------------------------------------------------------------------------------------------------------------------------R-----------------------------------------------------------------------------------------------------------------------------------------------------------------

>Harm-gammaREPAT19

---------MK-----SL-L----------V---F-M------L--L-------A-----A---A--A-----------A---A---------S---------------------------------------------------------------------------------------------------YVHSDTA--E--------A---------------------------------------------------------------------------------------------PEMTVTYEEV---DG---------------STVTDE-PV----TAVAL------------RN---------------------N---I---------SVG-T----------IGN--------S-------------------------------------------RLL--A-----------RSA------------------HIRG-AVAG-----G-R-WVQDITY----RT-----N-----S--------A--------------VR-ITA------------I---------R---V-----R-H-----------------V--G-T--PQG--AT-S--RITSGGVGSTFVTIRLQS-AVG--RGYHY-T-------IEI-Y---------------G--------------------------------------------------------------------------------------------------------------------------------------------------------------R-----------------------------------------------------------------------------------------------------------------------------------------------------------------

>Harm-gammaREPAT20

---------MK-----SL-L----------V---F-M------L--L-------A-----A---A--A-----------A---A---------S---------------------------------------------------------------------------------------------------YVHSDTA--E--------A---------------------------------------------------------------------------------------------PEMTVTYEEV---DG---------------STVTDE-PV----TAVAL------------RN---------------------N---I---------SVG-T----------IGN--------S-------------------------------------------RLL--A-----------RSA------------------HIRG-AVAG-----G-R-WVQDITY----RT-----N-----S--------A--------------VR-ITA------------I---------R---V-----R-H-----------------V--G-T--PQG--AT-S--RITSGGVGSTFVTIRLQS-AVG--RGYHY-T-------IEI-Y---------------G--------------------------------------------------------------------------------------------------------------------------------------------------------------R-----------------------------------------------------------------------------------------------------------------------------------------------------------------

>Harm-gammaREPAT21

---------MK-----SL-L----------V---F-M------L--L-------A-----A---A--A-----------A---A---------S---------------------------------------------------------------------------------------------------YVHSDTA--E--------A---------------------------------------------------------------------------------------------PEMTVTYEEV---DG---------------STVTDE-PV----TAVAL------------RN---------------------N---I---------SVG-T----------IGN--------S-------------------------------------------RLL--A-----------RSA------------------HIRG-AVAG-----G-R-WVQDITY----RT-----N-----S--------A--------------VR-ITA------------I---------R---V-----R-H-----------------V--G-T--PQG--AT-S--RITSGGVGSTFVTIRLQS-AVG--RGYHY-T-------IEI-Y---------------G--------------------------------------------------------------------------------------------------------------------------------------------------------------R-----------------------------------------------------------------------------------------------------------------------------------------------------------------

>Harm-gammaREPAT15

---------MK-----SF-L----------V---I-A------L--L-------A---VTA---A--A-----------S---V---------I---------------------------------------------------------------------------------------------------T-----H--E--------A---------------------------------------------------------------------------------------------P---------------------------------------------EP------------AA---------------------D-V-L---------PEG-S--ETQELQ--VAD--------A----------------------------------------D--EVP--M-----------STH-------------------SRN-PIAN-----Q-V-QTVNVRY----TG-----S-----S--------S--------------II-ILA------------V---------R---A-------------------------Y--G-S--GQG--AS-A--RVVEGYLGRNSITIQLQS-ARG--RGFHY-R-------IEI-W---------------G--------------------------------------------------------------------------------------------------------------------------------------------------------------R-----------------------------------------------------------------------------------------------------------------------------------------------------------------

>Harm-gammaREPAT16

---------MK-----SF-L----------V---I-A------L--L-------A---VTA---A--A-----------S---V---------I---------------------------------------------------------------------------------------------------T-----H--E--------A---------------------------------------------------------------------------------------------P---------------------------------------------EP------------AA---------------------D-V-L---------PEG-S--ETQELQ--VAD--------A----------------------------------------D--EVP--I-----------STH-------------------SRN-PIAN-----V-V-QTVDVRY----TG-----S-----N--------A--------------II-VLA------------V---------R---A-------------------------Y--G-S--RQG--AS-A--RVVEGALGRNSITIRLQS-ARG--RGFQY-R-------VEI-W---------------G--------------------------------------------------------------------------------------------------------------------------------------------------------------R-----------------------------------------------------------------------------------------------------------------------------------------------------------------

>Msex-betaREPAT1

---------MY------R-T----------T---L-V------V--L-------L-----A------V-----------F---L---------------C---------------------------------------------------------------------------T-------------KGEH-------------------------------------------------------------------------------------------------------------------------------------------------------------------------------------------------------L---------QVG-T----------SMN--------N----------------------------------------E--LIY--T-----------ENV------------------K-LS-AVPL-----T-T-RTKNVFF----NG-----N-----D--------T--------------RV-IKG------------I---------W---A-----I-D-----------------M--D-N--S-K--AK-A--TVTSGGVGFTFTNLKIKS-KRG--DGLNF-Q-------VQI-F---------------A--------------------------------------------------------------------------------------------------------------------------------------------------------------------------------------------------------------------------------------------------------------------------------------------------------------------------------

>Msex-betaREPAT10

---------MK------L-L----------------V------L--L-------L-----L---F--V-----------L---A---------------T-LSD-----------------------------------------------------------------------A------R---G-----N-------------------------------------------------------------------------------------------------------------------------------------------------------------------------------------------------------I---------SIG-D----------NFG--------A----------------------------------------K--LIY--H-----------ETR------------------H-KI-GIPL-----M-R-RDEKVTV----KG-----V-----G--------S--------------ER-IKS------------I---------L---V-----Q-D-----------------L--K-G----D--GE-S--YIMSGGVGHSSVTIKLKG-PKG--KGYKF-L-------IDV-Y---------------A--------------------------------------------------------------------------------------------------------------------------------------------------------------I-----------------------------------------------------------------------------------------------------------------------------------------------------------------

>Msex-betaREPAT11a

----MYKL--------------------------L-I------L--L-------C------------V-----------T---S---------------V---------------------------------------------------------------------------Y------S---K-----S-------------------------------------------------------------------------------------------------------------------------------------------------------------------------------------------------------V---------IVG-Q----------VYDYNGV----V----------------------------------------R--KVF--E-----------KVV------------------D-SS-SIPL-----I-K-REKEIDF----EL-----P-----T--------P-Q------------QT-IKG------------I---------A---I-----K-D-----------------L--D-N--G-L--AE-P--AITRGGLGFNFVKLKIKS-ERG--SGFKY-L-------VEI-Y---------------A--------------------------------------------------------------------------------------------------------------------------------------------------------------------------------------------------------------------------------------------------------------------------------------------------------------------------------

>Msex-betaREPAT12

---------MR------A-L-------TL-T---A-L------L--C-------L-----V---A--A-----------G---A---------------A---------------------------------------------------------------------------Q------S---H-----D-------------------------------------------------------------------------------------------------------------------------------------------------------------------------------------------------------L---------YLG-H----------ILY--------R----------------------------------------D--VPL--Y-----------TIN------------------EYKY-GFPL-----I-V-RTSIIEY-----P-EAGIK-----N--------F--------------AY-ITA------------I---------V---V-----K-DHY---------------T--D-G--N-G--GF-P--AVVAGGVGQRFAKIKLKS-QRS--HGFNF-T-------VTI-F---------------G--------------------------------------------------------------------------------------------------------------------------------------------------------------R-----------------------------------------------------------------------------------------------Y-----------------------------------------------------------------

>Msex-betaREPAT13

---------MY------N-LG---------L---C-V------L--V-------A-----V------V-----------T---V---------------V---------------------------------------------------------------------------A------A---N-----D-------------------------------------------------------------------------------------------------------------------------------------------------------------------------------------------------------L---------TVG-S---------Q--G--------G----------------------------------------K--KIH--D-----------ENL------------------Q----ASPA-----IWR-QDKNVTF----NA-----T-----E--------G--------------HL-ISR------------V---------I---I-----T-D-----------------LRPQ-K----D--GE-A--KIVDGGEGHKNVTIELKS-PAV-FRGFEF-N-------VQV-Y---------------A--------------------------------------------------------------------------------------------------------------------------------------------------------------T----P-DN--------N-------QQ--------SVSVQ----------------QQ-TSTQ-----------------------T---------------------------------------------------------------------------

>Msex-betaREPAT14

--MSISKR--------------------------C-F------L--L-------L-----F---F--V-----------L---T---------------I---------------------------------------------------------------------------S------C---Y-----D-------------------------------------------------------------------------------------------------------------------------------------------------------------------------------------------------------L---------YLG-H----------DFG--------A----------------------------------------R--MSY--------------KEV------------------K-RQ-GLLF-----F-K-RQANLSF----NY-----P-----V--------G--------------TN-ITG------------I---------V---C-----L-D-----------------R--TTE--K-L--GV-A--KIIEGGIGFNFVEIQLES-ERS--LSLRY-I-------VEI-Y---------------Q----------------------------------------------------------------------------QSLNKHDSNGKQVNIVYKFYSL------------------------------------------------------------------------------------------------------------------------------------------------------------------------------------------------------------------------------

>Msex-betaREPAT15

---------MKLS----L-L----------L---V-V------T--G-------I-----S---L----------------------------------A---------------------------------------------------------------------------Y------S---Y-----H-------------------------------------------------------------------------------------------------------------------------------------------------------------------------------------------------------M---------VIGNV----------NNR--------V----------------------------------------R--LYN--E-----------TVV------------------E-YN-AIPF-----M-K-RVKYYFV----AL-----P-----G--------N--------------EK-IQG------------I---------Q---A-----L-D-----------------N--L-K--S-T--SS-I--NITAGGVGHSFANLRLKS-ERG--RGINY-N-------IGV-Y-VSDNL-L----------------------------------------------------------------------------------------------------------------------------------------------------------------------------------------------------------------------------------------------------------------------------------------------------------------------------------------

>Msex-betaREPAT17

---------MM------L-I----------V---L-L------A--S-------I-----L---A--V-----------E---S------RIF------C---------------------------------------------------------------------------E------V---N-----K-------------------------------------------------------------------------------------------------------------------------------------------------------------------------------------------------------M---------PL-------------------------------------------------------------------------------------EVG------------------MWKS-NLRH-----M-D--SAVVEN-----T-----P-----V--------S--------------IS-EGQ------------V---------Y---V-----Y-ENYFPGYIIRYIHVDNLAT--R-S--C-G--AS-A--TIRDGGIGSQTVLIVLHA-DTN--QEIRS-I-------IDI-W---------------G-------E-K--------RP--GMKMKVT--RQPL--DT----EETKNLTSLYLFKDLRAVNHNK----GF---------------------------------------------------------------------------------------------------------------------------------------------------------------------------------------------------------------------------------------------------------

>Msex-betaREPAT2a

---------MK-S-I--Q-F----------M---C-L------T--L-------L-----I---Y--I-----------S---S---------------C---------------------------------------------------------------------------M------G---R-----D-------------------------------------------------------------------------------------------------------------------------------------------------------------------------------------------------------M---------MVG-T----------SYN--------A----------------------------------------Q--LVW--H-----------QKQSTLVFLH-----------E-----------------RVKEMFY----SD-----P-----S--------Q--------------K---SG------------V---------I---A-----R-D-----------------I--D-H--S-E--SS-A--IITAGGVGFTYVNLRFKS-ERG--SSLNY-E-------VEI-Y---------------V--------------------------------------------------------------------------------------------------------------------------------------------------------------------------------------------------------------------------------------------------------------------------------------------------------------------------------

>Msex-betaREPAT2b

---------MK-S-I--Q-F----------M---C-L------T--L-------L-----I---Y--I-----------S---S---------------C---------------------------------------------------------------------------M------G---R-----D-------------------------------------------------------------------------------------------------------------------------------------------------------------------------------------------------------M---------MVG-T----------SYN--------A----------------------------------------Q--LVW--H-----------QKA------------------E-YA-GIPF-----M-K-RVKEMFY----SD-----P-----S--------Q--------------KI-IKG------------V---------I---A-----R-D-----------------I--D-H--S-E--SS-A--IITAGGVGFTYVNLRFKS-ERG--SSLNY-E-------VEI-Y---------------V--------------------------------------------------------------------------------------------------------------------------------------------------------------------------------------------------------------------------------------------------------------------------------------------------------------------------------

>Msex-betaREPAT2c

---------MK-S-I--Q-F----------M---C-L------T--L-------L-----I---Y--I-----------S---S---------------C---------------------------------------------------------------------------M------G---R-----D-------------------------------------------------------------------------------------------------------------------------------------------------------------------------------------------------------M---------MVG-T----------SYN--------A----------------------------------------Q--LVW--H-----------QKA------------------E-YA-GIPF-----M-K-RVKEMFY----SD-----P-----S--------Q--------------KI-IKG------------V---------I---A-----R-D-----------------I--D-H--S-E--SS-A--IITAGGVGFTYVNLRFKS-ERG--SSLNY-E-------VEI-Y---------------V--------------------------------------------------------------------------------------------------------------------------------------------------------------------------------------------------------------------------------------------------------------------------------------------------------------------------------

>Msex-betaREPAT3

---------MK-Y-S--I-I----------L---F-V------V--L-------L-----A---I--I-----------C---G---------------V---------------------------------------------------------------------------Q------C---K-----D-------------------------------------------------------------------------------------------------------------------------------------------------------------------------------------------------------L---------IVG-T----------SFN--------N----------------------------------------R--LLF--Q-----------EKA------------------E-YN-AIPL-----K-K-RVKEVFF----SD-----P-----G--------Q--------------QI-IKG------------I---------I---A-----R-D-----------------L--D-H--T-D--AV-A--SITAGGVGFSYANVKLKS-ERG--SGLNY-Q-------IEF-Y---------------V--------------------------------------------------------------------------------------------------------------------------------------------------------------------------------------------------------------------------------------------------------------------------------------------------------------------------------

>Msex-betaREPAT4a

---------MR------L-V----------S---L-C------I--I-------F-----V---I--V-----------F---C---------------L---------------------------------------------------------------------------C---E--C---K-----D-------------------------------------------------------------------------------------------------------------------------------------------------------------------------------------------------------M---------IIG-D----------TVH--------R----------------------------------------K--MVF--H-----------QRV------------------K-DF-AIPF-----K-K-RIKTLSY----SD-----P-----E--------K--------------RM-IKG------------V---------A---A-----I-D-----------------N--D-F--S-H--AS-A--NITEGGVGYSYVTVRMKS-QRH--HPLNF-E-------VEI-Y---------------V--------------------------------------------------------------------------------------------------------------------------------------------------------------------------------------------------------------------------------------------------------------------------------------------------------------------------------

>Msex-betaREPAT4b

---------MR------L-V----------S---L-C------I--I-------F-----V---I--V-----------F---C---------------L---------------------------------------------------------------------------C---E--C---K-----D-------------------------------------------------------------------------------------------------------------------------------------------------------------------------------------------------------M---------IIG-D----------TVH--------R----------------------------------------K--MVF--H-----------QRV------------------K-DF-AIPF-----K-K-RIKTLSY----SD-----P-----E--------K--------------RM-IKG------------V---------A---A-----I-D-----------------N--D-F--S-H--AS-A--NITEGGVGYSYVTVRMKS-QRH--HPLNF-E-------VEI-Y---------------V--------------------------------------------------------------------------------------------------------------------------------------------------------------------------------------------------------------------------------------------------------------------------------------------------------------------------------

>Msex-betaREPAT5

---------MR-L----F-V----------I---F-C------L--L-------L-----A---I--I-----------T---T---------------V---------------------------------------------------------------------------V------G---E-----H-------------------------------------------------------------------------------------------------------------------------------------------------------------------------------------------------------L---------IMGDT----------NSN--------R----------------------------------------Q--LIH--H-----------TMA------------------Q-YH-AIPM-----M-K-RVKNVFY----NN-----Q-----E--------Q--------------RI-ISS------------I---------M---A-----Y-D-----------------N--L-N--S-K--AS-A--TVTAGGIGFSYVNIRLKS-ERG--KGLDY-D-------IGI-Y---------------S--------------------------------------------------------------------------------------------------------------------------------------------------------------------------------------------------------------------------------------------------------------------------------------------------------------------------------

>Msex-betaREPAT6

---------MK------I-E----------I-----------------------------L---V--V-----------L---C---------------VI--------------------------------------------------------------------------C------V---Y-VSAKD-------------------------------------------------------------------------------------------------------------------------------------------------------------------------------------------------------F---------VVG-T----------RTN--------N----------------------------------------L--LIS--T-----------EKV------------------I-VK-GYPF-----I-K-RDKDYIY----TD-----S-----K--------Q--------------KT-LKG------------I---------I---A-----R-D-----------------L--S-R--T-D--GE-V--SVKSGGVGASNVTLHLQS-ERG--EGLNY-L-------ILI-F---------------S---N-K--------------------------------------------------------------------------------------------------------------------------------------------------------------------------------------------------------------------------------------------------------------------------------------------------------------------------

>Msex-betaREPAT7a

---------MK------L-Y----------I---L-G------L--L-------L-----A---V--V-----------V---A---------------T---------------------------------------------------------------------------E------C---G-----H-------------------------------------------------------------------------------------------------------------------------------------------------------------------------------------------------------T---------FMG-T----------NVM--------R----------------------------------------R--QAH--H-----------AKA------------------K-FA-ANTF-----S-K-RIEYVSY----T----------------IPNSHN--------------QS-IQG------------I---------L---A-----Y-D-----------------L--T-N--S-G--AS-A--NVTSGGLGFNHVTLRLKS-DRG--KRLQY-D-------IYI-Y---------------V--------------------------------------------------------------------------------------------------------------------------------------------------------------------------------------------------------------------------------------------------------------------------------------------------------------------------------

>Msex-betaREPAT7b

---------MK------L-Y----------I---L-G------M--L-------L-----A---V--V-----------V---V---------------T---------------------------------------------------------------------------E------C---G-----H-------------------------------------------------------------------------------------------------------------------------------------------------------------------------------------------------------T---------FMG-T----------NVM--------R----------------------------------------R--QAH--H-----------ATV------------------K-FG-ANMF-----T-K-RIEYVNY----T----------------MPYSNN--------------QS-IQG------------I---------L---A-----Y-D-----------------L--T-N--S-G--AS-A--NVTSGGLGYNHVTLRLKS-DRS--KKLQY-D-------IYI-Y---------------V--------------------------------------------------------------------------------------------------------------------------------------------------------------------------------------------------------------------------------------------------------------------------------------------------------------------------------

>Msex-betaREPAT7c

---------MK------L-Y----------I---L-G------L--L-------L-----A---V--V-----------V---V---------------T---------------------------------------------------------------------------E------C---G-----H-------------------------------------------------------------------------------------------------------------------------------------------------------------------------------------------------------T---------FMG-T----------NVM--------R----------------------------------------R--QAH--Y-----------AKM------------------R-FG-ANTF-----S-K-RIEYFNY----T----------------VPHNLN--------------QS-IQG------------I---------L---A-----Y-D-----------------L--T-N--S-G--AS-A--NVTSGGMGFNHVTLRMKS-DRG--KKLQY-D-------IYI-Y---------------V--------------------------------------------------------------------------------------------------------------------------------------------------------------------------------------------------------------------------------------------------------------------------------------------------------------------------------

>Msex-betaREPAT8b

---------MF----A-K-I-SV-------------T------L--A-------L-----L---Y--L----A------S---F---------------A---------------------------------------------------------------------------E------S---N-----D-------------------------------------------------------------------------------------------------------------------------------------------------------------------------------------------------------I---------RLG-L---------VTPQ--------S----------------------------------------R--KIF--S-----------EIK------------------E----ADPA-----LWK-RTDDVIV----NA-----P-----N--------N--------------EF-ISA------------V---------Y---V-----T-D-----------------LRDD-K----D--GE-A--YIEAGGIGTKSVTVALKS-PTI-LRGYKF-L-------IEV-Y---------------G--------------------------------------------------------------------------------------------------------------------------------------------------------------N-------D--------P-----NANYFSKGIHSQHHY------------------DD-SQYA-----------------------R-------KY------------------------------------------------------------------

>Msex-betaREPAT9

---------MS------F-V----------V-P-I-L------A--A-------V-----V---V--I-----------A---S--------------------------------------------------------------------------------------------------C---N-----D-------------------------------------------------------------------------------------------------------------------------------------------------------------------------------------------------------L---------NLG-Y---------PDPT--------A----------------------------------------H--LIY--S-----------KVH------------------Q----ENPA-----IWV-RSETFRV----NC-----S-----S--------S--------------QV-ISA------------I---------R---I-----L-D-----------------LRQD-K----W--GE-A--YIKGGGIGQNSVTIELNS-PGV-FRGYNF-W-------VEV-Y---------------A--------------------------------------------------------------------------------------------------------------------------------------------------------------I----Q-TN--------H--FLYNHG--------------------------------------------------------------------KK------------------------------------------------------------------

>Msex-gammaREPAT1

---------MK-----TL-I----------F---L-V---------L-------A---VLG---A--------------N---A---------A---------------------------------------------------------------------------------------------------V-----V--K--------A---------------------------------------------------------------------------------------------P---------------------------------------------AA------------RS---------------------N---L---------NIG-Y----------IVP--------G----------------------------------------D--RLL--H-----------RSY------------------VYRP-AMAN-----S-M-QSENVSY----YG-----N-----R--------T--------------TR-ISA------------V---------Q---A-----Q-E-----------------V--G-A--SQY--AS-A--WIISGGIGQNNVTVRFQS-AVG--RGYYY-L-------LSV-W---------------G--------------------------------------------------------------------------------------------------------------------------------------------------------------R-----------------------------------------------------------------------------------------------------------------------------------------------------------------

>Msex-gammaREPAT2

---------MK-----TL-I----------F---L-V---------L-------A---VLG---A--------------N---A---------A---------------------------------------------------------------------------------------------------V-----V--K--------S---------------------------------------------------------------------------------------------P---------------------------------------------AA------------RS---------------------N---L---------NVG-F----------IAT--------G----------------------------------------D--RLL--H-----------SSY------------------VYRP-AMAN-----A-I-QSENVSF----NG-----N-----R--------T--------------TR-ITA------------V---------Q---A-----T-E-----------------V--G-A--TQF--AS-A--WIFSGGIGQNSVTVRFQS-SVG--RGYYY-L-------LRV-W---------------G--------------------------------------------------------------------------------------------------------------------------------------------------------------R-----------------------------------------------------------------------------------------------------------------------------------------------------------------

>Msex-gammaREPAT3

---------MK-----TL-I----------F---L-V---------L-------A---VLG---A--------------H---A---------A---------------------------------------------------------------------------------------------------V-----V--K--------A---------------------------------------------------------------------------------------------P---------------------------------------------AT------------RA---------------------N---L---------NVG-S----------IAT--------G----------------------------------------D--RLL--H-----------SST------------------VYRP-PRAN-----A-V-QSENVSF----YG-----N-----R--------T--------------TR-ISA------------V---------Q---A-----Q-E-----------------V--G-A--SQR--AS-A--WIISGGIGRNNVTVRFQS-AVG--RGYNY-I-------LRV-W---------------G--------------------------------------------------------------------------------------------------------------------------------------------------------------R-----------------------------------------------------------------------------------------------------------------------------------------------------------------

>Msex-gammaREPAT4

---------MK-----TL-I----------F---L-V---------L-------A---VLG---A--------------H---A---------A------------------------------------------------------------------------------------------------------------------------------------------------------------------------------------------------------------------------------------------------------------------------------------------------------N---L---------NVG-S----------IAA--------G----------------------------------------D--RLL--HL----------DST------------------VYRP-PRAN-----A-V-QSENVSF----YG-----N-----R--------N--------------TR-ITA------------V---------Q---A-----Q-E-----------------V--G-A--SQR--AS-A--RIISGGLGRNNVTVRFQS-AVG--RGYNY-V-------FRV-W---------------G--------------------------------------------------------------------------------------------------------------------------------------------------------------R-----------------------------------------------------------------------------------------------------------------------------------------------------------------

>Msex-gammaREPAT5

---------MK-----SL-I----------L---L-V---------L-------A---VLG---A--------------N---A---------A---------------------------------------------------------------------------------------------------S-----V--K--------T---------------------------------------------------------------------------------------------S---------------------------------------------AA------------RS---------------------S---F---------NAG-F----------IAV--------G----------------------------------------D--SLL--Q-----------RSI------------------VYRG-GRAN-----E-V-QTETVSF----NG-----N-----S--------T--------------TR-ITA------------V---------Q---A-----H-E-----------------Q--G-E-FSQR--PT-A--WMISGGIGLNNVTVKFKS-AVG--RGYYY-H-------LMV-W---------------G--------------------------------------------------------------------------------------------------------------------------------------------------------------R--------------------------------------------------------------------------------------------------------P--------------------------------------------------------

>Msex-gammaREPAT6

---------MK-----TL-I----------F---L-V---------L-------A---VLG---A--------------N---A---------A---------------------------------------------------------------------------------------------------S-----V--K--------T---------------------------------------------------------------------------------------------S---------------------------------------------AA------------RS---------------------S---F---------NAG-F----------IAV--------G----------------------------------------D--TLL--Q-----------RSI------------------VYRG-GRAN-----E-V-QTETVSF----NG-----N-----S--------T--------------TR-ITA------------V---------Q---A-----H-E-----------------Q--G-E-FSQR--PT-A--XMISGGIGLNNVTVKFKS-AVE--E------------------------------------------------------------------------------------------------------------------------------------------------------------------------------------------------------------------------------------------------------------------------------------------------------------------------------------------------------------------

>Msex-gammaREPAT7

---------MK-----TL-I----------F---L-V---------L-------A---VLG---A--------------N---A---------A---------------------------------------------------------------------------------------------------S-----V--K--------T---------------------------------------------------------------------------------------------S---------------------------------------------AA------------RS---------------------S---F---------NAG-F----------IAV--------G----------------------------------------D--TLL--Q-----------RSI------------------VYRG-GRAN-----E-V-QTETVSF----NG-----N-----S--------T--------------TR-ITA------------V---------Q---A-----H-E-----------------Q--G-E-FSQR--PT-A--WMISGGIGLNNVTVKFKS-AVG--RGYYY-H-------LMV-W---------------G--------------------------------------------------------------------------------------------------------------------------------------------------------------R--------------------------------------------------------------------------------------------------------P--------------------------------------------------------

>Sexi-alphaREPAT1

---------MR-----NI-I----------II--I-A------V--L------VA-----L---A------------D-C---A---------------V--------------------------------------------------------------------------IN------N---E---------------------------------------------------------------------------------------------------------------------------------------------------------------------------------------------------------------------------L----------VTT--------RGT--------A-RP-----------------------G--A--VIF--F-----------EES------------------YYVP-AVPD-----T-I-QRLDVVY----RG-----N-----S--------T--------------TY-ISQ------------V---------R---I-----E-E--------------------P-R--PNV--NF-NF-S----PLGNTYMQVRLAS-AVG--MPIQV-R-------IQF-L---------------G--------------------------------------------------------------------------------------------------H----------------------------------------YTDWP--------------------------------------------------------------------------------------------------------------------------------------------------------------------------------

>Sexi-alphaREPAT10

---------MR-----IF-I----------I---I-A------V--I------SA-----F---V--V------CY-G-D---I---------------I----NPS---------M------------------E--------P--M-------------SLV--Y--DDDDSSL------D---G---------------------------------------------------------------------------------------------------------------------------------------------------------------------------------------------------------------------------K----------LLS--------SSE---I---NL-RN--------------------QQW------III--S-----------EWQ------------------CHAP-AVAG-----Q-V-QHVGVQY----QG-----A-----S--------S--------------VT-INR------------L---------I---V-----T-Y-----------------T--Q-T--PAQ--VI-R--T----PLSTNAMNVTIVS-NSG--QLINA-T-------VTL-L---------------------------------------------------------------------------------------------------------------------------------------------------------------R--------------R----------------------------------------------------------------------------------------------------------------------------------------------------------------A

>Sexi-alphaREPAT11

---------MR-----TF-I----------I---I-T------V--L------SA-----L---V--A------CF-S-A---S---------------V----IPS---------E------------------E--------E--M----------H--TIM--Y--D--EKFP------N---G---------------------------------------------------------------------------------------------------------------------------------------------------------------------------------------------------------------------------T----------IVS--------QAE---L---DS-KYN--------TG------------------ALV--Y-----------QWH------------------CTAP-AISR-----Q-R-QTTILQY----LG-----N-----P--------T--------------EV-VTR------------IIFIHTGSTPQ---V-----T-Y------------------------------------T----PLYGNTVTITITS-QSG--AAMNS-V-------VQV-F---------------------------------------------------------------------------------------------------------------------------------------------------------------A--------------V-----------------------------------------------------------------------------------------------------------------------------------------------------------------

>Sexi-alphaREPAT12

---------MR-----SL-I----------I---I-A------V--L------SA-----L---A--A------CY-VNA-------------------V----HPT---------E------------------E-D----MSE-------------V-YTIV--F--D--DNSL------D---G---------------------------------------------------------------------------------------------------------------------------------------------------------------------------------------------------------------------------R----------VLS--------RAE---F---NA-RY---------------EG-W----------VLL--S-----------EAN------------------DFAP-AVEG-----Q-E-QSVEMFY----RA-----Q-----R--------G--------------VQ-ISM------------V---------S---I-----TAY-----------------V--S----IPE--IN-R--S----PLGKNYIEVTLKS-RIG--AGLNT-T-------IEL-F---------------------------------------------------------------------------------------------------------------------------------------------------------------R--------------R----------------------------------------------------------------------------------------------------------------------------------------------------Q------------

>Sexi-alphaREPAT13

---------MR-----SF-I----------I---I-T------V--L------SA-----L---A--A------CY-G-T-------------------V----LPS---------E------------------EK-----VSR--G----------Y--TMI--Y--D--DMSP------D---G---------------------------------------------------------------------------------------------------------------------------------------------------------------------------------------------------------------------------R----------IAS--------RVE---L---ES-IS---------------DG-Y----------ILL--S-----------WGA------------------LKAP-AVAG-----Q-I-QYISKVY----KA-----E-----P--------G--------------NT-ISS------------V---------V---I-----T-H-----------------S--I-P--PAL--VL-R--T----PLGRNSSKVSITS-YRG--QGINA-T-------IST-Y---------------------------------------------------------------------------------------------------------------------------------------------------------------G--------------R----------------------------------------------------------------------------------------------------------------------------------------------------E------------

>Sexi-alphaREPAT14

---------MK-----SF-I----------I---I-V------V--L------SA-----L---A--A------CY-DAA-------------------V----LPS---------E------------------E-E----MSQD------------F--TIV--F--D--DKSP------N---G---------------------------------------------------------------------------------------------------------------------------------------------------------------------------------------------------------------------------M----------VLS--------RAD---F---NA-RY---------------EH-F----------VLL--S-----------ESK------------------DFVP-AIKG-----Q-E-QSVTMLY----RA-----Q-----R--------D--------------VK-IGM------------I---------A---I-----TSH-----------------F--I-H--LPE--IV-R--S----PLGKNYIEVTLKS-RVG--QGLNT-T-------IQT-Y---------------------------------------------------------------------------------------------------------------------------------------------------------------R--------------W----------------------------------------------------------------------------------------------------------------------------------------------------K------CCQN--

>Sexi-alphaREPAT15

---------MK-----SF-M----------I---I-V------V--L------SA-----L---T--A-------Y-DAA-------------------V----LPS---------E------------------E-E----MSQ-------------V--TIV--F--D--DKSP------N---G---------------------------------------------------------------------------------------------------------------------------------------------------------------------------------------------------------------------------I----------VLS--------RAD---F---NA-RY---------------EH-F----------VLL--S-----------ESK------------------DSVP-AVEG-----Q-E-QSLTKVY----RA-----E-----S--------G--------------VK-ISM------------I---------S---I-----TAH-----------------F--S-H--LPE--IV-R--S----PLGKNYIEITLKS-RVG--EGLNT-T-------IQT-Y---------------------------------------------------------------------------------------------------------------------------------------------------------------R--------------W----------------------------------------------------------------------------------------------------------------------------------------------------K------------

>Sexi-alphaREPAT16

---------MK-----SF-I----------V---I-V------V--L------SA-----L---A--A------CY-DAA-------------------V----LPS---------E------------------K-E----MSQ-------------V--TIV--F--D--DKSP------N---G---------------------------------------------------------------------------------------------------------------------------------------------------------------------------------------------------------------------------M----------VLS--------RAD---F---NA-RY---------------EH-F----------ALL--S-----------ESK------------------DFVP-AVEG-----Q-E-QSLTKVY----RA-----E-----S--------G--------------VE-ISM------------I---------S---I-----TAH-----------------F--S-H--LPE--IV-R--S----PLGKNYIEITLKS-RVG--GGLNT-T-------IQT-Y---------------------------------------------------------------------------------------------------------------------------------------------------------------R--------------W----------------------------------------------------------------------------------------------------------------------------------------------------K------------

>Sexi-alphaREPAT17

---------MK-----SF-M----------I---I-V------V--L------SA-----L---V--A------CY-DAA-------------------V----LPS---------E--------------------E----MSQ-------------V--TIV--F--D--DKSP------N---G---------------------------------------------------------------------------------------------------------------------------------------------------------------------------------------------------------------------------M----------VLS--------RAD---F---NA-RY---------------EH-F----------VLL--S-----------ESK------------------DFVP-AVEG-----Q-E-QSLTKVY----RA-----E-----S--------G--------------VE-ISM------------I---------S---I-----TAH-----------------F--S-H--LPE--IV-R--S----PLGKNYIEITLKS-RVG--GGLNT-T-------IQT-Y---------------------------------------------------------------------------------------------------------------------------------------------------------------R--------------W----------------------------------------------------------------------------------------------------------------------------------------------------K------------

>Sexi-alphaREPAT18

---------MK-----SF-M----------I---I-V------V--L------SA-----L---V--A------CY-DAA-------------------V----LPS---------E--------------------E----MSQ-------------V--TIV--F--D--DKSP------N---G---------------------------------------------------------------------------------------------------------------------------------------------------------------------------------------------------------------------------M----------VLS--------RAD---F---NA-RY---------------EH-F----------VLL--S-----------ESK------------------DFVP-AVEG-----Q-D-QSLTKVY----RA-----E-----S--------G--------------VE-ISM------------I---------S---I-----TAH-----------------F--S-H--LPE--IV-R--S----PLGKNYIEITLKS-RVG--GGLNT-T-------IQT-Y---------------------------------------------------------------------------------------------------------------------------------------------------------------R--------------W----------------------------------------------------------------------------------------------------------------------------------------------------K------------

>Sexi-alphaREPAT19

---------MK-----SF-M----------I---I-V------V--L------SA-----L---A--V------CY-DAA-------------------V----LPS---------E------------------E-K----MSQ-------------V--TIV--F--D--DKSP------N---G---------------------------------------------------------------------------------------------------------------------------------------------------------------------------------------------------------------------------M----------VLS--------RAD---F---NA-RY---------------EH-F----------VLL--S-----------ESK------------------DFVP-AVEG-----Q-E-QSLTKVY----RA-----E-----S--------G--------------VE-ISM------------I---------S---I-----TAH-----------------F--S-H--LPE--IV-R--S----PLGKNYLEITLKP-RVG--EGLNT-T-------IQT-Y---------------------------------------------------------------------------------------------------------------------------------------------------------------R--------------W----------------------------------------------------------------------------------------------------------------------------------------------------K------------

>Sexi-alphaREPAT2

---------MR-----TS-I----------I---I-A------V--L------AA-----L---T--A------CH-D-A---A---------------T----PQS---------L------------------E--------E--R-------FPGY--KLV--YDVE--DVAQ------N---R---------------------------------------------------------------------------------------------------------------------------------------------------------------------------------------------------------------------------I----------TLS--------NGE--------V-RH-----------------------D--E--VLL--Y-----------EEE------------------IYKP-AIPG-----V-V-QTVVVSH----RA-----K-----P--------G--------------TW-ISR------------S---------S---L-----S-I-----------------Y--P-L--PSY--IH-T--Y----RLIPHMILIRISS-QVG--QGLVG-R-------IRV-F---------------G-IL-----------------------------------------------------------------------------------------------D----------------------------------------YYDKN--------------------------------------------------------------------------------------------------------------------------------------------------------------------------------

>Sexi-alphaREPAT20

---------MR-----NS-I----------I---I-T------L--L------SA-----L---A--V------CY-S-A---S---------------V-----SD---------T------------------E--------E--M---PKNL---I--YTL--Y--D--ANSP------N---G---------------------------------------------------------------------------------------------------------------------------------------------------------------------------------------------------------------------------T----------VVS--------LDD---F---SM-KS-------------------S---------SVL--F-----------TWR------------------YNVP-PIRG-----E-A-QRVEMKY----EG-----E-----P--------G--------------DR-VSS------------I---------M---V-----G-Q-----------------S--T-T--PAY--MY-Q--T----PLFNSYMDVHIVS-HPG--QGIDT-Y-------IAI--------------------------------------------------------------------------------------------------------------------------------------------------------------------------------R-----------------------------------------------------------------------------------------------------------------------------------------------------------------

>Sexi-alphaREPAT21

---------MR-----SF-I----------I---I-V------V--L------SA-----L---A--A------CY-G-AS------------------V----LPS---------E-D----------------E------MTG--E----------Y--TLV--Y--D--DMSP------E---G---------------------------------------------------------------------------------------------------------------------------------------------------------------------------------------------------------------------------R----------IMS--------TAE---L---NA-RS---------------DD-W----------ILV--S-----------EGI------------------VNAP-AAPG-----Q-V-QRIGTWY----QA-----D-----A--------G--------------VR-IER------------L---------A---L-----S-Y-----------------S--A-F--PAD--VY-H--S----PLGTNYLEVRITS-VPS--QRIYS-N-------ISA-F---------------------------------------------------------------------------------------------------------------------------------------------------------------R--------------R---------------------------------------------------------------------------------------------------------------------------------------------------AR------------

>Sexi-alphaREPAT22

---------MR-----SF-V----------I---I-V------V--L------SA-----L---A--A------CY-G-S---S---------------V----LPGKANMFSSFVR------------------E--------Q--M-------SKGW--ILE--HNEE--DAAQ------N---R---------------------------------------------------------------------------------------------------------------------------------------------------------------------------------------------------------------------------V----------TLT--------KGD--------T-SD-----------------------D--N--ILL--Y-----------QQT------------------CYVP-AIPY-----D-F-VRCDVVY----RG-----E-----P--------N--------------EK-IYR------------I---------S---TS----S-Y-----------------W--P-S--PSH--IY-V--D----GLLTNVGRVRMVS-KRG--ERLYS-T-------VKM-Y---------------GKII-----------------------------------------------------------------------------------------------E----------------------------------------DNSES--------------------------------------------------------------------------------------------------------------------------------------------------------------------------------

>Sexi-alphaREPAT23

---------MR-----SF-V----------I---I-V------V--L------SA-----L---A--A------CY-G-Y---S---------------V----LPGKANMFSSFVR------------------E--------Q--M-------SKGW--ILG--HNEE--DAAQ------N---R---------------------------------------------------------------------------------------------------------------------------------------------------------------------------------------------------------------------------V----------TLT--------KGE--------T-SD-----------------------D--Y--ILF--Y-----------QQT------------------CDVP-AIPN-----D-F-GRCDVVY----RG-----E-----P--------N--------------EK-IYR------------F---------E---TS----S-S-----------------F--L-P--SS---IY-T--E----GLGTNVGQVRLVS-QRG--EMLYA-I-------VRM-Y---------------GKII-----------------------------------------------------------------------------------------------E----------------------------------------DNSES--------------------------------------------------------------------------------------------------------------------------------------------------------------------------------

>Sexi-alphaREPAT24

---------MR-----SF-V----------I---I-V------V--L------SA-----L---A--A------CY-G-S---S---------------I----LPGKANMFTSFVR------------------E--------Q--M-------SKGW--ILG--HNEE--DAAQ------N---R---------------------------------------------------------------------------------------------------------------------------------------------------------------------------------------------------------------------------V----------TLT--------KGE--------T-SD-----------------------D--Y--ILF--Y-----------QQT------------------CDVP-AIPY-----D-F-GRCDVVY----RG-----E-----P--------N--------------EK-IYR------------F---------E---TS----S-S-----------------F--L-P--SS---IY-T--E----GLGTNVGKVRLVS-QRG--EMLYA-I-------VRM-Y---------------GKII-----------------------------------------------------------------------------------------------E----------------------------------------DNSES--------------------------------------------------------------------------------------------------------------------------------------------------------------------------------

>Sexi-alphaREPAT25

---------MR-----SF-I----------I---I-V------V--L------SA-----V---A--A------CY-G-S---S---------------I----LPGKANMFSSFVR------------------E--------Q--M-------SKGW--ILE--HNEE--DAAQ------N---R---------------------------------------------------------------------------------------------------------------------------------------------------------------------------------------------------------------------------V----------TLT--------KGE--------T-SD-----------------------D--Y--ILF--Y-----------QQT------------------CYVP-AIPY-----D-F-GRCDVVY----RG-----E-----P--------N--------------EK-IYS------------I---------Q---IT----S-Y-----------------W--P-P--PSY--IH-S--E----GLGTNVGRVRIVS-QWG--EMLYT-I-------VRM-Y---------------GKII-----------------------------------------------------------------------------------------------E----------------------------------------DNSES--------------------------------------------------------------------------------------------------------------------------------------------------------------------------------

>Sexi-alphaREPAT26

---------MR-----SF-V----------I---I-V------V--L------SA-----V---A--A------CY-G-S---S---------------I----LPGKANMFSSFVR------------------E--------Q--M-------SKGW--ILE--HNEE--DAAQ------N---R---------------------------------------------------------------------------------------------------------------------------------------------------------------------------------------------------------------------------V----------TLT--------KGE--------T-SD-----------------------D--Y--ILF--Y-----------QQT------------------CYVP-AIPY-----D-F-GRCDVVY----RG-----E-----P--------N--------------EK-IYS------------I---------Q---IT----S-S-----------------W--P-P--PSY--IH-S--E----GLGTNVGRVRIVS-QWG--EMLYT-I-------VRM-Y---------------GKII-----------------------------------------------------------------------------------------------E----------------------------------------DNSES--------------------------------------------------------------------------------------------------------------------------------------------------------------------------------

>Sexi-alphaREPAT27

---------MR-----SF-M----------I---V-V------V--L------SA-----L---A--A------CY-DAA-------------------V----LPS---------E------------------E-Q----MSQ-------------V---IV--F--D--EKSP------N---G---------------------------------------------------------------------------------------------------------------------------------------------------------------------------------------------------------------------------M----------VLS--------RAD---F---NA-RY---------------GG-F----------VLL--S-----------ESK------------------DFVP-AVEG-----Q-E-QSITKVY----RA-----E-----S--------D--------------IK-IDM------------T---------V---V-----TSF-----------------Y--N-P--LPE--II-H--S----PLGRNYVEITLIP-RVG--QELNT-Q-------IQI-Y---------------------------------------------------------------------------------------------------------------------------------------------------------------R--------------R----------------------------------------------------------------------------------------------------------------------------------------------------S----YN------

>Sexi-alphaREPAT28

---------MR-----SL-M----------I---I-V------V--L------SA-----L---A--A------CY-NAA-------------------V----LPS---------E------------------E-E----MSQ-------------VT--IV--F--D--EKSP------N---G---------------------------------------------------------------------------------------------------------------------------------------------------------------------------------------------------------------------------M----------VLS--------RAD---F---NA-RY---------------SG-F----------VLL--S-----------ESK------------------DFVP-AVEG-----E-E-QSVTKVY----RA-----E-----S--------D--------------IK-IGM------------T---------V---V-----YTY-----------------L--S-H--LPE--II-H--S----PLGKNYVEITLKP-RLG--EGLDT-T-------LEI-Y---------------------------------------------------------------------------------------------------------------------------------------------------------------R--------------S----------------------------------------------------------------------------------------------------------------------------------------------------A------------

>Sexi-alphaREPAT29

---------MR-----SL-M----------I---I-V------V--L------SA-----L---A--A------CY-DAA-------------------V----LQS---------E------------------E-E----MSQ-------------V---IV--F--D--EKSP------N---G---------------------------------------------------------------------------------------------------------------------------------------------------------------------------------------------------------------------------V----------VLF--------RAD---F---NA-RY---------------GS-Y----------ELV--S-----------KSN------------------DFLP-AVEG-----Q-E-QSITKVY----RA-----E-----S--------D--------------IK-INM------------F---------V---V-----TTH-----------------L--S-H--LPE--II-H--S----SLGKNYVEITLKPERLG--EGLHT-T-------LEI-Y---------------------------------------------------------------------------------------------------------------------------------------------------------------R--------------S----------------------------------------------------------------------------------------------------------------------------------------------------A------------

>Sexi-alphaREPAT3

---------MR-----IF-I----------I---I-T------V--L------SA-----L---A--A------CY-G-A---A---------------V----LPN---------D------------------E--------E--R-------NSNY--TLEPKNNED--DGVK------S---K---------------------------------------------------------------------------------------------------------------------------------------------------------------------------------------------------------------------------I----------STF--------GAE--------V-KD-----------------------T--DGLELL--Y-----------EQQ------------------FVRT-AVPN-----F-I-HNVDVLY----YG-----Q-----P--------P--------------RV-IKA------------A---------V---I-----Q-T-----------------F--P-T--ASW--IF-P--V----WHLDNYISIRISS-QRG--HGMNG-T-------VQI-Y---------------G-TL-----------------------------------------------------------------------------------------------------------------------------------------------------------------------------------------------------------------------------------------------------------------------------------------------------------------------------

>Sexi-alphaREPAT30

---------MR-----SL-M----------I---I-V------V--L------SA-----L---A--A------CY-DAA-------------------V----LQS---------E------------------E-E----MSQ-------------V---IV--F--D--EKSP------N---G---------------------------------------------------------------------------------------------------------------------------------------------------------------------------------------------------------------------------V----------VLF--------RAD---F---NA-RY---------------GS-Y----------ELV--S-----------KSN------------------DFLP-AVEG-----Q-E-QSITKVY----RA-----E-----S--------D--------------IK-INM------------F---------V---V-----TTH-----------------L--S-H--LPE--II-H--S----SLGKNYVEITLKPERLG--EGLHT-T-------LEI-Y---------------------------------------------------------------------------------------------------------------------------------------------------------------R--------------S----------------------------------------------------------------------------------------------------------------------------------------------------A------------

>Sexi-alphaREPAT4

---------MK-----YF-V----------I---I-A------V--L------SV-----L---I--D------CY-E-A---F---------------V----LLD---------R--QM--K---------MSE--------G--F-------------TIV--Y--D--DT-V------N---G---------------------------------------------------------------------------------------------------------------------------------------------------------------------------------------------------------------------------T----------IMS--------NAE---V---IA-RN--------------------DGW------RLL--Y-----------EVA------------------CTSP-AVVG-----R-V-KNSYMTY----RG-----P-----A--------N--------------TV-IGR------------I---------T---V-----T-P-----------------D--N-Q--PAN--VY-H--S----ALGTNNLEVRLMS-NIS--QGIQA-T-------VRL-Y---------------------------------------------------------------------------------------------------------------------------------------------------------------Q--------------R------------------------------------------------------------------------------------------------------------------SGAGKAFGEPAFHLMFFVLLYFGISTF-------------------T

>Sexi-alphaREPAT5

---------MR-----SF-I----------I---I-A------AVTL------SA-----L---A--A------CY-G-S---A---------------V----LPS---------E------------------E--------A--MQES-------H--TVL--Y--D--DKSP------N---G---------------------------------------------------------------------------------------------------------------------------------------------------------------------------------------------------------------------------R----------ILS--------NAE---L---HA-IEK--S----TTL------------------RLL--S-----------KWK------------------CDAP-SMPG-----H-R-QRVSVTF----DG-----P-----P--------GT-------------EI-LQY------------IVTYFGSA-T----I-----D-R------------------------------------T----PFSTNFMQINITS-AIG--ESVDA-T-------VQM-S---------------------------------------------------------------------------------------------------------------------------------------------------------------G--------------T-----------------------------------------------------------------------------------------------------------------------------------------------LN----------------

>Sexi-alphaREPAT6

---------MR-----SF-I----------I---I-A------A--L------SA-----L---A--A------CY-G-S---A---------------V----LPS---------E------------------E--------A--MQEG-------Y--TIL--Y--D--DKSP------N---G---------------------------------------------------------------------------------------------------------------------------------------------------------------------------------------------------------------------------R----------IMS--------STE---L---DA-RVK--D----MSY------------------GLL--S-----------VWQ------------------CDAP-AIPG-----H-K-QRVNVIY----EG-----S-----P--------GI-------------QI-LRF------------GVTPFNPD-IE---I-----E-F------------------------------------T----PFLTNFMQINVTY-PVG--IKADA-I-------IQM-F---------------------------------------------------------------------------------------------------------------------------------------------------------------G--------------I-----------------------------------------------------------------------------------------------------------------------------------------------QY----------------

>Sexi-alphaREPAT7

---------MK-----SL-I----------I---I-T------V--L------TA-----L---V--A------CY-E-A-----------------------------------------A-D---------LPE--------G--F-------------SLV--Y--D--DTAV------D---G---------------------------------------------------------------------------------------------------------------------------------------------------------------------------------------------------------------------------T----------IMS--------NRD---L---NE-RN--------------------QNF------RLV--Y-----------EAR------------------CTAS-AVVG-----R-S-KTSLVTY----RG-----P-----S--------N--------------VR-INR------------I---------V---I-----N-S-----------------P--I-Q---AN--VY-H--S----ALDTNNMEIRITS-NVS--RGVDT-T-------VFL-Y---------------------------------------------------------------------------------------------------------------------------------------------------------------Q--------------R------------------------------------------------------------------------------------------------------------------SGAGRSFGEPAFHLIISVIMYFGYSRL-------------------A

>Sexi-alphaREPAT8

---------MR-----SF-I----------I---I-V------V--L------SI-----L---A--A------CY-D-A---S---------------V----LSR---------Q------------------E--DPTA--E--Y----------V--TST--M--Y--E---------N---G---------------------------------------------------------------------------------------------------------------------------------------------------------------------------------------------------------------------------T----------IVY--------GED---FAFADL-RS-----------------------------GLL--S-----------EWN------------------CEAP-SIFG-----Q-S-QTISLQY----LG-----D-----G--------S--------------ER-ITR------------RVTSIYGD-PQ---I-----A-Q------------------------------------T----PLGGNIYEMTLTS-RPG--VRISA-R-------IYI-Y---------------------------------------------------------------------------------------------------------------------------------------------------------------G--------------S-----------------------------------------------------------------------------------------------------------------------------------------------------------------

>Sexi-alphaREPAT9

---------MK-----WF-I----------I---I-T------V--L------SA-----F---T--A------CY-G-V---S---------------I----PSS---------E------------------E--------E--F----------I--TTI--Y--D--ENSP------N---G---------------------------------------------------------------------------------------------------------------------------------------------------------------------------------------------------------------------------T----------IVS--------SRD---VELTNA-RY-----------------------------VLL--N-----------EWN------------------CEIP-SIFG-----Q-S-HTTSFIY----IG-----H-----A--------T--------------DR-IRK------------ITTSFLGS-PQ---V-----S-N------------------------------------P----TLGGSVFDITITT-RPG--TRTTA-R-------IAI-F---------------------------------------------------------------------------------------------------------------------------------------------------------------T--------------------------------------------------------------------------------------------------------------------------------------------------------------------------------

>Sexi-betaREPAT1

---------MS------L-W----------T---Q-V---------T-------V-----L------C-----------L---C---------------L---------------------------------------------------------------------------A------L---Y--NAED-------------------------------------------------------------------------------------------------------------------------------------------------------------------------------------------------------L---------HLG-S----------GVN--------A----------------------------------------Q--LGY--Q-----------EIV------------------K-LG-SVPL-----S-V-RTKNVFY----NS-----N-----N--------T--------------KT-IKS------------I---------S---A-----V-D-----------------I--D-K--S-K--AV-P--TVTAGGIGSSFVNVRLKS-EKG--KGLNY-L-------VQI-F---------------V--------------------------------------------------------------------------------------------------------------------------------------------------------------------------------------------------------------------------------------------------------------------------------------------------------------------------------

>Sexi-betaREPAT10

MA-------AK------V-L----------L---S-I------L--L-------L--S--I---S--V-----------S---L---------------V-LAR-----------------------------------------------------------------------A------R---G-----S-------------------------------------------------------------------------------------------------------------------------------------------------------------------------------------------------------L---------SIG-N----------TFG--------S----------------------------------------H--LIY--Q-----------ETH------------------Y-KY-AVPF-----I-V-RDGDITV----KG-----V-----N--------S--------------EI-IAA------------V---------T---V-----N-D-----------------L--Q-G----M--GL-C--YVKDGGIGQHNVTIHLES-SRG--HGYKF-L-------VEV-Y---------------A--------------------------------------------------------------------------------------------------------------------------------------------------------------I-----------------------------------------------------------------------------------------------------------------------------------------------------------------

>Sexi-betaREPAT11a

----MFKH--------------------------L-L------I--L-------C-----A---I--I-----------V---A---------------V---------------------------------------------------------------------------H------C---K-----S-------------------------------------------------------------------------------------------------------------------------------------------------------------------------------------------------------V---------IVG-Q----------TWEQNGV---------------------------------------------K--KVY--D-----------TIV------------------S-AN-SIPL-----F-K-RTKYVNF----EF-----P-----N--------Q-E------------TK-IKG------------I---------A---I-----K-D-----------------Q--D-N--G-L--AE-P--SITMGGLGFNFVNLKLKS-ERS--SGFTF-L-------IEI-Y---------------G--------------------------------------------------------------------------------------------------------------------------------------------------------------------------------------------------------------------------------------------------------------------------------------------------------------------------------

>Sexi-betaREPAT11b

----MFKE--------------------------I-V------I--L-------F-----G---V--I-----------V---L---------------V---------------------------------------------------------------------------Q------S---N-----S-------------------------------------------------------------------------------------------------------------------------------------------------------------------------------------------------------V---------FVG-H----------VYDQTGL----V----------------------------------------R--KVY--D-----------KMV------------------E-AT-GIPL-----I-K-REEEIFY----EY-----P-----A--------G-D------------QK-IKG------------I---------A---I-----K-D-----------------L--A-D--S-K--AQ-A--SITRGGLGFGFVNIKLKS-ERG--SGYKY-L-------IEV-F---------------A--------------------------------------------------------------------------------------------------------------------------------------------------------------------------------------------------------------------------------------------------------------------------------------------------------------------------------

>Sexi-betaREPAT12

---------MK------T-L-------SV-A---I-L------F--C-------L-----L---A--A-----------A---A---------------S---------------------------------------------------------------------------Q------S---H-----D-------------------------------------------------------------------------------------------------------------------------------------------------------------------------------------------------------L---------ILG-Q----------ATY--------G----------------------------------------D--VIV--F-----------KAD------------------EVKY-GFPF-----I-V-RTSIVEY-----P-EQGQK-----N--------Y--------------AY-IKA------------I---------Y---V-----K-DNE---------------R--D-G--S-G--GY-P--SISAGGIGQRFVKIKLKS-QRG--SGLNF-T-------ITI-Y---------------G--------------------------------------------------------------------------------------------------------------------------------------------------------------R-----------------------------------------------------------------------------------------------YI----------------------------------------------------------------

>Sexi-betaREPAT13a

---------MF------A-K----------V---V-A------F--L-------A-----L------V-----------A---V---------------A---------------------------------------------------------------------------A------A---A-----D-------------------------------------------------------------------------------------------------------------------------------------------------------------------------------------------------------L---------SVG-K---------M--G--------G----------------------------------------R--LIF--Y-----------KNF------------------S----ASPA-----IWK-QVQTIIV----NA-----T-----D--------D--------------AL-IRR------------V---------V---V-----I-D-----------------NRPE-K----D--GE-A--TVVDGGEGHNNVTIMLKS-PTV-FRGFDF-T-------VKV-Y---------------A--------------------------------------------------------------------------------------------------------------------------------------------------------------K----KRTT--------Q-----APQH--------PVSVG----------------TD-IQV----------------------------------------------------------------------------------------------------

>Sexi-betaREPAT13b

---------MF------A-K----------V---V-A------L--F-------G-----L------V-----------A---V---------------A---------------------------------------------------------------------------A------A---A-----D-------------------------------------------------------------------------------------------------------------------------------------------------------------------------------------------------------L---------CVG-R---------E--G--------G----------------------------------------Q--LIF--D-----------KNI------------------T----ASPA-----IWK-QIQNLTV----NA-----T-----D--------D--------------AL-ISR------------V---------V---V-----I-D-----------------NRPE-K----D--GE-A--KVVDGGEGHNNVTIELKS-PTV-FRGFDF-T-------VKV-F---------------A--------------------------------------------------------------------------------------------------------------------------------------------------------------S----EGTT--------Q-----APQH--------PVSVG----------------AD-TQV----------------------------------------------------------------------------------------------------

>Sexi-betaREPAT13c

---------MF------A-K----------V---V-A------L--F-------G-----L------V-----------A---V---------------A---------------------------------------------------------------------------A------A---A-----D-------------------------------------------------------------------------------------------------------------------------------------------------------------------------------------------------------L---------SVG-R---------E--G--------G----------------------------------------Q--LIF--D-----------KNI------------------T----ASPA-----IWK-QVQNLTV----NA-----T-----D--------D--------------AL-ISR------------V---------V---V-----I-D-----------------NRPE-K----D--GE-A--KVVDGGEGHNNVTIELKS-PTV-FRGFDF-T-------VKV-F---------------A--------------------------------------------------------------------------------------------------------------------------------------------------------------S----EGAT--------Q-----APQH--------PVSIG----------------AD-TQV----------------------------------------------------------------------------------------------------

>Sexi-betaREPAT13d

---------MF------A-K----------V---V-A------L--F-------G-----L------V-----------A---V---------------A---------------------------------------------------------------------------T------A---A-----D-------------------------------------------------------------------------------------------------------------------------------------------------------------------------------------------------------L---------SVG-R---------E--G--------G----------------------------------------Q--LIF--D-----------KNI------------------T----ASPA-----IWK-QIQNLTV----NA-----T-----D--------D--------------AL-ISR------------V---------V---V-----I-D-----------------NRPE-K----D--GE-A--KVVDGGEGHNNVTIELKS-PTV-FRGFDF-T-------VKV-F---------------A--------------------------------------------------------------------------------------------------------------------------------------------------------------S----EGTT--------Q-----APQH--------PVSVG----------------AD-TQV----------------------------------------------------------------------------------------------------

>Sexi-betaREPAT15

---------MKLL----I-V----------L---S-V------V--C-------L-----S---V----------------------------------V---------------------------------------------------------------------------Y------A---N-----H-------------------------------------------------------------------------------------------------------------------------------------------------------------------------------------------------------L---------IVGNV----------ANH--------V----------------------------------------V--LAN--Q-----------TRV------------------E-YN-AIPF-----I-K-RVKYYFY----TQ-----P-----D--------N--------------RL-IQG------------I---------Q---A-----L-D-----------------T--M-H--S-K--AS-I--NITAGGVGYSYVNLRLKS-ERG--RGLSY-D-------IGI-Y-VQDIFYH----------------------------------------------------------------------------------------------------------------------------------------------------------------------------------------------------------------------------------------------------------------------------------------------------------------------------------------

>Sexi-betaREPAT16

---------MK------S-F---LLLC---V---L-T------L--T-------I-----S---S--V---------------L---------------S---------------------------------------------------------------------------S------TYFFG-----E---------------------------------------------------------------------------------------------------------------------------------------------------------------------------------------------------------------------Q----------TKD--------T----------------------------------------R--IIY--Q-----------DKV------------------R-YE-AIPL-----K-K-RVKTYTY----NG-----T-----V------------------------P-IKA------------I---------S---C-----Y-D-----------------F--Q-N--S-E--AS-V--NITAGGIGYKYVELRLKS-QRS--YRLEY-A-------IEI-Y---------------A--------------------------------------------------------------------------------------------------------------------------------------------------------------------------------------------------------------------------------------------------------------------------------------------------------------------------------

>Sexi-betaREPAT17

---------MD------L-L----------L---I-V------L--L-------V-----S---V--T-----------D---A------KMF------C---------------------------------------------------------------------------K------K---P-----K-------------------------------------------------------------------------------------------------------------------------------------------------------------------------------------------------------A---------PL-------------------------------------------------------------------------------------ELG------------------EWEY-HLKH-----M-D--SAVVEN-----P-----P-----L--------S--------------IN-EGQ------------M---------Y---V-----Y-ENYFPNYTIKYIYVDNLAM--R-S--C-G--AS-A--AIKTGGIDSSYVVLVLHA-HIN--QEIRS-V-------IDI-W---------------G-------ERT--------TV--G---KST--ITEI--QP----DETTSKKLFHLFKHLRAVNHNR---------------------------------------------------------------------------------------------------------------------------------------------------------------------------------------------------------------------------------------------------------------

>Sexi-betaREPAT2

---------MK-C-V--L-A----------L---S-V------F--V-------L-----I---A--I-----------S---S---------------C---------------------------------------------------------------------------F------A---N-----D-------------------------------------------------------------------------------------------------------------------------------------------------------------------------------------------------------L---------IIG-T----------SYN--------A----------------------------------------R--LSW--Q-----------HKA------------------E-YM-GIPF-----K-K-RVKEVFY----SD-----P-----G--------Q--------------QI-IKG------------I---------I---A-----R-D-----------------I--D-H--T-S--GS-A--TVTAGGVGFSFANIRLKS-ERG--SGLNY-Q-------IEI-Y---------------V--------------------------------------------------------------------------------------------------------------------------------------------------------------------------------------------------------------------------------------------------------------------------------------------------------------------------------

>Sexi-betaREPAT3

---------MK-T-A--P-V----------F---L-V------V--L-------L-----A---I--I-----------C---A---------------V---------------------------------------------------------------------------H------C---K-----D-------------------------------------------------------------------------------------------------------------------------------------------------------------------------------------------------------L---------IVG-S----------SFN--------N----------------------------------------R--LIH--Q-----------EKA------------------E-YN-AIPL-----K-K-RVKEVFF----SD-----P-----G--------Q--------------QI-IKG------------I---------I---L-----R-D-----------------L--D-H--T-E--AI-P--SVTAGGVGFSYANIRLKS-ERG--SGLNY-Q-------IEF-Y---------------V--------------------------------------------------------------------------------------------------------------------------------------------------------------------------------------------------------------------------------------------------------------------------------------------------------------------------------

>Sexi-betaREPAT4

---------MR------F-V----------F---L-L------L--L-------V-----S---C--L-----------L---C---------------V---------------------------------------------------------------------------C-VP-VLC---S-----D-------------------------------------------------------------------------------------------------------------------------------------------------------------------------------------------------------M---------IVG-D----------TVH--------R----------------------------------------K--MIF--H-----------QRV------------------K-DF-AIPF-----K-K-RVKTLTF----SD-----P-----E--------K--------------RM-IKG------------I---------A---V-----I-D-----------------N--D-F--S-H--AS-A--NITDGGVGYSYVTVRMKS-QRH--HPLNF-E-------VEI-Y---------------V--------------------------------------------------------------------------------------------------------------------------------------------------------------------------------------------------------------------------------------------------------------------------------------------------------------------------------

>Sexi-betaREPAT5

---------MK-F----V-L----------V---L-F------A--C-------L-----F---V--I-----------V---T---------------C--------------------------------------------------------------------------------------D-----H-------------------------------------------------------------------------------------------------------------------------------------------------------------------------------------------------------L---------ILGNT----------NNN--------Q----------------------------------------N--MIY--H-----------TTA------------------H-YA-AIPF-----I-K-RVKNIFY----SG--------------------N--------------AV-INS------------I---------L---A-----Y-D-----------------N--K-H--T-N--AS-A--AVTAGGIGYTYVNLRLKS-ARG--KELDY-D-------IGI-Y---------------S--------------------------------------------------------------------------------------------------------------------------------------------------------------------------------------------------------------------------------------------------------------------------------------------------------------------------------

>Sexi-betaREPAT6

---------MK------C-V----------L-----------------------------I---T--VCLIL-------L---C---------------V---------------------------------------------------------------------------C------L---N--EGKD-------------------------------------------------------------------------------------------------------------------------------------------------------------------------------------------------------F---------IVG-T----------KAN--------N----------------------------------------L--LIS--T-----------QKL------------------K-YR-SIPL-----I-R-RDKDYTY----ID-----S-----K--------E--------------RI-IKG------------I---------I---A-----R-D-----------------L--S-R--T-D--TE-V--TVTSGGVGATNVTLHFQS-GRG--EELNY-L-------ILI-F---------------S---NIK--------------------------------------------------------------------------------------------------------------------------------------------------------------------------------------------------------------------------------------------------------------------------------------------------------------------------

>Sexi-betaREPAT7a

---------MK------V-V----------A---L-A------F--L-------L-----A---V--V-----------V---I---------------V---------------------------------------------------------------------------D------C---S-----H-------------------------------------------------------------------------------------------------------------------------------------------------------------------------------------------------------T---------FMG-T----------NVL--------R----------------------------------------P--LVY--H-----------HNA------------------E-YG-SKVF-----R-K-RVENLYY----SL-----P-----P---VPANAG--------------RS-IQG------------I---------L---A-----Y-D-----------------M--T-H--T-G--AT-A--NVTQGGLGFNFVNLRMKS-DRG--EKIHF-D-------VYI-Y---------------A--------------------------------------------------------------------------------------------------------------------------------------------------------------------------------------------------------------------------------------------------------------------------------------------------------------------------------

>Sexi-betaREPAT7b

---------MH------K-Y----------S--SI-A------I--I-------L-----A---V--V-----------V---A---------------S---------------------------------------------------------------------------E------C---G-----H-------------------------------------------------------------------------------------------------------------------------------------------------------------------------------------------------------S---------FLG-T----------NFL--------R----------------------------------------P--LVF--H-----------KEV------------------R-YP-AKVL-----Q-K-RIENFNF----TL-----P-----A---VPGVYS--------------RS-IQG------------I---------L---A-----Y-D-----------------L--T-H--S-G--AS-A--NVTKGGLGYNFVNMRMKS-DRY--KELHY-D-------IYI-Y---------------A--------------------------------------------------------------------------------------------------------------------------------------------------------------------------------------------------------------------------------------------------------------------------------------------------------------------------------

>Sexi-betaREPAT8a

---------MY----F-K-V----------I---L-V------L--S-------F-----V---Y--L-----------T---S---------------V---------------------------------------------------------------------------E------S---N-----D-------------------------------------------------------------------------------------------------------------------------------------------------------------------------------------------------------I---------RLG-F---------ATPY--------S----------------------------------------K--KIY--S-----------AIK------------------I----ANPA-----LWQ-REDDVLI----SS-----P-----N--------H--------------EL-IDA------------V---------Y---I-----T-D-----------------LRPE-K----D--GI-S--YIEAGGIGSKTVTIALKS-PSL-LRGYKF-A-------IEV-Y---------------A--------------------------------------------------------------------------------------------------------------------------------------------------------------Q----S-LHHGYGTNILPVQ---GAGYPRAG--LSGYG---------------------SQYP--GYPTAYSPVGVYP--N-SPIVG-------KK------------------------------------------------------------------

>Sexi-betaREPAT8b

---------MY----S-K-V----------L---V-T------L--F-------I-----F---Y--L-----------S---A---------------V---------------------------------------------------------------------------E------C---N-----D-------------------------------------------------------------------------------------------------------------------------------------------------------------------------------------------------------I---------HVG-Y---------LNSR--------S----------------------------------------K--KIY--S-----------ELK------------------E----AVPA-----LWK-RTDEITI----KA-----P-----S--------H--------------EV-IDS------------I---------H---V-----T-D-----------------LRDD-K----D--GE-A--YITGGGVGMTYVTIDLKS-PSI-LRGYKF-Q-------IEV-Y---------------G--------------------------------------------------------------------------------------------------------------------------------------------------------------S----R-SG--------F-----YDPIFGTG--NTNYGEG----------------QQ-QQYP-----------------------R-------KF------------------------------------------------------------------

>Sexi-gammaREPAT6

---------MK-----SV-F----------A---V-V------L--W-------AGC---V---A--V-----------G---Y---------AF----A---------------------------------------------------------------------------T------E---V-----DT-----S--V--------A---------------------------------------------------------------------------------------------D---------------------------------------------ET------------RS---------------------G---L---------ILG-S----------IGA--------D----------------------------------------D--RLL--S-----------NTN------------------HEVE-AALY-----V-V-QTQEVLI----RA-----V-----E--------Y--------------SN-ITS------------I---------R---V-----T-E-----------------V--G-T--SQE--PS-I--NILEGGLRSNFVRFLFRS-QKG--RAYSF-N-------VRI-H---------------G--------------------------------------------------------------------------------------------------------------------------------------------------------------T----------------------------------------------------------------------------------------------------------IGC----------------------------------------------------

>Sexi-gammaREPAT19

---------MK-----T--F----------A---V-V------L--L-------------V---A--I-----------A---A---------G-----V---------------------------------------------------------------------------Y------S---N-----EE-----Q--G--------A-------------------NMLNYDE-Q---SM--------------------------------EVSDQ-----------------------L---------------------------------------------AS------------RS---------------------S---Y---------GWG-V----------VGD--------D----------------------------------------D--RLI--S-----------STY------------------HEAY-DLPY-----L-I-NEQEVTY----RG-----N-----Q--------Y--------------TN-ISS------------I---------R---V-----S-S-----------------N--Y-G--NTT---Q-V--VLLDGGLNENYVTLGLKS-PRS--GGFGY-R-------IDI-F---------------A--------------------------------------------------------------------------------------------------------------------------------------------------------------T----------------------------------------------------------------------------------------------------------VNC--HNN-----------------------------------------------

>Sexi-gammaREPAT20

---------MK-----TV-L----------L---I-A------L--L-------A-----V---T--Y-----------V---Y---------AT----V---------------------------------------------------------------------------D-----------------L-----P--E--------A---------------------------------------------------------------------------------------------E---------------------------------------------FR---RPKVNASNLRL---------------------I---L---------GSK-S----------GS---------D----------------------------------------E--RLL--S-----------RTY------------------HYVE-GSHR-----I-A-HRQEILI----RL-----V-----D--------Y--------------AR-ITR------------V---------E---G-----W-P----------------------G--NPD--LE-L--TVLEGGFGQNFVKLGYNT-EEN--TFFSY-T-------LDV-Y---------------G--------------------------------------------------------------------------------------------------------------------------------------------------------------V----------------------------------------------------------------------------------------------------------LVCDD--------------------------------------------------

>Sexi-gammaREPAT21

---------MK-----I--L----------A---V-V------L--L-------------V---A--I-----------A---A---------G-----V---------------------------------------------------------------------------N------S---N-----EE----------------------------------------------------------------------------------------------------------------------------------------------------------------------------S---------------------S---Y---------GWG-V----------IGD--------N----------------------------------------D--RLI--S-----------STY------------------HEAY-DLPY-----L-I-NEQELTY----RG-----N-----Q--------Y--------------TN-ISS------------I---------R---V-----S-S-----------------N--Y-G--NTT---Q-V--VLLDGGLNENYVTLGLRS-PRS--GGFGC-R-------IDI-F---------------A--------------------------------------------------------------------------------------------------------------------------------------------------------------T----------------------------------------------------------------------------------------------------------VNC--HNN-----------------------------------------------

>Sexi-gammaREPAT22

---------MK-----TV-L----------L---I-A------L--L-------A-----V---T--Y-----------V---Y---------AT----V---------------------------------------------------------------------------D-----------------F-----S--E--------A---------------------------------------------------------------------------------------------E---------------------------------------------FR---RPELNASNLRL---------------------I---L---------GSK-S----------GS---------D----------------------------------------E--RLL--S-----------RTY------------------HYVE-GSHR-----I-A-HRQEILI----RL-----V-----D--------Y--------------AR-ITR------------V---------E---G-----W-P----------------------G--NPD--LE-L--TVLEGGFGQNFVKLGYVT-EEN--TFFSY-T-------LDV-Y---------------G--------------------------------------------------------------------------------------------------------------------------------------------------------------V----------------------------------------------------------------------------------------------------------LVCDD--------------------------------------------------

>Sexi-gammaREPAT23

---------MK-----I--L----------A---V-V------L--L-------------V---A--L-----------A---G---------G-----V---------------------------------------------------------------------------Y------S---N-----EE----------------------------------------------------------------------------------------------------------------------------------------------------------------------------S---------------------S---Y---------GWG-V----------IGD--------N----------------------------------------D--RLI--S-----------STY------------------HEAY-DLPY-----L-I-NEQELTY----RG-----N-----Q--------Y--------------TN-ISS------------I---------R---V-----S-S-----------------N--Y-G--NTT---Q-V--VLLDGGLNENYVTLGLRS-PRS--GGFGC-R-------IDI-F---------------A--------------------------------------------------------------------------------------------------------------------------------------------------------------T----------------------------------------------------------------------------------------------------------VNC--HNN-----------------------------------------------

>Sexi-gammaREPAT24

---------MK-----SV-L----------L---I-A------L--L-------A-----V---T--Y-----------V---Y---------AT----V---------------------------------------------------------------------------D-----------------L-----P--E--------A---------------------------------------------------------------------------------------------E---------------------------------------------FR---RPEVNASNLRL---------------------I---L---------GSK-S----------GS---------D----------------------------------------E--RLL--S-----------RTY------------------HYVE-GSHR-----I-A-HRQEILI----RL-----V-----D--------Y--------------AR-ITR------------V---------E---G-----W-P----------------------G--NPD--LE-L--TVLEGGFGQNFVKLGYVT-EEN--TFFSY-T-------LDV-Y---------------G--------------------------------------------------------------------------------------------------------------------------------------------------------------V----------------------------------------------------------------------------------------------------------LVCDD--------------------------------------------------

>Sexi-gammaREPAT25

---------MK-----I--L----------A---V-V------L--L-------------V---A--L-----------A---G---------G-----V---------------------------------------------------------------------------Y------S---N-----EE----------------------------------------------------------------------------------------------------------------------------------------------------------------------------S---------------------S---Y---------GWG-V----------IGD--------N----------------------------------------D--RLI--S-----------STY------------------HEAY-DLPY-----L-I-NEQELTY----RG-----N-----Q--------Y--------------TN-ISS------------I---------R---V-----S-S-----------------N--Y-G--NTT---Q-V--VLLDGGINENYVTLGLRS-PRS--GGFGC-R-------IDI-F---------------A--------------------------------------------------------------------------------------------------------------------------------------------------------------T----------------------------------------------------------------------------------------------------------VNC--HNN-----------------------------------------------

>Sexi-gammaREPAT26

---------MK-----SV-L----------L---I-A------L--L-------A-----V---T--Y-----------V---Y---------AT----V---------------------------------------------------------------------------D-----------------L-----P--E--------A---------------------------------------------------------------------------------------------E---------------------------------------------FR---RPEVNASNLRL---------------------I---L---------GSK-S----------VG---------D----------------------------------------D--RLL--S-----------KTY------------------HYVE-RSHR-----I-A-HRQEILI----RL-----V-----D--------Y--------------AR-ITR------------V---------E---G-----W-P----------------------G--NPD--LG-L--TVLEGGFGQNFVKLGYVT-EEN--MDFSY-T-------IDV-Y---------------G--------------------------------------------------------------------------------------------------------------------------------------------------------------V----------------------------------------------------------------------------------------------------------LVCDD--------------------------------------------------

>Sexi-gammaREPAT27

---------MK-----SV-L----------L---I-A------L--L-------A-----V---T--Y-----------V---C---------AT----V---------------------------------------------------------------------------D-----------------L-----P--E--------A---------------------------------------------------------------------------------------------E---------------------------------------------FR---RPEVNASNLRL---------------------I---L---------GSK-S----------GG---------E----------------------------------------D--RLL--S-----------RTY------------------HYVE-RSHR-----I-A-HRQEILI----RL-----V-----D--------Y--------------AR-ITR------------V---------E---G-----W-P----------------------G--YPD--LE-L--TVLEGGFGQNFVKLGYVT-EEN--MDFQY-T-------IDV-Y---------------G--------------------------------------------------------------------------------------------------------------------------------------------------------------V----------------------------------------------------------------------------------------------------------LVCDD--------------------------------------------------

>Sexi-gammaREPAT28

---------MK-----I--L----------A---V-V------L--L-------------V---A--I-----------A---A---------G-----V---------------------------------------------------------------------------Y------S---N-----EE-----Q--G--------A-------------------NMLNYDE-Q---TM--------------------------------EESDQ-----------------------L---------------------------------------------SS------------RS---------------------S---F---------GWG-V----------VGI--------D----------------------------------------D--RLI--S-----------RTY------------------HEAY-GYQY-----L-I-NEQELTY----RG-----N-----Q--------Y--------------TN-ISS------------I---------R---V-----Y-S-----------------N--Y-R--NTT---E-I--VLLDGGLNENYVTFGFRS-PRS--EGFGC-R-------IDI-Y---------------A--------------------------------------------------------------------------------------------------------------------------------------------------------------T----------------------------------------------------------------------------------------------------------INC--HNN-----------------------------------------------

>Sexi-gammaREPAT7

---------MK-----VF-L--------------V-I------L--S-------------A---A--V-----------V---T---------AG----V---------------------------------------------------------------------------T------T---N-----ER-----E--LGEVPEVP-AVPEITEVPKVPENKKLPLSD-LDKNE-LD---IPNLPEEDDLNVPDLPEDTDLDVPDGETKGDLDDEADL-----------------------A---------------------------------------------TA------------RN---------------------N---L---------SFG-T----------FGS--------N----------------------------------------D--RLM--S-----------RIT------------------HNVA-AASL-----L-I-HDQEITF----RG-----V-----R--------G--------------TN-ITA------------I---------R---V-----N-Q-----------------V--G-S--NNP---T-A--TRIAGGLGEEFVTIRIMS-GRG--RSFSY-T-------ILV-Y---------------A--------------------------------------------------------------------------------------------------------------------------------------------------------------E----------------------------------------------------------------------------------------------------------INC--SNN-----------------------------------------------

>Sexi-gammaREPAT29

---------MK-----I--F----------A---V-V------L--L-------------V---V--I-----------A---A---------G-----V---------------------------------------------------------------------------Y------S---N-----EE-----Q--G--------A-------------------NMLNYEE-KDPNIK--------------------------------MESDQ-----------------------M---------------------------------------------TT------------RP---------------------H---V----------WG-S----------YGP--------N----------------------------------------D--RLI--S-----------RTV------------------HVIS-ALTL-----F-T-NDQEVTY----RG-----H-----E--------K--------------TN-ITS------------I---------V---V-QENGL-N-----------------G--F-S----T---Q-V--DLLDGGLYHNFVTIGLKA-GRS--YTINK-D-------IRI-Y---------------A--------------------------------------------------------------------------------------------------------------------------------------------------------------T----------------------------------------------------------------------------------------------------------INC--FNN-----------------------------------------------

>Sexi-gammaREPAT30

---------MK-----DV-L----------L---I-A------L--L-------A-----V---T--L-----------G---C---------AT----A---------------------------------------------------------------------------D------S---G-----DL-----P--E--------P---------------------------------------------------------------------------------------------A---------------------------------------------AS---GSR-G--TDRS---------------------Q---L---------AIG-E----------IG---------S----------------------------------------S--RIL--S-----------STT------------------HRVP-SENG-----I-A-HTQDVLI----RL-----V-----E--------Y--------------AN-ITA------------I---------RG--I-----T-S----------------------F--ISA--AE-I--FLLEGGLGHNYVKLGFVT-EKG--FGYNY-A-------ITI-Y---------------G--------------------------------------------------------------------------------------------------------------------------------------------------------------D----------------------------------------------------------------------------------------------------------MVCKE--------------------------------------------------

>Sexi-gammaREPAT8

---------MK-----SI-F----------V---S-I------L--L-------AGC---V---A--V-----------G---Y---------AS----A---------------------------------------------------------------------------T------E---V-----EN-----S--I--------A---------------------------------------------------------------------------------------------E---------------------------------------------ET------------RS---------------------G---L---------SLG-S----------IGS--------G----------------------------------------D--RLL--S-----------NTN------------------HAVE-AALY-----V-M-QMQEVLI----RT-----V-----E--------Y--------------SK-ITS------------I---------R---V-----T-E-----------------V--G-T--SQN--AI-V--DIIDGGLTKNFVKLLFRS-EKG--RAYSF-N-------VRI-H---------------G--------------------------------------------------------------------------------------------------------------------------------------------------------------T----------------------------------------------------------------------------------------------------------IEC----------------------------------------------------

>Sexi-gammaREPAT9

---------MK-----TY-L----------A---V-T------L--L-------------V---A--V-----------V---T---------AG----V---------------------------------------------------------------------------T------T---N-----ER-----D--L--------AV--------VPEDKKLHLSD-F--------------IEREDLYVPDSPENTDLDVRELEIKGEEDEETNL-----------------------A---------------------------------------------TT------------RN---------------------N---L---------SLG-S----------FGS--------N----------------------------------------D--RLM--S-----------RIT------------------HNVA-AASL-----L-I-HDQEITF----RG-----V-----R--------G--------------TN-ITA------------I---------R---V-----S-Q-----------------V--G-S--NSP---T-A--TRIAGGLGSEFVTIRIRS-DRG--RAFSY-T-------ILV-Y---------------A--------------------------------------------------------------------------------------------------------------------------------------------------------------E----------------------------------------------------------------------------------------------------------INC--SNN-----------------------------------------------

>Sexi-gammaREPAT14

---------MK-----I--L----------A---V-V------L--L-------------V---A--I-----------A---A---------G-----V---------------------------------------------------------------------------Y------S---N-----EE-----Q--G--------A-------------------NMLK--E-Q---SM--------------------------------EKSDQ-----------------------L---------------------------------------------AS------------RS---------------------S---Y---------GWG-V----------VGD--------D----------------------------------------D--RLI--S-----------STY------------------HEAY-DLPY-----L-I-NEQELTY----RG-----N-----Q--------Y--------------TN-ISS------------I---------R---V-----S-S-----------------N--Y-G--NTT---Q-V--VLLDGGLNENYVTLGLRS-PRS--GGFGY-R-------IDI-F---------------A--------------------------------------------------------------------------------------------------------------------------------------------------------------T----------------------------------------------------------------------------------------------------------VNC--HNN-----------------------------------------------

>Sexi-gammaREPAT15

---------MK-----I--L----------A---V-V------L--L-------------V---A--I-----------A---A---------G-----V---------------------------------------------------------------------------Y------S---N-----EE-----Q--G--------A-------------------NMLK--------------------------------------------SDQ-----------------------L---------------------------------------------AS------------RS---------------------S---Y---------GWG-V----------VGD--------D----------------------------------------D--RLI--S-----------STY------------------HEAY-DLPY-----L-I-NEQELTY----RG-----N-----Q--------Y--------------TN-ISS------------I---------R---V-----S-S-----------------N--Y-G--NTT---Q-V--VLLDGGLNENYVTLGLRS-PRS--GGFGY-R-------IDI-F---------------A--------------------------------------------------------------------------------------------------------------------------------------------------------------T----------------------------------------------------------------------------------------------------------VNC--HNN-----------------------------------------------

>Sexi-gammaREPAT16

---------MK-----SV-L----------L---I-A------L--L-------A-----V---T--Y-----------V---Y---------AA----V---------------------------------------------------------------------------D-----------------L-----S--G--------A---------------------------------------------------------------------------------------------E---------------------------------------------FK---RPELNASKLRL---------------------I---L---------GSK-S----------GG---------D----------------------------------------E--RLL--S-----------RTY------------------HYVE-GSHR-----I-A-HRQEILI----RL-----V-----D--------Y--------------AR-ITR------------V---------E---G-----W-P----------------------G--NPD--LE-L--TVLEGGLGQSFIKLGYVT-EEN--TFFSY-T-------LDV-Y---------------G--------------------------------------------------------------------------------------------------------------------------------------------------------------V----------------------------------------------------------------------------------------------------------LVCDD--------------------------------------------------

>Sexi-gammaREPAT17

---------MK-----I--L----------A---V-V------L--L-------------V---A--I-----------A---A---------G-----V---------------------------------------------------------------------------Y------S---N-----EE-----Q--G--------A-------------------NMLNYDE-Q---SI--------------------------------EESDQ-----------------------L---------------------------------------------AS------------RS---------------------S---F---------GWG-V----------VGD--------D----------------------------------------D--RLI--S-----------STY------------------HEAY-DLPY-----L-I-NEQEVTY----RG-----N-----Q--------Y--------------TN-ISS------------I---------R---V-----S-S-----------------N--Y-G--NTT---Q-V--VLQGGGLNENYVTLGLRS-PRS--GGFGC-R-------IDI-F---------------A--------------------------------------------------------------------------------------------------------------------------------------------------------------T----------------------------------------------------------------------------------------------------------VNC--HNN-----------------------------------------------

>Sexi-gammaREPAT18

---------MK-----I--F----------A---V-V------L--L-------------V---A--I-----------A---A---------G-----V---------------------------------------------------------------------------Y------S---N-----EE-----Q--G--------A-------------------NMLNYDK-K---SM--------------------------------EKSDQ-----------------------L---------------------------------------------AS------------RS---------------------S---Y---------GWG-V----------VGD--------D----------------------------------------D--RLI--S-----------STY------------------HEAY-DLPY-----L-I-NEQEVTY----RG-----N-----Q--------Y--------------TN-ISS------------I---------R---V-----S-S-----------------N--Y-G--NTT---Q-V--VLLGGGINENYVTLGLRS-PRS--GGFGC-R-------IDI-F---------------A--------------------------------------------------------------------------------------------------------------------------------------------------------------T----------------------------------------------------------------------------------------------------------VNC--HNN-----------------------------------------------

>Sexi-gammaREPAT1

---------MK-----SL-I----------L---V-A------V--L-------A---ALA---V--C-----------N---D---------A---------------------------------------------------------------------------------------------------A-----A-LQ--------E---------------------------------------------------------------------------------------------P---------------------------------------------AF------------RA---------------------N---L---------YQG-A----------IRP--------G----------------------------------------D--RLL--H-----------RNY------------------YYKS-PIAN-----A-V-QYQDITY----RG-----N-----S--------S--------------TR-ISY------------I---------Q---V-----T-E-----------------V--G-Y--TQW--GI-P--SLRAGGVNFNHATIRLTS-QRG--YGYYY-R-------VEI-W---------------G--------------------------------------------------------------------------------------------------------------------------------------------------------------R-----------------------------------------------------------------------------------------------------------------------------------------------------------------

>Sexi-gammaREPAT2

---------MK-----SL-I----------L---V-A------V--L-------A---VLT---V--C-----------N---D---------A---------------------------------------------------------------------------------------------------A-----T-VH--------E---------------------------------------------------------------------------------------------P---------------------------------------------AF------------RA---------------------N---L---------YQG-S----------IRP--------G----------------------------------------D--RLL--H-----------RNY------------------YVKN-AIPN-----A-V-QYQDINY----RG-----N-----S--------T--------------TR-ISY------------I---------Q---A-----T-E-----------------V--G-Y--SQR--GI-P--SLRSGGVNFNHATIRLTS-QRG--MGYYY-R-------VEI-W---------------G--------------------------------------------------------------------------------------------------------------------------------------------------------------R-----------------------------------------------------------------------------------------------------------------------------------------------------------------

>Sexi-gammaREPAT3

---------MK-----SL-I----------L---V-A------V--L-------A---ALA---V--C-----------N---D---------A---------------------------------------------------------------------------------------------------A-----A-LQ--------E---------------------------------------------------------------------------------------------P---------------------------------------------AF------------RA---------------------N---L---------YQG-S----------IRP--------G----------------------------------------D--RLL--H-----------SNY------------------YVKN-AIPN-----A-S-QWQEINY----NG-----S-----S--------R--------------TR-ISY------------I---------Q---V-----T-E-----------------V--G-Y--SQR--GI-P--SLMAGGVNYNYARIRLTT-QRG--MGYYY-R-------VEI-W---------------G--------------------------------------------------------------------------------------------------------------------------------------------------------------R-----------------------------------------------------------------------------------------------------------------------------------------------------------------

>Sexi-gammaREPAT4

---------MK-----SL-I----------L---V-A------V--L-------A---ALA---V--C-----------N---D---------A---------------------------------------------------------------------------------------------------A-----A-LQ--------E---------------------------------------------------------------------------------------------P---------------------------------------------AF------------RA---------------------N---L---------YQG-A----------IRP--------G----------------------------------------D--RLL--H-----------SNY------------------YVKN-AIPN-----A-S-QWQDINY----NG-----S-----S--------R--------------TR-ISY------------I---------Q---V-----T-E-----------------V--G-Y--SQR--GI-P--SLMAGGVNYNYARIRLTT-QRG--MGYYY-R-------VEI-W---------------G--------------------------------------------------------------------------------------------------------------------------------------------------------------R-----------------------------------------------------------------------------------------------------------------------------------------------------------------

>Sexi-gammaREPAT5

---------MK-----SL-I----------L---V-A------V--L-------A---VLT---V--C-----------N---D---------A---------------------------------------------------------------------------------------------------A-----T-VH--------E---------------------------------------------------------------------------------------------P---------------------------------------------AF------------RA---------------------N---L---------YQG-A----------IRP--------G----------------------------------------D--RLL--H-----------SNY------------------YVRN-PVPN-----I-S-QSQEVNY----RG-----N-----S--------T--------------TR-ISY------------I---------Q---A-----T-E-----------------V--G-Y--SQR--GI-P--SLVGGGVNYNFARIRLTT-QRG--MGYYY-R-------VEI-W---------------G--------------------------------------------------------------------------------------------------------------------------------------------------------------R-----------------------------------------------------------------------------------------------------------------------------------------------------------------

>Sexi-gammaREPAT10

---------MK-----SL-L----------L---V-V------L--L-------A-----V---A--A-----------S---A---------A---------------------------------------------------------------------------------------------------IVSQDIA--E--------A---------------------------------------------------------------------------------------------PEAVVEYEEVLYIDE---------------AEVAEI-EAGSRSVEIES------------RR---------------------N---L---------SVG-T----------IGN--------S-------------------------------------------RLL--V-----------TSQ------------------HIRG-AIAN-----T-I-IVQNITF----NF-----A-----A--------S--------------IR-ISA------------I---------R---V-----T-R-----------------V--G-A--SQN--AT-P--SIASGGLNRNFVTIRLQS-ARG--RGYNY-R-------IQI-Y---------------G--------------------------------------------------------------------------------------------------------------------------------------------------------------R-----------------------------------------------------------------------------------------------------------------------------------------------------------------

>Sexi-gammaREPAT11

---------MK-----GL-L----------L---V-V------L--L-------A-----A---A--A-----------S---A---------A---------------------------------------------------------------------------------------------------IVSQDIA--E--------A---------------------------------------------------------------------------------------------PEAIVTYENILYSDD---------------AEVAEI-EAGSRSVEIET------------RR---------------------N---L---------SVG-T----------IGN--------S-------------------------------------------RLL--V-----------SSQ------------------HIRG-AIAN-----T-I-IVQNITF----NF-----A-----A--------S--------------IR-ISA------------I---------R---V-----T-R-----------------V--G-A--SQN--AT-P--SIVSGGLNRNFVTIRLQS-ARG--RGYNY-R-------IQI-Y---------------G--------------------------------------------------------------------------------------------------------------------------------------------------------------R-----------------------------------------------------------------------------------------------------------------------------------------------------------------

>Sexi-gammaREPAT12

---------MK-----SL-L----------L---V-V------L--L-------A-----A---A--A-----------S---A---------A---------------------------------------------------------------------------------------------------IVSQDIA--E--------A---------------------------------------------------------------------------------------------PEAVVEYESILYSDE---------------AEVSEV-EAGSRSVDIET------------RR---------------------N---L---------SVG-T----------IGN--------S-------------------------------------------RLL--V-----------TSQ------------------HIRG-AIAN-----T-I-IVQNITF----NF-----A-----A--------S--------------IR-ISA------------I---------R---V-----M-R-----------------V--G-A--SQN--AT-P--SIASGGLNRNFVTIRLQS-ARG--RGYNY-R-------IQI-Y---------------G--------------------------------------------------------------------------------------------------------------------------------------------------------------R-----------------------------------------------------------------------------------------------------------------------------------------------------------------

>Sexi-gammaREPAT13

---------MK-----SL-L----------L---V-V------M--L-------A-----A---A--A-----------S---A---------A---------------------------------------------------------------------------------------------------IVSQDIA--E--------A---------------------------------------------------------------------------------------------PDAIVEYEEILYIDE---------------AEVAAV-EALSGGVEIET------------RR---------------------N---L---------SVG-T----------IGN--------S-------------------------------------------RLL--T-----------TRY------------------QARG-AVPN-----T-V-IVQNIVF----NF-----P-----A--------S--------------IR-IAA------------I---------R---V-----T-R-----------------M--G-A--SQN--TT-P--SIVYGGLNRNFVTIRLQS-ARG--RGYNY-R-------IQI-Y---------------G--------------------------------------------------------------------------------------------------------------------------------------------------------------R-----------------------------------------------------------------------------------------------------------------------------------------------------------------

>Sfru-alphaREPAT1

---------MR-----SF-V----------I---I-T------V--L------SA-----L---A--A------CY-G-A---A---------------V----LPR---------G------------------K--------E--N-------YTED--KVIQKYNKD--DDGL------N---R---------------------------------------------------------------------------------------------------------------------------------------------------------------------------------------------------------------------------I----------TML--------GEQMRY-----V-ES-----------------------D--DDLVFL--H-----------E--------------------FLKN-AVPD-----A-I-ENVDILY----YA-----P-----P--------S--------------TV-IKT------------I---------L---V-----T-S-----------------S--P-P--PSS--VF-P--V----WHLDNYISVRISS-QRG--CALNG-T-------VRF-Y---------------G-TL-----------------------------------------------------------------------------------------------------------------------------------------------------------------------------------------------------------------------------------------------------------------------------------------------------------------------------

>Sfru-alphaREPAT10

---------MR-----NF-I---------II---I-A------V--L------TA-----L---V--A------CY-G-D---A---------------I----TES---------L------------------E--------E--M-------------SLV--Y--DEEDTSM------N---G---------------------------------------------------------------------------------------------------------------------------------------------------------------------------------------------------------------------------R----------LLS--------SAD---V---DS-RN--------------------TTW------VVI--S-----------EWQ------------------LSAP-AVAG-----Q-V-QRVGVMY----EG-----A-----A--------N--------------VT-INR------------L---------T---V-----T-Y-----------------T--E-T--PAA--VD-R--S----PLSTNVMNVMIRS-HAG--QQVNA-T-------VTI-L---------------------------------------------------------------------------------------------------------------------------------------------------------------R--------------R----------------------------------------------------------------------------------------------------------------------------------------------------------------A

>Sfru-alphaREPAT11

---------MR-----SF-I----------I---I-T------V--L------SA-----L---A--A------CY-A-A---S---------------V----VPS---------E------------------E--------E--M----------V--IVV--Y--D--ENFP------N---G---------------------------------------------------------------------------------------------------------------------------------------------------------------------------------------------------------------------------T----------IVT--------KAE---L---DS-RFN--------SG------------------IVI--A-----------SWM------------------ANAP-AIAR-----Q-S-QVTTLQY----FG-----D-----------------------------EV-VRR------------IVYYNWGN-PS---M-----T-Y------------------------------------T----PLLSNNMQITITT-RIG--VELRS-T-------VEL-Q---------------------------------------------------------------------------------------------------------------------------------------------------------------L--------------------------------------------------------------------------------------------------------------------------------------------------------------------------------

>Sfru-alphaREPAT12

---------MK-----TF-I----------V---I-A------L--L------SA-----L---A--A------CC-SAA-------------------V----LPN---------E------------------E-D----MSG--G----------V--TIV--Y--D--DESP------N---G---------------------------------------------------------------------------------------------------------------------------------------------------------------------------------------------------------------------------R----------VMS--------RND---F---GS-IL---------------DNYW----------VLL--S-----------EAV------------------INVP-AVPG-----K-V-QKAGMTY----QA-----P-----P--------G--------------VK-ISR------------I---------S---I-----TLY-----------------T--N-L--PDD--VY-K--S----PVGGEYIEALITS-RPG--DVING-T-------IAA-Y---------------------------------------------------------------------------------------------------------------------------------------------------------------R--------------R-------------------------------------------------------------------------------------------------------------------------------------------------YN-E------------

>Sfru-alphaREPAT13

---------MR-----SF-I----------I---I-T------V--L------SA-----L---A--A------CY-G-A-------------------N----LPS---------E------------------E------FSK--G----------F--TIV--F--D--DSAP------N---G---------------------------------------------------------------------------------------------------------------------------------------------------------------------------------------------------------------------------R----------TMS--------PVD---M---KI-SW---------------EG-W----------TLL--S-----------QGT------------------YIVP-AIAG-----E-V-QSITKVF----QP-----E-----A--------D--------------VR-ISG------------F---------S---I-----A-Y-----------------N--E-Y--ALP--SY----T----PLPNNAMNITLTS-IPG--EGIHC-F-------ISI-Y---------------------------------------------------------------------------------------------------------------------------------------------------------------S-------------------------------------------------------------------------------------------------------------------------------------------------------------------Q------------

>Sfru-alphaREPAT14

---------MR-----SF-I----------I---I-T------V--L------SA-----L---A--A------CY-A-AA------------------V----LPS---------E------------------E------MSE--G----------Y--SLV--F--D--DMSP------E---G---------------------------------------------------------------------------------------------------------------------------------------------------------------------------------------------------------------------------R----------IVS--------NAE---L---NA-RS---------------SD-W----------ILI--S-----------EGV------------------VSAP-AVLG-----Q-V-QRVGTWH----QA-----D-----L--------G--------------VR-IER------------L---------A---L-----A-Y-----------------G--A-F--PAD--VQ-H--T----PLGSNFLEVRIIS-APS--QRVSA-T-------ISA-Y---------------------------------------------------------------------------------------------------------------------------------------------------------------R--------------R---------------------------------------------------------------------------------------------------------------------------------------------------AF------------

>Sfru-alphaREPAT15

---------MK-----TF-I----------T---I-A------L--L------SA-----L---A--A------CC-S-A---S---------------V----FPS---------K------------------E--------E--L-------ARGV--TVV--ENAD--EAEL------Q---K---------------------------------------------------------------------------------------------------------------------------------------------------------------------------------------------------------------------------FLS--------THP--------NRQ--------A-LN-----------------------E--S--VVL--F-----------ETT------------------FFVP-AIPG-----E-V-VTHRIEY----IG-----G-----P--------N--------------TK-MTG------------S---------Q---R-----T-H-----------------L--P-PL-PAN--VH-S--S----GWLTNHLVKTFTS-QVG--GNIIG-N-------VKF-Y---------------G--------------------------------------------------------------------------------------------------H----------------------------------------KYGY---------------------------------------------------------------------------------------------------------------------------------------------------------------------------------

>Sfru-alphaREPAT16

---------MK-----TF-I----------I---I-A------L--L------SA-----L---A--A------CC-S-A---S---------------V----LPS---------E---------------------------E--M-------AEGI--TVV--ENAD--EAEL------Q---K---------------------------------------------------------------------------------------------------------------------------------------------------------------------------------------------------------------------------FLS--------THP--------GRL--------V-E------------------------E--H--V----F-----------ESH------------------FSVP-AIPG-----E-V-VTHRIEY----IG-----G-----P--------D--------------TV-MTR------------T---------F---R-----S-M-----------------F--P-PL-PVE--VT-T--E----GWLTNRLVMTLTS-EVG--GSLVG-N-------VIF-Y---------------G--------------------------------------------------------------------------------------------------H----------------------------------------KYGF---------------------------------------------------------------------------------------------------------------------------------------------------------------------------------

>Sfru-alphaREPAT17

---------MK-----TF-I----------I---I-A------L--L------SA-----L---A--A------CC-S-A---S---------------V----LPS---------E------------------K--------E--M-------AEGI--TVV--ENAD--EAEL------Q---K---------------------------------------------------------------------------------------------------------------------------------------------------------------------------------------------------------------------------FLS--------THP--------GHP--------L-E------------------------E--T--V----F-----------ESH------------------FSVP-AIPG-----K-V-VTHRIEY----IG-----G-----P--------D--------------TV-MTR------------T---------F---R-----S-M-----------------F--P-PL-PAE--VT-T--E----GWLTNRLVMTLTS-EVG--GSLVG-N-------VQF-F---------------G--------------------------------------------------------------------------------------------------Y----------------------------------------KH-FS--------------K-----------------------------------------------------------------------------------------------------------------------------------------------------------------

>Sfru-alphaREPAT18

---------MK-----TF-I----------I---I-A------L--L------SA-----L---A--A------CC-S-A---S---------------V----LPS---------E------------------K--------E--M-------AEGI--TVV--ENAD--EAEL------Q---K---------------------------------------------------------------------------------------------------------------------------------------------------------------------------------------------------------------------------FLS--------THP--------ERQ--------A-A------------------------E--R--VFL--F-----------ESH------------------FSVP-AIPG-----E-T-VTHRIEY----IG-----G-----P--------D--------------TV-LTG------------T---------H---R-----T-S-----------------L--P-PL-PAE--VT-T--E----GWLTNRLVMTLTS-EVG--GSLVG-S-------VQF-F---------------G--------------------------------------------------------------------------------------------------Y----------------------------------------KYGF---------------------------------------------------------------------------------------------------------------------------------------------------------------------------------

>Sfru-alphaREPAT19

---------MK-----TF-I----------I---I-A------L--L------SA-----L---A--A------CC-S-A---S---------------V----LPS---------K------------------K--------E--M-------AEGL--TVV--ENAD--EAEL------Q---K---------------------------------------------------------------------------------------------------------------------------------------------------------------------------------------------------------------------------LLS--------THP--------ERQ--------A-A------------------------E--R--VFL--F-----------ETD------------------FSVP-AIPG-----E-T-VTHRIEY----IG-----G-----P--------D--------------TV-LTG------------T---------H---R-----T-S-----------------L--P-PL-PAE--VT-T--E----GWLTNRLVMTLTS-EVG--GSLVG-N-------VKF-L---------------G--------------------------------------------------------------------------------------------------Y----------------------------------------KYGR---------------------------------------------------------------------------------------------------------------------------------------------------------------------------------

>Sfru-alphaREPAT2

---------MR-----SF-I----------I---I-T------V--L------SA-----L---A--A------CY-G-A---A---------------V----LPS---------G------------------D--------E--N-------YTED--QVIQKYNRD--DDVQ------N---R---------------------------------------------------------------------------------------------------------------------------------------------------------------------------------------------------------------------------I----------TML--------GEQKRY-----V-ES-----------------------D--DDLVFL--H-----------E--------------------FLKN-AVPD-----A-I-ENVDILY----YA-----Q-----P--------N--------------TV-IKT------------I---------L---V-----T-S-----------------S--P-P--PSS--VF-P--V----WHLDNYISVRISS-QRG--CALNG-T-------VRF-Y---------------G-TL-----------------------------------------------------------------------------------------------------------------------------------------------------------------------------------------------------------------------------------------------------------------------------------------------------------------------------

>Sfru-alphaREPAT20

---------MK-----TF-I----------I---I-A------L--L------SA-----L---A--A------CC-S-A---S---------------V----LPS----------------------------E--------E--M-------AGGI--TVV--ENAD--EVEL------Q---K---------------------------------------------------------------------------------------------------------------------------------------------------------------------------------------------------------------------------FLS--------T-----------RQ--------A-SN-----------------------E--T--ELL--F-----------ESN------------------FFVP-AIPG-----E-A-VVHRIEY----IG-----A-----P--------N--------------TK-ITT------------T---------H---R-----N-H-----------------F--P-PL-PSL--VH-T--A----GWLTNNLVMTLTS-QVG--GSMLG-S-------VKI-Y---------------G--------------------------------------------------------------------------------------------------Y----------------------------------------VHGN---------------------------------------------------------------------------------------------------------------------------------------------------------------------------------

>Sfru-alphaREPAT21

---------MK-----TF-I----------I---I-A------L--L------SA-----L---A--A------CC-S-A---S---------------V----LPS---------K------------------K--------E--M-------AEGL--TVV--ENAD--EAEL------Q---K---------------------------------------------------------------------------------------------------------------------------------------------------------------------------------------------------------------------------LLS--------THP--------ERQ--------A-A------------------------E--R--VFL--F-----------ETD------------------FSVP-AIPG-----E-T-VTHRIEY----IG-----G-----P--------D--------------TV-LTG------------T---------H---R-----T-S-----------------L--P-PL-PAE--VT-T--E----GWLTNRLVMTLTS-EVG--GSLVG-N-------VKF-L---------------G--------------------------------------------------------------------------------------------------Y----------------------------------------KYGR---------------------------------------------------------------------------------------------------------------------------------------------------------------------------------

>Sfru-alphaREPAT22

---------MK-----TF-I----------I---I-A------L--L------SA-----L---A--A------CC-S-A---S---------------V----LPS----------------------------E--------E--M-------AGGI--TVV--ENAD--EAEL------Q---K---------------------------------------------------------------------------------------------------------------------------------------------------------------------------------------------------------------------------FLS--------T-----------RQ--------A-SN-----------------------E--T--ELL--F-----------ESN------------------FFVP-AIPG-----E-A-VVHRIEY----IG-----A-----P--------N--------------TK-ITT------------T---------H---R-----N-H-----------------F--P-PL-PSL--VH-T--A----GWLTNNLVMTLTS-QVG--GSMLG-S-------VKI-Y---------------G--------------------------------------------------------------------------------------------------Y----------------------------------------VHGN---------------------------------------------------------------------------------------------------------------------------------------------------------------------------------

>Sfru-alphaREPAT23

---------MK-----TF-I----------I---I-A------L--L------SA-----L---A--A------CC-S-A---S---------------V----LPS----------------------------E--------E--M-------AGGI--TVV--ENAD--EAEL------Q---K---------------------------------------------------------------------------------------------------------------------------------------------------------------------------------------------------------------------------LLS--------THP--------ERQ--------A-A------------------------E--R--VFL--F-----------ETD------------------FSVP-AIPG-----E-T-VTHRIEY----IG-----G-----P--------D--------------TV-LTG------------T---------H---R-----T-S-----------------L--P-PL-PAE--VT-T--E----GWLTNRLVMTLTS-EVG--GSLVG-N-------VKF-L---------------G--------------------------------------------------------------------------------------------------Y----------------------------------------KYGR---------------------------------------------------------------------------------------------------------------------------------------------------------------------------------

>Sfru-alphaREPAT24

---------MK-----TF-I----------T---I-A------L--L------SA-----L---A--A------CC-S-A---S---------------V----LPS---------K------------------E--------E--L-------ARGV--TVV--ENAD--EAEL------Q---K---------------------------------------------------------------------------------------------------------------------------------------------------------------------------------------------------------------------------FLS--------T-----------RQ--------A-SN-----------------------E--T--ELL--F-----------ESN------------------FFVP-AIPG-----E-A-VVHRIEY----IG-----A-----P--------N--------------TV-ITT------------T---------H---R-----N-H-----------------F--Q-PL-PSL--VH-T--A----GWLTNNLVMTLTS-QVG--GSMLG-S-------VKI-Y---------------G--------------------------------------------------------------------------------------------------Y----------------------------------------VHGN---------------------------------------------------------------------------------------------------------------------------------------------------------------------------------

>Sfru-alphaREPAT25

---------MK-----SF-I----------I---I-A------V--L------SA-----L---A--A------CH-G-A---A---------------V----PPSEADMFSSFIQ------------------E--------Q--M-------LNGF--TVV--HSEE--DAAL------N---R---------------------------------------------------------------------------------------------------------------------------------------------------------------------------------------------------------------------------V----------TLS--------KGE--------T-RD-----------------------D--H--VLL--F-----------ERA------------------CYVP-AVPN-----D-V-VRCDVVY----YG-----E-----P--------N--------------VT-IWS------------V---------S---IS----F-D-----------------W--P-P--PAN--FN-I--T----GVQTNVGRVRLVS-QRG--EHLHT-I-------VRM-F---------------GKIT-----------------------------------------------------------------------------------------------E----------------------------------------DNNES--------------------------------------------------------------------------------------------------------------------------------------------------------------------------------

>Sfru-alphaREPAT26

---------MK-----SF-I----------I---I-A------V--L------SA-----L---A--A------CH-G-A---A---------------V----PPSEADMFSSFIQ------------------E--------Q--M-------LNGF--TVV--HSEE--DAAL------N---R---------------------------------------------------------------------------------------------------------------------------------------------------------------------------------------------------------------------------V----------TLS--------KGE--------T-RD-----------------------D--D--VLF--F-----------ERL------------------CYVP-AVPN-----D-V-VRCDVVY----YG-----E-----P--------N--------------VR-ISR------------V---------S---IS----F-S-----------------W--P-P--AAN--TS-I--T----GLSTNVVRVRFVS-QRD--EHLHL-T-------VRM-F---------------GKIT-----------------------------------------------------------------------------------------------E----------------------------------------DNNES--------------------------------------------------------------------------------------------------------------------------------------------------------------------------------

>Sfru-alphaREPAT27

---------MK-----SF-I----------I---I-A------V--L------SA-----L---A--A------CH-G-A---A---------------V----PPSEADMFSSFIQ------------------E--------Q--M-------LNGF--TVV--HSEE--DAAL------N---R---------------------------------------------------------------------------------------------------------------------------------------------------------------------------------------------------------------------------V----------TLS--------KGE--------T-RD-----------------------D--H--FIF--R-----------E-S------------------CFVP-AIPN-----Q-S-VNCEVVY----EG-----E-----P--------N--------------VR-IRR------------V---------T---VT----S-I-----------------R--P-S--PAY--IY-I--S----GFSTNVGLVRLES-QIG--GPLQS-V-------VRL-F---------------G----------------------------------------------------------------------------------------------------------------------------------------------EL--------------------------------------------------------------------------------------------------------------------------------------------------------------------------------

>Sfru-alphaREPAT28

---------MK-----SF-I----------I---I-A------V--L------SA-----L---A--A------CH-G-A---A---------------V----PPSEADMFSSFIQ------------------E--------Q--M-------YPNS--TVE--YNEE--D--Q------N-------------------------------------------------------------------------------------------------------------------------------------------------------------------------------------------------------------------------------I----------TFS--------KGE--------V-GE-----------------------D--Q--ILL--L-----------TRE------------------IHVP-AEPG-----K-T-QYANLIH----RS-----P-----A--------N--------------IR-ITR------------V---------R--WRY----G-K-----------------L--E-K--PAD--MG-I--E----GMETRVVHISYTS-FEG--DGIHV-F-------LEV-F---------------GRIV-----------------------------------------------------------------------------------------------E----------------------------------------N------------------------------------------------------------------------------------------------------------------------------------------------------------------------------------

>Sfru-alphaREPAT3

---------MK-----GF-F----------I---I-A------V--L------SA-----L---I--A------SY-D-A---F---------------V----LPN---------R--LMSGS---------VHD--------G--F-------------TLV--Y--D--DTSV------N---G---------------------------------------------------------------------------------------------------------------------------------------------------------------------------------------------------------------------------T----------IMT--------NAD---V---EA-KN--------------------DEW------RLL--Y-----------EVV------------------CTAP-AAFG-----R-I-KNSFVTY----QG-----P-----T--------T--------------TV-VGR------------V---------T---I-----T-P-----------------D--I-H--PAN--VF-H--T----ALGTNYMEVRLMS-NIS--QPIEA-T-------VRM-Y---------------------------------------------------------------------------------------------------------------------------------------------------------------Q--------------R------------------------------------------------------------------------------------------------------------------NGAGKAFGEPAFHLMIFVLIYFGISSF-------------------T

>Sfru-alphaREPAT4

---------MR-----SF-I----------I---I-T------V--L------SA-----L---A--A------SY-C-T---A---------------D----VWS---------E------------------K--------E--MQER-------Y--NIM--Y--N--EEST------N---G---------------------------------------------------------------------------------------------------------------------------------------------------------------------------------------------------------------------------K----------MVS--------DAA---L---NEIISK--N----LTI--------------------I--A-----------QWN------------------CQTP-AQPN-----K-V-QRVGVVY----DG-----P-----P--------DN-------------KI-ILF------------IVTAHKPL-PK---I-----D-Y------------------------------------H----YSSLHAISINITA-PIG--LRINS-T-------VYV-C---------------------------------------------------------------------------------------------------------------------------------------------------------------G--------------V---------------------------------------------------------------------------------------------------------------------------------------------VEQN----------------

>Sfru-alphaREPAT5

---------MR-----SF-I----------I---I-T------V--L------SA-----L---A--A------CY-G-A---A---------------V----LPS---------EK-----------------P--------E-GY----------I--TMV--F--D--EESP------N---G---------------------------------------------------------------------------------------------------------------------------------------------------------------------------------------------------------------------------R----------VVS--------EAE---L---NN-W----------MA-PN---------------GWY--Y--------EVSTWK------------------CEAP-PIYG-----E-V-QHFSVKY----EG-----D-----P--------DV-------------LI-KSY------------SISYYPPA-PV---I-----H-Y------------------------------------S----PIGTNFMEVNITT-LVG--REIHT-T-------VIL-Y---------------------------------------------------------------------------------------------------------------------------------------------------------------S--------------S-----------------------------------------------------------------------------------------------------------------------------------------------------------------

>Sfru-alphaREPAT6

---------MR-----SF-I----------I---I-T------V--L------SA-----L---A--A------CY-G-A---A---------------V----LPS---------E------------------P--------E-GY----------I--TMV--F--D--EESP------N---G---------------------------------------------------------------------------------------------------------------------------------------------------------------------------------------------------------------------------R----------VES--------DSE---L---NK-WAT--DEWILLTE-PN---------------RLI--Y-----------RWK------------------CEAP-SIDG-----E-E-QYFSVRY----ED-----D-----P--------SV-------------RI-SKF------------SISYFPPR-PF---I-----K-D------------------------------------S----GLGKHFMDVNVTT-GIG--REIRS-T-------IKM-Y---------------------------------------------------------------------------------------------------------------------------------------------------------------RRMLETNIMYLETD-T-----------------------------------------------------------------------------------------------------------------------------------------------------------------

>Sfru-alphaREPAT7

---------MK-----TL-T----------I---I-T------V--L------SA-----L---V--A------CY-E-A-----------------------------------------SGE---------LPG--------G--Y-------------SLV--Y--D--DTA-------D---G---------------------------------------------------------------------------------------------------------------------------------------------------------------------------------------------------------------------------L----------ELS--------NTE---I---NA-RN--------------------QNY------KIV--Y-----------EVV------------------AGAP-TAIG-----R-V-KNSVVSY----RG-----P-----N--------N--------------VK-INR------------L---------V---I-----T-N-----------------H--I-T--PAS--VV-H--T----ALDQSNMEVRFTS-NLS--QGISS-T-------VQM-Y---------------------------------------------------------------------------------------------------------------------------------------------------------------Q--------------R------------------------------------------------------------------------------------------------------------------SGAGRAFGEPAFHLLIVVLMYFGYSQL-------------------A

>Sfru-alphaREPAT8

---------MR-----SF-I----------I---I-T------V--L------LA-----L---V--A------CC-S-A---T---------------A----LSR---------Q------------------E--D-----G--Y----------I--TTT--E--Y--E---------N---G---------------------------------------------------------------------------------------------------------------------------------------------------------------------------------------------------------------------------T----------IVN--------GAD---LEFSNV-IT------------------------------LL--S-----------EWR------------------CEAP-SIAG-----Q-S-QTITLKY----VG-----D-----T--------S--------------ER-IVR------------RVTSIFGD-PH---V-----Q-Q------------------------------------T----PLGSNIYEMTLTS-RPG--VRISA-S-------FHI-Y---------------------------------------------------------------------------------------------------------------------------------------------------------------G--------------------------------------------------------------------------------------------------------------------------------------------------------------------------------

>Sfru-alphaREPAT9

---------MR-----SF-I----------I---I-T------V--L------SA-----L---A--A------CC-S-A---A---------------V----LED---------E------------------E--------K--M----------I--HIV--Y--D--EQFP------N---G---------------------------------------------------------------------------------------------------------------------------------------------------------------------------------------------------------------------------T----------IVS--------DAE---F---DA-RFK--------SV------------------TPL--A-----------EWR------------------ALSP-AISR-----Q-E-QTMGLTY----TG-----D-----P--------G--------------DL-ISR------------ITFTQYGK-PR---V-----H-Y------------------------------------T----QLMRSTISIVITS-RSG--VGLNS-L-------VQI-F---------------------------------------------------------------------------------------------------------------------------------------------------------------T--------------------------------------------------------------------------------------------------------------------------------------------------------------------------------

>Sfru-betaREPAT1

---------MS------L-L----------T---Q-V---------T-------L-----L------C-----------L---C---------------L---------------------------------------------------------------------------S------L---Y--NAAD-------------------------------------------------------------------------------------------------------------------------------------------------------------------------------------------------------L---------QLG-S----------NAN--------A----------------------------------------Q--LGY--Q-----------EIV------------------K-LG-SVPL-----S-T-RTKNVFY----NS-----N-----N--------T--------------KT-IKS------------I---------S---A-----V-D-----------------I--D-K--S-K--AV-A--TITAGGIGSSFVNVRLKS-EKG--KGLNY-L-------VQI-F---------------V--------------------------------------------------------------------------------------------------------------------------------------------------------------------------------------------------------------------------------------------------------------------------------------------------------------------------------

>Sfru-betaREPAT10

MA-------AK------V-L----------L---S-V------L--L-------L--S--I---S--V-----------C---L---------------V-LGR-----------------------------------------------------------------------A------R---G-----S-------------------------------------------------------------------------------------------------------------------------------------------------------------------------------------------------------L---------SIG-N----------TFG--------S----------------------------------------H--LIY--Q-----------ETH------------------Y-KY-GIPF-----I-V-RDEDITV----KG-----V-----N--------N--------------EI-IEA------------V---------V---V-----N-D-----------------L--Q-G----D--GL-S--YIKTGGVGQHDVTIHLES-SRG--HGYKF-L-------VEV-Y---------------A--------------------------------------------------------------------------------------------------------------------------------------------------------------MKYYK-------------------------------------------------------------------------------------------------------------------------------------------------------------

>Sfru-betaREPAT11a

----MFKH--------------------------L-L------I--L-------C-----A---I--V-----------A---A---------------V---------------------------------------------------------------------------H------C---K-----S-------------------------------------------------------------------------------------------------------------------------------------------------------------------------------------------------------V---------IVG-Q----------TWEQAGV---------------------------------------------R--KVY--E-----------TTV------------------S-AN-SIPF-----F-K-REKFVNF----EF-----P-----D--------Q-M------------SK-IKG------------I---------A---I-----K-D-----------------L--D-N--G-L--AE-P--SITMGGLGFNFANLKLKS-ERG--SGFNF-L-------IEI-Y---------------G--------------------------------------------------------------------------------------------------------------------------------------------------------------------------------------------------------------------------------------------------------------------------------------------------------------------------------

>Sfru-betaREPAT11b

----MFKE--------------------------I-V------I--L-------C-----G---V--I-----------V---L---------------V---------------------------------------------------------------------------Q------S---N-----S-------------------------------------------------------------------------------------------------------------------------------------------------------------------------------------------------------V---------FVG-H----------VYDQTGL----V----------------------------------------R--KVY--D-----------KMV------------------E-ST-GIPL-----I-K-REEEIYY----EY-----P-----A--------G-D------------QK-IKG------------I---------A---I-----K-D-----------------L--T-D--S-K--AQ-A--SITRGGLGFGFVNIKLKS-ERG--SGYKY-L-------IEI-F---------------A--------------------------------------------------------------------------------------------------------------------------------------------------------------------------------------------------------------------------------------------------------------------------------------------------------------------------------

>Sfru-betaREPAT12

---------MR------T-L-------SA-A---I-L------L--C-------V-----L---A--A-----------A---A---------------S---------------------------------------------------------------------------Q------S---H-----D-------------------------------------------------------------------------------------------------------------------------------------------------------------------------------------------------------L---------ILG-Q----------ATY--------G----------------------------------------D--VIV--F-----------KAD------------------EFKY-GFPL-----F-V-RTSIVEY-----P-EQGQK-----N--------Y--------------AY-IKA------------I---------Y---V-----K-DNE---------------R--D-G--T-G--GY-P--SISAGGIGQRFVKIKLKS-QRG--SGLNF-T-------ITI-Y---------------G--------------------------------------------------------------------------------------------------------------------------------------------------------------R-----------------------------------------------------------------------------------------------YV----------------------------------------------------------------

>Sfru-betaREPAT13a

---------MY------T-K----------V---V-A------L--F-------G-----L------V-----------A---V---------------A---------------------------------------------------------------------------A------A---A-----D-------------------------------------------------------------------------------------------------------------------------------------------------------------------------------------------------------L---------SVG-R---------E--G--------G----------------------------------------Q--LIF--D-----------KNV------------------T----ASPA-----IWK-QVQNLTV----NA-----T-----D--------D--------------GL-ISR------------V---------V---V-----I-D-----------------NRPK-K----D--GE-A--KVVDGGVGHNDVTIELKS-PTV-FRGFDF-T-------VKV-Y---------------A--------------------------------------------------------------------------------------------------------------------------------------------------------------S----EGAA--------Q-----FPQH--------PVPVG----------------AS-KDG----------------------------------------------------------------------------------------------------

>Sfru-betaREPAT13b

---------MY------T-K----------V---V-A------L--F-------G-----L------V-----------A---V---------------A---------------------------------------------------------------------------A------A---A-----D-------------------------------------------------------------------------------------------------------------------------------------------------------------------------------------------------------L---------SVG-R---------E--G--------G----------------------------------------R--LIF--D-----------KNV------------------T----ASPA-----IWK-QVQNLTV----NA-----T-----D--------D--------------GL-ISR------------V---------V---V-----I-D-----------------NRPE-K----D--GE-A--KVVDGGVGHNDVTIELKS-PTV-FRGFDF-T-------VKV-Y---------------A--------------------------------------------------------------------------------------------------------------------------------------------------------------S----EGAA--------Q-----FPQH--------PVPVG----------------AD-TKV----------------------------------------------------------------------------------------------------

>Sfru-betaREPAT15

---------MKSL----L-V----------L---F-V------V--C-------L-----S---V----------------------------------V---------------------------------------------------------------------------Y------A---N-----H-------------------------------------------------------------------------------------------------------------------------------------------------------------------------------------------------------L---------IVGNI----------ANR--------V----------------------------------------I--LAN--Q-----------ARV------------------E-YN-AIPF-----I-K-RVKYFFY----SQ-----P-----D--------N--------------RL-IQG------------I---------Q---A-----L-D-----------------T--L-H--S-K--AS-I--NITAGGVGYSYVNLRLKS-ERG--RGLAY-D-------IGI-Y-VQDSFYN----------------------------------------------------------------------------------------------------------------------------------------------------------------------------------------------------------------------------------------------------------------------------------------------------------------------------------------

>Sfru-betaREPAT16

---------MK------S-F---LIFC---V---F-I------L--T-------L-----S---S--V---------------L---------------S---------------------------------------------------------------------------S------TYFFG-----E---------------------------------------------------------------------------------------------------------------------------------------------------------------------------------------------------------------------Q----------TKK--------T----------------------------------------R--IIY--Q-----------DKV------------------R-YE-AIPL-----K-K-RVKTYTY----NG-----T-----V------------------------P-IKA------------I---------S---C-----Y-D-----------------F--Q-N--S-E--AS-V--NITAGGIGYKYVELRLKS-QRS--YRLDY-A-------IEI-Y---------------A--------------------------------------------------------------------------------------------------------------------------------------------------------------------------------------------------------------------------------------------------------------------------------------------------------------------------------

>Sfru-betaREPAT17

---------MD------L-L----------L---I-V------L--L-------A-----I---V--T-----------D---A------KMI------C---------------------------------------------------------------------------K------K---T-----K-------------------------------------------------------------------------------------------------------------------------------------------------------------------------------------------------------V---------PL-------------------------------------------------------------------------------------ELG------------------KWRY-DLKH-----M-D--SAVVEN-----P-----P-----V--------S--------------IS-EGQ------------M---------Y---V-----Y-ENYFPNYTIKYIHVDNLAI--R-S--C-G--AS-A--VMKTGGIGSSYVLIVLHA-HIN--EEIRS-V-------VDI-W---------------G-------ERKGGKEG---NV--D---KLP--RSET--KP----GELKTNQSLHLFKHLRAVNHNS----AY---------------------------------------------------------------------------------------------------------------------------------------------------------------------------------------------------------------------------------------------------------

>Sfru-betaREPAT2a

---------MK-C-V--V-A----------L---S-V------F--V-------L-----I---S--I-----------S---S---------------C---------------------------------------------------------------------------L------A---S-----D-------------------------------------------------------------------------------------------------------------------------------------------------------------------------------------------------------L---------IIG-T----------TYN--------A----------------------------------------R--LTW--Q-----------QKA------------------D-YM-GIPF-----K-K-RVKEVFY----SD-----P-----G--------Q--------------QI-IKG------------I---------V---A-----R-D-----------------I--D-H--T-S--GS-A--TVTAGGVGFSFANIRLKS-ERG--NGLNY-Q-------IEI-Y---------------V--------------------------------------------------------------------------------------------------------------------------------------------------------------------------------------------------------------------------------------------------------------------------------------------------------------------------------

>Sfru-betaREPAT2b

---------MK-C-V--V-V----------L---S-V------F--V-------L-----I---S--I-----------S---S---------------C---------------------------------------------------------------------------L------A---S-----D-------------------------------------------------------------------------------------------------------------------------------------------------------------------------------------------------------L---------IIG-T----------TYN--------A----------------------------------------R--LTW--Q-----------QKA------------------E-YM-GIPF-----K-K-RVKEVFY----SD-----P-----G--------Q--------------QI-IKG------------I---------V---A-----R-D-----------------I--D-H--T-S--GS-A--TVTAGGVGFSFANIRLKS-ERG--NGLNY-Q-------IEI-Y---------------V--------------------------------------------------------------------------------------------------------------------------------------------------------------------------------------------------------------------------------------------------------------------------------------------------------------------------------

>Sfru-betaREPAT3

---------MK-T-A--P-V----------F---L-V------V--L-------L-----S---I--I-----------C---V---------------V---------------------------------------------------------------------------H------C---K-----D-------------------------------------------------------------------------------------------------------------------------------------------------------------------------------------------------------L---------IVG-S----------SFN--------N----------------------------------------R--LIH--Q-----------EKA------------------E-YN-AIPL-----K-K-RVKEVFF----SD-----P-----G--------Q--------------QI-IKG------------I---------I---L-----R-D-----------------L--D-H--T-E--AI-P--SVTAGGVGFSYANIRLKS-ERG--SGLNY-Q-------IEI-Y---------------V--------------------------------------------------------------------------------------------------------------------------------------------------------------------------------------------------------------------------------------------------------------------------------------------------------------------------------

>Sfru-betaREPAT4

---------MR------F-V----------F---L-L------L--L-------V-----L---C--L-----------L---C---------------V---------------------------------------------------------------------------S-VP-VLC---S-----D-------------------------------------------------------------------------------------------------------------------------------------------------------------------------------------------------------M---------IVG-D----------TVH--------R----------------------------------------K--MIF--H-----------QRV------------------K-DF-AIPF-----K-K-RVKTLTF----SD-----P-----E--------K--------------RM-IKG------------V---------A---V-----I-D-----------------N--D-F--S-H--AS-A--NITEGGVGYHYVTVRMKS-QRH--HPLNF-E-------VEI-Y---------------V--------------------------------------------------------------------------------------------------------------------------------------------------------------------------------------------------------------------------------------------------------------------------------------------------------------------------------

>Sfru-betaREPAT5

---------MK-F----V-L----------L---L-F------A--T-------I-----F---V--V-----------A---T---------------C--------------------------------------------------------------------------------------D-----H-------------------------------------------------------------------------------------------------------------------------------------------------------------------------------------------------------L---------ILGNT----------NNN--------Q----------------------------------------N--MIY--H-----------TTA------------------H-YT-AIPF-----I-K-RVKNIFY----SG--------------------N--------------SI-INS------------I---------M---A-----Y-D-----------------N--K-H--T-N--AS-A--AVTAGGIGYTYVNLRLKS-ERG--KELDY-D-------IGI-Y---------------A--------------------------------------------------------------------------------------------------------------------------------------------------------------------------------------------------------------------------------------------------------------------------------------------------------------------------------

>Sfru-betaREPAT6

---------MK------C-V----------L-----------------------------I---T--VCLIL-------L---C---------------V---------------------------------------------------------------------------C------L---N--EGKD-------------------------------------------------------------------------------------------------------------------------------------------------------------------------------------------------------F---------IAG-T----------KAN--------N----------------------------------------L--LIS--T-----------QKL------------------K-YR-SIPL-----I-R-RDKDYTY----ID-----S-----K--------E--------------RI-IKG------------I---------I---A-----R-D-----------------L--S-R--T-D--TE-V--TVTSGGVGATNVTLHFQS-GRG--EELNY-L-------ILI-F---------------S---NIK--------------------------------------------------------------------------------------------------------------------------------------------------------------------------------------------------------------------------------------------------------------------------------------------------------------------------

>Sfru-betaREPAT7a

---------MK------V-V----------A---L-A------F--L-------L-----A---V--V-----------V---I---------------V---------------------------------------------------------------------------D------C---T-----H-------------------------------------------------------------------------------------------------------------------------------------------------------------------------------------------------------T---------FMG-T----------NVL--------R----------------------------------------P--LIY--H-----------HSA------------------E-YG-SKVF-----R-K-RVENLYY----SL-----P-----P---VPANAG--------------RS-IQG------------I---------L---A-----F-D-----------------M--T-H--T-G--AT-A--NVTQGGLGFNFVNLRMKS-DRG--EKIHF-D-------VYI-Y---------------A--------------------------------------------------------------------------------------------------------------------------------------------------------------------------------------------------------------------------------------------------------------------------------------------------------------------------------

>Sfru-betaREPAT7b

---------MH------I-Y----------S---L-A------I--V-------L-----A---V--V-----------V---V---------------S---------------------------------------------------------------------------E------C---G-----H-------------------------------------------------------------------------------------------------------------------------------------------------------------------------------------------------------T---------FLG-T----------SVL--------R----------------------------------------P--LVF--H-----------KEV------------------R-YP-AKVL-----V-K-RIENFNF----TL-----P-----S---VPGVYN--------------RS-IQG------------I---------L---A-----Y-D-----------------L--T-N--S-G--AS-A--NVTQGGLGHNFVNIRMKS-DRY--KEIHY-D-------IYI-Y---------------A--------------------------------------------------------------------------------------------------------------------------------------------------------------------------------------------------------------------------------------------------------------------------------------------------------------------------------

>Sfru-betaREPAT8a

---------MY----F-K-V----------A---L-V------L--S-------L-----V---Y--L-----------T---S---------------V---------------------------------------------------------------------------E------S---N-----D-------------------------------------------------------------------------------------------------------------------------------------------------------------------------------------------------------I---------MLG-F---------ATPY--------S----------------------------------------K--KIY--S-----------AIK------------------T----ANPA-----LWQ-REDDVLI----ST-----P-----N--------H--------------EL-IDA------------V---------Y---I-----T-D-----------------LRPE-K----D--GI-S--YIEAGGIGSKTVTIALKS-PSI-LRGYKF-A-------IEV-Y---------------A--------------------------------------------------------------------------------------------------------------------------------------------------------------Q----S-LHHGYGTNILPVQ---GAGYPRAG--LSGYG---------------------SQYP--GYPTAYSGVGVYPQHN-Y---G-------KK------------------------------------------------------------------

>Sfru-betaREPAT8b

---------MY----S-K-V----------L---V-T------L--F-------V-----L---Y--L-----------S---T---------------V---------------------------------------------------------------------------E------C---N-----D-------------------------------------------------------------------------------------------------------------------------------------------------------------------------------------------------------I---------HIG-Y---------TNTR--------S----------------------------------------R--KVY--S-----------ELK------------------E----ADPA-----LWR-RTDEVTI----KT-----P-----P--------H--------------EV-IDS------------I---------H---V-----S-D-----------------LREE-K----D--GE-A--YITSGGVGMPYVTIALKS-PTI-LRGYKF-Q-------IEV-Y---------------A--------------------------------------------------------------------------------------------------------------------------------------------------------------S----S-TA--------S-----NEGLFSK---SPYFGDV----------------QQ-QQVS-----------------------R-------KF------------------------------------------------------------------

>Sfru-gammaREPAT6

---------MH-----SV-F----------M---I-V------L--L-------VGC---V---A--V-----------S---Y---------TT----A---------------------------------------------------------------------------F------E---D-----DF-----T--D--------G---------------------------------------------------------------------------------------------I---------------------------------------------ET------------RR---------------------G---L---------SLG-S----------IGG--------D----------------------------------------D--RLL--S-----------NTN------------------HASD-GAAF-----V-V-QTQEVVI----RP-----V-----S--------Y--------------TN-ITA------------I---------Q---V-----T-E-----------------V--G-G--AHN--AQ-V--SVVDGGIGKNFVRLSFRS-EIG--RGYSY-N-------VRI-H---------------G--------------------------------------------------------------------------------------------------------------------------------------------------------------R----------------------------------------------------------------------------------------------------------MGC----------------------------------------------------

>Sfru-gammaREPAT7

---------MK-----SI-F----------V---I-V------L--L-------AGC---V---A--V-----------G---Y---------TS----A---------------------------------------------------------------------------T------E---V-----EP-----S--V--------E---------------------------------------------------------------------------------------------S---------------------------------------------AV------------RS---------------------G---L---------SLG-S----------IGP--------D----------------------------------------N--RLL--S-----------NTN------------------HAVE-AAVF-----V-V-QRQEVLI----RT-----V-----S--------Y--------------SK-ITS------------I---------Q---V-----S-E-----------------V--G-T--PVN--ST-V--SLIDGGLNRNFVKFLFQS-QVG--RAYSY-N-------VRI-H---------------G--------------------------------------------------------------------------------------------------------------------------------------------------------------L----------------------------------------------------------------------------------------------------------IEC----------------------------------------------------

>Sfru-gammaREPAT16

---------MK-----SV-T----------L---V-V------L--L-------A-----V---T--L-----------G---Y---------AA----V---------------------------------------------------------------------------D------A---A-----SR-----P--Q--------A---------------------------------------------------------------------------------------------L---------------------------------------------FT---EPDDNLEKPRV---------------------L---L---------ASK-N----------SD---------N----------------------------------------D--RLL--S-----------STY------------------HRVD-RSHR-----V-A-HNQEIVI----RL-----V-----E--------Y--------------AR-ITR------------V---------E---G-----W-G----------------------G--S-D--SV-L--KVLDGGLGHNFIKLGYGT-EEE--MDFNY-R-------FYV-Y---------------G--------------------------------------------------------------------------------------------------------------------------------------------------------------Y----------------------------------------------------------------------------------------------------------LVCDD--------------------------------------------------

>Sfru-gammaREPAT17

---------MK-----I--L----------A---L-L------L--L-------------V---A--V-----------A---A---------A-----V---------------------------------------------------------------------------N------S---Y-----EG-----Q--E--------A-------------------NVY-YEQLLE---V--------------------------------EESGR----------------------------------------------------------------------S------------HL---------------------NI--F---------QFG-T----------IGP--------D----------------------------------------D--RLI--S-----------DTT------------------HEAY-GIEY-----H-V-NDHELTY----RG-----S-----E--------N--------------TN-ITG------------I---------I---V-----H-E-----------------Y--L-S--NYT---A-V--TIQDGGIHRNYVTISLKG-RRS--YHVGC-R-------IGI-F---------------A--------------------------------------------------------------------------------------------------------------------------------------------------------------S----------------------------------------------------------------------------------------------------------INC--SNM-----------------------------------------------

>Sfru-gammaREPAT18

---------MM-----M--L--------------V-V------L--L-------------V---T--I-----------A---A---------A-----V---------------------------------------------------------------------------H------T---N-----EG-----Q--E--------A-------------------KLLNYDA-QDSHHF--------------------------------TDADE-----------------------M---------------------------------------------GR------------AS---------------------R---R---------QWG-S----------VGP--------D----------------------------------------D--RLI--S-----------TTV------------------HTME-ALSF-----F-I-NYQEVTY----RG-----L-----E--------N--------------TN-ITM------------I---------R---T-----T-E-----------------P--W-G--S-T---E-V--VLLDGGLYHNFVTLGFKS-ERS--SSMTNMA-------IDI-F---------------A--------------------------------------------------------------------------------------------------------------------------------------------------------------T----------------------------------------------------------------------------------------------------------INC--SNN-----------------------------------------------

>Sfru-gammaREPAT19

---------MK-----SV-T----------L---V-V------L--L-------A-----V---T--L-----------G---Y---------AA----V---------------------------------------------------------------------------D------G---D-----DL-----L--E--------A---------------------------------------------------------------------------------------------P---------------------------------------------ST---GPE-GISTRNG---------------------I---L---------SIG-D----------D----------N----------------------------------------G--RLF--S-----------STT------------------HQVL-SSPN-----V-L-HKQEVLI----RL-----V-----E--------Y--------------AN-ITG------------I---------R---A-----N-P----------------------S--STL--GV-V--TLIDGGLGHNYVKLGFNS-QKN--LGYLY-R-------ILV-Y---------------G--------------------------------------------------------------------------------------------------------------------------------------------------------------S----------------------------------------------------------------------------------------------------------LVCDD--------------------------------------------------

>Sfru-gammaREPAT1

---------MK-----SL-I----------L---V-A------V--L-------A---ALA---V--F-----------N---D---------A---------------------------------------------------------------------------------------------------A-----T-LQ--------N---------------------------------------------------------------------------------------------G---------------------------------------------AF------------RA---------------------N---M---------YQG-G----------IRP--------G----------------------------------------D--RLL--H-----------SNY------------------YYKN-PITN-----A-V-QYQDITY----RG-----N-----A--------S--------------TR-ISY------------I---------Q---A-----T-E-----------------V--G-Y--TQW--GI-P--SLRAGGVNYNYATIRLTS-QRG--YGYYY-R-------VEI-W---------------G--------------------------------------------------------------------------------------------------------------------------------------------------------------R-----------------------------------------------------------------------------------------------------------------------------------------------------------------

>Sfru-gammaREPAT2

---------MK-----SL-I----------L---V-A------V--L-------A---ALT---I--C-----------N---D---------A---------------------------------------------------------------------------------------------------A-----S-LQ--------N---------------------------------------------------------------------------------------------G---------------------------------------------AF------------RA---------------------N---M---------YQG-A----------IRP--------G----------------------------------------D--RIL--H-----------SNY------------------YTKG-PIPN-----A-V-QYQDITY----RG-----N-----A--------S--------------TR-ISY------------I---------Q---V-----T-E-----------------V--G-Y--TQW--GI-P--SLRAGGVNFNHATIRLTS-QRG--YGYYY-R-------VEI-W---------------G--------------------------------------------------------------------------------------------------------------------------------------------------------------R-----------------------------------------------------------------------------------------------------------------------------------------------------------------

>Sfru-gammaREPAT3

---------MK-----SL-I----------L---V-A------V--L-------A---ALA---V--C-----------N---D---------A---------------------------------------------------------------------------------------------------A-----S-LE--------S---------------------------------------------------------------------------------------------P---------------------------------------------AF------------RA---------------------N---L---------YQG-A----------IRP--------G----------------------------------------D--RLL--H-----------SNY------------------YYKN-PITN-----A-V-QYQDITY----RG-----N-----A--------S--------------TR-ISY------------I---------Q---A-----T-E-----------------V--G-Y--TQW--GI-P--SLRAGGVNFNHATIRLTS-QRG--YGYYY-R-------VEI-W---------------G--------------------------------------------------------------------------------------------------------------------------------------------------------------R-----------------------------------------------------------------------------------------------------------------------------------------------------------------

>Sfru-gammaREPAT4

---------MK-----SL-I----------L---V-A------V--L-------A---ALT---I--C-----------N---D---------A---------------------------------------------------------------------------------------------------A-----S-LQ--------N---------------------------------------------------------------------------------------------P---------------------------------------------AF------------RA---------------------N---L---------YQG-A----------IRP--------G----------------------------------------D--RLL--H-----------SNY------------------YYKN-PITN-----A-V-QYQDITY----RG-----N-----A--------S--------------TR-ISY------------I---------Q---A-----T-E-----------------V--G-Y--TQW--GI-P--SLRAGGVNFNHATIRLTS-QRG--YGYYY-R-------VEI-W---------------G--------------------------------------------------------------------------------------------------------------------------------------------------------------R-----------------------------------------------------------------------------------------------------------------------------------------------------------------

>Sfru-gammaREPAT5

---------MK-----SL-I----------L---V-A------V--L-------A---ALT---I--C-----------N---D---------A---------------------------------------------------------------------------------------------------A-----S-LQ--------N---------------------------------------------------------------------------------------------P---------------------------------------------AF------------RA---------------------N---L---------YQG-A----------IRP--------G----------------------------------------D--RLL--H-----------SNY------------------YYKN-PITN-----A-V-QYQDITY----RG-----N-----A--------S--------------TR-ISY------------I---------Q---A-----T-E-----------------V--G-Y--TQW--GI-P--SLRAGGVNFNHATIRLTS-QRG--YGYYY-R-------VEI-W---------------G--------------------------------------------------------------------------------------------------------------------------------------------------------------R-----------------------------------------------------------------------------------------------------------------------------------------------------------------

>Sfru-gammaREPAT8

---------MK-----SL-F----------L---A-V------L--L-------V-----A---V--A-----------S---A---------A---------------------------------------------------------------------------------------------------TINEEIV--D--------D---------------------------------------------------------------------------------------------PNLFETSEEFDLSDDPDVSNDRDVADMTDVADVTEV-PV----SEV-T------------SP---------------------S---IDAATCC---SLG-A-----------RS--------G-------------------------------------------RLL--V-----------RSN------------------QVRH-GKPN-----V-T-YIQNLTF----RT-----G-----A--------P---------------R-INA------------I---------Q---I-----I-H-----------------I--G-A--MQG--AS-A--SIVSGGLNRNHVTIRLQS-ARG--RGYNF-R-------ILI-Y---------------G--------------------------------------------------------------------------------------------------------------------------------------------------------------R-----------------------------------------------------------------------------------------------------------------------------------------------------------------

>Sfru-gammaREPAT9

---------MK-----SL-F----------L---A-V------L--L-------V-----A---V--A-----------S---A---------A---------------------------------------------------------------------------------------------------TINGEIV--D--------D---------------------------------------------------------------------------------------------PNLLETSEEFDLSDDLDVSNDPDVADMTDVADVTEV-PV----SEV-T------------TP---------------------S---IDAATCC---SVG-A-----------RS--------G-------------------------------------------RVL--V-----------RSN------------------HVRR-GKPN-----V-T-YIQNITF----RT-----G-----T--------R---------------R-ITA------------I---------Q---I-----I-H-----------------I--G-A--VQG--AR-A--SIVSGGLNRNHVTIRLQS-ARG--RGYNF-K-------IVI-Y---------------G--------------------------------------------------------------------------------------------------------------------------------------------------------------R-----------------------------------------------------------------------------------------------------------------------------------------------------------------

>Sfru-gammaREPAT10

---------MK-----IL-L----------L---F-V------L--I-------T-----T---A--G-----------S---A---------SILDYEV---------------------------------------------------------------------------T------D---N-----DD-----Q--E--------D----------------------------------------------------------------------YTMTPDRDYTPDYYEEEMMVRISP---------------------------------------------TT------------TAVNIDEWAELFRMAVEEAKIHRN---L---------SLG-T----------IGA--------N----------------------------------------D--ILL--S-----------KTH------------------HFKS-DGDT-----N-V--AEDVYF----CG-----L-----P--------T--------------TV-ITA------------I---------R---G-----G-S-----------------V-------SEN--AR-P--AVLAGGVGHNNVCIRLHS-SKG--RGYQY-C-------IEI-Y---------------G--------------------------------------------------------------------------------------------------------------------------------------------------------------R---------------------------------------------------------------------------------------------------------K-------------------------------------------------------

>Sfru-gammaREPAT11

---------MK-----SL-L----------L---V-V------L--L-------A-----A---A--A-----------T---A---------A---------------------------------------------------------------------------------------------------IVHQDVT--D--------A---------------------------------------------------------------------------------------------PEAVVEYEEILYSDE---------------AEVAEV-EAGSRSVEIES------------RR---------------------N---L---------SVG-T----------IGN--------S-------------------------------------------RLL--A-----------NTQ------------------HTRN-AVAN-----T-I-IVQNITF----NF-----A-----S--------S--------------IR-IAA------------I---------R---V-----T-R-----------------V--G-A--SQN--AT-P--SIVSGGLNRNFVTIRITS-ARG--RGYNY-R-------IQI-Y---------------G--------------------------------------------------------------------------------------------------------------------------------------------------------------R-----------------------------------------------------------------------------------------------------------------------------------------------------------------

>Sfru-gammaREPAT12

---------MK-----SL-L----------L---V-V------L--L-------A-----A---A--A-----------T---A---------A---------------------------------------------------------------------------------------------------IVHQDIT--E--------A---------------------------------------------------------------------------------------------PEGIVEYEEILYNDE---------------AEVAEV-EAGSRSVEIES------------RR---------------------N---L---------SVG-T----------IGN--------S-------------------------------------------RLL--A-----------NTQ------------------HIRG-AVAN-----T-I-IVQNITF----NF-----A-----S--------S--------------IR-IAA------------I---------R---V-----T-R-----------------V--G-A--SQN--AT-P--SIVSGGLNRNFVTIRITS-ARG--RGYNY-R-------IQI-Y---------------G--------------------------------------------------------------------------------------------------------------------------------------------------------------R-----------------------------------------------------------------------------------------------------------------------------------------------------------------

>Sfru-gammaREPAT13

---------MK-----SL-L----------L---V-V------L--L-------A-----A---A--A-----------T---A---------A---------------------------------------------------------------------------------------------------IVHQDIA--E--------A---------------------------------------------------------------------------------------------PEAVVEYENILYSDE---------------AEVAEV-EAGSRSVEIES------------RR---------------------N---L---------SVG-T----------IGN--------S-------------------------------------------RLL--A-----------NTQ------------------HIRG-AVAN-----T-I-IVQNITF----NF-----A-----S--------S--------------IR-IAA------------I---------R---V-----T-R-----------------V--G-A--SQN--AT-P--SIVSGGLNRNFVTIRITS-ARG--RGYNY-R-------IQI-Y---------------G--------------------------------------------------------------------------------------------------------------------------------------------------------------R-----------------------------------------------------------------------------------------------------------------------------------------------------------------

>Sfru-gammaREPAT14

---------MK-----SL-F----------L---V-V------L--L-------A-----A---A--A-----------T---A---------A---------------------------------------------------------------------------------------------------IVSQDVA--E--------A---------------------------------------------------------------------------------------------PEGVVEYEEILYSDE---------------AEMAEVEEAGSRSMEIES------------RR---------------------N---L---------SVG-T----------IGN--------S-------------------------------------------RLL--A-----------NTQ------------------HIRG-AVAN-----T-I-IVQNITF----NF-----A-----S--------S--------------IR-IAA------------I---------R---V-----T-R-----------------V--G-A--SQN--AT-P--SIVSGGLNRNFVTIRITS-ARG--RGYNY-R-------IQI-Y---------------G--------------------------------------------------------------------------------------------------------------------------------------------------------------R-----------------------------------------------------------------------------------------------------------------------------------------------------------------

>Sfru-gammaREPAT15

---------MK-----SL-L----------L---V-V------L--L-------A-----A---A--A-----------T---A---------A---------------------------------------------------------------------------------------------------IVSQDVA--E--------A---------------------------------------------------------------------------------------------PEGVVEYEEILYSDE---------------TELAEVEEAGSRSVEIES------------RR---------------------N---L---------SVG-T----------IGN--------S-------------------------------------------RLL--A-----------NTQ------------------HIRG-AVAN-----T-I-IVQNITF----NF-----A-----S--------S--------------IR-IAA------------I---------R---V-----T-R-----------------V--G-A--SQN--AT-P--SIVSGGLNRNFVTIRITS-ARG--RGYNY-R-------IQI-Y---------------G--------------------------------------------------------------------------------------------------------------------------------------------------------------R-----------------------------------------------------------------------------------------------------------------------------------------------------------------

>Slitt-alphaREPAT1

---------MR-----SF-I----------I---I-A------V--L------SA-----L---A--A------CY-D-C---A---------------V--------------------------------------------------------------------------IN------N---E---------------------------------------------------------------------------------------------------------------------------------------------------------------------------------------------------------------------------V----------VTT--------RGT--------A-RE-----------------------G--D--IIF--F-----------EKT------------------YYVP-AEPL-----S-V-QRLDVAY----YG-----N-----S--------T--------------TY-INQ------------I---------R---I-----T-E--------------------P-R--LNPLGSV-TS-S----ALGKTFIQLRISS-VVG--GFLHL-T-------VQF-L---------------G--------------------------------------------------------------------------------------------------Y----------------------------------------YTDSP--------------------------------------------------------------------------------------------------------------------------------------------------------------------------------

>Slitt-alphaREPAT10

---------MK-----NF-I----------I---I-A------V--L------SA-----L---V--A------CY-A-D------------------------IPS---------L------------------E--------P--M-------------SLV--Y--DEDDTCM------D---G---------------------------------------------------------------------------------------------------------------------------------------------------------------------------------------------------------------------------K----------PMS--------SAD---V---NL-RN--------------------TTW------VVI--S-----------EWQ------------------CNAP-AVAG-----L-V-QRVNVHY----EG-----A-----P--------N--------------LR-INR------------L---------I---V-----T-Y-----------------T--V-T--PAA--VD-W--S----PLSTNVMTVTIRS-HAG--QQVNA-N-------VTL-L---------------------------------------------------------------------------------------------------------------------------------------------------------------R--------------R----------------------------------------------------------------------------------------------------------------------------------------------------------------A

>Slitt-alphaREPAT11

---------MR-----SF-I----------I---I-T------V--L------SA-----L---A--A------CC-S-A---S---------------V----LPN---------E------------------E--------E--M----------I--TVV--Y--D--ENFP------N---G---------------------------------------------------------------------------------------------------------------------------------------------------------------------------------------------------------------------------T----------IVS--------KEE---L---DS-RFS--------RV------------------VAE--S-----------SWV------------------CSVP-AISR-----Q-S-QVITMQY----LG-----D-----P--------R--------------DL-IRR------------IVHSYSGN-PT---M-----S-H------------------------------------S----PLFSNNMLVTITS-RSG--QGINS-R-------IEL-Y---------------------------------------------------------------------------------------------------------------------------------------------------------------L--------------------------------------------------------------------------------------------------------------------------------------------------------------------------------

>Slitt-alphaREPAT12

---------MR-----TF-I----------I---I-A------L--L------SA-----L---A--A------CY-GAA-------------------V----LPS---------E------------------K-V----MSE--G----------I--TIV--Y--D--EDSP------N---G---------------------------------------------------------------------------------------------------------------------------------------------------------------------------------------------------------------------------R----------IMS--------RSD----------TV---------------DNLF----------VLL--S-----------ESS------------------INVP-AIPG-----K-V-QKAGITY----QA-----P-----E--------G--------------VR-ISR------------V---------S---I-----TLY-----------------A--N-L--PGD--TQ-N--S----PLGSDYVESHVTS-RPG--DAINS-SSTGFIDTIK------------------------------------------------------------------------------------------------------------------------------------------------------------------------------------------------------------------------------------------------------------------------------------------------------------------------------------E-Q------------

>Slitt-alphaREPAT13

---------MR-----TF-I----------I---I-T------V--L------SA-----L---A--A------CY-S-AA------------------D----LPS---------E------------------D------MSR--G----------Y--TVL--Y--D--DMSP------E---G---------------------------------------------------------------------------------------------------------------------------------------------------------------------------------------------------------------------------R----------IVS--------SAE---L---NS-KS---------------DF-W----------IPI--S-----------EAS------------------LVAP-AVAG-----Q-R-QYITKVF----QA-----A-----P--------G--------------VI-ING------------L---------S---V-----T-Y-----------------S--K-T--PAY--SY-R--S----PLGSDSVIVSITS-HPG--DEIFS-K-------ITV-Y---------------------------------------------------------------------------------------------------------------------------------------------------------------R--------------R----------------------------------------------------------------------------------------------------------------------------------------------------P------------

>Slitt-alphaREPAT14

---------MR-----SF-I----------I---I-T------V--L------SA-----L---A--A------CY-GAA-------------------V----LPS---------E------------------E-E----M---------------V--TII--Y--D--DKSP------N---G---------------------------------------------------------------------------------------------------------------------------------------------------------------------------------------------------------------------------I----------IMS--------RAD---F---NA-SS---------------QG-W----------VLL--S-----------KTN------------------DFVP-AVEG-----Q-V-QSVGVFY----RA-----N-----F--------G--------------VR-IDR------------L---------V---I-----TTY-----------------G--H-L--IPD--VN-R--T----PLGSNKMIVILTS-RPG--ELIRS-T-------IET-Y---------------------------------------------------------------------------------------------------------------------------------------------------------------K--------------R----------------------------------------------------------------------------------------------------------------------------------------------------DEISS--------

>Slitt-alphaREPAT15

---------MR-----TF-I----------I---I-T------V--L------LA-----L---A--A------CY-S-AA------------------D----LPS---------E------------------D------MSG--G----------Y--TVL--Y--D--DLSP------E---G---------------------------------------------------------------------------------------------------------------------------------------------------------------------------------------------------------------------------R----------IVS--------SAE---L---NA-KS---------------NI-W----------IPI--T-----------EAT------------------FEAP-AVAG-----Q-V-QHITKSF----KA-----P-----P--------G--------------VV-INA------------L---------V---V-----S-Y-----------------S--E-K--PME--LY-L--G----PLGTNSMIVSITS-HPG--DEISS-T-------LSI-Y---------------------------------------------------------------------------------------------------------------------------------------------------------------S-------------------------------------------------------------------------------------------------------------------------------------------------------------------Y------------

>Slitt-alphaREPAT16

---------MR-----SF-I----------I---I-T------V--L------SV-----L---A--A------CY-GAA-------------------V----LPS---------E------------------E-E----M---------------V--TII--Y--D--DKSP------N---G---------------------------------------------------------------------------------------------------------------------------------------------------------------------------------------------------------------------------I----------IMS--------RAD---F---DA-RS---------------QG-W----------VLL--S-----------EAN------------------DLIP-AVEG-----Q-V-QSIRMVY----RA-----Q-----R--------G--------------VK-ISM------------L---------S---V-----SAR-----------------L--S-G--IPE--VD-N--T----PLGRNYIDVKVTS-RLG--LGLNS-T-------ISV-Y---------------------------------------------------------------------------------------------------------------------------------------------------------------R--------------W----------------------------------------------------------------------------------------------------------------------------------------------------D------------

>Slitt-alphaREPAT17

---------MR-----TF-I----------I---I-T------V--L------SA-----L---A--A------CY-S-AA------------------D----LPS---------E------------------D------MSG--G----------Y--TVL--Y--D--DMSP------E---G---------------------------------------------------------------------------------------------------------------------------------------------------------------------------------------------------------------------------K----------IVS--------SAE---L---NS-KS---------------DF-W----------IPI--S-----------EAS------------------LVAP-AVAG-----Q-R-QYITKVF----QA-----A-----P--------G--------------VI-ING------------L---------S---V-----T-Y-----------------S--K-T--PAY--SY-R--S----PLGSDSVIVSITS-HPG--DEISS-K-------ITI-Y---------------------------------------------------------------------------------------------------------------------------------------------------------------R--------------R----------------------------------------------------------------------------------------------------------------------------------------------------P------------

>Slitt-alphaREPAT18

---------MR-----SF-I----------I---I-T------V--L------SA-----L---A--A------CY-GAA-------------------V----LPS---------E------------------E-E----M---------------V--TII--Y--D--DKSP------N---G---------------------------------------------------------------------------------------------------------------------------------------------------------------------------------------------------------------------------I----------IMS--------RAD---F---DA-RS---------------QG-W----------VLL--S-----------EAN------------------DLIP-AVEG-----E-V-QSIRMVY----RA-----Q-----R--------G--------------VK-ISM------------L---------S---V-----SAR-----------------L--S-G--IPE--VD-N--T----PLGRNYIDVKVTS-RLG--LGLNS-T-------ISV-Y---------------------------------------------------------------------------------------------------------------------------------------------------------------R--------------W----------------------------------------------------------------------------------------------------------------------------------------------------D------------

>Slitt-alphaREPAT19

---------MR-----TF-I----------I---I-T------V--L------SA-----L---A--A------CY-S-AA------------------D----LPS---------E------------------D------MSG--G----------Y--TVL--Y--D--DMSP------E---G---------------------------------------------------------------------------------------------------------------------------------------------------------------------------------------------------------------------------R----------IVS--------SAE---L---NS-KS---------------DF-W----------IPI--S-----------EAS------------------LVAP-AVAG-----Q-R-QYITKVF----QA-----A-----P--------G--------------VI-ING------------L---------S---V-----T-Y-----------------S--K-T--PAY--SY-R--S----PLGSDSVIVSITS-HPG--DDISS-K-------ITI-Y---------------------------------------------------------------------------------------------------------------------------------------------------------------R--------------R----------------------------------------------------------------------------------------------------------------------------------------------------P------------

>Slitt-alphaREPAT2

---------MR-----SF-I----------I---I-A------V--L------SA-----L---A--A------CY-G-T---P---------------V----LPS---------A------------------E--------E--T-------YPEY--KMIHKYNKE--DDAQ------N---R---------------------------------------------------------------------------------------------------------------------------------------------------------------------------------------------------------------------------I----------TML--------REEMRH-------DN-----------------------DRQNGEVLL--D-----------------------------------T-AVPKDCDSYV-I-QYVDILY----YG-----I-----P--------P--------------TV-ITT------------V---------E---I-----T-S-----------------S--P-P--PSF--VL-P--I----WHLDNYISIRISS-QRG--LGLNG-T-------VQI-F---------------G-TL-----------------------------------------------------------------------------------------------------------------------------------------------------------------------------------------------------------------------------------------------------------------------------------------------------------------------------

>Slitt-alphaREPAT20

---------MR-----SF-I----------I---I-T------V--L------SA-----L---A--A------CY-GAA-------------------V----LPS---------E------------------E-E----M---------------V--TII--Y--D--DKSP------N---G---------------------------------------------------------------------------------------------------------------------------------------------------------------------------------------------------------------------------I----------IMS--------RAD---F---DA-RS---------------QG-W----------VLL--S-----------EAN------------------DLIP-AVEG-----E-V-QSIRMVY----RA-----Q-----R--------G--------------VK-ISM------------L---------S---V-----SAR-----------------L--S-G--IPE--VD-N--T----PLGRNYIDVKVTS-RLG--LGLNS-T-------ISV-Y---------------------------------------------------------------------------------------------------------------------------------------------------------------R--------------W----------------------------------------------------------------------------------------------------------------------------------------------------D------------

>Slitt-alphaREPAT21

---------MR-----SF-I----------I---V-T------V--L------SA-----L---A--A------SY-N-A---T---------------V----LPD---------A------------------E--------N--M---PSNF---I--YTL--Y--D--GNSP------N---G---------------------------------------------------------------------------------------------------------------------------------------------------------------------------------------------------------------------------T----------IVS--------LND---I---AI-KS-------------------S---------FLL--S-----------TWR------------------HNAQ-PFPG-----E-E-QRIEMKY----EG-----L-----P--------G--------------DR-ITT------------I---------M---V-----G-H-----------------S--T-T--PAN--IY-H--S----PLFDYYIDMQIIS-HRG--QGIDT-Y-------VAI-Y---------------------------------------------------------------------------------------------------------------------------------------------------------------G--------------R-----------------------------------------------------------------------------------------------------------------------------------------------------------------

>Slitt-alphaREPAT22

---------MR-----SF-I----------I---I-T------V--L------SA-----L---A--A------CY-G-VA------------------V----PPS---------E------------------E------VSG--G----------Y--TIL--F--D--NISP------E---G---------------------------------------------------------------------------------------------------------------------------------------------------------------------------------------------------------------------------R----------IVS--------SAE---L---SA-GS---------------DR-W----------VKL--S-----------ESI------------------LNVP-AVPG-----Q-V-QRIGTKY----QG-----D-----A--------S--------------VV-ITR------------W---------S---V-----T-------------------S--D-V--PTA--SY-S-FS----PFGSNLMEVTIQS-HPG--NGINS-T-------VRI-Y---------------------------------------------------------------------------------------------------------------------------------------------------------------T--------------D----------------------------------------------------------------------------------------------------------------------------------------------------L------------

>Slitt-alphaREPAT23

---------MK-----SF-I----------I---L-T------V--L------SA-----L---A--A------CY-G-AA------------------V----VPS---------E------------------E------MSE--G----------Y--TMI--F--D--EMSP------D---G---------------------------------------------------------------------------------------------------------------------------------------------------------------------------------------------------------------------------R----------IVP--------NAE---L---NA-KS---------------SE-W----------ILI--S-----------EGV------------------VNAP-AVPG-----Q-V-QRVGTWH----QA-----D-----L--------G--------------VR-IER------------L---------A---L-----S-Y-----------------G--A-F--PAD--VQ-H--T----PLGSNFLEVKIIS-GLG--QRVAA-T-------ISV-Y---------------------------------------------------------------------------------------------------------------------------------------------------------------R--------------R---------------------------------------------------------------------------------------------------------------------------------------------------GC----------C-

>Slitt-alphaREPAT24

---------MR-----SF-I----------I---I-T------V--L------SA-----L---A--A------CY-G-A---A---------------V----TSK---------V------------------D--------E--M-------TPIF--TRT--YTYP--D-LT------N---K---------------------------------------------------------------------------------------------------------------------------------------------------------------------------------------------------------------------------T----------MSL--------DDE--------D-LP-----------------------P--I--VTL--F-----------EKE------------------HFVP-AVPG-----E-V-QTLLVIYEETERT-----I-----A--------I--------------TH-LHV------------S---------L---F-----A-I--------------------P-R--HML--RY-T--S----EFGAS---VYIES-QVG--DPIRA-I-------VKV-S---------------G--------------------------------------------------------------------------------------------------R----------------------------------------VYD----------------------------------------------------------------------------------------------------------------------------------------------------------------------------------

>Slitt-alphaREPAT25

---------MR-----SF-I----------I---I-V------V--L------SA-----L---A--A------CY-G-A---A---------------V----LPNEATMFSPFIQ------------------E--------Q--I-------LDGW--TVE--NNEE--DVAQ------N---R---------------------------------------------------------------------------------------------------------------------------------------------------------------------------------------------------------------------------V----------TLS--------KGE--------A-RN-----------------------D--L--ILL--F-----------ERT------------------CYVL-AIPN-----D-V-ARCDVVY----YG-----E-----P--------N--------------IR-ISR------------I---------S---VN----S-N-----------------W--P-V--PAY--VH-V--S----GLSTNVGRVRLTS-RIG--EGLHA-V-------VRM-F---------------GIIN-----------------------------------------------------------------------------------------------E----------------------------------------DNNES--------------------------------------------------------------------------------------------------------------------------------------------------------------------------------

>Slitt-alphaREPAT26

---------MR-----SF-I----------I---I-T------V--L------SA-----L---A--T------CY-S-A---A---------------V----PPSEEGMISRFAL------------------E--------Q--M-------YPDF--TVE--YNEK--D--Q------N---R---------------------------------------------------------------------------------------------------------------------------------------------------------------------------------------------------------------------------V----------VFS--------KGE--------A-RN-----------------------E--Q--ILL--L-----------TRE------------------IHVP-AVPG-----E-T-QYAHLIH----RS-----S-----S--------N--------------VR-ITR------------V---------V--WIP----L-T-----------------W--P-L--PTI--IS-I--K----GIPSRVVEISCTS-FVG--ESIHL-M-------FYV-Y---------------GEVI-----------------------------------------------------------------------------------------------E----------------------------------------N------------------------------------------------------------------------------------------------------------------------------------------------------------------------------------

>Slitt-alphaREPAT27

---------MR-----SF-I----------I---I-L------V--L------SA-----L---A--A------CH-G-A---A---------------V----LPDEANMFSSFIR------------------E--------Q--M-------LDGW--TLE--HNEE--DAAQ------N---R---------------------------------------------------------------------------------------------------------------------------------------------------------------------------------------------------------------------------V----------TMN--------KGE--------A-RD-----------------------D--F--IVL--Y-----------ERP------------------CIVP-AIPN-----D-I-VSCEVVY----HG-----E-----P--------N--------------VR-IMR------------L---------S---VS----Y-F-----------------W--P-P--PAH--ID-A--S----GLLTNVGRVKLTS-HIG--ERLQA-V-------VRM-Y---------------GVII-----------------------------------------------------------------------------------------------E----------------------------------------DNN----------------------------------------------------------------------------------------------------------------------------------------------------------------------------------

>Slitt-alphaREPAT3

---------MK-----SF-I----------I---I-A------V--L------SA-----L---V--A------SY-E-A---F---------------V----LPS---------R--VMSGRHDAFVSSRQMTD--------D--F-------------TLV--Y--D--DTAV------N---G---------------------------------------------------------------------------------------------------------------------------------------------------------------------------------------------------------------------------T----------IMT--------SAE---V---ET-KN--------------------DGW------RLL--Y-----------EVS------------------CSAP-AAVG-----R-V-KNSFVTY----QG-----P-----T--------N--------------TV-INR------------I---------T---I-----T-P-----------------Q--I-H--PAN--VF-H--S----ALGTNNMEVRLIS-NVS--QPIQA-T-------VRL-Y---------------------------------------------------------------------------------------------------------------------------------------------------------------Q-------------QR------------------------------------------------------------------------------------------------------------------SGAGKVFGEPAFHLMFFVLIYIGISNF-------------------T

>Slitt-alphaREPAT4

---------MR-----SF-I----------I---I-T------V--L------SA-----F---A--A------CY-G-V---S---------------D----LMN---------E---------------------------E--VQER-------Y--TML--Y--D--EKTP------N---G---------------------------------------------------------------------------------------------------------------------------------------------------------------------------------------------------------------------------R----------IVS--------DTE---L---NK-ITE--E----ISI--------------------L--A-----------QWY------------------CQAP-ALPD-----K-Q-QRVGVMY----EG-----P-----P--------GT-------------TM-VVY------------LVFYHPPE-PK---I-----E-H------------------------------------S----KWGSNFLSINITS-PIG--ISVDA-V-------VAL-Y---------------------------------------------------------------------------------------------------------------------------------------------------------------G--------------K---------------------------------------------------------------------------------------------------------------------------------------------IPK-----------------

>Slitt-alphaREPAT5

---------MR-----TF-I----------I---I-T------V--L------SA-----L---A--A------CC-S-A---S---------------I----LPS---------E------------------E--------E--M----------I--TVV--Y--D--DESP------N---G---------------------------------------------------------------------------------------------------------------------------------------------------------------------------------------------------------------------------R----------ILS--------NAE---L---HN-S-IIPNNFVLLAEY-----------------GII--Y-----------RWR------------------CDAP-SING-----Q-V-QRVGVRY----DG-----D-----P--------NV-------------QI-SQV------------SVTYYSPA-PR---I-----D-R------------------------------------S----PLGRNFMEVNITT-PVG--REVWS-T-------VRVYY---------------------------------------------------------------------------------------------------------------------------------------------------------------R--------------A-----------------------------------------------------------------------------------------------------------------------------------------------------------------

>Slitt-alphaREPAT6

---------MK-----SL-I----------L---I-T------V--L------SA-----L---I--A------CY-E-A-----------------------------------------SGE---------LPG--------G--Y-------------SVV--Y--D--DTT-------D---G---------------------------------------------------------------------------------------------------------------------------------------------------------------------------------------------------------------------------N----------VMS--------NTE---I---NA-RN--------------------QNF------KVI--Y-----------EVV------------------ATAS-ASIG-----R-V-KNSFVTY----QG-----P-----N--------N--------------VR-VDR------------L---------V---I-----T-N-----------------H--V-Q--PAA--VY-H--S----TLGTNNMEVRFTS-NVS--QPISS-T-------VLM-Y---------------------------------------------------------------------------------------------------------------------------------------------------------------Q--------------R------------------------------------------------------------------------------------------------------------------SGAGRAFGEPAFHLLVLVLMYFGYSQL-------------------A

>Slitt-alphaREPAT7

---------MR-----SF-I----------I---I-T---------V------LA-----L---A--A------CC-N-A---S---------------A----LSR---------Q------------------E--DLSE--V--Y----------V--TTT--E--Y--E---------N---G---------------------------------------------------------------------------------------------------------------------------------------------------------------------------------------------------------------------------T----------IVD--------GAD---FALSNV-AS------------------------------LL--S-----------EWH------------------CEAP-SVVG-----Q-S-QTITLKY----LG-----D-----S--------S--------------ER-ITR------------RVTTIYGQ-PQ---V-----T-Q------------------------------------T----PLGSNVYEMTLTS-RLG--ERISA-S-------FYI-Y---------------------------------------------------------------------------------------------------------------------------------------------------------------A--------------------------------------------------------------------------------------------------------------------------------------------------------------------------------

>Slitt-alphaREPAT8

---------MR-----SF-I----------V---I-A------V--L------SA-----L---A--T------CY-G-A---S---------------V----VPS---------E------------------D--------E--Y----------I--TTI--Y--D--EKSP------N---G---------------------------------------------------------------------------------------------------------------------------------------------------------------------------------------------------------------------------T----------IVS--------NGD---IEFRNE-RS-----------------------------SIF--A-----------QWD------------------SVIP-SSMD-----R---HTITFMY----VG-----D-----P--------R--------------DI-ISR------------LETAISGE-PE-I-V-----Y-R------------------------------------S----PMNTNIFQMILMT-RAG--RRTSA-R-------IQI-S---------------------------------------------------------------------------------------------------------------------------------------------------------------R--------------------------------------------------------------------------------------------------------------------------------------------------------------------------------

>Slitt-alphaREPAT9

---------MR-----SF-I----------I---I-T------V--L------SA-----L---A--A------CY-S-A---T---------------D----LPS---------E------------------E--------E--M----------Y--TIV--Y--D--EKYP------N---G---------------------------------------------------------------------------------------------------------------------------------------------------------------------------------------------------------------------------T----------IVS--------DAE---I---DS-RFK--------SV------------------IPL--A-----------EWR------------------CRAP-AISR-----R-Q-QTITVAY----HG-----D-----P--------R--------------DI-ITR------------TTFSTSGN-PR---I-----T-Q------------------------------------T----PLMRSSILITITS-RSG--VGIDS-L-------VQI-F---------------------------------------------------------------------------------------------------------------------------------------------------------------T--------------------------------------------------------------------------------------------------------------------------------------------------------------------------------

>Slitt-betaREPAT1

---------MS------L-L----------T---Q-V---------T-------I-----F------C-----------L---C---------------L---------------------------------------------------------------------------A------L---Y--NAED-------------------------------------------------------------------------------------------------------------------------------------------------------------------------------------------------------L---------QLG-S----------GVN--------A----------------------------------------Q--LGY--Q-----------EIV------------------K-LG-SVPL-----T-V-RTKNVFY----NS-----N-----N--------T--------------KT-IKS------------I---------S---A-----V-D-----------------I--D-K--S-K--AV-P--TITAGGIGSSFVNVRLKS-EKG--KALNY-L-------VQI-F---------------V--------------------------------------------------------------------------------------------------------------------------------------------------------------------------------------------------------------------------------------------------------------------------------------------------------------------------------

>Slitt-betaREPAT10

MA-------VK------A-L----------L---S-I------L--L-------L--S--I---S--V-----------T---L---------------V-LAR-----------------------------------------------------------------------G------R---G-----S-------------------------------------------------------------------------------------------------------------------------------------------------------------------------------------------------------L---------SIG-N----------TFG--------S----------------------------------------H--LIY--Q-----------ETH------------------Y-KY-GIPF-------V-RDEDITV----KG-----V-----N--------N--------------EI-IEA------------V---------V---V-----N-D-----------------L--E-G----N--GL-C--YIRTGGVGHHDVTIHLES-SRG--HGYKF-L-------VEV-Y---------------A--------------------------------------------------------------------------------------------------------------------------------------------------------------KKYYK-------------------------------------------------------------------------------------------------------------------------------------------------------------

>Slitt-betaREPAT11a

----MFKH--------------------------L-L------I--L-------C-----A---I--V-----------V---A---------------V---------------------------------------------------------------------------H------C---K-----S-------------------------------------------------------------------------------------------------------------------------------------------------------------------------------------------------------V---------IVG-Q----------TWEQNGV---------------------------------------------R--KVY--E-----------TTV------------------S-AN-SIPL-----F-K-RTKYVNF----EF-----P-----D--------Q-I------------SK-IKG------------I---------A---I-----K-D-----------------Q--D-N--G-L--AE-P--SITMGGLGFNFANLKLKS-ERG--SGFNF-L-------IEI-Y---------------G--------------------------------------------------------------------------------------------------------------------------------------------------------------------------------------------------------------------------------------------------------------------------------------------------------------------------------

>Slitt-betaREPAT11b

----MFKE--------------------------I-V------I--L-------F-----G---A--I-----------V---L---------------V---------------------------------------------------------------------------Q------S---N-----S-------------------------------------------------------------------------------------------------------------------------------------------------------------------------------------------------------V---------FVG-H----------VYDQTGL----V----------------------------------------R--KVY--D-----------KMV------------------E-AT-GIPL-----I-K-REEEIFY----EY-----P-----A--------G-D------------QK-IKG------------I---------A---I-----K-D-----------------L--T-D--S-K--AQ-A--SITRGGLGFGFVNIKLKS-ERG--SGYKY-L-------IEV-F---------------A--------------------------------------------------------------------------------------------------------------------------------------------------------------------------------------------------------------------------------------------------------------------------------------------------------------------------------

>Slitt-betaREPAT12

---------MR------T-S-------SV-A---I-L------L--C-------L-----L---A--A-----------A---A---------------S---------------------------------------------------------------------------Q------S---H-----D-------------------------------------------------------------------------------------------------------------------------------------------------------------------------------------------------------L---------ILG-Q----------ATY--------G----------------------------------------D--VIV--F-----------KAD------------------EVKY-GFPL-----F-V-RTSIVEY-----P-EQGQK-----N--------Y--------------AY-IKA------------I---------Y---V-----K-DNE---------------R--D-G--S-G--GY-P--SISAGGIGQRFVKIKLKS-QRG--SGLNF-T-------VTI-Y---------------G--------------------------------------------------------------------------------------------------------------------------------------------------------------R-----------------------------------------------------------------------------------------------YV----------------------------------------------------------------

>Slitt-betaREPAT13a

---------MF------T-K----------V---V-A------L--F-------C-----L------V-----------A---V---------------A---------------------------------------------------------------------------A------A---T-----D-------------------------------------------------------------------------------------------------------------------------------------------------------------------------------------------------------L---------TVG-K---------R--E--------G----------------------------------------R--LIF--Y-----------TTI------------------T----ARPT-----IWK-QVQNLTV----NA-----T-----D--------D--------------AL-ITR------------V---------V---V-----V-D-----------------NRPQ-K----D--GV-A--KLVDGGVGHNNVTIELKS-PTV-FRGFNF-T-------VKV-F---------------E--------------------------------------------------------------------------------------------------------------------------------------------------------------K----KVAM--------Q-----KPRH--------PVFVG----------------AN-TQK----------------------------------------------------------------------------------------------------

>Slitt-betaREPAT13b

---------MF------T-K----------V---V-A------F--F-------G-----L------V-----------A---V---------------A---------------------------------------------------------------------------A------A---A-----D-------------------------------------------------------------------------------------------------------------------------------------------------------------------------------------------------------L---------SVG-K---------E--G--------G----------------------------------------Q--LIF--D-----------KNV------------------T----ASPA-----IWK-QVQNLTV----NA-----T-----D--------D--------------AL-ITR------------V---------V---V-----I-D-----------------NRPE-K----D--GE-A--KVVDGGVGHNNITIELKS-PTV-FRGFDF-T-------VKV-F---------------T--------------------------------------------------------------------------------------------------------------------------------------------------------------N----EGAV--------Q-----VPQH--------PVSVG----------------AN-TQV----------------------------------------------------------------------------------------------------

>Slitt-betaREPAT15

---------MKLI----L-V----------C---L-I------V--C-------L-----S---V----------------------------------V---------------------------------------------------------------------------Y------A---N-----H-------------------------------------------------------------------------------------------------------------------------------------------------------------------------------------------------------L---------IVGNV----------ANR--------V----------------------------------------V--LAN--Q-----------ARV------------------E-YN-AIPF-----I-K-RVKYFFY----TQ-----P-----D--------N--------------RL-IQG------------I---------Q---A-----L-D-----------------T--M-H--S-K--AS-I--NITAGGVGYSYVNLRLKS-ERG--RGLAY-D-------IGI-Y-VQDLFYH----------------------------------------------------------------------------------------------------------------------------------------------------------------------------------------------------------------------------------------------------------------------------------------------------------------------------------------

>Slitt-betaREPAT16

---------MR------S-F---LIFC---V---F-I------L--T-------L-----S---S--V---------------L---------------S---------------------------------------------------------------------------S------TYFFG-----E---------------------------------------------------------------------------------------------------------------------------------------------------------------------------------------------------------------------Q----------TKN--------T----------------------------------------R--IIY--Q-----------DKV------------------R-YE-AIPL-----K-K-RVKTYSY----NG-----T-----V------------------------P-IKA------------I---------S---C-----Y-D-----------------F--Q-N--S-E--AS-V--NITAGGVGYKYVELRLKS-QRS--YRLDY-A-------IEI-Y---------------A--------------------------------------------------------------------------------------------------------------------------------------------------------------------------------------------------------------------------------------------------------------------------------------------------------------------------------

>Slitt-betaREPAT17

---------MD------L-L----------L---I-V------L--L-------A-----I---I--T-----------D---A------KMI------C---------------------------------------------------------------------------K------K---T-----K-------------------------------------------------------------------------------------------------------------------------------------------------------------------------------------------------------A---------PL-------------------------------------------------------------------------------------ELG------------------EWRY-DLKH-----M-D--SAVVEN-----P-----P-----V--------S--------------IN-EGQ------------M---------Y---V-----Y-ENYFPNYTIKYIHVDNLAI--R-S--C-G--AS-A--MMRTGGIGSSYVLIVLHA-HIN--EEIRS-V-------VDI-W---------------G-------ERK--------KVITE---ELP--RTPL--KP----DEMTSLKSLYLFKHLRAVNHNS----AY---------------------------------------------------------------------------------------------------------------------------------------------------------------------------------------------------------------------------------------------------------

>Slitt-betaREPAT2

---------MK-C-V--L-A----------L---S-V------F--V-------V-----I---A--I-----------S---S---------------C---------------------------------------------------------------------------L------A---S-----D-------------------------------------------------------------------------------------------------------------------------------------------------------------------------------------------------------L---------IIG-T----------SYN--------A----------------------------------------R--LAW--Q-----------HKS------------------D-YM-GIPF-----K-K-RVKEVFY----SD-----P-----G--------Q--------------QI-IKG------------I---------I---A-----R-D-----------------I--D-H--T-S--GS-A--TVTAGGVGFSFANIRLKS-ERG--SGLNY-Q-------IEI-Y---------------V--------------------------------------------------------------------------------------------------------------------------------------------------------------------------------------------------------------------------------------------------------------------------------------------------------------------------------

>Slitt-betaREPAT3

---------MK-T-A--P-V----------F---L-V------V--L-------L-----S---I--I-----------C---V---------------V---------------------------------------------------------------------------H------C---K-----D-------------------------------------------------------------------------------------------------------------------------------------------------------------------------------------------------------L---------IVG-S----------SFN--------N----------------------------------------R--LIH--Q-----------EKA------------------E-YN-AIPL-----K-K-RVKEVFF----SD-----P-----G--------Q--------------QI-IKG------------I---------I---L-----R-D-----------------L--D-H--T-E--AI-P--SVTAGGVGFSYANIRLKS-ERG--SGLNY-Q-------VEI-Y---------------V--------------------------------------------------------------------------------------------------------------------------------------------------------------------------------------------------------------------------------------------------------------------------------------------------------------------------------

>Slitt-betaREPAT4

---------MR------F-V----------F---L-L------L--L-------V-----L---C--L-----------L---C---------------V---------------------------------------------------------------------------S-VP-VLC---S-----D-------------------------------------------------------------------------------------------------------------------------------------------------------------------------------------------------------M---------IVG-D----------TVH--------R----------------------------------------K--MIF--H-----------QRI------------------K-DF-AIPF-----K-K-RVKTLTF----SD-----P-----E--------K--------------RM-IKG------------V---------A---V-----V-D-----------------N--D-F--S-H--AS-A--NITEGGVGYSYVTVRMKS-QRH--HPLNF-E-------MEI-Y---------------V--------------------------------------------------------------------------------------------------------------------------------------------------------------------------------------------------------------------------------------------------------------------------------------------------------------------------------

>Slitt-betaREPAT5

---------MK-L----I-L----------V---L-F------A--S-------I-----F---V--V-----------A---T---------------C--------------------------------------------------------------------------------------D-----H-------------------------------------------------------------------------------------------------------------------------------------------------------------------------------------------------------L---------ILGNT----------NNN--------Q----------------------------------------N--MIY--H-----------TTS------------------H-YS-AIPF-----I-K-RVKNVFY----SG--------------------S--------------SV-INS------------I---------M---A-----Y-D-----------------N--K-H--T-N--AS-A--AVTAGGIGYTYVNLRLKS-ERG--KELDY-D-------IGI-Y---------------S--------------------------------------------------------------------------------------------------------------------------------------------------------------------------------------------------------------------------------------------------------------------------------------------------------------------------------

>Slitt-betaREPAT6

---------MK------C-V----------L-----------------------------I---T--VCLIL-------L---R---------------V---------------------------------------------------------------------------C------L---N--EGKD-------------------------------------------------------------------------------------------------------------------------------------------------------------------------------------------------------F---------IVG-T----------KAN--------N----------------------------------------L--LIS--T-----------QKL------------------K-YR-SIPL-----I-R-RDKDYTY----ID-----S-----K--------E--------------RI-IKG------------I---------I---A-----R-D-----------------L--S-R--T-D--TD-V--TVTSGGVGATNVTLHFQS-GRG--EELNY-L-------ILI-F---------------S---NIK--------------------------------------------------------------------------------------------------------------------------------------------------------------------------------------------------------------------------------------------------------------------------------------------------------------------------

>Slitt-betaREPAT7a

---------MK------V-V----------A---L-A------F--L-------L-----A---I--V-----------V---I---------------V---------------------------------------------------------------------------D------C---S-----H-------------------------------------------------------------------------------------------------------------------------------------------------------------------------------------------------------T---------FMG-T----------NVL--------R----------------------------------------P--LIY--H-----------HNV------------------E-YG-SKVF-----R-K-RVENLYY----SL-----P-----A---VPANAG--------------RS-IQG------------I---------L---A-----F-D-----------------M--T-H--T-G--AT-A--NVTQGGLGFNFVNLRMKS-DRG--EKIHF-D-------VYI-Y---------------A--------------------------------------------------------------------------------------------------------------------------------------------------------------------------------------------------------------------------------------------------------------------------------------------------------------------------------

>Slitt-betaREPAT7b

---------MH------I-S----------S---I-A------L--V-------L-----A---M--I-----------V---V---------------S---------------------------------------------------------------------------E------C---G-----H-------------------------------------------------------------------------------------------------------------------------------------------------------------------------------------------------------T---------FLG-T----------SVQ--------R----------------------------------------P--LVF--H-----------KEV------------------R-YP-ARGF-----Q-K-RIENVNF----TL-----P-----H---VSDIYL--------------RK-IQG------------I---------L---A-----Y-D-----------------L--T-N--S-G--AS-A--NVTKGGLGYNFVNIRLKS-DRN--KEIRY-D-------INI-Y---------------A--------------------------------------------------------------------------------------------------------------------------------------------------------------------------------------------------------------------------------------------------------------------------------------------------------------------------------

>Slitt-betaREPAT8a

---------MF----F-K-V----------T---L-L------V--S-------L-----L---Y--L-----------T---S---------------V---------------------------------------------------------------------------E------S---N-----D-------------------------------------------------------------------------------------------------------------------------------------------------------------------------------------------------------I---------RLG-F---------ATPY--------S----------------------------------------K--KIY--S-----------AIK------------------I----ANPA-----LWQ-REDDVLI----SS-----P-----N--------H--------------EL-IDA------------V---------Y---I-----T-D-----------------LRPE-K----D--GI-A--YIEAGGIGSKTVTIALKS-PSL-LRGYKF-A-------IEV-Y---------------A--------------------------------------------------------------------------------------------------------------------------------------------------------------Q----S-LHHGYGTNILPIQ---GAAYPRAG--LSGYG---------------------SQYP--GYPTAYSGVGVYP--NNY---G-------KK------------------------------------------------------------------

>Slitt-betaREPAT8b

---------MF----A-K-V----------L---F-T------L--F-------V-----L---Y--L-----------S---T---------------V---------------------------------------------------------------------------Q------C---N-----D-------------------------------------------------------------------------------------------------------------------------------------------------------------------------------------------------------I---------RLG-Y---------TNLR--------S----------------------------------------I--KIY--S-----------ELK------------------E----ADPA-----LWK-RTDEVTI----KA-----P-----P--------H--------------EF-IDS------------I---------H---V-----S-D-----------------LRED-K----D--GE-A--YITSGGIGMPYVTISLKS-PTI-LRGYKF-Q-------IEV-Y---------------A--------------------------------------------------------------------------------------------------------------------------------------------------------------S----P-LA--------S-----NEGLFSKG--SFNYGDA----------------QQVQQYP-----------------------R-------KF------------------------------------------------------------------

>Slitt-betaREPAT9

---------MG------F-R----------I---V-I------F--N-------Y-----I---T--L-----------V---S---------------I---------------------------------------------------------------------------T------C---H-----D-------------------------------------------------------------------------------------------------------------------------------------------------------------------------------------------------------F---------FIG-S---------PDSS--------S----------------------------------------K--LI---------------KGN------------------E----LKPA-----VWI-RSDILSV----NC-----S-----G--------N--------------DV-ISS------------V---------Q---I-----M-D-----------------LIPE-K----L--SE-A--HLKSGGIGSKFVVIELRS-SG--YEGLRF-L-------VRM-F---------------V--------------------------------------------------------------------------------------------------------------------------------------------------------------F----P--S--------S--LRYRYM--------------------------------------------------------------------KI------------------------------------------------------------------

>Slitt-gammaREPAT5

---------MK-----S--F----------V---I-V------L--L-------AGC---A---I--V-----------G---Y---------SS----A---------------------------------------------------------------------------T------E---V-----EP-----S--V--------K---------------------------------------------------------------------------------------------S---------------------------------------------EA------------RG---------------------G---L---------SLG-T----------LGP--------D----------------------------------------D--RLL--S-----------NTN------------------HAVE-AAVY-----V-V-QSQEVLI----RT-----V-----E--------Y--------------SN-ITS------------I---------H---V-----S-E-----------------V--G-T--TTN--AS-V--SVIDGGLRTNFVKFLFRS-QKG--RAYSY-N-------IRI-H---------------G--------------------------------------------------------------------------------------------------------------------------------------------------------------I----------------------------------------------------------------------------------------------------------IEC----------------------------------------------------

>Slitt-gammaREPAT6

---------MK-----IF-I----------V---L-V------L--L-------------A---A--A-----------A---A---------AG----V---------------------------------------------------------------------------T------N---N-----DR-----E--Q--------AA--------ALENKELHIPDTVDNNE-LD---VP---EKNELNVPDRPDNTELDVPEDEI-GEEDEETGL-----------------------A---------------------------------------------AA------------RN---------------------N---L---------SLG-T----------FGA--------D----------------------------------------D--RLM--S-----------RIT------------------HNVA-ATTL-----L-I-HDQEVTF----RG-----T-----R--------G--------------TN-ITA------------I---------R---V-----N-Q-----------------V--G-S--NTP---I-A--TRVAGGLGSEFVTIRIQS-GRG--RAFSY-T-------ILV-Y---------------A--------------------------------------------------------------------------------------------------------------------------------------------------------------A----------------------------------------------------------------------------------------------------------INC--SNN-----------------------------------------------

>Slitt-gammaREPAT13

---------MK-----SF-M----------F---V-V------L--L-------A-----V---T--L-----------G---Y---------AT----V---------------------------------------------------------------------------D------A---A-----KR-----P--E--------A---------------------------------------------------------------------------------------------Q---------------------------------------------FT---GPGKTTSKAHV---------------------L---L---------ASK-N----------SV---------N----------------------------------------D--RIF--S-----------STY------------------HSVD-RSHR-----I-A-HNQEILI----RL-----V-----D--------F--------------AK-ITR------------V---------E---G-----W-A----------------------G--A-D--SV-L--KVLDGGLGHNFIKLGYNT-EED--KFFSY-R-------VDV-Y---------------G--------------------------------------------------------------------------------------------------------------------------------------------------------------V----------------------------------------------------------------------------------------------------------LECDD--------------------------------------------------

>Slitt-gammaREPAT14

---------MK-----V--L----------A---V-V------L--LVAVA---------V---A--V-----------A---A---------A-----V---------------------------------------------------------------------------N------S---Y-----ET-----Q--E--------A-------------------SVF-YEE-PD---M--------------------------------EEPDP----------------------------------------------------------------------SVSR---------RL---------------------H---Y---------QWG-S----------VGP--------D----------------------------------------D--RLI--S-----------ATY------------------HEAY-GMEY-----Q-I-NDQEVTY----RG-----S-----E--------K--------------TN-ITS------------I---------I---V-----R-E-----------------Y--L-S--NYT---T-V--TILDGGLYRNHVTLGLKG-LRS--LQIGC-H-------ISI-Y---------------A--------------------------------------------------------------------------------------------------------------------------------------------------------------T----------------------------------------------------------------------------------------------------------VNC--SNM-----------------------------------------------

>Slitt-gammaREPAT15

---------MK-----I--L----------A---V-V------L--L----ML-------V---T--I-----------A---A---------G-----V---------------------------------------------------------------------------N------S---N-----LE-----Q--E--------A-------------------NVLNSSE-QDPNKM--------------------------------VESDL-----------------------M---------------------------------------------AS------------R----------------------T---A----------WG-V----------RGP--------D----------------------------------------D--RLI--S-----------TTI------------------QSAD-AFTF-----F-V-NDQEVTY----RG-----H-----E--------T--------------TN-ITR------------I---------D---I-----R-M-----------------R--F-G--DIT---Q-V--DLLDGGLYHNYVTLRFTC-QRS--SSMYN-V-------IDI-Y---------------A--------------------------------------------------------------------------------------------------------------------------------------------------------------T----------------------------------------------------------------------------------------------------------INC--HNN-----------------------------------------------

>Slitt-gammaREPAT16

---------MK-----GF-L----------L---I-I------L--L-------A-----V---T--H-----------G---Y---------AT----V---------------------------------------------------------------------------D------G---E-----DL-----P--E--------A---------------------------------------------------------------------------------------------L---------------------------------------------LG---GPE-GINTFSG---------------------S---L---------SLG-D----------T----------N----------------------------------------G--RMM--S-----------STS------------------HSVE-RSMN-----V-A-HTQEVLV----RL-----V-----E--------Y--------------AN-ITG------------I---------R---V-----T-P----------------------T--GNT--VA-P--SLIDGGLGHNYVKLGFTS-QLN--LGYVY-R-------IVI-Y---------------G--------------------------------------------------------------------------------------------------------------------------------------------------------------S----------------------------------------------------------------------------------------------------------LVCDD--------------------------------------------------

>Slitt-gammaREPAT1

---------MK-----SL-I----------L---V-A------V--L-------A---ALT---V--C-----------N---D---------A---------------------------------------------------------------------------------------------------A-----T-LQ--------N---------------------------------------------------------------------------------------------A---------------------------------------------AF------------RA---------------------N---L---------YQG-G----------IRP--------G----------------------------------------D--RLL--H-----------SNY------------------YIKN-PIAN-----A-V-QYQDITY----RG-----N-----S--------S--------------TR-ISY------------I---------Q---V-----T-E-----------------V--G-Y--TQW--GI-P--SLRSGGVNYSYATIRLTS-QRG--YGYYY-R-------VEI-W---------------G--------------------------------------------------------------------------------------------------------------------------------------------------------------R-----------------------------------------------------------------------------------------------------------------------------------------------------------------

>Slitt-gammaREPAT2

---------MK-----SL-I----------L---V-A------V--L-------A---ALT---V--C-----------N---D---------A---------------------------------------------------------------------------------------------------A-----T-LQ--------N---------------------------------------------------------------------------------------------P---------------------------------------------AY------------RA---------------------N---L---------YQG-A----------IRP--------G----------------------------------------D--RLL--H-----------SNY------------------YYKN-PIAN-----A-V-QYQDITY----RG-----N-----S--------S--------------TR-ISY------------I---------Q---V-----T-E-----------------V--G-Y--TQW--GI-P--SLRSGGVNFQHATIRLTS-PRG--YGYYY-R-------VEI-W---------------G--------------------------------------------------------------------------------------------------------------------------------------------------------------R-----------------------------------------------------------------------------------------------------------------------------------------------------------------

>Slitt-gammaREPAT3

---------MK-----SL-I----------L---V-A------V--L-------A---ALT---V--C-----------N---D---------A---------------------------------------------------------------------------------------------------A-----T-LQ--------N---------------------------------------------------------------------------------------------P---------------------------------------------AY------------RA---------------------N---L---------YQG-A----------IRP--------G----------------------------------------D--RLL--H-----------RNY------------------YYKN-PIAN-----A-V-QYQDITY----RG-----N-----S--------S--------------TR-ISY------------I---------Q---V-----T-E-----------------V--G-Y--TQW--GI-P--SLRSGGVNFNHATIRLTS-QRG--YGYYY-R-------VEI-W---------------G--------------------------------------------------------------------------------------------------------------------------------------------------------------R-----------------------------------------------------------------------------------------------------------------------------------------------------------------

>Slitt-gammaREPAT4

---------MK-----SL-I----------L---V-A------V--L-------A---ALT---V--C-----------N---D---------A---------------------------------------------------------------------------------------------------A-----T-LQ--------N---------------------------------------------------------------------------------------------P---------------------------------------------AY------------RA---------------------N---L---------YQG-A----------IRP--------G----------------------------------------D--RIL--H-----------RNY------------------YIKN-PIAN-----A-V-QYQDITY----RG-----N-----S--------S--------------TR-ISY------------I---------Q---V-----T-E-----------------V--G-Y--TQW--GI-P--SLRSGGVNFNHATIRLTS-QRG--YGYYY-R-------VEI-W---------------G--------------------------------------------------------------------------------------------------------------------------------------------------------------R-----------------------------------------------------------------------------------------------------------------------------------------------------------------

>Slitt-gammaREPAT7

---------MK-----SL-F----------L---I-V------L--L-------V-----V---A--A-----------S---T---------A---------------------------------------------------------------------------------------------------VINQETA--A--------D---------------------------------------------------------------------------------------------PDLVETFEEFDLGDDPDISYEPD------VADVTEV-PI----SDV-T------------SP---------------------S---IDVATCC---SVG-A-----------RS--------G-------------------------------------------KLL--V-----------RSN------------------HVRR-GKLN-----V-T-YVQNVTF----RT-----G-----S--------H---------------L-ITA------------I---------R---I-----T-H-----------------I--G-P--MQG--SR-A--SIIAGGLNRNFVTIRLQS-ARG--RGYKF-R-------IQI-Y---------------G--------------------------------------------------------------------------------------------------------------------------------------------------------------R-----------------------------------------------------------------------------------------------------------------------------------------------------------------

>Slitt-gammaREPAT8

---------MK-----GL-L----------L---L-V------L--L-------A-----A---A--A-----------S---A---------SILNYEV---------------------------------------------------------------------------T------E---E-----DN-----Q--E--------E----------------------------------------------------------------------DTLSPDLDDTPDHYEEEVMLRISP---------------------------------------------TP------------TATDIDEWAELFRMAVEEGMIHRN---L---------SMG-N----------IGA--------H----------------------------------------D--ILL--S-----------KTH------------------HIKN-DGDT-----N-V--SEDVYF----RC-----C-----P--------T--------------TVIITA------------I---------R---G-----G-S-----------------V-------SEN--VR-P--SVVSGGVGHNHVCIRLHS-QKG--HGYRY-C-------IEI-Y---------------G--------------------------------------------------------------------------------------------------------------------------------------------------------------R---------------------------------------------------------------------------------------------------------K-------------------------------------------------------

>Slitt-gammaREPAT9

---------MK-----SL-L----------I---V-V------L--L-------A-----A---A--A-----------S---A---------A---------------------------------------------------------------------------------------------------IVSQDIA--E--------A---------------------------------------------------------------------------------------------PEAVITYESIMYSDD---------------AEVAEV-EAGSKSVDIET------------RR---------------------N---L---------SVG-T----------IGN--------S-------------------------------------------RLL--A-----------NTQ------------------HIRG-AIAN-----T-I-RVLNATF----NF-----A-----S--------S--------------IR-IAA------------I---------R---V-----T-R-----------------V--G-A--SQN--AT-P--SIVSGGLNRNFVTIRITS-ARG--RGFNY-R-------IQI-Y---------------G--------------------------------------------------------------------------------------------------------------------------------------------------------------R-----------------------------------------------------------------------------------------------------------------------------------------------------------------

>Slitt-gammaREPAT10

---------MK-----SL-L----------V---V-V------L--L-------A-----A---A--A-----------S---A---------A---------------------------------------------------------------------------------------------------IVSQDIA--D--------A---------------------------------------------------------------------------------------------PEAVITYESIMYSDD---------------AEVAEV-EAGSKSVDIET------------RR---------------------N---L---------SVG-T----------IGN--------S-------------------------------------------RLL--A-----------NTQ------------------HIRG-AIAN-----T-I-LVHNATF----NF-----A-----S--------S--------------IR-IAA------------I---------R---V-----T-R-----------------V--G-A--SQN--AT-P--SIVSGGLNRNFVTIRITS-ARG--RGFNY-R-------IQI-Y---------------G--------------------------------------------------------------------------------------------------------------------------------------------------------------R-----------------------------------------------------------------------------------------------------------------------------------------------------------------

>Slitt-gammaREPAT11

---------MK-----SL-L----------V---V-V------L--L-------A-----A---V--A-----------S---A---------A---------------------------------------------------------------------------------------------------IVSQDIA--D--------A---------------------------------------------------------------------------------------------PEAVITYESIMYSDD---------------AEVAEV-EAGSKSVDIET------------RR---------------------N---L---------SVG-T----------IGN--------S-------------------------------------------RLL--A-----------NTQ------------------HIRG-AIAN-----T-I-LVHNATF----NF-----A-----S--------S--------------IR-IAA------------I---------R---V-----T-R-----------------V--G-A--SQN--AT-P--SIVSGGLNRNFVTIRITS-ARG--RGFNY-R-------IQI-Y---------------G--------------------------------------------------------------------------------------------------------------------------------------------------------------R-----------------------------------------------------------------------------------------------------------------------------------------------------------------

>Slitt-gammaREPAT12

---------MK-----SL-L----------L---V-V------L--L-------A-----A---A--A-----------S---A---------A---------------------------------------------------------------------------------------------------IVSQDIA--E--------A---------------------------------------------------------------------------------------------PEAVVTYESIMYSDD---------------AEVAEV-EAASKSVDIET------------RR---------------------N---L---------SVG-T----------IGN--------S-------------------------------------------RLL--A-----------NTQ------------------HIRG-AIAN-----T-I-LVHNATF----NF-----A-----S--------S--------------IR-IAA------------I---------R---V-----T-R-----------------V--G-A--SQN--AT-P--SIVSGGLNRNFVTIRITS-ARG--RGFNY-R-------IQI-Y---------------G--------------------------------------------------------------------------------------------------------------------------------------------------------------R-----------------------------------------------------------------------------------------------------------------------------------------------------------------

>Slitu-alphaREPAT1

---------MR-----SF-I----------I---I-A------V--L------SA-----L---A--A------CY-D-C---A---------------V--------------------------------------------------------------------------IN------N---E---------------------------------------------------------------------------------------------------------------------------------------------------------------------------------------------------------------------------V----------VTT--------KGT--------A-RE-----------------------G--D--VIF--F-----------EKT------------------YYVP-AEPF-----S-I-QRLDVAY----YG-----N-----S--------T--------------TY-INQ------------I---------R---I-----I-E--------------------P-R--LNPIGSV-TS-S----ALGKTFMQLRISS-VVG--GFLHL-T-------VQI-L---------------G--------------------------------------------------------------------------------------------------Y----------------------------------------YTDSP--------------------------------------------------------------------------------------------------------------------------------------------------------------------------------

>Slitu-alphaREPAT10

---------MR-----NF-I----------I---I-A------V--L------SA-----L---V--A------CY-A-D------------------------IPS---------L------------------E--------P--M-------------SLV--Y--DEDDTSI------D---G---------------------------------------------------------------------------------------------------------------------------------------------------------------------------------------------------------------------------K----------LMS--------SAD---V---NL-KN--------------------TAW------VVI--S-----------EWQ------------------CNAP-AVAG-----L-V-QRVRVHY----DG-----A-----P--------N--------------LT-INR------------L---------I---V-----T-Y-----------------T--V-T--PAA--VD-W--T----PLSTNVMTVIIRS-HAG--QQVNA-N-------VTL-L---------------------------------------------------------------------------------------------------------------------------------------------------------------R--------------R----------------------------------------------------------------------------------------------------------------------------------------------------------------A

>Slitu-alphaREPAT11

---------MR-----SF-I----------I---I-T------V--L------SA-----L---V--A------CY-S-A---S---------------V----LPS---------E------------------E--------E--M----------I--TVI--Y--D--ENFP------N---G---------------------------------------------------------------------------------------------------------------------------------------------------------------------------------------------------------------------------T----------IVS--------KEE---L---DS-RFT--------RS------------------GLV--S-----------SWV------------------CTVP-AISR-----Q-S-QVITLQY----LG-----D-----S--------R--------------DL-VRR------------VVHSYTGS-PT---M-----T-H------------------------------------S----PLITNNMFVTITS-RTG--QEINS-R-------VEL-Y---------------------------------------------------------------------------------------------------------------------------------------------------------------L--------------Y-----------------------------------------------------------------------------------------------------------------------------------------------------------------

>Slitu-alphaREPAT12

---------MR-----TF-I----------I---I-T------V--L------SA-----L---A--A------CY-GAA-------------------V----LPS---------E------------------K-V----MSE--G----------I--TIV--Y--D--VDSP------N---G---------------------------------------------------------------------------------------------------------------------------------------------------------------------------------------------------------------------------R----------VMS--------RSD----------IL---------------DPLF----------FLL--S-----------EST------------------INVP-AIPG-----E-V-QKAGITY----QA-----P-----V--------G--------------VT-IGR------------V---------T---I-----TLY-----------------A--N-L--PAD--TQ-N--S----PLGSNYVESLVTS-RPG--HAING-S-------IAV-Y---------------------------------------------------------------------------------------------------------------------------------------------------------------R----------------------------------------------------------------------------------------------------------------------------------------------------------------WA-D------------

>Slitu-alphaREPAT13

---------MR-----AF-I----------I---I-T------M--L------SA-----L---A--A------CY-GGA-------------------V----LPS---------E------------------K-V----MSE--G----------I--TIV--Y--D--VDSP------N---G---------------------------------------------------------------------------------------------------------------------------------------------------------------------------------------------------------------------------R----------VMS--------RSD----------IL---------------DTLF----------VLL--S-----------EST------------------INVP-AIPG-----E-V-QKAGITY----QA-----P-----V--------G--------------VT-IGR------------V---------S---I-----TLY-----------------A--Y-L--PAD--TQ-N--S----PLGSNYVESLVTS-RPG--DAING-S-------IAV-Y---------------------------------------------------------------------------------------------------------------------------------------------------------------R----------------------------------------------------------------------------------------------------------------------------------------------------------------WA-D------------

>Slitu-alphaREPAT14

---------MR-----TF-I----------I---I-T------V--L------SA-----L---A--A------CY-S-AA------------------D----LPS---------E------------------E------ISG--G----------Y--TVL--Y--D--DMSP------E---G---------------------------------------------------------------------------------------------------------------------------------------------------------------------------------------------------------------------------R----------IVS--------SAE---L---N---A---------------NV-W----------IPI--S-----------EAS------------------LLAP-AVAG-----Q-R-QYITKVF----QA-----T-----P--------G--------------VI-ING------------L---------S---V-----T-Y-----------------S--K-T--PAY--SY-H--S----SLGSDSMIVSITS-HPG--DEISS-K-------ITV-Y---------------------------------------------------------------------------------------------------------------------------------------------------------------R--------------R----------------------------------------------------------------------------------------------------------------------------------------------------P------------

>Slitu-alphaREPAT15

---------MR-----SF-I----------I---I-T------V--L------SA-----F---A--A------CY-SAA-------------------V----LPR---------E------------------E-E----M---------------V--TII--Y--D--DKSP------N---G---------------------------------------------------------------------------------------------------------------------------------------------------------------------------------------------------------------------------T----------ILS--------RAD---F---NA-RS--------------HQG-W----------VLL--S-----------ETN------------------DFVP-AVEG-----Q-V-QSVGVFY----RA-----N-----F--------G--------------VR-IDR------------L---------V---I-----TTY-----------------G--H-L--IPD--VD-R--T----PLGSNKMIVILTS-RPG--ELIRS-T-------IET-Y---------------------------------------------------------------------------------------------------------------------------------------------------------------K--------------R----------------------------------------------------------------------------------------------------------------------------------------------------DNISS--------

>Slitu-alphaREPAT16

---------MR-----TF-I----------I---I-T------V--L------SA-----L---A--A------CY-S-AA------------------D----LPN---------E------------------E------ISG--G----------Y--TVL--Y--D--DMSP------E---G---------------------------------------------------------------------------------------------------------------------------------------------------------------------------------------------------------------------------R----------IVS--------GAE---L---NA-KS---------------VV-W----------VPI--S-----------EAA------------------FEAP-AVAG-----Q-V-QHITKLF----KA-----A-----P--------G--------------VI-INA------------L---------I---V-----S-Y-----------------S--K-T--PME--SY-R--S----PLGSDSMIVSITS-HPG--DEISS-T-------VSI-F---------------------------------------------------------------------------------------------------------------------------------------------------------------S--------------R----------------------------------------------------------------------------------------------------------------------------------------------------H------------

>Slitu-alphaREPAT17

---------MR-----SF-I----------I---I-T------V--L------SA-----L---V--A------CY-GAA-------------------V----LPS---------E------------------E-E----M---------------V--TII--Y--D--DKSP------N---G---------------------------------------------------------------------------------------------------------------------------------------------------------------------------------------------------------------------------I----------IMS--------RAD---F---NA-RS---------------QG-W----------VLL--S-----------EAN------------------DLIP-AVEG-----Q-E-QSIRMVY----RA-----Q-----R--------G--------------VK-INM------------L---------S---V-----SAR-----------------L--S-G--IPE--VN-N--T----PLGRNYIDVKVTS-RPG--LGLNS-T-------ISV-Y---------------------------------------------------------------------------------------------------------------------------------------------------------------R--------------W----------------------------------------------------------------------------------------------------------------------------------------------------D------------

>Slitu-alphaREPAT18

---------MR-----TF-I----------I---I-T------V--L------SA-----L---A--A------CY-S-AA------------------D----LPN---------E------------------E------ISG--G----------Y--TVL--Y--D--DMSP------E---G---------------------------------------------------------------------------------------------------------------------------------------------------------------------------------------------------------------------------R----------IVS--------SAE---L---N---A---------------NV-W----------IPI--S-----------EAT------------------LLAP-AVAG-----Q-R-QYITKVF----QA-----A-----P--------G--------------VI-ING------------L---------S---V-----T-Y-----------------S--K-T--PAY--SY-R--S----SLGSDSMIVSITS-HPG--DEISS-K-------ITV-Y---------------------------------------------------------------------------------------------------------------------------------------------------------------R--------------R----------------------------------------------------------------------------------------------------------------------------------------------------P------------

>Slitu-alphaREPAT19

---------MR-----SF-I----------I---I-T------V--L------SA-----L---V--A------CY-GAA-------------------V----LPS---------E------------------E-E----M---------------V--TII--Y--D--DKSP------N---G---------------------------------------------------------------------------------------------------------------------------------------------------------------------------------------------------------------------------I----------IMS--------RAD---F---SA-RS---------------QG-W----------VLL--S-----------EAN------------------DLIP-AVEG-----Q-E-QSIRMVY----RA-----Q-----R--------G--------------VK-INM------------L---------S---V-----SAR-----------------L--S-G--IPE--VN-N--T----PLGRNYIDVKVTS-RPG--LGLNS-T-------ISV-Y---------------------------------------------------------------------------------------------------------------------------------------------------------------R--------------W----------------------------------------------------------------------------------------------------------------------------------------------------D------------

>Slitu-alphaREPAT2

---------MR-----SF-I----------I---I-A------V--L------SA-----L---A--A------SY-G-T---P---------------V----LPS---------G------------------E--------E--T-------YPEY--KMIHKYNKE--DDFQ------N---R---------------------------------------------------------------------------------------------------------------------------------------------------------------------------------------------------------------------------I----------TIL--------REGMRH-------DN-----------------------DRQDGLVLL--D-----------EQR------------------FIRT-AIPD-----V-I-QHVDILY----YG-----K-----P--------P--------------TV-ITT------------V---------E---I-----T-S-----------------S--P-P--PSL--VF-P--V----WHLDNYISIRISS-QRG--LGMNA-T-------VQI-F---------------G-TL-----------------------------------------------------------------------------------------------------------------------------------------------------------------------------------------------------------------------------------------------------------------------------------------------------------------------------

>Slitu-alphaREPAT20

---------MR-----SF-I----------I---V-T------V--L------SA-----L---V--A------SY-N-A---S---------------V----LPD---------A------------------E--------Q--M---PSNF---I--YTV--Y--D--GNSP------N---G---------------------------------------------------------------------------------------------------------------------------------------------------------------------------------------------------------------------------T----------IVA--------LND---I---TI-KS-------------------S---------FLL--S-----------TWR------------------NNVQ-PIPG-----E-E-QRIEMKY----EG-----I-----P--------G--------------DR-ITT------------I---------M---V-----G-H-----------------S--T-T--PTN--IY-H--S----PLFDYYIDMQIVS-HRG--QGIDI-Y-------VAI-Y---------------------------------------------------------------------------------------------------------------------------------------------------------------G--------------R-----------------------------------------------------------------------------------------------------------------------------------------------------------------

>Slitu-alphaREPAT21

---------MR-----SF-I----------I---I-T------V--L------SA-----L---A--A------CY-G-VA------------------V----PPS---------E------------------E------VYG--G----------Y--TLL--F--D--DISP------E---G---------------------------------------------------------------------------------------------------------------------------------------------------------------------------------------------------------------------------R----------IVS--------SAE---L---NA-GS---------------DR-W----------VKL--S-----------ESI------------------LDVP-AVPG-----H-V-QRIGTKY----EG-----D-----A--------S--------------VV-ITR------------W---------S---V-----T-------------------S--D-V--PTA--SY-S-FS----PFGSNLMEVTIQS-HLG--NRIYS-T-------VTI-Y------------GVI------------------------------------------------------------------------------------------------------------------------------------------------L--------------N----------------------------------------------------------------------------------------------------------------------------------------------------K------------

>Slitu-alphaREPAT22

---------MR-----SF-I----------I---I-T------V--L------SA-----L---A--A------CY-S-A-------------------D----LPS---------E------------------A------MTE--D----------Y--TMV--F--D--EMSP------N---R---------------------------------------------------------------------------------------------------------------------------------------------------------------------------------------------------------------------------K----------IAS--------NAE---F---HA-NS---------------DG-W----------ILI--S-----------RLK------------------LY------------K-T-KTKTTIY----IK-----H-----R--------K--------------IR-PHRRQWGKYPGGWQHC---------H---V-----V-W-----------------Q--C-L--TSD--RE-K--D----QPSGHEM-----C-QPG------------------V-L---------------------------------------------------------------------------------------------------------------------------------------------------------------K--------------C----------------------------------------------------------------------------------------------------------------------------------------------------F------------

>Slitu-alphaREPAT23

---------MR-----SF-I----------I---I-T------V--L------SA-----L---A--A------CY-G-AA------------------V----VPS---------E------------------E------MSE--G----------C--TMI--F--D--EMSP------D---G---------------------------------------------------------------------------------------------------------------------------------------------------------------------------------------------------------------------------R----------IVP--------NAE---L---NA-KS---------------SE-W----------ILI--S-----------EGV------------------VNAP-AVPG-----Q-V-QRVGTWH----QA-----D-----L--------G--------------VR-IER------------L---------A---L-----S-Y-----------------G--A-F--PAD--VQ-H--T----PLGSNFLEVKIIS-ALG--QRVAA-T-------ISV-Y---------------------------------------------------------------------------------------------------------------------------------------------------------------R--------------R---------------------------------------------------------------------------------------------------------------------------------------------------GC----------C-

>Slitu-alphaREPAT24

---------MR-----SF-I----------I---I-V------V--L------SA-----L---A--A------CY-G-A---A---------------V----LPGEANMFSSFIQ------------------E--------Q--M-------LDGW--TVE--HNEE--DVAQ------N---R---------------------------------------------------------------------------------------------------------------------------------------------------------------------------------------------------------------------------V----------TLS--------KGD--------A-RD-----------------------D--L--IFL--F-----------ERA------------------CYVP-ATPN-----D-V-GRCDVVY----YG-----E-----P--------N--------------IR-IRR------------I---------T---IS----S-S-----------------W--P-P--PAY--SH-T--S----GLTTNVGRVRLIS-QIG--ERLHA-V-------VRM-Y---------------GLIN-----------------------------------------------------------------------------------------------E----------------------------------------DNNES--------------------------------------------------------------------------------------------------------------------------------------------------------------------------------

>Slitu-alphaREPAT25

---------MR-----SF-I----------I---I-V------V--L------SA-----L---A--A------CY-G-A---A---------------V----LPGEANMFSSFIQ------------------E--------Q--M-------LDGW--TVE--NNEE--DAAQ------N---R---------------------------------------------------------------------------------------------------------------------------------------------------------------------------------------------------------------------------V----------TMS--------KGE--------A-RD-----------------------D--L--IFL--F-----------ERA------------------CYVP-ATPN-----D-V-GRFDVVY----YG-----E-----P--------N--------------VS-IRR------------F---------S---IS----F-T-----------------W--P-V--PAH--IY-I--S----GLLSNVGRVRLTS-HIG--ERLQA-V-------VRI-Y---------------GLII-----------------------------------------------------------------------------------------------E----------------------------------------DNNES--------------------------------------------------------------------------------------------------------------------------------------------------------------------------------

>Slitu-alphaREPAT26

---------MT-----SF-I----------I---I-T------V--L------SA-----L---A--A------CY-G-A---V---------------V----PPSEENMISRFAL------------------E--------Q--M-------YPDF--TVE--YNEK--Y--E------N---R---------------------------------------------------------------------------------------------------------------------------------------------------------------------------------------------------------------------------V----------AFS--------KGE--------A-RN-----------------------D--Q--ILL--L-----------TRE------------------IHVP-AVPG-----E-T-QYADLIH----RS-----S-----S--------N--------------VR-ITR------------V---------V--WIP----L-T-----------------W--P-L--PTL--IS-I--K----GVGTDVVHISCTS-FVG--ENIHV-L-------FSL-Y---------------GEII-----------------------------------------------------------------------------------------------E----------------------------------------N------------------------------------------------------------------------------------------------------------------------------------------------------------------------------------

>Slitu-alphaREPAT27

---------MR-----SF-I----------I---I-L------V--L------SA-----L---A--A------CH-G-A---A---------------V----LPDESNMFSSFIR------------------E--------Q--M-------LDGW--TVE--HNEE--DAVQ------N---R---------------------------------------------------------------------------------------------------------------------------------------------------------------------------------------------------------------------------V----------TMN--------KGE--------A-RD-----------------------D--L--IFL--Y-----------ESP------------------CIVP-ALPN-----D-I-VTCEVVY----HG-----E-----P--------N--------------VS-IRR------------F---------S---VS----Y-F-----------------W--P-P--PAH--ID-V--S----GFLSNVGRVKLTS-HIG--ERLQA-V-------IGI-Y---------------GVIN-----------------------------------------------------------------------------------------------E----------------------------------------DNN----------------------------------------------------------------------------------------------------------------------------------------------------------------------------------

>Slitu-alphaREPAT3

---------MK-----SF-I----------V---I-I------V--L------SA-----L---V--A------SY-E-A---F---------------V----LPS---------R--VISGRHDAFVSSRQMTD--------A--F-------------TLV--Y--D--DTAV------N---G---------------------------------------------------------------------------------------------------------------------------------------------------------------------------------------------------------------------------T----------IMT--------SAE---V---EA-KN--------------------DGW------RLL--Y-----------EVS------------------CSAP-AAVG-----R-V-KNSFVTY----QG-----P-----T--------N--------------TV-IDR------------I---------T---I-----T-P-----------------Q--I-N--PAN--VY-H--S----ALGTNNMEVRLLS-NIS--QPIQA-T-------VRL-Y---------------------------------------------------------------------------------------------------------------------------------------------------------------Q-------------QR------------------------------------------------------------------------------------------------------------------SGAGKAFGEPAFHLMFFVLIYFGISNF-------------------T

>Slitu-alphaREPAT4

---------MR-----SF-I----------I---I-T------V--L------SV-----L---A--A------CY-G-F---A---------------D----LIN---------E---------------------------E--VQER-------Y--TML--Y--D--EKSP------N---G---------------------------------------------------------------------------------------------------------------------------------------------------------------------------------------------------------------------------R----------IVS--------DAE---L---NK-ISE--E----FSI--------------------L--A-----------QWY------------------CQSP-ALPD-----K-N-QRVGVMY----EG-----P-----P--------GT-------------TM-VIY------------LVFYHPPK-PK---I-----E-H------------------------------------S----KWGSNFLSINITS-PIG--ISVDA-V-------VAL-Y---------------------------------------------------------------------------------------------------------------------------------------------------------------G--------------N---------------------------------------------------------------------------------------------------------------------------------------------FPK-----------------

>Slitu-alphaREPAT5

---------MR-----SF-I----------F---I-T------V--L------SA-----L---T--A------CY-G-A---S---------------V----LPD---------N------------------N--------E--M----------I--TVV--Y--D--DEAP------N---G---------------------------------------------------------------------------------------------------------------------------------------------------------------------------------------------------------------------------R----------IVS--------DAE---F---KN-SVIIPDNWVLLSD------------------GII--Y-----------RWR------------------CDAP-PINN-----Q-V-QRVGVRY----EG-----D-----P--------SI-------------RI-GQV------------SVSYFPPA-PH---I-----A-Y------------------------------------G----QLGNYFMEVNITT-PVG--REVRS-T-------VGMYY---------------------------------------------------------------------------------------------------------------------------------------------------------------R--------------A-----------------------------------------------------------------------------------------------------------------------------------------------------------------

>Slitu-alphaREPAT6

---------MK-----SL-I----------L---I-T------V--L------SA-----L---V--A------CY-E-A-----------------------------------------SAE---------LPG--------G--Y-------------SVV--Y--D--DTV-------D---G---------------------------------------------------------------------------------------------------------------------------------------------------------------------------------------------------------------------------N----------VMP--------NTE---I---NA-RN--------------------QNF------RLI--Y-----------EVV------------------AVAS-AAIG-----R-V-KNSFVTY----QG-----P-----N--------N--------------VR-VDR------------L---------V---I-----T-N-----------------H--V-Q--PAS--VF-H--S----TLGTNNMEVRFVS-NVS--QPISS-T-------VRM-Y---------------------------------------------------------------------------------------------------------------------------------------------------------------Q--------------R------------------------------------------------------------------------------------------------------------------SGAGRAFGEPAFHLVVLVLMYFGFSQL-------------------A

>Slitu-alphaREPAT7

---------MR-----SF-I----------I---I-T------G--L------LA-----L---A--V------CC-N-A---S---------------A----LTR---------Q------------------E--DPSE--V--Y----------V--TTT--E--Y--E---------N---G---------------------------------------------------------------------------------------------------------------------------------------------------------------------------------------------------------------------------T----------IVD--------GAD---FAFSNV-GS------------------------------LL--S-----------EWH------------------CEAP-SIVG-----Q-S-QTITLKY----LG-----D-----S--------S--------------ER-ITR------------RVTSIYGD-PQ---L-----I-Q------------------------------------T----PLGSNVYEMTLTS-RPG--VRIAA-S-------FYI-Y---------------------------------------------------------------------------------------------------------------------------------------------------------------G--------------------------------------------------------------------------------------------------------------------------------------------------------------------------------

>Slitu-alphaREPAT8

---------MR-----SF-I----------I---I-A------V--L------SA-----L---A--A------CY-G-A---S---------------V----VPS---------E------------------D--------G--Y----------I--TTI--Y--D--EKSP------N---G---------------------------------------------------------------------------------------------------------------------------------------------------------------------------------------------------------------------------T----------IVS--------NGD---IEFTNE-RS-----------------------------TIY--A-----------QWE------------------SEIP-TSID-----R---HTVSFMY----VG-----N-----P--------S--------------DI-ITS------------IRTGIYGE-PE-I-V-----Y-R------------------------------------S----PLNTNIFQMILRT-RAG--QRMRA-T-------IQI-S---------------------------------------------------------------------------------------------------------------------------------------------------------------R--------------------------------------------------------------------------------------------------------------------------------------------------------------------------------

>Slitu-alphaREPAT9

---------MR-----SF-I----------I---I-T------V--L------SA-----L---A--A------C--S-V---A---------------D----LPS---------E------------------E--------E--M----------Y--TIV--Y--D--EKYP------N---G---------------------------------------------------------------------------------------------------------------------------------------------------------------------------------------------------------------------------T----------IVS--------DAE---F---DS-RFK--------SV------------------IPL--A-----------EWR------------------CRSP-AISR-----R-Q-QTVTVAY----HG-----D-----P--------R--------------DI-ITR------------TTFSQSGN-PR---I-----S-Q------------------------------------T----PLMRNSILITITS-RSG--VGIDS-L-------VQI-F---------------------------------------------------------------------------------------------------------------------------------------------------------------T--------------------------------------------------------------------------------------------------------------------------------------------------------------------------------

>Slitu-betaREPAT1

---------MS------L-L----------T---Q-I---------T-------I-----L------C-----------L---C---------------L---------------------------------------------------------------------------A------L---Y--NAED-------------------------------------------------------------------------------------------------------------------------------------------------------------------------------------------------------L---------QLG-S----------GVN--------A----------------------------------------Q--LGY--Q-----------EIV------------------K-LG-SVPL-----T-V-RTKNVFY----NS-----N-----N--------T--------------KT-IKS------------I---------S---A-----V-D-----------------I--D-K--S-K--AV-P--TITAGGIGSSFVNVRLKS-EKG--KALNY-L-------VQI-F---------------V--------------------------------------------------------------------------------------------------------------------------------------------------------------------------------------------------------------------------------------------------------------------------------------------------------------------------------

>Slitu-betaREPAT10

MA-------VK------A-L----------L---SII------L--L-------L--S--I---S--V-----------S---L---------------V-LAR-----------------------------------------------------------------------G------R---G-----S-------------------------------------------------------------------------------------------------------------------------------------------------------------------------------------------------------L---------SIG-N----------TFG--------S----------------------------------------H--LIY--Q-----------ETH------------------Y-KY-GIPF-----I-V-RDEDITV----KG-----V-----N--------N--------------EI-IEA------------V---------V---V-----N-D-----------------L--E-G----N--GL-C--YIKTGGVGHHDVTIHLES-SRG--HGYKF-L-------VEV-Y---------------A--------------------------------------------------------------------------------------------------------------------------------------------------------------KKYYK-------------------------------------------------------------------------------------------------------------------------------------------------------------

>Slitu-betaREPAT11a

----MFKH--------------------------L-L------I--L-------C-----A---I--V-----------V---A---------------V---------------------------------------------------------------------------H------C---K-----S-------------------------------------------------------------------------------------------------------------------------------------------------------------------------------------------------------V---------IVG-Q----------TWEQNGV---------------------------------------------R--KVY--E-----------TTV------------------S-AN-SIPL-----F-K-RTKYVNF----EF-----P-----D--------Q-I------------SK-IKG------------I---------A---I-----K-D-----------------Q--D-N--G-L--AE-P--SITMGGLGFNFANLKLKS-ERG--SGFNF-L-------IEI-Y---------------G--------------------------------------------------------------------------------------------------------------------------------------------------------------------------------------------------------------------------------------------------------------------------------------------------------------------------------

>Slitu-betaREPAT11b

----MFKE--------------------------I-V------I--L-------F-----C---V--V-----------V---L---------------V---------------------------------------------------------------------------Q------S---N-----S-------------------------------------------------------------------------------------------------------------------------------------------------------------------------------------------------------V---------FVG-H----------VYDQTGL----V----------------------------------------R--KVY--D-----------KMV------------------E-AT-GIPL-----I-K-REEEIYY----EY-----P-----A--------G-D------------QK-IKG------------I---------A---I-----K-D-----------------L--T-D--S-K--AQ-A--SITRGGLGFGFVNIKLKS-ERG--SGYKY-L-------IEV-F---------------A--------------------------------------------------------------------------------------------------------------------------------------------------------------------------------------------------------------------------------------------------------------------------------------------------------------------------------

>Slitu-betaREPAT12

---------MR------T-S-------SI-A---I-L------L--C-------L-----L---A--A-----------A---A---------------S---------------------------------------------------------------------------Q------S---H-----D-------------------------------------------------------------------------------------------------------------------------------------------------------------------------------------------------------L---------ILG-Q----------ATY--------G----------------------------------------D--VIV--F-----------KAD------------------EVKY-GFPL-----F-V-RTSIVEY-----P-EQGQK-----N--------Y--------------AY-IKA------------I---------Y---V-----K-DNE---------------R--D-G--S-G--GY-P--SISAGGIGQRFVKIKLKS-QRG--SGLNF-T-------VTI-Y---------------G--------------------------------------------------------------------------------------------------------------------------------------------------------------R-----------------------------------------------------------------------------------------------YV----------------------------------------------------------------

>Slitu-betaREPAT13a

---------MF------T-K----------V---V-A------L--F-------C-----L------V-----------A---V---------------A---------------------------------------------------------------------------A------A---T-----D-------------------------------------------------------------------------------------------------------------------------------------------------------------------------------------------------------L---------TVG-K---------R--D--------G----------------------------------------R--LIF--Y-----------TTI------------------T----ARPT-----IWK-QVQNLTV----NA-----T-----D--------D--------------AL-ITR------------V---------V---V-----I-D-----------------NRPE-K----D--GE-A--RVVDGGVGHNNVTIELKS-PTV-FRGFNF-T-------VKV-F---------------E--------------------------------------------------------------------------------------------------------------------------------------------------------------R----KVGE--------Q-----NRRH--------PVFVG----------------TD-TQK----------------------------------------------------------------------------------------------------

>Slitu-betaREPAT13b

---------MF------T-K----------V---V-A------L--F-------G-----L------V-----------A---L---------------A---------------------------------------------------------------------------A------A---A-----D-------------------------------------------------------------------------------------------------------------------------------------------------------------------------------------------------------L---------SVG-K---------E--G--------G----------------------------------------Q--LIF--D-----------KNV------------------T----ASPA-----IWK-QVQNLTV----NA-----T-----D--------D--------------AL-ITR------------V---------V---V-----I-D-----------------NRPE-K----D--GE-A--KVVDGGVGHNNITIELKS-PTV-FRGFDF-T-------VKV-F---------------A--------------------------------------------------------------------------------------------------------------------------------------------------------------N----EGAV--------Q-----VPQH--------PVSVG----------------AN-TQV----------------------------------------------------------------------------------------------------

>Slitu-betaREPAT15

---------MKLI----L-V----------G---L-V------V--C-------L-----S---V----------------------------------V---------------------------------------------------------------------------Y------A---N-----H-------------------------------------------------------------------------------------------------------------------------------------------------------------------------------------------------------L---------IVGNV----------ANR--------V----------------------------------------V--LAN--Q-----------ARV------------------E-YN-AIPF-----I-K-RVKYFFY----TQ-----P-----D--------N--------------RL-IQG------------I---------Q---A-----L-D-----------------T--M-H--S-K--AS-I--NITAGGVGYSYVNLRLKS-ERG--RGLAY-D-------IGI-Y-VQDLFYH----------------------------------------------------------------------------------------------------------------------------------------------------------------------------------------------------------------------------------------------------------------------------------------------------------------------------------------

>Slitu-betaREPAT16

---------MR------S-F---LIFC---V---F-I------L--T-------L-----S---S--V---------------L---------------S---------------------------------------------------------------------------S------TYFFG-----E---------------------------------------------------------------------------------------------------------------------------------------------------------------------------------------------------------------------Q----------TKN--------T----------------------------------------R--IIY--Q-----------DKV------------------R-YE-AIPL-----K-K-RVKTYSY----NG-----T-----V------------------------P-IKA------------I---------S---C-----Y-D-----------------Y--Q-N--S-E--AS-V--NITAGGIGYKYVELRLKS-QRS--YRLDY-A-------IEI-Y---------------A--------------------------------------------------------------------------------------------------------------------------------------------------------------------------------------------------------------------------------------------------------------------------------------------------------------------------------

>Slitu-betaREPAT17

---------MD------L-L----------L---I-V------L--L-------A-----I---I--T-----------D---A------KMI------C---------------------------------------------------------------------------K------K---T-----K-------------------------------------------------------------------------------------------------------------------------------------------------------------------------------------------------------A---------PL-------------------------------------------------------------------------------------ELG------------------EWRY-DLKH-----M-D--SAVVEN-----P-----P-----V--------S--------------IN-EGQ------------M---------Y---V-----Y-ENYFPNYTIKYIHVDNLAI--R-S--C-G--AS-A--AMRTGGIGSSYVLIVLHA-HIN--EEIRS-V-------VDI-W---------------G-------ERK--------KVITE---ELP--RTPL--KP----DEMTSLKSLYLFKHLRAVNHNS----AY---------------------------------------------------------------------------------------------------------------------------------------------------------------------------------------------------------------------------------------------------------

>Slitu-betaREPAT2

---------MK-C-V--L-A----------L---S-V------F--V-------V-----I---A--I-----------S---S---------------C---------------------------------------------------------------------------L------A---S-----D-------------------------------------------------------------------------------------------------------------------------------------------------------------------------------------------------------L---------IIG-T----------SYN--------A----------------------------------------R--LAW--Q-----------HKS------------------D-YM-GIPF-----K-K-RVKEVFY----SD-----P-----G--------Q--------------QI-IKG------------I---------I---A-----R-D-----------------I--D-H--T-S--GS-A--TVTAGGVGFSFANIRLKS-ERG--SGLNY-Q-------IEI-Y---------------V--------------------------------------------------------------------------------------------------------------------------------------------------------------------------------------------------------------------------------------------------------------------------------------------------------------------------------

>Slitu-betaREPAT3

---------MK-T-A--P-V----------F---L-V------V--L-------L-----S---I--I-----------C---V---------------V---------------------------------------------------------------------------H------C---K-----D-------------------------------------------------------------------------------------------------------------------------------------------------------------------------------------------------------L---------IVG-S----------SFN--------N----------------------------------------R--LIH--Q-----------EKA------------------E-YN-AIPL-----K-K-RVKEVFF----SD-----P-----G--------Q--------------QI-IKG------------I---------I---L-----R-D-----------------L--D-H--T-E--AI-P--SVTAGGVGFSYANIRLKS-ERG--SGLNY-Q-------VEI-Y---------------V--------------------------------------------------------------------------------------------------------------------------------------------------------------------------------------------------------------------------------------------------------------------------------------------------------------------------------

>Slitu-betaREPAT4

---------MR------F-V----------F---L-L------L--L-------V-----L---C--L-----------L---C---------------V---------------------------------------------------------------------------S-VP-VLC---S-----D-------------------------------------------------------------------------------------------------------------------------------------------------------------------------------------------------------M---------IVG-D----------TVH--------R----------------------------------------K--MIF--H-----------QRI------------------K-DF-AIPF-----K-K-RVKTLTF----SD-----P-----E--------K--------------RK-IKG------------V---------A---V-----V-D-----------------N--D-F--S-H--AS-A--NITEGGVGYSYVTVRMKS-QRH--HPLNF-E-------VEI-Y---------------V--------------------------------------------------------------------------------------------------------------------------------------------------------------------------------------------------------------------------------------------------------------------------------------------------------------------------------

>Slitu-betaREPAT5

---------MK-L----V-L----------V---L-F------A--S-------I-----F---V--V-----------A---T---------------C--------------------------------------------------------------------------------------D-----H-------------------------------------------------------------------------------------------------------------------------------------------------------------------------------------------------------L---------ILGNT----------NNN--------Q----------------------------------------N--MIY--H-----------TTS------------------H-YS-AIPF-----I-K-RVKNVFY----SG--------------------S--------------SV-INS------------I---------M---A-----Y-D-----------------N--K-H--T-N--AS-A--AVTAGGIGYTYVNLRLKS-ERG--KELDY-D-------IGI-Y---------------S--------------------------------------------------------------------------------------------------------------------------------------------------------------------------------------------------------------------------------------------------------------------------------------------------------------------------------

>Slitu-betaREPAT6

---------MK------C-V----------L-----------------------------I---T--VCLIL-------L---C---------------V---------------------------------------------------------------------------C------L---N--EGKD-------------------------------------------------------------------------------------------------------------------------------------------------------------------------------------------------------F---------IVG-T----------KAN--------N----------------------------------------L--LIS--T-----------QKL------------------K-YR-SIPL-----I-R-RDKDYTY----ID-----S-----K--------E--------------RI-IKG------------I---------I---A-----R-D-----------------L--S-R--T-D--TE-V--TVTSGGVGATNVTLHFQS-GRG--EELNY-L-------ILI-F---------------S---NIK--------------------------------------------------------------------------------------------------------------------------------------------------------------------------------------------------------------------------------------------------------------------------------------------------------------------------

>Slitu-betaREPAT7a

---------MK------V-V----------A---L-A------F--L-------L-----A---I--V-----------V---I---------------V---------------------------------------------------------------------------D------C---S-----H-------------------------------------------------------------------------------------------------------------------------------------------------------------------------------------------------------T---------FMG-T----------NVL--------R----------------------------------------P--LIY--H-----------HNV------------------E-YG-SKVF-----R-K-RVENLYY----SL-----P-----A---VPANAG--------------RS-IQG------------I---------L---A-----F-D-----------------M--T-H--T-G--AS-A--NVTQGGLGFNFVNLRMKS-DRG--EKIHF-D-------VYI-Y---------------A--------------------------------------------------------------------------------------------------------------------------------------------------------------------------------------------------------------------------------------------------------------------------------------------------------------------------------

>Slitu-betaREPAT7b

---------MH------I-S----------S---V-A------L--V-------L-----A---M--I-----------V---V---------------S---------------------------------------------------------------------------E------C---G-----H-------------------------------------------------------------------------------------------------------------------------------------------------------------------------------------------------------T---------FLG-T----------SVQ--------R----------------------------------------P--LVF--H-----------KEV------------------R-YP-ARGF-----Q-K-RIENVNF----TL-----P-----H---VSDIYL--------------RN-IQG------------I---------L---A-----Y-D-----------------L--T-N--S-G--AS-A--NVTKGGLGYNFVNIRLKS-DRN--KEIRY-D-------INI-Y---------------A--------------------------------------------------------------------------------------------------------------------------------------------------------------------------------------------------------------------------------------------------------------------------------------------------------------------------------

>Slitu-betaREPAT8a

---------MF----F-K-V----------T---L-L------V--S-------L-----L---Y--L-----------T---S---------------V---------------------------------------------------------------------------E------S---N-----D-------------------------------------------------------------------------------------------------------------------------------------------------------------------------------------------------------I---------RLG-F---------ATPY--------S----------------------------------------K--KIY--S-----------AIK------------------I----ANPA-----LWQ-REDDVLI----SS-----P-----N--------H--------------EL-IDA------------V---------Y---I-----T-D-----------------LRPE-K----D--GI-A--YIEAGGIGSKTVTIALKS-PSL-LRGYKF-A-------IEV-Y---------------A--------------------------------------------------------------------------------------------------------------------------------------------------------------Q----S-LHHGYGTNILPIQ---GAAYPRAG--LSGYG---------------------SQYP--GYPTAYSGVGVYP--NNY---G-------KK------------------------------------------------------------------

>Slitu-betaREPAT8f

---------MY----A-K-V----------L---L-T------L--F-------V-----L---Y--L-----------S---T---------------V---------------------------------------------------------------------------Q------C---N-----D-------------------------------------------------------------------------------------------------------------------------------------------------------------------------------------------------------I---------RIG-Y---------TNLR--------S----------------------------------------I--KIY--S-----------ELK------------------E----ADPA-----LWK-RTEEVTI----KA-----P-----P--------H--------------EF-IDS------------I---------H---V-----S-D-----------------LRED-K----D--GE-A--YITSGGIGMPYVTISLKS-PTI-LRGYKF-Q-------IEV-Y---------------A--------------------------------------------------------------------------------------------------------------------------------------------------------------S----P-LV--------S-----NEGLFSKG--SFNYGEA----------------QQVQQYP-----------------------R-------KF------------------------------------------------------------------

>Slitu-betaREPAT9

---------MG------F-R----------I---V-I------F--N-------Y-----I---T--L-----------V---S---------------I---------------------------------------------------------------------------T------C---H-----D-------------------------------------------------------------------------------------------------------------------------------------------------------------------------------------------------------F---------FIG-S---------PDSS--------S----------------------------------------K--LI---------------KGN------------------E----LKPA-----IWI-RSDILSV----NC-----S-----G--------N--------------DV-ISS------------V---------Q---I-----M-D-----------------LIPE-K----L--SE-A--RLKSGGIGSKFVVIELRS-SD--YEGLRF-V-------VRM-F---------------V--------------------------------------------------------------------------------------------------------------------------------------------------------------F----P--S--------S--LRHRYM--------------------------------------------------------------------KI------------------------------------------------------------------

>Slitu-gammaREPAT5

---------MK-----SI-F----------V---I-V------L--W-------AGC---A---I--V-----------G---Y---------SS----A---------------------------------------------------------------------------T------Q---V-----E----------------------------------------------------------------------------------------------------------------N---------------------------------------------EA------------RG---------------------G---L---------SLG-T----------LGP--------D----------------------------------------D--RLL--S-----------NTN------------------HAVE-AAVY-----V-V-QTQEVLI----RT-----V-----Q--------Y--------------SS-ITS------------I---------H---V-----S-E-----------------V--G-T--TTN--AS-V--SIIEGGLRTNFVKFLFQS-QKG--RAYSY-N-------IRI-H---------------G--------------------------------------------------------------------------------------------------------------------------------------------------------------T----------------------------------------------------------------------------------------------------------IEC----------------------------------------------------

>Slitu-gammaREPAT6

---------MK-----IF-I----------V---L-V------V--L-------------A---A--A-----------A---A---------AG----V---------------------------------------------------------------------------T------N---N-----DR-----E--Q--------AA--------ALENKELHLPDIVDNNE-LD---VP---EKNEPNVPDRPDNTELDVPEDEI-GEEDEETGL-----------------------A---------------------------------------------AA------------RN---------------------N---L---------SLG-T----------FGA--------D----------------------------------------D--RLM--S-----------RIT------------------HNVA-AATL-----L-I-HDQEITF----RG-----T-----R--------G--------------TN-ITA------------I---------R---V-----N-Q-----------------V--G-T--NTP---I-A--TRVAGGLGSEFVTIRIQS-GRG--RAFSY-T-------ILV-Y---------------A--------------------------------------------------------------------------------------------------------------------------------------------------------------A----------------------------------------------------------------------------------------------------------INC--SNN-----------------------------------------------

>Slitu-gammaREPAT13

---------MK-----SF-M----------F---V-V------L--L-------A-----V---T--L-----------G---Y---------AT----V---------------------------------------------------------------------------D------A---A-----KR-----P--E--------A---------------------------------------------------------------------------------------------R---------------------------------------------FT---GPENITSKAHV---------------------L---L---------ASK-N----------SV---------N----------------------------------------D--RIL--S-----------STY------------------HSVD-RSHR-----I-A-HNQEILI----RL-----V-----E--------F--------------AR-ITR------------V---------E---G-----W-A----------------------G--T-D--SV-L--KVLDGGLGHNFIKLGYNT-EEH--KSFSY-R-------VDV-Y---------------G--------------------------------------------------------------------------------------------------------------------------------------------------------------V----------------------------------------------------------------------------------------------------------LECDD--------------------------------------------------

>Slitu-gammaREPAT14

---------MK-----V--L----------A---V-V------L--L-------------V---A--V-----------A---A---------A-----V---------------------------------------------------------------------------N------S---Y-----EA-----Q--E--------A-------------------NVF-YEE-LD---M--------------------------------EESDP----------------------------------------------------------------------SVAR---------RL---------------------H---Y---------QWG-S----------VGP--------D----------------------------------------D--RLI--S-----------ATY------------------HEAY-GMEY-----Q-I-NDQEVTY----RG-----N-----E--------K--------------TN-ITS------------I---------I---V-----R-E-----------------Y--L-S--NYT---T-V--TILDGGLYRNHVTLGLKG-LRS--LQIGC-H-------ISI-Y---------------A--------------------------------------------------------------------------------------------------------------------------------------------------------------T----------------------------------------------------------------------------------------------------------VNC--SNM-----------------------------------------------

>Slitu-gammaREPAT15

---------MK-----I--L----------A-----V------L--L----ML-------V---A--I-----------A---A---------G-----V---------------------------------------------------------------------------N------S---N-----LE-----Q--E--------A-------------------NVLNYNE-QDSNKM--------------------------------VESDL-----------------------M---------------------------------------------AS------------RQ---------------------I---V----------WG-V----------DGP--------D----------------------------------------D--RLI--S-----------STI------------------HSAD-AFNF-----L-I-NDQEVTY----RG-----H-----E--------T--------------TN-ITR------------I---------V---V-----L-K-----------------R--V-G--DIT---Q-V--DLVDGGLYHNFVTIRFTC-QRS--YSMYN-I-------IDI-F---------------A--------------------------------------------------------------------------------------------------------------------------------------------------------------T----------------------------------------------------------------------------------------------------------INC--HNN-----------------------------------------------

>Slitu-gammaREPAT16

---------MK-----GF-L----------L---I-I------L--L-------A-----V---T--Y-----------G---Y---------AT----V---------------------------------------------------------------------------D------G---E-----DL-----P--E--------A---------------------------------------------------------------------------------------------P---------------------------------------------LG---GPE-GISTLSG---------------------S---L---------SLG-D----------T----------N----------------------------------------G--RML--S-----------STS------------------HTVG-SSIN-----I-I-HTQEVLV----RL-----V-----E--------Y--------------AN-ITG------------I---------R---V-----T-P----------------------T--NNM--NAVP--SLINGGLGHNYVNLRFTS-QLS--LGYGY-R-------IMV-Y---------------G--------------------------------------------------------------------------------------------------------------------------------------------------------------S----------------------------------------------------------------------------------------------------------FVC----------------------------------------------------

>Slitu-gammaREPAT1

---------MK-----SL-I----------L---V-A------V--L-------A---ALT---V--C-----------N---D---------A---------------------------------------------------------------------------------------------------A-----T-LQ--------N---------------------------------------------------------------------------------------------A---------------------------------------------AF------------RA---------------------N---L---------YQG-G----------IRP--------G----------------------------------------D--RLL--H-----------SNY------------------YYKN-PIAN-----A-V-QYQDITY----RG-----N-----S--------S--------------TR-ISY------------I---------Q---V-----T-E-----------------V--G-Y--TQW--GI-P--SLRSGGVNYSYATIRLTS-QRG--YGYYY-R-------VEI-W---------------G--------------------------------------------------------------------------------------------------------------------------------------------------------------R-----------------------------------------------------------------------------------------------------------------------------------------------------------------

>Slitu-gammaREPAT2

---------MK-----SL-I----------L---I-A------V--L-------A---ALT---V--C-----------N---D---------A---------------------------------------------------------------------------------------------------A-----A-LQ--------N---------------------------------------------------------------------------------------------P---------------------------------------------AY------------RA---------------------N---L---------YQG-A----------IRP--------G----------------------------------------D--RLL--H-----------SNY------------------YYKN-PIAN-----A-V-QYQDITY----RG-----N-----S--------S--------------TR-ISY------------I---------Q---V-----T-E-----------------V--G-Y--TQW--GI-P--SLRSGGVNFNHATIRLTS-QRG--YGYYY-R-------VEI-W---------------G--------------------------------------------------------------------------------------------------------------------------------------------------------------R-----------------------------------------------------------------------------------------------------------------------------------------------------------------

>Slitu-gammaREPAT3

---------MK-----SL-I----------L---V-A------V--L-------A---ALT---V--C-----------N---D---------A---------------------------------------------------------------------------------------------------A-----T-LQ--------N---------------------------------------------------------------------------------------------P---------------------------------------------AY------------RA---------------------N---L---------YQG-A----------IRP--------G----------------------------------------D--RVL--H-----------SNY------------------YIKN-PIPN-----A-V-QYQDITY----RG-----N-----S--------T--------------TR-ISY------------I---------Q---V-----T-E-----------------V--G-Y--TQW--GI-P--SLRSGGVNFNHATIRLTS-QRG--YGYYY-R-------VEI-W---------------G--------------------------------------------------------------------------------------------------------------------------------------------------------------R-----------------------------------------------------------------------------------------------------------------------------------------------------------------

>Slitu-gammaREPAT4

---------MK-----SL-I----------L---V-A------V--L-------A---ALT---V--C-----------N---D---------A---------------------------------------------------------------------------------------------------A-----T-LQ--------N---------------------------------------------------------------------------------------------P---------------------------------------------AY------------RA---------------------N---L---------YQG-A----------IRP--------G----------------------------------------D--RIL--H-----------SNY------------------YIKN-PIAN-----A-V-QYQDITY----RG-----N-----S--------S--------------TR-ISY------------I---------Q---V-----T-E-----------------V--G-Y--TQW--GI-P--SLRSGGVNFNHATIRLTS-QRG--YGYYY-R-------VEI-W---------------G--------------------------------------------------------------------------------------------------------------------------------------------------------------R-----------------------------------------------------------------------------------------------------------------------------------------------------------------

>Slitu-gammaREPAT7

---------MK-----SL-F----------L---I-V------L--L-------V-----V---A--A-----------S---T---------A---------------------------------------------------------------------------------------------------VINQETA--A--------D---------------------------------------------------------------------------------------------PHLVETFEEFDLGDDPDISYEPD------VADVTDV-PI----GEV-T------------SP---------------------S---IDAATCC---SIG-A-----------RS--------G-------------------------------------------KLL--A-----------RSN------------------HVRR-GKPN-----V-T-YVQNVTF----RT-----G-----S--------H---------------L-ITA------------I---------H---I-----T-H-----------------I--G-P--MQG--SR-A--SIIAGGLNRNFVTIKLQS-ARG--RGYNF-R-------IRI-Y---------------G--------------------------------------------------------------------------------------------------------------------------------------------------------------R-----------------------------------------------------------------------------------------------------------------------------------------------------------------

>Slitu-gammaREPAT8

---------MK-----VL-L----------L---L-V------L--L-------A-----A---A--A-----------S---A---------SILDYEV---------------------------------------------------------------------------T------D---E-----DN-----Q--E--------E----------------------------------------------------------------------DTVSPDLDDTPDHYEEDVMLRISP---------------------------------------------TP------------TATDIDEWAELFRMAVEEGMIHRN---L---------SLG-N----------IGA--------R----------------------------------------D--ILL--S-----------KTH------------------HFKS-DGDT-----N-V--SEDIYF----RC-----C-----P--------T--------------TIIITA------------I---------R---G-----G-S-----------------V-------SEN--VR-P--SVVSGGVGHNHVCIRLHS-QKG--HGYRY-C-------IEI-Y---------------G--------------------------------------------------------------------------------------------------------------------------------------------------------------R---------------------------------------------------------------------------------------------------------K-------------------------------------------------------

>Slitu-gammaREPAT9

---------MK-----SL-L----------L---V-V------L--L-------A-----A---A--A-----------S---A---------A---------------------------------------------------------------------------------------------------IVSQDIA--E--------A---------------------------------------------------------------------------------------------PEAVVTYESIMYSDD---------------AEVAEI-EAASRSVEIET------------RR---------------------N---L---------SVG-T----------IGN--------S-------------------------------------------RLL--A-----------NTQ------------------HIRG-AIAN-----T-I-LVHNATF----NF-----A-----S--------S--------------IR-IAA------------I---------R---V-----T-R-----------------V--G-A--SQN--AT-P--SIVSGGLNRNFVTIRITS-ARG--RGFNY-R-------IQI-Y---------------G--------------------------------------------------------------------------------------------------------------------------------------------------------------R-----------------------------------------------------------------------------------------------------------------------------------------------------------------

>Slitu-gammaREPAT10

---------MK-----SL-L----------V---V-V------L--L-------A-----A---A--A-----------S---A---------A---------------------------------------------------------------------------------------------------IVSQDIA--E--------A---------------------------------------------------------------------------------------------PEAVVTYESVMYSDD---------------AEVAEV-EAGSKSVDIET------------RR---------------------N---L---------SVG-T----------IGN--------S-------------------------------------------RLL--A-----------NTQ------------------HIRG-AIAN-----T-I-LVHNATF----NF-----A-----S--------S--------------IR-IAA------------I---------R---V-----T-R-----------------V--G-A--SQN--AT-P--SIVSGGLNRNFVTIRITS-ARG--RGFNY-R-------IQI-Y---------------G--------------------------------------------------------------------------------------------------------------------------------------------------------------R-----------------------------------------------------------------------------------------------------------------------------------------------------------------

>Slitu-gammaREPAT11

---------MK-----SL-L----------V---V-V------L--L-------A-----A---A--A-----------S---A---------A---------------------------------------------------------------------------------------------------IVSQDIA--E--------A---------------------------------------------------------------------------------------------PEAVVTYESIMYSDD---------------AEVAEI-EAASRSVEIES------------RR---------------------N---L---------SVG-T----------IGN--------S-------------------------------------------RLL--A-----------NTQ------------------HIRG-AIAN-----T-I-LVHNATF----NF-----A-----S--------S--------------IR-IAA------------I---------R---V-----T-R-----------------V--G-A--SQN--AT-P--SIVSGGLNRNFVTIRITS-ARG--RGFNY-R-------IQI-Y---------------G--------------------------------------------------------------------------------------------------------------------------------------------------------------R-----------------------------------------------------------------------------------------------------------------------------------------------------------------

>Slitu-gammaREPAT12

---------MK-----SL-L----------V---V-V------L--L-------A-----A---A--A-----------S---A---------A---------------------------------------------------------------------------------------------------IVSQDIA--E--------A---------------------------------------------------------------------------------------------PEAVITYESIMYSDD---------------AEVAEI-EAGSKSVEIES------------RR---------------------N---L---------SVG-T----------IGN--------S-------------------------------------------RLL--A-----------NTQ------------------HIRG-AIAN-----T-I-LVHNATF----NF-----A-----S--------S--------------IR-IAA------------I---------R---V-----T-R-----------------V--G-A--SQN--AT-P--SIVSGGLNRNFVTIRITS-ARG--RGFNY-R-------IQI-Y---------------G--------------------------------------------------------------------------------------------------------------------------------------------------------------R-----------------------------------------------------------------------------------------------------------------------------------------------------------------

>Tuta-betaREPAT11

----MLSK--------------------------V-V------I--L-------L-----A---L--V-----------A---I---------------V---------------------------------------------------------------------------Y------G---K-----S-------------------------------------------------------------------------------------------------------------------------------------------------------------------------------------------------------V---------IVG-Q----------IFDHSGQ----V----------------------------------------R--KVY--D-----------KMA------------------E-KN-AVPL-----L-K-REEEIQF----EY-----P-----L--------G-D------------QR-IKG------------I---------A---I-----K-D-----------------L--D-N--G-L--AE-P--SIVRGGLGFNFVTIKLKS-ERG--SGYRF-L-------IEI-Y---------------A--------------------------------------------------------------------------------------------------------------------------------------------------------------------------------------------------------------------------------------------------------------------------------------------------------------------------------

>Tuta-betaREPAT12

---------MR------A-I-------TL-L---T-F------F--C-------I-----L---A--A-----------A---S---------------C---------------------------------------------------------------------------Q------S---H-----N-------------------------------------------------------------------------------------------------------------------------------------------------------------------------------------------------------F---------YIG-T----------ASY--------G----------------------------------------D--VVL--Y-----------KSV------------------EVKY-AFPF-----F-V-RTTDVKF-----P-ELGKA-----S--------Y--------------AI-ITA------------I---------Y---V-----H-DNY---------------I--D-G--S-G--GY-A--TIRAGGVGQRFVNVHLKS-QRG--GGFNF-T-------VTI-Y---------------G--------------------------------------------------------------------------------------------------------------------------------------------------------------R-----------------------------------------------------------------------------------------------YY----------------------------------------------------------------

>Tuta-betaREPAT13

---------MY-------------------------L------R--L-------V-----LTIAL--V-----------A---S---------------V---------------------------------------------------------------------------T------C---K-----D-------------------------------------------------------------------------------------------------------------------------------------------------------------------------------------------------------F---------EQG-Q---------NATE--------A----------------------------------------R--KIF--D-----------EKR------------------E----ASPA-----VWR-QVDKVTV----NT-----T-----D--------T--------------EV-ISR------------V---------V---V-----T-D-----------------LRPD-K----D--GE-A--SIVDGGEGNQTVTIELKS-PTV-LRGYTF-Q-------IEV-Y---------------A----------------------------------------------------------------------------------------------------------------------------------------------------------------------V--------P-----DYGHV-----STPAQVG----------------DD------------------------------------KP------------------------------------------------------------------

>Tuta-betaREPAT8

---------MF------K-I----------T---S-L------L--L-------V-----L---Y--V-----------S---S---------------S---------------------------------------------------------------------------Q------A---N-----D-------------------------------------------------------------------------------------------------------------------------------------------------------------------------------------------------------I---------RLG-H---------ASPS--------S----------------------------------------R--KIF--S-----------ELR------------------E----ASPA-----IWT-RNDDVIV----TT-----V-----G--------T--------------EV-ISA------------I---------Y---V-----T-D-----------------LRED-K----T--GE-A--YIESGGIGMKTVTIGLRS-PSI-LRGYKF-Q-------IEV-Y---------------A------D-------------------------------------------------------------------------------------------------------------------------------------------------------D----S-TA--------R-----YYGHA-KG--GYPYPAGGLSGAYGVKGGATTYYDD-GQFA-----------------------R-------KY------------------------------------------------------------------

>Tuta-betaREPAT1

---------MK------S-L----------F---L-F------L--V-------V-----A------V-----------Y---C---------------I---------------------------------------------------------------------------S------L---N--KADD-------------------------------------------------------------------------------------------------------------------------------------------------------------------------------------------------------L---------QMG-N----------GVG--------S----------------------------------------T--LAY--L-----------ENI------------------K-LS-GIPF-----K-T-RTKNVFF----SS-----D-----K--------N--------------QI-IKA------------I---------S---A-----I-D-----------------Q--K-R--S-G--AK-A--TVTAGGVGSTFVNIKLKS-ERS--EGLDY-Q-------VQI-Y---------------A--------------------------------------------------------------------------------------------------------------------------------------------------------------------------------------------------------------------------------------------------------------------------------------------------------------------------------

>Tuta-betaREPAT15

----MVKIVSILA----L-V----------F---L-A------A--C-------L--------------------------------------------V---------------------------------------------------------------------------T------A---D-----H-------------------------------------------------------------------------------------------------------------------------------------------------------------------------------------------------------L---------VIGNV----------ANR--------V----------------------------------------Q--LAK--H-----------ELV------------------E-YN-AFPF-----M-K-RVKMYFY----SG-----P-----D--------Y--------------KK-IQG------------I---------Q---A-----L-D-----------------T--L-H--S-K--AS-I--NITAGGIGHSFVNMRMKS-ERG--RGLSY-D-------IGI-Y-VNPDF-Q----------------------------------------------------------------------------------------------------------------------------------------------------------------------------------------------------------------------------------------------------------------------------------------------------------------------------------------

>Tuta-betaREPAT16

---------MK------I-I---VYLC---I---L-L------S--A-------L-----T---C--V---------------M---------------C---------------------------------------------------------------------------K------TYFYG-----Q---------------------------------------------------------------------------------------------------------------------------------------------------------------------------------------------------------------------K----------TDE--------T----------------------------------------R--LIN--R-----------EKL------------------V-YF-AVPW-----R-R-RIEYYAY----ND-----T-----L--------Q--------------RP-IKA------------I---------D---C-----H-D-----------------Y--Q-R--T-D--AT-V--NITEGGVGYPYVSLLFQS-QRS--YGLEY-A-------IEI-F---------------A--------------------------------------------------------------------------------------------------------------------------------------------------------------------------------------------------------------------------------------------------------------------------------------------------------------------------------

>Tuta-betaREPAT17

---------MD------A-L----------V---L-V------L--L-------L-----S---A--L-----------A---A-AVRGTRLF------C---------------------------------------------------------------------------D------R---D-----Q-------------------------------------------------------------------------------------------------------------------------------------------------------------------------------------------------------L---------PL-------------------------------------------------------------------------------------EEG------------------EWSR-DYAH-----L-D--TATIEN-----P-----P-----V--------T--------------VL-EGQ------------I---------Y---V-----Y-ENFFPNRTIKYMHVDNLAI--K-S--C-G--AS-A--AIKAGGIGTSGVMVVLHA-GPH--QEIRS-V-------IDI-W---------------G-------TRL--------TP--R---RPQHRRAML--VPPEDLEQMKKLRSKYLAVS-RTVNHNKYNDTDF----D----------------------------------------------------------------------------------D-----------------------------------------------------------------------------------------------------------------------------------------------------------------

>Tuta-betaREPAT2

---------MK-S-I--V-I----------L---S-F------V--V-------L-----I---C--V-----------S---S---------------S---------------------------------------------------------------------------L------A---K-----D-------------------------------------------------------------------------------------------------------------------------------------------------------------------------------------------------------L---------IVG-T----------SFN--------N----------------------------------------R--LLW--Q-----------QKA------------------E-YN-AIPF-----K-K-RVKEVFF----SD-----P-----G--------Q--------------QI-IKG------------I---------I---A-----R-D-----------------L--D-H--T-E--AS-A--TVTAGGVGFSYANIRLKS-DRG--SGLNY-Q-------IEI-Y---------------T--------------------------------------------------------------------------------------------------------------------------------------------------------------------------------------------------------------------------------------------------------------------------------------------------------------------------------

>Tuta-betaREPAT3

---------MK-T-V--P-V----------L---C-I------V--L-------L-----A---L--V-----------G---D---------------S---------------------------------------------------------------------------Y------C---K-----D-------------------------------------------------------------------------------------------------------------------------------------------------------------------------------------------------------L---------IVG-T----------SFN--------N----------------------------------------R--LLW--Q-----------QKA------------------E-YN-AIPL-----K-K-RVKEVFF----SD-----P-----G--------Q--------------QI-IKG------------I---------I---A-----R-D-----------------L--D-H--T-E--AE-A--TVTAGGVGFSYANIRLKS-ERG--SGLNY-Q-------IEI-Y---------------S--------------------------------------------------------------------------------------------------------------------------------------------------------------------------------------------------------------------------------------------------------------------------------------------------------------------------------

>Tuta-betaREPAT4

---------MR------Q-V----------L---L-I------L--V-------L-----S---V--V-----------V---Y---------------V---------------------------------------------------------------------------C------A---Y-----D-------------------------------------------------------------------------------------------------------------------------------------------------------------------------------------------------------M---------IIG-D----------TVH--------R----------------------------------------K--MVF--H-----------QRV------------------K-DF-AIPF-----K-K-RIKTLTY----MD-----P-----E--------K--------------RV-IKG------------V---------A---A-----V-D-----------------N--D-F--S-L--AS-A--NITDGGVGSSHVTVRMKS-QRH--HPLNF-E-------IEI-Y---------------V--------------------------------------------------------------------------------------------------------------------------------------------------------------------------------------------------------------------------------------------------------------------------------------------------------------------------------

>Tuta-betaREPAT5a

---------MK------F-E----------I---L-I------L--L-------L-----G---C--V---------------L---------------A---------------------------------------------------------------------------N------C---G-----H-------------------------------------------------------------------------------------------------------------------------------------------------------------------------------------------------------M---------KIG-E----------IEG--------R----------------------------------------T--LMH--E-----------TVA------------------Q-YP-GVEN-----R-R-RLKYVVY----TD-----L-----Q--------M--------------RT-IRG------------I---------D---A-----H-D-----------------N--M-E--S-G--AM-A--NILAGGIGYPFVTMKLRS-ERS--KGLDY-A-------IKI-FA--------------*--------------------------------------------------------------------------------------------------------------------------------------------------------------------------------------------------------------------------------------------------------------------------------------------------------------------------------

>Tuta-betaREPAT5b

---------MK-------------------T---S-V------V--L-------A-----A------I-----------L---T---------------I---------------------------------------------------------------------------A------S---V--KSEN-------------------------------------------------------------------------------------------------------------------------------------------------------------------------------------------------------L---------TLG-E----------THC--------N----------------------------------------H--VIY--L-----------QKI------------------K-FV-GIPF-----V-K-RDTDIFF----NS-----K-----Y--------N--------------EK-ISR------------I---------E---A-----I-E-----------------K--H-P--T-K--AT-A--IITAGGLGQGYVNIHLKS-ERG--SGLYY-K-------ITI-Y---------------S--------------------------------------------------------------------------------------------------------------------------------------------------------------------------------------------------------------------------------------------------------------------------------------------------------------------------------

>Tuta-betaREPAT6

---------MK------L-V----------I---S-V------T--L-------L-----V---L--I-----------L---C---------------V---------------------------------------------------------------------------S------I---Y--EAKD-------------------------------------------------------------------------------------------------------------------------------------------------------------------------------------------------------L---------LVG-S----------REN--------K----------------------------------------R--LIS--T-----------QKV------------------V-NK-GTPL-----L-I-KEQDYTY----TD-----P-----K--------Q--------------RI-IKA------------I---------I---A-----R-D-----------------L--S-R--T-D--TI-A--EVKSGGEGASHVTIHFKS-GRG--QNINY-L-------ILL-F---------------A-----E--------------------------------------------------------------------------------------------------------------------------------------------------------------------------------------------------------------------------------------------------------------------------------------------------------------------------

>Tuta-betaREPAT7a

---------MK------L-F----------I---L-F------L--V-------T-----F---V--M-----------V---M---------------V---------------------------------------------------------------------------E------S---L-----H-------------------------------------------------------------------------------------------------------------------------------------------------------------------------------------------------------M---------FAG-S----------NVM--------R----------------------------------------N--LVM--H-----------KEL------------------K-YG-ARKS-----Q-N-RVEFYNY----TL-----PFDNLQR--------D--------------QY-IAG------------L---------L---V-----F-D-----------------K--M-N--S-G--AT-V--EVVSGGLGYNYLQLRMMS-EKG--KPLKY-D-------INV-Y---------------T--------------------------------------------------------------------------------------------------------------------------------------------------------------------------------------------------------------------------------------------------------------------------------------------------------------------------------

>Tuta-betaREPAT7b

---------MK------V-V----------I---L-A------L--I-------V-----G---L--V-----------A---V---------------A---------------------------------------------------------------------------V------Y---GD----H-------------------------------------------------------------------------------------------------------------------------------------------------------------------------------------------------------T---------FMG-T----------NMG--------R----------------------------------------R--LAH--N-----------AKV------------------R-YS-SKLM-----Q-K-RVEYWNY----NF-----P-----A--------G--------------TQ-IAG------------I---------L---A-----Y-D-----------------E--L-K--S-G--AI-A--NVTAGGIGYNFVQLQMKS-SRG--KAIRY-N-------VYL-Y---------------H--------------------------------------------------------------------------------------------------------------------------------------------------------------------------------------------------------------------------------------------------------------------------------------------------------------------------------

>Tuta-betaREPAT7c

---------MK------V-L----------L---S-A------L--V-------L-----C---L--V-----------A---V---------------T---------------------------------------------------------------------------V------Y---S-----E-------------------------------------------------------------------------------------------------------------------------------------------------------------------------------------------------------T------GFLFDG-N----------SVY--------T----------------------------------------K--LVR--K-----------EKV------------------R-FT-SD-----------KVEEWHY----VS-----P-----G--------A--------------KS-IQG------------I---------L---A-----Y-D-----------------E--T-N--S-R--AQ-V--NVTNGGTGYGYVTLAFKA-GQGQ-KKIYF-D-------VFI-Y----------TL---A--------------------------------------------------------------------------------------------------------------------------------------------------------------------------------------------------------------------------------------------------------------------------------------------------------------------------------

>Tuta-betaREPAT7d

---------MK------L------------T---L-A------F--F-------V-----V---I--L-----------A---V---------------V---------------------------------------------------------------------------N------C---K-----H-------------------------------------------------------------------------------------------------------------------------------------------------------------------------------------------------------T---------FMG-T----------NVQ--------R----------------------------------------P--LLE--H-----------KVI------------------K-YS-ANII-----T-K-RIEKFHY----TV-----P-----Q-----KHFG--------------QV-IQG------------I---------L---A-----Y-D-----------------Q--T-N--T-G--AS-A--NVTNGGLGANTVTLRMKS-DRG--KPIHF-D-------VYI-Y---------------A--------------------------------------------------------------------------------------------------------------------------------------------------------------------------------------------------------------------------------------------------------------------------------------------------------------------------------

>Tuta-gammaREPAT1

---------MK-----AI-L----------V---L-CM--------L-------A---ALA---L--------------T---E---------A---------------------------------------------------------------------------------------------------S-----VA-R--------A---------------------------------------------------------------------------------------------P---------------------------------------------AT------------RS---------------------N---L---------SLG-Y----------IQY--------G----------------------------------------D--RLL--A-----------RQY------------------LAKG-ALPN-----A-I-QYQDLIY----RG-----N-----A--------T--------------TR-ISA------------I---------Q---A-----M-E-----------------V--G-Y--TQW--AS-A--WVVAGGIGWNSATVRVQS-ARG--YGYNY-Q-------IDI-W---------------G--------------------------------------------------------------------------------------------------------------------------------------------------------------R-----------------------------------------------------------------------------------------------------------------------------------------------------------------

>Tuta-gammaREPAT2

---------MK-----TT-L----------L---L-C------T--L-------A---VLA---I--------------T---Q---------A---------------------------------------------------------------------------------------------------S-----V--I--------K---------------------------------------------------------------------------------------------Q---------------------------------------------A-------------TN---------------------D---L---------ILG-A----------IQP--------G----------------------------------------D--RLLY-K-----------QIQ------------------MYQE-AIPM-----Q-S-HIHDLIF----IG-----N-----E--------T--------------SV-IHA------------I---------H---A-----I-E-----------------E--G-E--FQY--AS-I--SLRSGGLGTNKAVLRARA-RQG--FGYNY-W-------VSI-Y---------------G--------------------------------------------------------------------------------------------------------------------------------------------------------------T-------------------------------------------------------------------------------------------------------D---------------------------------------------------------

>Tuta-gammaREPAT3

---------MK-----TT-L----------L---L-C------T--L-------A---VLA---I--------------T---Q---------A---------------------------------------------------------------------------------------------------S-----V--I--------K---------------------------------------------------------------------------------------------Q---------------------------------------------A-------------TN---------------------D---L---------ILG-A----------IQP--------G----------------------------------------D--RLL--R-----------RLQ------------------MYQE-AIPM-----Q-S-HIHDLIF----IG-----N-----E--------T--------------SV-IHA------------I---------H---A-----I-E-----------------E--G-E--FQY--AS-M--SLRSGGLGTNKAVLRARA-RQG--FGYNY-W-------VSI-Y---------------G--------------------------------------------------------------------------------------------------------------------------------------------------------------T-------------------------------------------------------------------------------------------------------D---------------------------------------------------------

>Tuta-gammaREPAT4

---------MK-----TT-L----------L---L-C------T--L-------A---VLA---I--------------T---Q---------A---------------------------------------------------------------------------------------------------S-----V--I--------K---------------------------------------------------------------------------------------------Q---------------------------------------------A-------------RN---------------------D---L---------ILG-A----------VQP--------G----------------------------------------D--RLL--R-----------RQV------------------MYQE-AIPM-----Q-S-HIHDLIF----IG-----N-----E--------T--------------NV-IHA------------I---------Q---A-----I-E-----------------E--G-E--PQF--GS-M--TLRAGGLGTNLAVLRAHS-RQG--FGYNY-W-------VSI-Y---------------G--------------------------------------------------------------------------------------------------------------------------------------------------------------T-------------------------------------------------------------------------------------------------------D---------------------------------------------------------

>Tuta-gammaREPAT5

---------MK-----TA-L----------I---L-C------T--L-------A---ALA---I--------------S---Q---------A---------------------------------------------------------------------------------------------------S-----V--V--------K---------------------------------------------------------------------------------------------Q---------------------------------------------A-------------RA---------------------N---L---------ILG-S----------VRP--------G----------------------------------------D--RLL--Q-----------SAV------------------LYQP-AIAN-----R-V-QTQDLIF----RG-----N-----V--------S--------------SR-ISS------------I---------Q---A-----R-E-----------------V--G-A--SQY--AQ-V--RLISGGLNRSNATLRATS-RVS--YGFNY-Q-------IQI-W---------------G--------------------------------------------------------------------------------------------------------------------------------------------------------------R-----------------------------------------------------------------------------------------------------------------------------------------------------------------
